# Recurrent mutations in topoisomerase 2a cause a novel mutator phenotype in human cancers

**DOI:** 10.1101/2020.05.22.111666

**Authors:** Arnoud Boot, Steven G. Rozen

## Abstract

Topoisomerases are essential for genome stability. Here, we link the p.K743N mutation in topoisomerase TOP2A to a previously undescribed mutator phenotype in human cancers. This phenotype primarily generates a distinctive pattern of duplications of 2 to 4 base pairs and deletions of 6 to 8 base pairs, which we call ID_TOP2A. All tumors carrying the TOP2A p.K743N mutation showed ID_TOP2A, which was absent in all of 12,269 other tumors. We also report evidence of structural variation associated with TOP2A p.K743N. All tumors with ID_TOP2A mutagenesis had several indels in known cancer genes, including frameshift mutations in *PTEN* and *TP53* and an in-frame activating mutation in *BRAF*. Thus, ID_TOP2A mutagenesis almost certainly contributed to tumorigenesis in these tumors. This is the first report of topoisomerase-associated mutagenesis in human cancers, and sheds further light on TOP2A’s role in genome maintenance. We also postulate that tumors showing ID_TOP2A mutagenesis might be especially sensitive to topoisomerase inhibitors.

## Main text

Type II topoisomerases are essential for genome maintenance. They untangle DNA by creating and then repairing double strand breaks to allow entangled DNA strands to pass through each other and thus untangle [1-3]. During this process, the 5’ ends of the breaks are bound to the topoisomerase with a phosphotyrosyl bond. After the breaks have been repaired, the phosphotyrosyl bond is cleaved, releasing the topoisomerase-DNA complex [4]. Humans have 2 genes encoding type II topoisomerases: *TOP2A* on chromosome 17 and *TOP2B* on chromosome 3 (reference [5]). TOP2A and TOP2B function similarly, but TOP2A is mainly expressed in proliferating cells, whereas TOP2B is also expressed in quiescent cells [6]. Topoisomerases have been widely studied as therapeutic targets, with several topoisomerase inhibitors, such as etoposide and doxorubicin, being commonly used chemotherapeutics [5, 7]. Additionally, upregulation of topoisomerases in cancer has been shown to be a marker of poor prognosis [8, 9]. However, to date, there have been no reports of somatically altered type II topoisomerases playing a role in human tumorigenesis.

To study whether defects in TOP2A or TOP2B contribute to carcinogenesis, we looked for hotspot mutations in 23,829 tumors [10]. TOP2B showed a clear hotspot at p.R651, and TOP2A showed several hotspots in various protein domains (Fig. 1A, 1B). TOP2B p.R651 and most recurrent TOP2A mutations were in hypermutated tumors (Fig. 1C, Supplementary Table S1). TOP2A p.K743N on the other hand, was observed in 4 gastric cancers (GCs) and 1 cholangiocarcinoma (CCA_TH_19), none of which were hypermutated. Strikingly, all GCs carrying TOP2A p.K743N showed elevated levels of small insertions and deletions (indels). The indel spectra of the 4 GCs were partly composed of a distinct pattern that was previously identified as indel mutational signature ID17, which is characterized by duplications of 2 to 4 base pairs (bp) in non-repetitive sequences [10] (Fig. 1D). Among 12,273 tumors with indels analyzed by Alexandrov et al., TOP2A p.K743N and ID17 occurred only in these 4 GCs (p = 1.06×10^−15^, two-sided Fisher’s exact test). Alexandrov et al. did not analyze indels in CCA_TH_19. Furthermore, the COSMIC database of cancer mutations contained 2 additional tumors carrying TOP2A p.K743N, and WES was available for these [11, 12]. Re-alignment and indel calling in CCH_TH_19 and the 2 tumors in COSMIC revealed ID17 in all three (Fig. 1D).

**Fig. 1:**
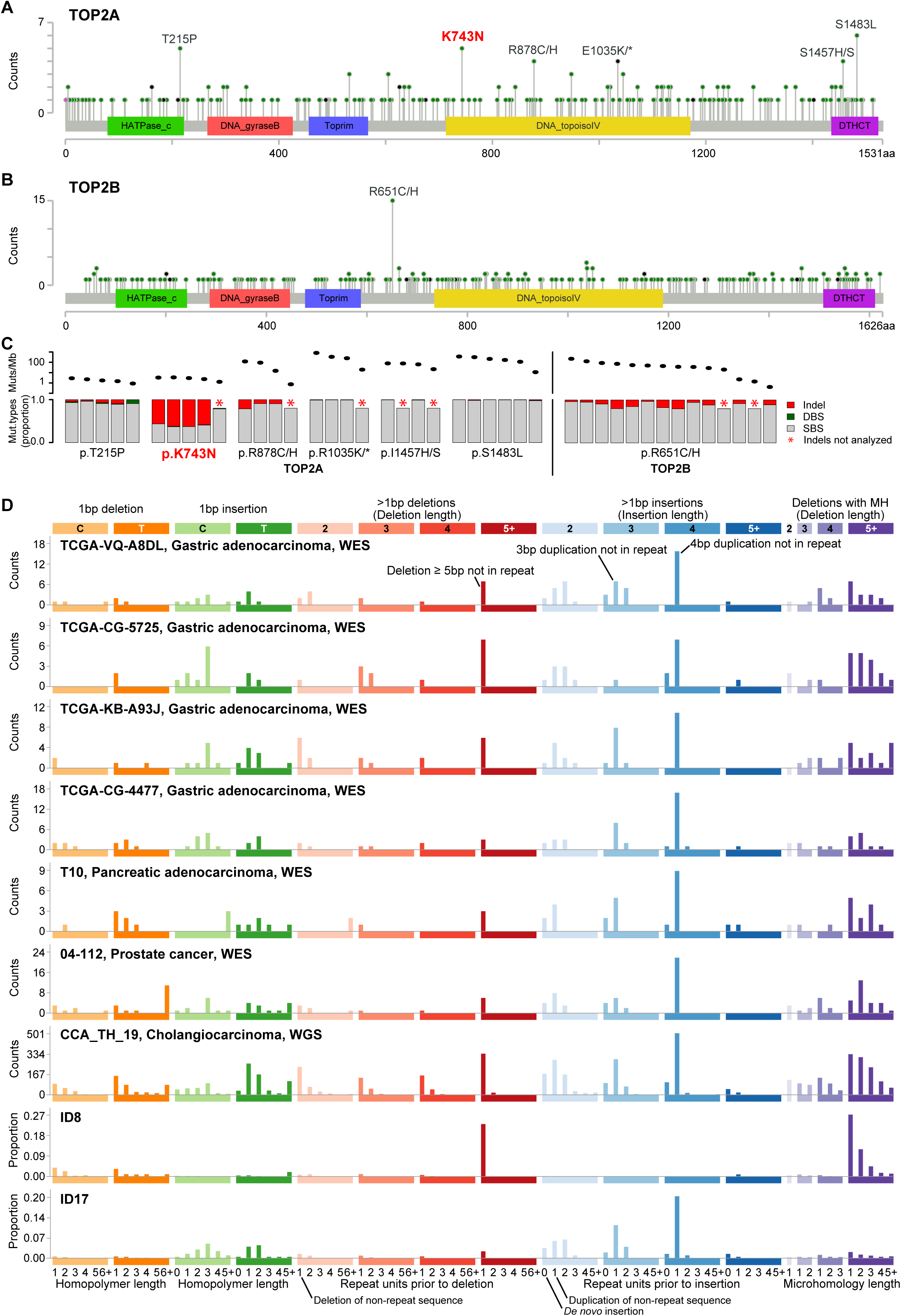
TOP2A and TOP2B hotspot mutations and associated mutagenesis. Recurrent amino acid substitutions in TOP2A (**A**) and TOP2B (**B**) (figure generated using MutationMapper [21]). **C**, Mutagenesis profiles of TOP2A and TOP2B hotspot carriers as reported by Alexandrov et al. Ovals in top sub-panel show combined mutation load of indels, DBSs and SBSs; bars in bottom panel show the proportions of indels (red), DBSs (green) and SBSs (gray). **D**, Indel mutation spectra of tumors carrying the TOP2A p.K743N substitution, and the ID8 and ID17 indel mutational signatures [10]. Abbreviations: bp: base pair, DBS: Doublet base substitution, indels: small insertions and deletions, MH: microhomology, Mut: mutation, SBS: Single base substitution, WES: Whole exome sequencing, WGS: whole-genome sequencing.

Hypothesizing that ID17 is related to DNA cleavage by TOP2A, and because TOP2A cleavage is enriched in highly transcribed regions [13], we examined ID17 mutagenesis in relation to transcriptional activity. Consistent with this, indel mutagenesis in CCA_TH_19 increased with transcriptional activity (p = 6.13×10^−53^, one-sided Cochran-Armitage test, Supplementary Fig. S1). Notably, in addition to duplications of 2 to 4 bp not in repetitive sequences, all other classes of indels were also positively correlated with transcriptional activity, suggesting that they also stemmed from TOP2A p.K743N mutagenesis (FDR < 0.05 for all classes, one-sided Cochran-Armitage test, Fig. 2A). As with CCA_TH_19, all whole-exome sequenced samples also showed higher density of indel mutagenesis in more highly transcribed regions (p = 0.016, one-sided sign-test, Fig. 2B).

**Fig. 2:**
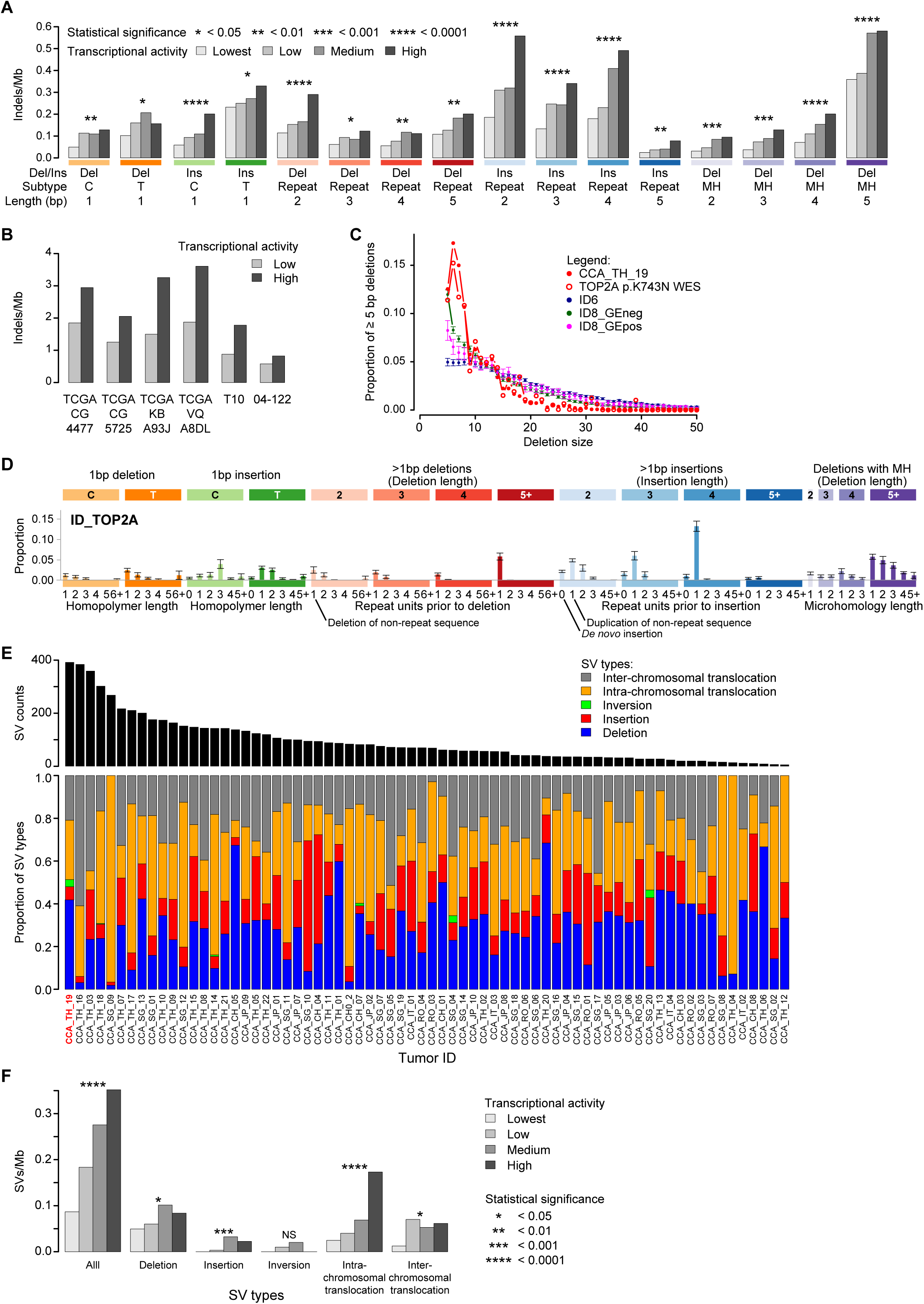
Characterization of ID_TOP2A. **A**, All types of indels show increased density of mutagenesis with increased transcriptional activity in CCA_TH_19. **B**, All WES samples also show higher indel density in highly transcribed regions (p = 0.016, one-sided sign-test). Because of low indel counts in WES data, mutations were split into low (light gray) and high (dark gray) expression bins. **C**, Comparison of size distributions of ≥ 5 bp deletions in TOP2A p.K743N carriers, in tumors with high counts of ID6 deletions, in tumors with high counts of ID8 deletions with increasing mutation density in highly transcribed genes (ID8_GEpos), and in tumors with high counts of ID8 deletions without this characteristic (ID8_GEneg). Please see Online Methods for details. Compared to each of the other groups of tumors, TOP2A p.K743N carriers were enriched for 6 to 8 bp deletions (p < 0.0001 for each of the pairwise comparisons, two-sided Wilcoxon rank sum tests). For ID6, ID8_GEneg and ID8_GEpos, means and 95% confidence intervals are plotted. For display purposes, all TOP2A p.K743N WES data was combined, statistical comparisons between the groups was performed using the individual data points. **D**, The ID_TOP2A mutational signature. Consensus indel signature observed in tumors carrying TOP2A p.K743N. Each bar shows the mean proportion of indels in a given indel class; error bars show standard error of the mean. **E**, Total numbers and composition of SVs in 70 CCAs [16]. **F**, In CCA_TH_19, all types of SVs except inversions correlate with transcriptional activity, which is similar to indels in this tumor. *, **, ***, **** indicate Benjamini-Hochberg false discovery rates of < 0.05, < 0.01, < 0.001, and < 0.0001, based on p values from two-sided Cochran-Armitage tests for trend. Abbreviations: bp: base pair, Del: deletion, Ins: insertion, indels: small insertions and deletions, Mb: megabase, MH: microhomology, NS: not significant, SV: structural variation, WES: Whole exome sequencing.

We also observed that the ID17 pattern of duplications in the TOP2A p.K743N tumors always co-occurred with a pattern of ≥ 5 bp deletions that resembled indel mutational signature ID8, which consists almost entirely of ≥ 5 bp deletions. Indeed, in 3 of the indel spectra of the 4 p.K743N GCs, Alexandrov et al., assigned the ≥ 5 bp deletions to ID8 [10]. Furthermore, the numbers of ID17-like mutations and ≥ 5 bp deletions were correlated in the p.K743N tumors. This suggested that the signature of TOP2A p.K743N might in fact be a combination of ID17-like mutations and ≥ 5 bp deletions. This raised two additional questions, because ID8 occurs in the majority of tumors lacking TOP2A p.K743N and occurs across most cancer types.

First, we considered whether ID8 might represent indels caused by TOP2A absent the p.K743N substitution. To investigate this, we compared the ≥ 5 bp deletions in the TOP2A p.K743N carriers to those in all WGS tumors with indels analyzed by Alexandrov et al [10]. These deletions were positively correlated with transcriptional activity in only 124 tumors (Supplementary Table S2). Most of these tumors showed ID8 mutagenesis (82.6%), and 70.2% were liver cancers (Supplementary Table S2). However, in the vast majority of tumors with ID8, transcriptional activity and ID8 density were uncorrelated. Thus, in most tumors, ID8 mutagenesis seems unrelated to TOP2A activity.

Second, we considered whether the grouping together of all ≥ 5 bp deletions as part of categorizing indels might have obscured important differences in deletion size distributions stemming from different mutational processes. To investigate this, we compared the sizes of ≥ 5 bp deletions in TOP2A p.K743N tumors to the sizes of these deletions in tumors with many ID6 or ID8 mutations, as these two signatures consist mainly of ≥ 5 bp deletions. We separately analyzed ID8-high tumors with or without increased ID8 mutagenesis in highly transcribed regions (ID8_GE8pos and ID8_GEneg, respectively, Fig. 2C, Supplementary Table S2). We observed marked enrichment for 6 to 8 bp deletions in TOP2A p.K743N carriers as compared to the other tumors (p = 7.49×10^−6^, 1.94×10^−5^ and 1.97×10^−5^ for ID6, ID8_GEpos and ID8_GEneg respectively, two-sided Wilcoxon rank-sum tests).

To summarize, in addition to the duplications of 2 to 4 bp that were previously reported as ID17, TOP2A p.K743N also induces substantial numbers of 6 to 8 bp deletions, which previous deletion categorizations did not separate from other ID8 deletions [10]. We have shown that these 6 to 8 bp deletions correlate with transcriptional activity, as do all the other major classes of indels observed in TOP2A p.K743N carriers, which is consistent with known mechanistic properties of TOP2A. Thus, we define a novel indel mutational signature incorporating ID17 and features of ID8, which we call ID_TOP2A (Fig. 2D, Supplementary Table S3).

Importantly, every tumor with ID_TOP2A mutagenesis had several insertions and deletions in known cancer driver genes (Supplementary Table S4). Indeed, 6 of the 7 tumors had frameshift mutations in the key tumor suppressor genes *PTEN* and *TP53*. Interestingly, we also observed a 15 bp in-frame deletion in *BRAF*, that has previously been reported to be oncogenic [14]. Thus, ID_TOP2A mutagenesis almost certainly contributed to tumorigenesis in these tumors.

As TOP2A creates and repairs double-strand breaks, and arrested TOP2A cleavage complexes have previously been shown to induce genomic rearrangements [15], we also investigated whether TOP2A p.K743N might induce large-scale, structural variants (SVs). Because SVs are strongly depleted in exons, and WGS was available only for CCA_TH_19, we compared the SVs previously reported for this tumor with the SVs in the other CCAs in the same cohort [16]. CCA_TH_19 had the highest SV-load, consisting mainly of deletions (41.5%) and inter-chromosomal translocations (27.6%, Fig. 2E). Similar to the indels, SVs were also enriched in highly transcribed regions (p = 2.56×10^−12^, one-sided Cochran-Armitage test, Fig. 2F). Enrichment in more highly transcribed regions was also observed for SV deletions, inter-chromosomal translocations, intra-chromosomal translocations and insertions (FDRs 1.93×10^−2^, 4.00×10^−9^, 1.33×10^−2^ and 1.24×10^−3^ respectively, one-sided Cochran-Armitage tests, Fig. 2F). This suggests that the SVs observed in CCA_TH_19, are also associated with TOP2A activity.

Although we have only observed this pattern of indel mutagenesis in human tumors carrying the TOP2A p.K743N substitution, a TOP2 double-mutant (p.F1025Y and p.R1128G) in yeast (*Saccharomyces cerevisiae*) was reported to have a similar mutator phenotype. This suggests that other TOP2A alterations might induce mutator phenotypes similar to ID_TOP2A [17]. Interestingly, this yeast model also showed a strong increase in sensitivity to the topoisomerase II inhibitor etoposide [17]. As we observed in a human TOP2A p.K743N tumor, the yeast double mutants showed strong increases in SVs. This fits with previous observations that trapped TOP2 complexes in yeast result in increased SVs [18].

As more human cancers are sequenced, more somatic TOP2A alterations that induce ID_TOP2A-like mutator phenotypes may be discovered. Human TOP2A p.K743N is located within the DNA cleavage domain of TOP2A, and neither yeast TOP2 p.F1025 nor p.R1128 is located within the orthologous domain. Furthermore, neither of the amino acids altered in TOP2 p.F1025 or p.R1128 is conserved in human TOP2A or TOP2B. Moreover, TOP2 overexpression in yeast also resulted in indels similar to both ID_TOP2A and the indels observed in the p.F1025 and p.R1128 double mutant [17]. Given that such different alterations, as well as overexpression of wild-type TOP2, generated similar indel mutator phenotypes suggests that still other TOP2A alterations might result in similar mutagenesis.

In conclusion, we identified a novel indel mutator phenotype that we propose is induced by TOP2A p.K743N substitutions. This phenotype generates a characteristic pattern of indels that we name ID_TOP2A, which consists of *de novo* duplications of 2 to 4 bp and deletions of 6 to 8 bp. Likely ID_TOP2A mutations in key cancer drivers such as *BRAF, PTEN*, and *TP53*, show that ID_TOP2A almost certainly contributed to tumorigenesis. Data from a yeast model suggest that presence of ID_TOP2A might indicate increased tumor vulnerability to topoisomerase II inhibitors such as etoposide [17].

## Methods

### Data sources

We used published mutation spectra from 23,829 tumors (https://www.synapse.org/#!Synapse:syn11726601) [10]. Variant calls for all 2780 WGS samples from the ICGC/TCGA Pan-Cancer Analysis of Whole Genomes Consortium and gene expression data for a subset of these were obtained from the ICGC data portal (https://dcc.icgc.org/releases/current/Projects/) [19]. Sequencing reads from samples 04-112 and 10T were kindly provided by Professors Peter S. Nelson and Fergus J. Couch [11, 12]. Sequencing reads from CCA_TH_19, for which Alexandrov et al. had not previously analyzed indels, were downloaded from EGA (EGAS00001001653). SV calls for 70 whole-genome sequenced cholangiocarcinomas were obtained from supplementary information of the associated publication [16].

### Reanalysis of raw sequencing data

Read alignment, variant calling and filtering were performed as described previously [20], except that reads were aligned to GRCh38.p7. For analysis of SV occurrence as a function of transcriptional activity, the transcriptional activity at the location of the SV breakpoint with the highest transcriptional activity was taken.

### Mutational signature analysis

We used the classification for indel mutational signatures as proposed previously [10]; for details see https://www.synapse.org/#!Synapse:syn11801742. Mutational signatures were plotted using ICAMSv2.1.2.9000 (github.com/steverozen/ICAMS).

### Correlation between transcriptional activity and mutagenesis in WGS data

For each tumor type, genes were assigned to 1 of 4 gene expression bins. Subsequently for every sample, variants were grouped by these same expression bins by linking the genomic position of the variant to the genes assigned to expression bins. Mutation density was reported as events per Mb to compensate for differences in size of the total gene expression bins. Cochran-Armitage test for trend was performed to determine statistical significance. As not all ICGC projects have RNA-sequencing data available, for some ICGC cohorts we used RNA-sequencing data from similar cohorts. For details, see Supplementary Table S2.

### Definition of ID6-high and ID8-high tumors in WGS data

For comparison of the size distributions of ≥ 5 bp deletions in ID6-high and ID8-high tumors, we classified tumors using the existing indel signature assignments [10]. ID6-high tumors were those with > the median of counts of ID6 mutations across tumors that had > 0 ID6 mutations. ID8-high tumors were selected analogously. In addition, in selecting ID8-high tumors, tumors with > 0 ID6 mutations were excluded.

### Statistics

Statistical analyses were performed in R v.3.6.3; multiple testing correction was done according to Benjamini-Hochberg method using the p.adjust function. Cochran-Armitage tests were performed using the DescTools package.

## Acknowledgements

We thank Sue Jinks-Robertson (Duke University Medical Center) for sharing preliminary data from her TOP2 yeast data that led us to the discovery of the link between ID17 and TOP2A. We also thank professors Peter S. Nelson and Fergus J. Couch for kindly providing sequencing data and Willie Yu for technical support. This work was supported by a Khoo Postdoctoral Fellowship Award (Duke-NUS-KPFA/2018/0027) to AB and MOH-000032/MOH-CIRG18may-0004 to SGR.

## Author contributions

AB and SGR both contributed to the conception, analysis, writing, and editing of the manuscript.

## Supplementary Material

Sup.Fig. S1: **Density of indels correlates with transcriptional activity in CCA_TH_19**. In-transcript indel spectra for 4 expression-level bins. Association of increased mutational density with increased transcript level is consistent with the known enrichment of TOP2A activity in highly transcribed regions [13].

Sup.Table S1: **Tumors carrying TOP2A or TOP2B hotspot mutations**.

Sup.Table S2: **ID8 mutagenesis (**≥ **5 bp deletions) as a function of transcriptional activity in WGS data**.

Sup.Table S3: **The ID_TOP2A indel mutational signature**.

Sup.Table S4: **Coding indels in known cancer genes in ID_TOP2A tumors**. Somatic indels in cancer driver genes listed in the Cancer Gene Census.

