## Supplementary for "Recurrent mutations in topoisomerase 2a cause a novel mutator phenotype in human cancers"

Sup.Fig_S1 ____ 2
Sup.Table_S1 ____ 3
Sup.Table_S2 ____ 4
Sup.Table_S3 ____ 44
Sup.Table_S4 ____ 48

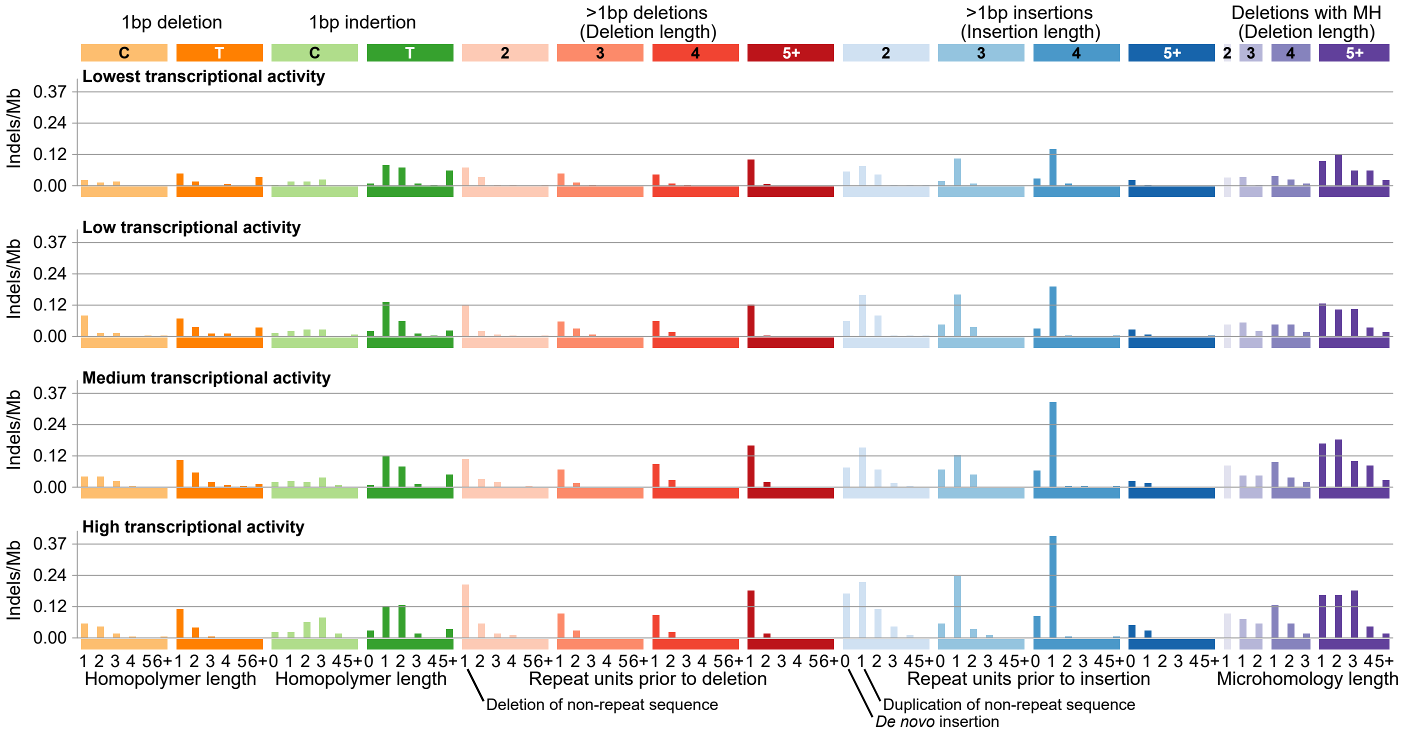

1bp deletion
1bp indertion
>1bp deletions (Deletion length)
>1bp insertions (Insertion length)
Deletions with MH (Deletion length)
C T C T 2 3 4 5+ 2 3 4 5+ 2 3 4 5+
Lowest transcriptional activity
Low transcriptional activity
Medium transcriptional activity
High transcriptional activity
Indels/Mb
0.37
0.24
0.12
0.00
Homopolymer length
Homopolymer length
Repeat units prior to deletion
Repeat units prior to insertion
Microhomology length
Deletion of non-repeat sequence
Duplication of non-repeat sequence
De novo insertion

Supplementary Table S1: Tumors carrying TOP2A or TOP2B hotspot mutations

| | | Sample | | | Mutation load | | | |
|---|---|---|---|---|---|---|---|---|
| Gene | Substitution | CancerType | Sample | DataType | SBS | DBS | ID | Hypermutator phenotype |
| TOP2A | p.T215P | Ovary-AdenoCa | TCGA-04-1343 | WES | 132 | 4 | 6 | |
| TOP2A | p.T215P | Ovary-AdenoCa | TCGA-23-1123 | WES | 108 | 1 | 3 | |
| TOP2A | p.T215P | Ovary-AdenoCa | TCGA-13-1411 | WES | 79 | 2 | 6 | |
| TOP2A | p.T215P | Ovary-AdenoCa | TCGA-23-1027 | WES | 66 | 1 | 7 | |
| TOP2A | p.T215P | Ovary-AdenoCa | TCGA-25-1328 | WES | 40 | 4 | 0 | |
| **TOP2A** | **p.K743N** | **Stomach-AdenoCa** | **TCGA-CG-4477** | **WES** | **72** | **0** | **93** | |
| **TOP2A** | **p.K743N** | **Stomach-AdenoCa** | **TCGA-VQ-A8DL** | **WES** | **64** | **2** | **107** | |
| **TOP2A** | **p.K743N** | **Stomach-AdenoCa** | **TCGA-KB-A93J** | **WES** | **55** | **0** | **93** | |
| **TOP2A** | **p.K743N** | **Stomach-AdenoCa** | **TCGA-CG-5725** | **WES** | **48** | **1** | **69** | |
| **TOP2A** | **p.K743N** | **Biliary-AdenoCa** | **CCA_TH_19** | **WGS** | **3767** | **72** | **NA** | |
| TOP2A | p.R878H | Uterus-AdenoCa | TCGA-EO-A3KX | WES | 4772 | 3 | 1301 | DNA mismatch repair deficiency |
| TOP2A | p.R878C | Stomach-AdenoCa | TCGA-BR-4184 | WES | 4216 | 2 | 425 | DNA mismatch repair deficiency |
| TOP2A | p.R878H | Prost-AdenoCa | TCGA-J9-A52C | WES | 682 | 0 | 70 | DNA mismatch repair deficiency |
| TOP2A | p.R878C | Lung-AdenoCa | LUAD-E00163 | WES | 37 | 0 | NA | |
| TOP2A | p.E1035* | ColoRect-AdenoCa | SP80615 | WGS | 2426469 | 3656 | 19514 | POLE exonuclease domain deficiency |
| TOP2A | p.E1035* | ColoRect-AdenoCa | TCGA-AG-A002 | WES | 16961 | 16 | 75 | POLE exonuclease domain deficiency |
| TOP2A | p.E1035* | Uterus-AdenoCa | TCGA-AP-A056 | WES | 12419 | 17 | 76 | POLE exonuclease domain deficiency |
| TOP2A | p.E1035K | Skin-BCC | 5-PT023-T1 | WES | 975 | 0 | NA | UV-melanoma |
| TOP2A | p.P1457H | Skin-Melanoma | TCGA-BF-A1Q0 | WES | 3932 | 17 | 5 | UV-melanoma |
| TOP2A | p.P1457S | Skin-BCC | 5-VS024-T1 | WES | 3883 | 0 | NA | UV-melanoma |
| TOP2A | p.P1457H | Skin-Melanoma | TCGA-EE-A183 | WES | 3144 | 25 | 4 | UV-melanoma |
| TOP2A | p.P1457S | Skin-BCC | 5-PT013-T1 | WES | 1083 | 0 | NA | UV-melanoma |
| TOP2A | p.S1483L | CNS-GBM | TCGA-06-5416 | WES | 18226 | 16 | 48 | POLE exonuclease domain deficiency |
| TOP2A | p.S1483L | DLBC | TCGA-DU-6392 | WES | 15643 | 13 | 235 | DNA mismatch repair deficiency |
| TOP2A | p.S1483L | Uterus-AdenoCa | TCGA-AX-A05Z | WES | 10628 | 7 | 30 | POLE exonuclease domain deficiency |
| TOP2A | p.S1483L | Uterus-AdenoCa | TCGA-AP-A1E0 | WES | 8399 | 6 | 30 | POLE exonuclease domain deficiency |
| TOP2A | p.S1483L | Uterus-AdenoCa | TCGA-AJ-A5DW | WES | 5621 | 1 | 27 | POLE exonuclease domain deficiency |
| TOP2A | p.S1483L | Transitional-cell-carcinoma | TCGA-XF-A9T3 | WES | 544 | 4 | 9 | |
| TOP2B | p.R651H | Uterus-AdenoCa | TCGA-FI-A2D0 | WES | 11400 | 8 | 442 | DNA mismatch repair deficiency |
| TOP2B | p.R651H | ColoRect-AdenoCa | TCGA-CK-4951 | WES | 6442 | 22 | 313 | DNA mismatch repair deficiency |
| TOP2B | p.R651H | Stomach-AdenoCa | TCGA-BR-4184 | WES | 4216 | 2 | 425 | DNA mismatch repair deficiency |
| TOP2B | p.R651H | ColoRect-AdenoCa | TCGA-NH-A5IV | WES | 3072 | 2 | 769 | DNA mismatch repair deficiency |
| TOP2B | p.R651H | ColoRect-AdenoCa | TCGA-G4-6628 | WES | 2438 | 1 | 438 | DNA mismatch repair deficiency |
| TOP2B | p.R651H | ColoRect-AdenoCa | TCGA-AA-A01R | WES | 2542 | 1 | 86 | DNA mismatch repair deficiency |
| TOP2B | p.R651H | ColoRect-AdenoCa | TCGA-AA-A022 | WES | 1935 | 1 | 416 | DNA mismatch repair deficiency |
| TOP2B | p.R651H | Stomach-AdenoCa | TCGA-HF-A5NB | WES | 1571 | 1 | 409 | DNA mismatch repair deficiency |
| TOP2B | p.R651H | Uterus-AdenoCa | TCGA-EC-A24G | WES | 1797 | 0 | 107 | DNA mismatch repair deficiency |
| TOP2B | p.R651H | Uterus-AdenoCa | TCGA-AX-A063 | WES | 1324 | 1 | 176 | DNA mismatch repair deficiency |
| TOP2B | p.R651H | Biliary-AdenoCa | BD135T | WES | 1062 | 4 | NA | DNA mismatch repair deficiency |
| TOP2B | p.R651H | Panc-AdenoCa | TCGA-HV-A7OP | WES | 110 | 0 | 10 | |
| TOP2B | p.R651C | Liver-HCC | BCM703T | WES | 74 | 0 | NA | |
| TOP2B | p.R651H | CNS-Medullo | SP78805 | WGS | 1164 | 4 | 143 | |
| TOP2B | p.R651H | CNS-LGG | ICGC_MB64 | WES | 13 | 0 | NA | |

Supplementary Table S2: ID8 mutagenesis (≥ 5 bp deletions) as a function of transcriptional activity in WGS data

| | | | Indel spectrum and indel signature composition | | | | | Mutation density by gene expression | | | | Cochran Armitage test for trend, two.sided | | | |
|---|---|---|---|---|---|---|---|---|---|---|---|---|---|---|---|
| Sample | tissue | GE data used* | Total indels | ID6 | ID8 | ≥ 5 bp deletions (n) | Median size of ≥ 5 bp deletions (bp)# | Lowest GE | Low GE | Medium GE | High GE | Z | pval | FDR (BH adjusted) | Direction of trend |
| LIHC-US::SP49124 | Liver | LIHC-US | 1284 | 0 | 707 | 297 | 16 | 0.123 | 0.178 | 0.342 | 0.502 | -8.7206 | 2.77E-18 | 4.01E-15 | Increasing |
| LIRI-JP::SP99197 | Liver | LIRI-JP | 1726 | 299 | 635 | 333 | 13.5 | 0.165 | 0.208 | 0.293 | 0.614 | -8.5007 | 1.88E-17 | 1.37E-14 | Increasing |
| BLCA-US::SP975 | Bladder | BLCA-US | 1363 | 0 | 1039 | 353 | 17 | 0.156 | 0.216 | 0.453 | 0.497 | -8.3237 | 8.53E-17 | 4.12E-14 | Increasing |
| LIRI-JP::SP107076 | Liver | LIRI-JP | 1736 | 0 | 842 | 362 | 15 | 0.181 | 0.243 | 0.353 | 0.562 | -7.5103 | 5.90E-14 | 2.14E-11 | Increasing |
| BLCA-US::SP1305 | Bladder | BLCA-US | 1830 | 0 | 904 | 350 | 15 | 0.144 | 0.290 | 0.397 | 0.436 | -6.6628 | 2.69E-11 | 7.79E-09 | Increasing |
| LINC-JP::SP98853 | Liver | LIHC-US | 934 | 0 | 123 | 77 | 12 | 0.022 | 0.042 | 0.071 | 0.191 | -6.4765 | 9.39E-11 | 2.27E-08 | Increasing |
| LIRI-JP::SP112248 | Liver | LIRI-JP | 722 | 0 | 182 | 89 | 17 | 0.029 | 0.035 | 0.118 | 0.168 | -6.3557 | 2.07E-10 | 4.30E-08 | Increasing |
| LIRI-JP::SP107025 | Liver | LIRI-JP | 615 | 0 | 182 | 86 | 15 | 0.016 | 0.055 | 0.099 | 0.162 | -6.1299 | 8.79E-10 | 1.59E-07 | Increasing |
| LINC-JP::SP98891 | Liver | LIHC-US | 613 | 0 | 121 | 48 | 13 | 0.006 | 0.026 | 0.045 | 0.132 | -6.0771 | 1.22E-09 | 1.97E-07 | Increasing |
| LIRI-JP::SP112290 | Liver | LIRI-JP | 1041 | 0 | 193 | 95 | 13 | 0.032 | 0.048 | 0.121 | 0.162 | -5.7728 | 7.80E-09 | 1.13E-06 | Increasing |
| LIRI-JP::SP99069 | Liver | LIRI-JP | 812 | 78 | 206 | 119 | 15 | 0.038 | 0.073 | 0.150 | 0.180 | -5.7134 | 1.11E-08 | 1.46E-06 | Increasing |
| LIRI-JP::SP112205 | Liver | LIRI-JP | 642 | 0 | 113 | 45 | 17.5 | 0.010 | 0.013 | 0.067 | 0.093 | -5.4831 | 4.18E-08 | 5.05E-06 | Increasing |
| LIRI-JP::SP112178 | Liver | LIRI-JP | 1091 | 0 | 231 | 98 | 13 | 0.045 | 0.045 | 0.118 | 0.168 | -5.2968 | 1.18E-07 | 1.31E-05 | Increasing |
| LIRI-JP::SP99049 | Liver | LIRI-JP | 853 | 0 | 228 | 102 | 13 | 0.041 | 0.058 | 0.115 | 0.174 | -5.2755 | 1.32E-07 | 1.37E-05 | Increasing |
| MALY-DE::SP116670 | Blood | MALY-DE | 1207 | 0 | 0 | 33 | 11 | 0.006 | 0.009 | 0.009 | 0.075 | -5.2428 | 1.58E-07 | 1.53E-05 | Increasing |
| LINC-JP::SP98867 | Liver | LIHC-US | 817 | 0 | 159 | 60 | 13.5 | 0.016 | 0.034 | 0.071 | 0.120 | -5.1353 | 2.82E-07 | 2.55E-05 | Increasing |
| LIRI-JP::SP99011 | Liver | LIRI-JP | 859 | 0 | 153 | 82 | 14 | 0.025 | 0.055 | 0.083 | 0.151 | -5.0888 | 3.60E-07 | 3.07E-05 | Increasing |
| LIRI-JP::SP98973 | Liver | LIRI-JP | 1701 | 268 | 229 | 173 | 14 | 0.124 | 0.065 | 0.191 | 0.278 | -5.0773 | 3.83E-07 | 3.08E-05 | Increasing |
| LIRI-JP::SP98955 | Liver | LIRI-JP | 1530 | 0 | 320 | 163 | 14 | 0.067 | 0.113 | 0.194 | 0.209 | -5.0654 | 4.08E-07 | 3.11E-05 | Increasing |
| LIRI-JP::SP107136 | Liver | LIRI-JP | 879 | 0 | 195 | 91 | 13 | 0.032 | 0.063 | 0.089 | 0.162 | -4.9778 | 6.43E-07 | 4.66E-05 | Increasing |
| LIRI-JP::SP107004 | Liver | LIRI-JP | 704 | 0 | 141 | 79 | 17 | 0.029 | 0.048 | 0.083 | 0.145 | -4.9459 | 7.58E-07 | 5.23E-05 | Increasing |
| LIRI-JP::SP50113 | Liver | LIRI-JP | 1163 | 0 | 156 | 84 | 15 | 0.035 | 0.050 | 0.083 | 0.156 | -4.8575 | 1.19E-06 | 7.84E-05 | Increasing |
| LIRI-JP::SP98997 | Liver | LIRI-JP | 961 | 0 | 145 | 85 | 13 | 0.029 | 0.048 | 0.115 | 0.122 | -4.7979 | 1.60E-06 | 9.72E-05 | Increasing |
| LICA-FR::SP97685 | Liver | LICA-FR | 506 | 0 | 160 | 85 | 13 | 0.020 | 0.059 | 0.076 | 0.131 | -4.7972 | 1.61E-06 | 9.72E-05 | Increasing |
| LIRI-JP::SP112190 | Liver | LIRI-JP | 1085 | 0 | 222 | 101 | 14 | 0.038 | 0.068 | 0.115 | 0.151 | -4.6389 | 3.50E-06 | 2.03E-04 | Increasing |
| LIRI-JP::SP112051 | Liver | LIRI-JP | 773 | 0 | 150 | 77 | 14 | 0.035 | 0.040 | 0.083 | 0.139 | -4.6224 | 3.79E-06 | 2.12E-04 | Increasing |
| LIRI-JP::SP99257 | Liver | LIRI-JP | 1066 | 0 | 210 | 94 | 12.5 | 0.035 | 0.055 | 0.124 | 0.127 | -4.6050 | 4.13E-06 | 2.22E-04 | Increasing |
| LIRI-JP::SP99233 | Liver | LIRI-JP | 457 | 0 | 88 | 42 | 14 | 0.006 | 0.028 | 0.045 | 0.087 | -4.5828 | 4.59E-06 | 2.38E-04 | Increasing |
| BLCA-US::SP1491 | Bladder | BLCA-US | 657 | 0 | 304 | 117 | 15 | 0.052 | 0.092 | 0.108 | 0.194 | -4.5013 | 6.75E-06 | 3.38E-04 | Increasing |
| LIRI-JP::SP99313 | Liver | LIRI-JP | 544 | 0 | 280 | 114 | 15 | 0.032 | 0.105 | 0.105 | 0.168 | -4.4841 | 7.32E-06 | 3.50E-04 | Increasing |
| LINC-JP::SP98881 | Liver | LIHC-US | 821 | 0 | 190 | 84 | 11 | 0.035 | 0.065 | 0.064 | 0.167 | -4.4798 | 7.47E-06 | 3.50E-04 | Increasing |
| LIRI-JP::SP50169 | Liver | LIRI-JP | 782 | 0 | 119 | 53 | 14 | 0.003 | 0.038 | 0.080 | 0.070 | -4.4668 | 7.94E-06 | 3.60E-04 | Increasing |
| LIRI-JP::SP99241 | Liver | LIRI-JP | 864 | 0 | 208 | 90 | 15 | 0.032 | 0.058 | 0.115 | 0.122 | -4.4080 | 1.04E-05 | 4.58E-04 | Increasing |
| LIRI-JP::SP112310 | Liver | LIRI-JP | 738 | 0 | 134 | 41 | 16 | 0.010 | 0.015 | 0.067 | 0.064 | -4.3730 | 1.23E-05 | 5.23E-04 | Increasing |
| LIRI-JP::SP112133 | Liver | LIRI-JP | 656 | 0 | 132 | 65 | 16 | 0.016 | 0.043 | 0.086 | 0.093 | -4.3477 | 1.38E-05 | 5.70E-04 | Increasing |
| LIRI-JP::SP107080 | Liver | LIRI-JP | 786 | 0 | 322 | 118 | 18 | 0.067 | 0.075 | 0.102 | 0.203 | -4.3025 | 1.69E-05 | 6.80E-04 | Increasing |
| LIRI-JP::SP107101 | Liver | LIRI-JP | 1223 | 0 | 199 | 87 | 14 | 0.032 | 0.068 | 0.080 | 0.145 | -4.2559 | 2.08E-05 | 8.10E-04 | Increasing |
| LIRI-JP::SP112211 | Liver | LIRI-JP | 881 | 0 | 136 | 54 | 11 | 0.016 | 0.045 | 0.032 | 0.122 | -4.2514 | 2.12E-05 | 8.10E-04 | Increasing |
| LIRI-JP::SP107047 | Liver | LIRI-JP | 810 | 0 | 133 | 51 | 15 | 0.019 | 0.028 | 0.054 | 0.098 | -4.2165 | 2.48E-05 | 9.22E-04 | Increasing |
| MALY-DE::SP116676 | Blood | MALY-DE | 1873 | 0 | 0 | 48 | 10 | 0.013 | 0.025 | 0.020 | 0.084 | -4.1969 | 2.71E-05 | 9.81E-04 | Increasing |
| LIRI-JP::SP99021 | Liver | LIRI-JP | 534 | 0 | 103 | 48 | 16 | 0.019 | 0.028 | 0.041 | 0.104 | -4.1833 | 2.87E-05 | 1.02E-03 | Increasing |
| LIRI-JP::SP99073 | Liver | LIRI-JP | 732 | 0 | 112 | 50 | 13 | 0.025 | 0.018 | 0.057 | 0.098 | -4.1585 | 3.20E-05 | 1.11E-03 | Increasing |
| OV-AU::SP101931 | Ovary | OV-AU | 896 | 344 | 226 | 136 | 12 | 0.169 | 0.107 | 0.072 | 0.062 | 4.1123 | 3.92E-05 | 1.32E-03 | Decreasing |
| LINC-JP::SP98849 | Liver | LIHC-US | 308 | 0 | 59 | 27 | 11 | 0.006 | 0.010 | 0.035 | 0.060 | -4.0822 | 4.46E-05 | 1.47E-03 | Increasing |
| LIRI-JP::SP50149 | Liver | LIRI-JP | 706 | 0 | 145 | 51 | 14 | 0.010 | 0.030 | 0.080 | 0.064 | -4.0776 | 4.55E-05 | 1.47E-03 | Increasing |
| LIRI-JP::SP98902 | Liver | LIRI-JP | 868 | 0 | 213 | 99 | 15 | 0.041 | 0.068 | 0.115 | 0.133 | -4.0449 | 5.23E-05 | 1.65E-03 | Increasing |

| | | | | | | | | | | | | | | | |
|---|---|---|---|---|---|---|---|---|---|---|---|---|---|---|---|
| LIRI-JP::SP112210 | Liver | LIRI-JP | 924 | 0 | 134 | 57 | 13 | 0.025 | 0.033 | 0.054 | 0.110 | -4.0248 | 5.70E-05 | 1.76E-03 | Increasing |
| LIRI-JP::SP99125 | Liver | LIRI-JP | 496 | 0 | 85 | 46 | 13 | 0.006 | 0.035 | 0.057 | 0.070 | -3.9185 | 8.91E-05 | 2.69E-03 | Increasing |
| MALY-DE::SP116715 | Blood | MALY-DE | 3491 | 0 | 0 | 43 | 11 | 0.026 | 0.011 | 0.029 | 0.072 | -3.8732 | 1.07E-04 | 3.18E-03 | Increasing |
| LICA-FR::SP97681 | Liver | LICA-FR | 698 | 0 | 102 | 41 | 12 | 0.010 | 0.026 | 0.032 | 0.075 | -3.8443 | 1.21E-04 | 3.51E-03 | Increasing |
| RECA-EU::SP103715 | Kidney | RECA-EU | 617 | 0 | 97 | 35 | 9.5 | 0.010 | 0.019 | 0.028 | 0.070 | -3.8180 | 1.35E-04 | 3.83E-03 | Increasing |
| LIRI-JP::SP50147 | Liver | LIRI-JP | 808 | 0 | 124 | 44 | 13 | 0.022 | 0.025 | 0.029 | 0.104 | -3.7933 | 1.49E-04 | 4.14E-03 | Increasing |
| LIRI-JP::SP112192 | Liver | LIRI-JP | 685 | 0 | 146 | 57 | 15 | 0.019 | 0.035 | 0.073 | 0.081 | -3.7620 | 1.69E-04 | 4.61E-03 | Increasing |
| BRCA-US::SP13036 | Breast | BRCA-US | 961 | 0 | 173 | 34 | 10 | 0.017 | 0.008 | 0.041 | 0.069 | -3.7482 | 1.78E-04 | 4.78E-03 | Increasing |
| LIHC-US::SP49417 | Liver | LIHC-US | 679 | 0 | 107 | 47 | 12 | 0.028 | 0.018 | 0.045 | 0.102 | -3.7303 | 1.91E-04 | 4.97E-03 | Increasing |
| LIRI-JP::SP112194 | Liver | LIRI-JP | 703 | 0 | 97 | 43 | 13 | 0.013 | 0.025 | 0.054 | 0.070 | -3.7294 | 1.92E-04 | 4.97E-03 | Increasing |
| LIRI-JP::SP112252 | Liver | LIRI-JP | 927 | 0 | 104 | 52 | 13 | 0.022 | 0.028 | 0.060 | 0.087 | -3.7236 | 1.96E-04 | 5.00E-03 | Increasing |
| LIRI-JP::SP99061 | Liver | LIRI-JP | 1317 | 0 | 118 | 56 | 13 | 0.013 | 0.043 | 0.070 | 0.075 | -3.7010 | 2.15E-04 | 5.37E-03 | Increasing |
| PACA-AU::SP71480 | Pancreas | PACA-AU | 578 | 0 | 0 | 13 | 13.5 | 0.000 | 0.000 | 0.015 | 0.027 | -3.6930 | 2.22E-04 | 5.45E-03 | Increasing |
| LIRI-JP::SP98975 | Liver | LIRI-JP | 380 | 0 | 71 | 27 | 14 | 0.010 | 0.008 | 0.038 | 0.052 | -3.6744 | 2.38E-04 | 5.71E-03 | Increasing |
| LIRI-JP::SP112219 | Liver | LIRI-JP | 1000 | 0 | 211 | 88 | 15 | 0.045 | 0.055 | 0.095 | 0.127 | -3.6726 | 2.40E-04 | 5.71E-03 | Increasing |
| UCEC-US::SP92931 | Endometrium | UCEC-US | 531 | 0 | 74 | 29 | 11 | 0.006 | 0.018 | 0.033 | 0.058 | -3.6298 | 2.84E-04 | 6.63E-03 | Increasing |
| LIRI-JP::SP107104 | Liver | LIRI-JP | 407 | 0 | 75 | 34 | 14 | 0.013 | 0.020 | 0.029 | 0.075 | -3.6129 | 3.03E-04 | 6.97E-03 | Increasing |
| STAD-US::SP84491 | Stomach | STAD-US | 3278 | 0 | 0 | 90 | 12 | 0.049 | 0.050 | 0.108 | 0.122 | -3.6065 | 3.10E-04 | 7.03E-03 | Increasing |
| LINC-JP::SP98875 | Liver | LIHC-US | 817 | 0 | 127 | 62 | 14 | 0.022 | 0.039 | 0.087 | 0.078 | -3.6006 | 3.18E-04 | 7.04E-03 | Increasing |
| LIRI-JP::SP112265 | Liver | LIRI-JP | 557 | 0 | 86 | 41 | 16 | 0.013 | 0.028 | 0.041 | 0.075 | -3.5984 | 3.20E-04 | 7.04E-03 | Increasing |
| LIRI-JP::SP112196 | Liver | LIRI-JP | 793 | 0 | 130 | 66 | 15 | 0.029 | 0.050 | 0.054 | 0.116 | -3.5469 | 3.90E-04 | 8.19E-03 | Increasing |
| LIRI-JP::SP50107 | Liver | LIRI-JP | 989 | 0 | 125 | 62 | 13 | 0.013 | 0.060 | 0.057 | 0.093 | -3.5522 | 3.82E-04 | 8.19E-03 | Increasing |
| LIRI-JP::SP50167 | Liver | LIRI-JP | 621 | 0 | 138 | 76 | 17 | 0.035 | 0.055 | 0.070 | 0.122 | -3.5475 | 3.89E-04 | 8.19E-03 | Increasing |
| LIRI-JP::SP98941 | Liver | LIRI-JP | 910 | 0 | 99 | 40 | 13 | 0.013 | 0.020 | 0.057 | 0.058 | -3.5312 | 4.14E-04 | 8.57E-03 | Increasing |
| LIRI-JP::SP112273 | Liver | LIRI-JP | 1315 | 0 | 0 | 51 | 14 | 0.016 | 0.040 | 0.048 | 0.087 | -3.5220 | 4.28E-04 | 8.75E-03 | Increasing |
| LIRI-JP::SP112168 | Liver | LIRI-JP | 872 | 0 | 135 | 53 | 11.5 | 0.019 | 0.043 | 0.041 | 0.098 | -3.5129 | 4.43E-04 | 8.93E-03 | Increasing |
| LIRI-JP::SP107071 | Liver | LIRI-JP | 781 | 0 | 165 | 69 | 14 | 0.032 | 0.053 | 0.054 | 0.122 | -3.4854 | 4.91E-04 | 9.76E-03 | Increasing |
| OV-AU::SP101558 | Ovary | OV-AU | 534 | 237 | 259 | 127 | 19 | 0.153 | 0.102 | 0.057 | 0.078 | 3.4728 | 5.15E-04 | 1.01E-02 | Decreasing |
| MALY-DE::SP116697 | Blood | MALY-DE | 9419 | 0 | 0 | 315 | 12 | 0.172 | 0.247 | 0.168 | 0.359 | -3.4186 | 6.29E-04 | 1.22E-02 | Increasing |
| LUAD-US::SP56168 | Lung | LUAD-US | 598 | 0 | 88 | 31 | 12 | 0.000 | 0.033 | 0.029 | 0.054 | -3.4105 | 6.48E-04 | 1.24E-02 | Increasing |
| LIRI-JP::SP99053 | Liver | LIRI-JP | 413 | 0 | 58 | 23 | 13 | 0.003 | 0.018 | 0.019 | 0.052 | -3.3914 | 6.95E-04 | 1.31E-02 | Increasing |
| STAD-US::SP105673 | Stomach | STAD-US | 1382 | 606 | 190 | 237 | 13 | 0.302 | 0.143 | 0.196 | 0.149 | 3.3717 | 7.47E-04 | 1.39E-02 | Decreasing |
| LIRI-JP::SP99065 | Liver | LIRI-JP | 485 | 0 | 64 | 28 | 12 | 0.003 | 0.020 | 0.035 | 0.046 | -3.3669 | 7.60E-04 | 1.40E-02 | Increasing |
| LINC-JP::SP98887 | Liver | LIHC-US | 936 | 0 | 120 | 58 | 12 | 0.022 | 0.044 | 0.061 | 0.090 | -3.3485 | 8.13E-04 | 1.47E-02 | Increasing |
| STAD-US::SP105261 | Stomach | STAD-US | 544 | 0 | 77 | 22 | 13 | 0.009 | 0.006 | 0.026 | 0.048 | -3.3388 | 8.41E-04 | 1.51E-02 | Increasing |
| LIRI-JP::SP50181 | Liver | LIRI-JP | 503 | 0 | 70 | 30 | 14 | 0.006 | 0.020 | 0.035 | 0.052 | -3.3298 | 8.69E-04 | 1.54E-02 | Increasing |
| LIRI-JP::SP107119 | Liver | LIRI-JP | 729 | 0 | 79 | 39 | 13 | 0.016 | 0.025 | 0.035 | 0.075 | -3.3041 | 9.53E-04 | 1.66E-02 | Increasing |
| LIHC-US::SP98297 | Liver | LIHC-US | 758 | 0 | 121 | 49 | 10 | 0.028 | 0.024 | 0.052 | 0.090 | -3.2920 | 9.95E-04 | 1.72E-02 | Increasing |
| LIRI-JP::SP107012 | Liver | LIRI-JP | 30199 | 0 | 0 | 34 | 8 | 0.016 | 0.020 | 0.025 | 0.075 | -3.2727 | 1.07E-03 | 1.82E-02 | Increasing |
| LINC-JP::SP98865 | Liver | LIHC-US | 288 | 0 | 51 | 21 | 11 | 0.006 | 0.013 | 0.016 | 0.054 | -3.2646 | 1.10E-03 | 1.85E-02 | Increasing |
| BRCA-UK::SP116379 | Breast | BRCA-US | 294 | 0 | 80 | 15 | 9 | 0.003 | 0.005 | 0.016 | 0.037 | -3.2490 | 1.16E-03 | 1.93E-02 | Increasing |
| LIRI-JP::SP107056 | Liver | LIRI-JP | 929 | 0 | 123 | 53 | 14 | 0.019 | 0.043 | 0.048 | 0.087 | -3.2404 | 1.19E-03 | 1.94E-02 | Increasing |
| LIRI-JP::SP50179 | Liver | LIRI-JP | 514 | 0 | 49 | 21 | 12 | 0.010 | 0.008 | 0.019 | 0.052 | -3.2405 | 1.19E-03 | 1.94E-02 | Increasing |
| RECA-EU::SP102783 | Kidney | RECA-EU | 1202 | 0 | 192 | 67 | 9 | 0.033 | 0.040 | 0.057 | 0.100 | -3.2197 | 1.28E-03 | 2.07E-02 | Increasing |
| PRAD-UK::SP114992 | Prostate | PRAD-US | 1420 | 534 | 586 | 278 | 17 | 0.317 | 0.242 | 0.192 | 0.191 | 3.2114 | 1.32E-03 | 2.10E-02 | Decreasing |
| LIRI-JP::SP112180 | Liver | LIRI-JP | 677 | 0 | 100 | 42 | 11 | 0.022 | 0.030 | 0.019 | 0.098 | -3.2051 | 1.35E-03 | 2.13E-02 | Increasing |
| OV-US::SP67609 | Ovary | OV-US | 1517 | 1041 | 476 | 332 | 14 | 0.383 | 0.249 | 0.260 | 0.220 | 3.1944 | 1.40E-03 | 2.19E-02 | Decreasing |
| LIRI-JP::SP50097 | Liver | LIRI-JP | 1433 | 0 | 130 | 62 | 14 | 0.025 | 0.043 | 0.073 | 0.081 | -3.1743 | 1.50E-03 | 2.30E-02 | Increasing |
| COAD-US::SP16958 | CRC | COAD-US | 2334 | 0 | 0 | 56 | 10 | 0.027 | 0.038 | 0.056 | 0.091 | -3.1731 | 1.51E-03 | 2.30E-02 | Increasing |
| OV-AU::SP102045 | Ovary | OV-AU | 2008 | 1334 | 674 | 460 | 13 | 0.421 | 0.388 | 0.311 | 0.274 | 3.1552 | 1.60E-03 | 2.42E-02 | Decreasing |

| | | | | | | | | | | | | | | | |
|---|---|---|---|---|---|---|---|---|---|---|---|---|---|---|---|
| OV-US::SP64546 | Ovary | OV-US | 577 | 309 | 219 | 118 | 21 | 0.160 | 0.084 | 0.076 | 0.075 | 3.1354 | 1.72E-03 | 2.57E-02 | Decreasing |
| BLCA-US::SP1132 | Bladder | BLCA-US | 847 | 0 | 107 | 49 | 9 | 0.009 | 0.050 | 0.053 | 0.067 | -3.1235 | 1.79E-03 | 2.64E-02 | Increasing |
| LIRI-JP::SP112208 | Liver | LIRI-JP | 882 | 0 | 134 | 65 | 14 | 0.032 | 0.033 | 0.095 | 0.070 | -3.1172 | 1.83E-03 | 2.67E-02 | Increasing |
| MALY-DE::SP124969 | Blood | MALY-DE | 1765 | 0 | 0 | 30 | 13 | 0.000 | 0.018 | 0.020 | 0.045 | -3.1112 | 1.86E-03 | 2.70E-02 | Increasing |
| LIRI-JP::SP107037 | Liver | LIRI-JP | 741 | 0 | 136 | 53 | 12.5 | 0.019 | 0.030 | 0.083 | 0.052 | -3.1041 | 1.91E-03 | 2.74E-02 | Increasing |
| PAEN-AU::SP102567 | Pancreas | PAEN-AU | 239 | 0 | 34 | 12 | 12 | 0.005 | 0.003 | 0.000 | 0.030 | -3.0964 | 1.96E-03 | 2.79E-02 | Increasing |
| PRAD-UK::SP114898 | Prostate | PRAD-US | 997 | 0 | 0 | 15 | 9 | 0.000 | 0.008 | 0.018 | 0.030 | -3.0863 | 2.03E-03 | 2.83E-02 | Increasing |
| LIRI-JP::SP50115 | Liver | LIRI-JP | 462 | 0 | 73 | 36 | 13 | 0.010 | 0.025 | 0.045 | 0.052 | -3.0855 | 2.03E-03 | 2.83E-02 | Increasing |
| LIHC-US::SP109301 | Liver | LIHC-US | 755 | 0 | 81 | 31 | 12 | 0.009 | 0.018 | 0.042 | 0.048 | -3.0787 | 2.08E-03 | 2.87E-02 | Increasing |
| PACA-CA::SP117974 | Pancreas | PACA-CA | 681 | 0 | 0 | 11 | 11 | 0.000 | 0.003 | 0.008 | 0.022 | -3.0699 | 2.14E-03 | 2.93E-02 | Increasing |
| MALY-DE::SP59452 | Blood | MALY-DE | 1400 | 0 | 0 | 23 | 9 | 0.006 | 0.009 | 0.014 | 0.039 | -3.0669 | 2.16E-03 | 2.93E-02 | Increasing |
| LIRI-JP::SP50163 | Liver | LIRI-JP | 538 | 0 | 77 | 24 | 13 | 0.006 | 0.013 | 0.032 | 0.041 | -3.0593 | 2.22E-03 | 2.98E-02 | Increasing |
| LIRI-JP::SP107121 | Liver | LIRI-JP | 1000 | 0 | 153 | 64 | 12 | 0.032 | 0.040 | 0.073 | 0.087 | -3.0531 | 2.27E-03 | 3.01E-02 | Increasing |
| BTCA-SG::SP117621 | Biliary | BTCA-SG | 875 | 0 | 0 | 20 | 11 | 0.009 | 0.013 | 0.012 | 0.056 | -3.0438 | 2.34E-03 | 3.08E-02 | Increasing |
| PRAD-CA::SP112941 | Prostate | PRAD-US | 245 | 0 | 32 | 14 | 11 | 0.000 | 0.008 | 0.015 | 0.030 | -3.0342 | 2.41E-03 | 3.15E-02 | Increasing |
| LIHC-US::SP49651 | Liver | LIHC-US | 803 | 445 | 221 | 134 | 11 | 0.161 | 0.107 | 0.100 | 0.066 | 3.0264 | 2.48E-03 | 3.20E-02 | Decreasing |
| LINC-JP::SP98889 | Liver | LIHC-US | 813 | 0 | 133 | 53 | 12 | 0.013 | 0.052 | 0.055 | 0.072 | -3.0136 | 2.58E-03 | 3.31E-02 | Increasing |
| LIRI-JP::SP112152 | Liver | LIRI-JP | 460 | 0 | 0 | 15 | 10.5 | 0.003 | 0.008 | 0.016 | 0.035 | -2.9949 | 2.75E-03 | 3.49E-02 | Increasing |
| PACA-CA::SP125752 | Pancreas | PACA-CA | 812 | 0 | 0 | 24 | 14 | 0.004 | 0.014 | 0.017 | 0.038 | -2.9903 | 2.79E-03 | 3.51E-02 | Increasing |
| KIRC-US::SP42154 | Kidney | KIRC-US | 682 | 0 | 83 | 45 | 8.5 | 0.018 | 0.030 | 0.046 | 0.069 | -2.9850 | 2.84E-03 | 3.53E-02 | Increasing |
| PACA-AU::SP76867 | Pancreas | PACA-AU | 562 | 0 | 0 | 17 | 12.5 | 0.004 | 0.005 | 0.015 | 0.030 | -2.9839 | 2.85E-03 | 3.53E-02 | Increasing |
| LIRI-JP::SP50119 | Liver | LIRI-JP | 620 | 0 | 130 | 59 | 12.5 | 0.019 | 0.048 | 0.070 | 0.070 | -2.9778 | 2.90E-03 | 3.57E-02 | Increasing |
| LIRI-JP::SP112170 | Liver | LIRI-JP | 941 | 0 | 131 | 51 | 12 | 0.025 | 0.038 | 0.038 | 0.093 | -2.9663 | 3.01E-03 | 3.64E-02 | Increasing |
| LIRI-JP::SP50109 | Liver | LIRI-JP | 565 | 0 | 98 | 47 | 13 | 0.013 | 0.033 | 0.070 | 0.046 | -2.9668 | 3.01E-03 | 3.64E-02 | Increasing |
| LIRI-JP::SP112160 | Liver | LIRI-JP | 570 | 0 | 64 | 20 | 13 | 0.010 | 0.008 | 0.019 | 0.046 | -2.9406 | 3.28E-03 | 3.93E-02 | Increasing |
| MALY-DE::SP116654 | Blood | MALY-DE | 456 | 0 | 0 | 11 | 12 | 0.000 | 0.002 | 0.009 | 0.021 | -2.9258 | 3.44E-03 | 4.08E-02 | Increasing |
| LINC-JP::SP98845 | Liver | LIHC-US | 9355 | 0 | 0 | 29 | 10 | 0.009 | 0.021 | 0.029 | 0.054 | -2.9185 | 3.52E-03 | 4.15E-02 | Increasing |
| PACA-CA::SP125713 | Pancreas | PACA-CA | 1918 | 1019 | 468 | 377 | 15 | 0.362 | 0.299 | 0.290 | 0.226 | 2.9102 | 3.61E-03 | 4.21E-02 | Decreasing |
| BRCA-US::SP5559 | Breast | BRCA-US | 1275 | 0 | 182 | 74 | 10 | 0.028 | 0.066 | 0.072 | 0.095 | -2.9084 | 3.63E-03 | 4.21E-02 | Increasing |
| CLLE-ES::SP96882 | Blood | CLLE-ES | 222 | 0 | 42 | 22 | 9 | 0.008 | 0.012 | 0.013 | 0.042 | -2.8923 | 3.82E-03 | 4.39E-02 | Increasing |
| LIHC-US::SP49385 | Liver | LIHC-US | 632 | 0 | 146 | 57 | 14 | 0.035 | 0.034 | 0.055 | 0.096 | -2.8906 | 3.85E-03 | 4.39E-02 | Increasing |
| LUAD-US::SP55235 | Lung | LUAD-US | 3235 | 0 | 0 | 74 | 14 | 0.041 | 0.057 | 0.062 | 0.114 | -2.8765 | 4.02E-03 | 4.48E-02 | Increasing |
| LUSC-US::SP57538 | Lung | LUSC-US | 1633 | 0 | 258 | 55 | 10 | 0.026 | 0.034 | 0.072 | 0.068 | -2.8778 | 4.00E-03 | 4.48E-02 | Increasing |
| BRCA-US::SP7171 | Breast | BRCA-US | 358 | 0 | 101 | 23 | 12 | 0.010 | 0.011 | 0.022 | 0.048 | -2.8784 | 4.00E-03 | 4.48E-02 | Increasing |
| OV-US::SP64296 | Ovary | OV-US | 1005 | 472 | 402 | 209 | 18 | 0.245 | 0.170 | 0.132 | 0.157 | 2.8715 | 4.08E-03 | 4.52E-02 | Decreasing |
| GBM-US::SP23925 | Brain | GBM-US | 1759 | 0 | 0 | 34 | 5 | 0.014 | 0.017 | 0.042 | 0.051 | -2.8602 | 4.23E-03 | 4.62E-02 | Increasing |
| BRCA-US::SP7378 | Breast | BRCA-US | 202 | 0 | 78 | 13 | 15 | 0.003 | 0.005 | 0.013 | 0.032 | -2.8615 | 4.22E-03 | 4.62E-02 | Increasing |
| BLCA-US::SP1781 | Bladder | BLCA-US | 822 | 0 | 0 | 25 | 15.5 | 0.009 | 0.011 | 0.039 | 0.036 | -2.8503 | 4.37E-03 | 4.73E-02 | Increasing |
| LIRI-JP::SP50129 | Liver | LIRI-JP | 582 | 0 | 69 | 26 | 10 | 0.006 | 0.020 | 0.025 | 0.046 | -2.8275 | 4.69E-03 | 0.0504 | Increasing |
| LIRI-JP::SP112304 | Liver | LIRI-JP | 600 | 54 | 76 | 47 | 12 | 0.025 | 0.030 | 0.041 | 0.081 | -2.8221 | 4.77E-03 | 0.0506 | Increasing |
| LIRI-JP::SP50117 | Liver | LIRI-JP | 603 | 0 | 102 | 43 | 12 | 0.025 | 0.025 | 0.035 | 0.081 | -2.8216 | 4.78E-03 | 0.0506 | Increasing |
| BLCA-US::SP96136 | Bladder | BLCA-US | 1156 | 0 | 442 | 163 | 15 | 0.092 | 0.145 | 0.154 | 0.188 | -2.8087 | 4.97E-03 | 0.0523 | Increasing |
| PACA-CA::SP125724 | Pancreas | PACA-CA | 615 | 0 | 0 | 14 | 12 | 0.004 | 0.006 | 0.006 | 0.029 | -2.8045 | 5.04E-03 | 0.0526 | Increasing |
| CESC-US::SP107595 | Cervix | CESC-US | 718 | 220 | 0 | 36 | 11 | 0.054 | 0.025 | 0.020 | 0.012 | 2.7959 | 5.18E-03 | 0.0536 | Decreasing |
| CESC-US::SP13072 | Cervix | CESC-US | 314 | 53 | 0 | 29 | 13 | 0.014 | 0.014 | 0.037 | 0.050 | -2.7917 | 5.24E-03 | 0.0539 | Increasing |
| LIRI-JP::SP107084 | Liver | LIRI-JP | 635 | 0 | 93 | 34 | 11 | 0.010 | 0.028 | 0.035 | 0.052 | -2.7623 | 5.74E-03 | 0.0559 | Increasing |
| LIRI-JP::SP107111 | Liver | LIRI-JP | 671 | 0 | 76 | 34 | 12.5 | 0.013 | 0.025 | 0.032 | 0.058 | -2.7623 | 5.74E-03 | 0.0559 | Increasing |
| LIRI-JP::SP107141 | Liver | LIRI-JP | 256 | 0 | 41 | 23 | 12.5 | 0.013 | 0.013 | 0.013 | 0.058 | -2.7708 | 5.59E-03 | 0.0559 | Increasing |
| RECA-EU::SP102759 | Kidney | RECA-EU | 1304 | 0 | 209 | 94 | 9.5 | 0.040 | 0.080 | 0.068 | 0.120 | -2.7739 | 5.54E-03 | 0.0559 | Increasing |
| MALY-DE::SP116690 | Blood | MALY-DE | 1862 | 0 | 0 | 32 | 12 | 0.000 | 0.020 | 0.026 | 0.042 | -2.7744 | 5.53E-03 | 0.0559 | Increasing |

| | | | | | | | | | | | | | | | |
|---|---|---|---|---|---|---|---|---|---|---|---|---|---|---|---|
| BTCA-SG::SP117556 | Biliary | BTCA-SG | 802 | 0 | 0 | 24 | 16.5 | 0.012 | 0.013 | 0.028 | 0.050 | -2.7643 | 5.70E-03 | 0.0559 | Increasing |
| LIRI-JP::SP112241 | Liver | LIRI-JP | 555 | 0 | 98 | 32 | 14 | 0.016 | 0.015 | 0.038 | 0.052 | -2.7726 | 5.56E-03 | 0.0559 | Increasing |
| LIRI-JP::SP99029 | Liver | LIRI-JP | 462 | 0 | 49 | 18 | 10 | 0.000 | 0.018 | 0.016 | 0.035 | -2.7663 | 5.67E-03 | 0.0559 | Increasing |
| OV-US::SP59950 | Ovary | OV-US | 1160 | 352 | 186 | 123 | 13 | 0.137 | 0.114 | 0.082 | 0.064 | 2.7548 | 5.87E-03 | 0.0568 | Decreasing |
| LIRI-JP::SP99019 | Liver | LIRI-JP | 642 | 0 | 89 | 38 | 11 | 0.019 | 0.015 | 0.057 | 0.046 | -2.7498 | 5.96E-03 | 0.0573 | Increasing |
| LIRI-JP::SP50135 | Liver | LIRI-JP | 461 | 0 | 91 | 40 | 10 | 0.013 | 0.023 | 0.067 | 0.035 | -2.7470 | 6.01E-03 | 0.0574 | Increasing |
| LIRI-JP::SP99189 | Liver | LIRI-JP | 241 | 0 | 36 | 12 | 16 | 0.003 | 0.005 | 0.013 | 0.029 | -2.7360 | 6.22E-03 | 0.0589 | Increasing |
| LIRI-JP::SP98909 | Liver | LIRI-JP | 860 | 0 | 133 | 55 | 14 | 0.022 | 0.045 | 0.054 | 0.075 | -2.7028 | 6.88E-03 | 0.0647 | Increasing |
| KIRC-US::SP37432 | Kidney | KIRC-US | 477 | 0 | 73 | 32 | 9 | 0.007 | 0.021 | 0.043 | 0.039 | -2.6857 | 7.24E-03 | 0.0677 | Increasing |
| OV-AU::SP101666 | Ovary | OV-AU | 520 | 161 | 168 | 90 | 21 | 0.111 | 0.061 | 0.054 | 0.052 | 2.6829 | 7.30E-03 | 0.0678 | Decreasing |
| LIRI-JP::SP50185 | Liver | LIRI-JP | 529 | 0 | 97 | 35 | 13 | 0.022 | 0.018 | 0.029 | 0.070 | -2.6744 | 7.49E-03 | 0.0691 | Increasing |
| BRCA-US::SP6825 | Breast | BRCA-US | 985 | 216 | 0 | 51 | 13 | 0.034 | 0.021 | 0.060 | 0.074 | -2.6686 | 7.62E-03 | 0.0699 | Increasing |
| RECA-EU::SP103408 | Kidney | RECA-EU | 712 | 0 | 65 | 28 | 9 | 0.010 | 0.016 | 0.026 | 0.045 | -2.6641 | 7.72E-03 | 0.0704 | Increasing |
| LINC-JP::SP98851 | Liver | LIHC-US | 122 | 0 | 22 | 11 | 16 | 0.003 | 0.008 | 0.003 | 0.036 | -2.6608 | 7.80E-03 | 0.0706 | Increasing |
| LUAD-US::SP56303 | Lung | LUAD-US | 807 | 114 | 181 | 96 | 12 | 0.054 | 0.079 | 0.088 | 0.125 | -2.6558 | 7.91E-03 | 0.0710 | Increasing |
| PACA-AU::SP108123 | Pancreas | PACA-AU | 355 | 0 | 68 | 13 | 12 | 0.004 | 0.003 | 0.012 | 0.023 | -2.6552 | 7.93E-03 | 0.0710 | Increasing |
| LIHC-US::SP49531 | Liver | LIHC-US | 1105 | 0 | 0 | 48 | 12 | 0.032 | 0.029 | 0.039 | 0.090 | -2.6526 | 7.99E-03 | 0.0711 | Increasing |
| PRAD-UK::SP114991 | Prostate | PRAD-US | 1337 | 477 | 557 | 250 | 17 | 0.276 | 0.216 | 0.179 | 0.176 | 2.6448 | 8.17E-03 | 0.0719 | Decreasing |
| LIRI-JP::SP107032 | Liver | LIRI-JP | 94 | 0 | 28 | 11 | 11.5 | 0.003 | 0.005 | 0.010 | 0.029 | -2.6444 | 8.18E-03 | 0.0719 | Increasing |
| PRAD-UK::SP114996 | Prostate | PRAD-US | 1359 | 488 | 566 | 253 | 17 | 0.276 | 0.221 | 0.183 | 0.176 | 2.6396 | 8.30E-03 | 0.0725 | Decreasing |
| MALY-DE::SP116612 | Blood | MALY-DE | 907 | 0 | 0 | 22 | 10 | 0.006 | 0.011 | 0.012 | 0.036 | -2.6357 | 8.40E-03 | 0.0728 | Increasing |
| RECA-EU::SP102741 | Kidney | RECA-EU | 724 | 0 | 136 | 44 | 9.5 | 0.020 | 0.028 | 0.037 | 0.065 | -2.6337 | 8.44E-03 | 0.0728 | Increasing |
| OV-AU::SP101634 | Ovary | OV-AU | 587 | 355 | 232 | 144 | 18 | 0.159 | 0.100 | 0.108 | 0.072 | 2.6300 | 8.54E-03 | 0.0728 | Decreasing |
| BRCA-UK::SP2144 | Breast | BRCA-US | 381 | 165 | 190 | 85 | 17 | 0.048 | 0.063 | 0.082 | 0.111 | -2.6299 | 8.54E-03 | 0.0728 | Increasing |
| LIRI-JP::SP107041 | Liver | LIRI-JP | 392 | 0 | 52 | 19 | 10 | 0.000 | 0.015 | 0.029 | 0.023 | -2.6272 | 8.61E-03 | 0.0730 | Increasing |
| PACA-AU::SP71084 | Pancreas | PACA-AU | 468 | 0 | 0 | 11 | 10.5 | 0.000 | 0.008 | 0.003 | 0.023 | -2.6160 | 8.90E-03 | 0.0750 | Increasing |
| PACA-CA::SP125771 | Pancreas | PACA-CA | 1436 | 0 | 0 | 24 | 10 | 0.011 | 0.008 | 0.017 | 0.038 | -2.6086 | 9.09E-03 | 0.0762 | Increasing |
| STAD-US::SP85287 | Stomach | STAD-US | 1053 | 0 | 0 | 23 | 11 | 0.012 | 0.011 | 0.020 | 0.048 | -2.6040 | 9.22E-03 | 0.0768 | Increasing |
| LIRI-JP::SP50123 | Liver | LIRI-JP | 860 | 0 | 0 | 44 | 12 | 0.022 | 0.025 | 0.054 | 0.058 | -2.5969 | 9.41E-03 | 0.0779 | Increasing |
| LIRI-JP::SP112224 | Liver | LIRI-JP | 576 | 0 | 100 | 38 | 12 | 0.019 | 0.025 | 0.035 | 0.064 | -2.5889 | 9.63E-03 | 0.0789 | Increasing |
| MELA-AU::SP124341 | Skin | SKCM-US | 365 | 0 | 0 | 12 | 13 | 0.007 | 0.003 | 0.009 | 0.033 | -2.5887 | 9.63E-03 | 0.0789 | Increasing |
| UCEC-US::SP92268 | Endometrium | UCEC-US | 446 | 0 | 0 | 16 | 11 | 0.006 | 0.008 | 0.017 | 0.035 | -2.5858 | 9.72E-03 | 0.0791 | Increasing |
| PRAD-CA::SP112817 | Prostate | PRAD-US | 1107 | 0 | 0 | 11 | 8 | 0.000 | 0.003 | 0.021 | 0.015 | -2.5836 | 9.78E-03 | 0.0792 | Increasing |
| PRAD-UK::SP114903 | Prostate | PRAD-US | 1042 | 0 | 0 | 15 | 10.5 | 0.000 | 0.011 | 0.018 | 0.025 | -2.5783 | 9.93E-03 | 0.0800 | Increasing |
| LIRI-JP::SP99045 | Liver | LIRI-JP | 835 | 0 | 138 | 55 | 12 | 0.029 | 0.040 | 0.051 | 0.081 | -2.5690 | 1.02E-02 | 0.0817 | Increasing |
| LIRI-JP::SP112130 | Liver | LIRI-JP | 498 | 0 | 56 | 23 | 14.5 | 0.019 | 0.005 | 0.016 | 0.058 | -2.5639 | 1.03E-02 | 0.0825 | Increasing |
| PACA-CA::SP125805 | Pancreas | PACA-CA | 438 | 0 | 0 | 14 | 9 | 0.000 | 0.006 | 0.017 | 0.019 | -2.5546 | 1.06E-02 | 0.0842 | Increasing |
| PACA-CA::SP117568 | Pancreas | PACA-CA | 796 | 0 | 0 | 22 | 13 | 0.011 | 0.008 | 0.011 | 0.038 | -2.5474 | 1.09E-02 | 0.0855 | Increasing |
| LIRI-JP::SP98927 | Liver | LIRI-JP | 689 | 0 | 110 | 49 | 14 | 0.022 | 0.043 | 0.035 | 0.081 | -2.5408 | 1.11E-02 | 0.0858 | Increasing |
| LIHC-US::SP49379 | Liver | LIHC-US | 657 | 0 | 0 | 35 | 10 | 0.016 | 0.024 | 0.039 | 0.054 | -2.5435 | 1.10E-02 | 0.0858 | Increasing |
| LIHC-US::SP49469 | Liver | LIHC-US | 686 | 0 | 136 | 63 | 13 | 0.025 | 0.055 | 0.071 | 0.072 | -2.5407 | 1.11E-02 | 0.0858 | Increasing |
| HNSC-US::SP30213 | Head&Neck | HNSC-US | 325 | 0 | 55 | 18 | 9 | 0.009 | 0.008 | 0.016 | 0.040 | -2.5381 | 1.11E-02 | 0.0860 | Increasing |
| LICA-FR::SP97747 | Liver | LICA-FR | 587 | 0 | 113 | 51 | 14.5 | 0.026 | 0.028 | 0.050 | 0.065 | -2.5330 | 1.13E-02 | 0.0868 | Increasing |
| LIRI-JP::SP107060 | Liver | LIRI-JP | 596 | 0 | 94 | 27 | 10 | 0.003 | 0.025 | 0.032 | 0.035 | -2.5289 | 1.14E-02 | 0.0869 | Increasing |
| LIRI-JP::SP112141 | Liver | LIRI-JP | 849 | 90 | 128 | 69 | 14 | 0.035 | 0.058 | 0.057 | 0.098 | -2.5300 | 1.14E-02 | 0.0869 | Increasing |
| PAEN-AU::SP106808 | Pancreas | PAEN-AU | 156 | 0 | 28 | 13 | 10 | 0.000 | 0.005 | 0.011 | 0.021 | -2.5231 | 1.16E-02 | 0.0875 | Increasing |
| LIHC-US::SP49119 | Liver | LIHC-US | 624 | 0 | 68 | 20 | 13 | 0.003 | 0.018 | 0.019 | 0.036 | -2.5227 | 1.16E-02 | 0.0875 | Increasing |
| PAEN-AU::SP102547 | Pancreas | PAEN-AU | 137 | 0 | 25 | 14 | 13 | 0.000 | 0.005 | 0.013 | 0.021 | -2.5180 | 1.18E-02 | 0.0882 | Increasing |
| OV-US::SP64036 | Ovary | OV-US | 793 | 297 | 407 | 178 | 20 | 0.199 | 0.142 | 0.142 | 0.104 | 2.5151 | 1.19E-02 | 0.0885 | Decreasing |
| OV-AU::SP101610 | Ovary | OV-AU | 577 | 254 | 296 | 127 | 20 | 0.124 | 0.109 | 0.087 | 0.057 | 2.5006 | 1.24E-02 | 0.0917 | Decreasing |

| | | | | | | | | | | | | | | | |
|---|---|---|---|---|---|---|---|---|---|---|---|---|---|---|---|
| UCEC-US::SP95406 | Endometrium | UCEC-US | 307 | 0 | 54 | 15 | 12 | 0.000 | 0.010 | 0.027 | 0.017 | -2.4878 | 1.29E-02 | 0.0946 | Increasing |
| LIHC-US::SP49157 | Liver | LIHC-US | 716 | 0 | 81 | 34 | 11 | 0.019 | 0.018 | 0.039 | 0.054 | -2.4585 | 1.40E-02 | 0.1022 | Increasing |
| LUSC-US::SP59245 | Lung | LUSC-US | 2056 | 363 | 243 | 97 | 10 | 0.134 | 0.058 | 0.065 | 0.074 | 2.4485 | 1.43E-02 | 0.1030 | Decreasing |
| RECA-EU::SP102718 | Kidney | RECA-EU | 780 | 0 | 135 | 52 | 9 | 0.023 | 0.035 | 0.048 | 0.065 | -2.4500 | 1.43E-02 | 0.1030 | Increasing |
| BTCA-SG::SP117930 | Biliary | BTCA-SG | 1682 | 0 | 0 | 31 | 11 | 0.015 | 0.027 | 0.032 | 0.056 | -2.4521 | 1.42E-02 | 0.1030 | Increasing |
| KIRC-US::SP36218 | Kidney | KIRC-US | 319 | 0 | 47 | 20 | 9.5 | 0.007 | 0.013 | 0.015 | 0.039 | -2.4502 | 1.43E-02 | 0.1030 | Increasing |
| PRAD-UK::SP111127 | Prostate | PRAD-US | 377 | 0 | 0 | 13 | 10 | 0.000 | 0.013 | 0.006 | 0.030 | -2.4366 | 1.48E-02 | 0.1059 | Increasing |
| LIRI-JP::SP107052 | Liver | LIRI-JP | 464 | 0 | 53 | 28 | 14 | 0.010 | 0.023 | 0.025 | 0.046 | -2.4295 | 1.51E-02 | 0.1070 | Increasing |
| BRCA-US::SP7291 | Breast | BRCA-US | 321 | 0 | 91 | 11 | 11 | 0.003 | 0.005 | 0.009 | 0.026 | -2.4275 | 1.52E-02 | 0.1070 | Increasing |
| LUAD-US::SP50592 | Lung | LUAD-US | 343 | 0 | 49 | 23 | 12 | 0.006 | 0.022 | 0.016 | 0.043 | -2.4289 | 1.51E-02 | 0.1070 | Increasing |
| LIRI-JP::SP98965 | Liver | LIRI-JP | 719 | 0 | 80 | 32 | 13.5 | 0.013 | 0.023 | 0.035 | 0.046 | -2.4219 | 1.54E-02 | 0.1071 | Increasing |
| LIRI-JP::SP98985 | Liver | LIRI-JP | 769 | 0 | 84 | 34 | 12.5 | 0.016 | 0.023 | 0.035 | 0.052 | -2.4220 | 1.54E-02 | 0.1071 | Increasing |
| LIRI-JP::SP99109 | Liver | LIRI-JP | 824 | 0 | 99 | 36 | 11 | 0.029 | 0.015 | 0.025 | 0.075 | -2.4241 | 1.53E-02 | 0.1071 | Increasing |
| PACA-AU::SP70202 | Pancreas | PACA-AU | 868 | 423 | 189 | 156 | 14 | 0.139 | 0.152 | 0.108 | 0.084 | 2.4119 | 1.59E-02 | 0.1096 | Decreasing |
| KIRC-US::SP34452 | Kidney | KIRC-US | 1304 | 0 | 219 | 76 | 9 | 0.044 | 0.051 | 0.083 | 0.088 | -2.4056 | 1.61E-02 | 0.1109 | Increasing |
| STAD-US::SP105018 | Stomach | STAD-US | 4714 | 3163 | 888 | 1018 | 17 | 0.969 | 0.869 | 0.788 | 0.807 | 2.3933 | 1.67E-02 | 0.1142 | Decreasing |
| LIRI-JP::SP98915 | Liver | LIRI-JP | 426 | 0 | 0 | 14 | 12 | 0.003 | 0.010 | 0.013 | 0.029 | -2.3808 | 1.73E-02 | 0.1176 | Increasing |
| LIHC-US::SP49334 | Liver | LIHC-US | 487 | 0 | 70 | 31 | 11 | 0.006 | 0.026 | 0.045 | 0.030 | -2.3684 | 1.79E-02 | 0.1198 | Increasing |
| RECA-EU::SP103507 | Kidney | RECA-EU | 824 | 0 | 76 | 38 | 9 | 0.007 | 0.031 | 0.045 | 0.035 | -2.3670 | 1.79E-02 | 0.1198 | Increasing |
| LINC-JP::SP98883 | Liver | LIHC-US | 744 | 0 | 89 | 31 | 13 | 0.009 | 0.026 | 0.035 | 0.042 | -2.3684 | 1.79E-02 | 0.1198 | Increasing |
| LIRI-JP::SP99149 | Liver | LIRI-JP | 607 | 0 | 91 | 43 | 15 | 0.016 | 0.038 | 0.041 | 0.058 | -2.3678 | 1.79E-02 | 0.1198 | Increasing |
| HNSC-US::SP31174 | Head&Neck | HNSC-US | 495 | 0 | 76 | 27 | 14.5 | 0.006 | 0.025 | 0.033 | 0.035 | -2.3595 | 1.83E-02 | 0.1213 | Increasing |
| MELA-AU::SP124357 | Skin | SKCM-US | 316 | 0 | 53 | 18 | 11 | 0.007 | 0.010 | 0.019 | 0.033 | -2.3551 | 1.85E-02 | 0.1213 | Increasing |
| LIRI-JP::SP98981 | Liver | LIRI-JP | 486 | 0 | 75 | 23 | 10.5 | 0.006 | 0.020 | 0.019 | 0.041 | -2.3571 | 1.84E-02 | 0.1213 | Increasing |
| LINC-JP::SP98873 | Liver | LIHC-US | 758 | 0 | 122 | 55 | 13 | 0.032 | 0.039 | 0.055 | 0.078 | -2.3506 | 1.87E-02 | 0.1213 | Increasing |
| LIRI-JP::SP107010 | Liver | LIRI-JP | 634 | 0 | 103 | 39 | 10 | 0.022 | 0.030 | 0.022 | 0.075 | -2.3510 | 1.87E-02 | 0.1213 | Increasing |
| LUAD-US::SP52667 | Lung | LUAD-US | 1182 | 0 | 142 | 32 | 12 | 0.050 | 0.022 | 0.013 | 0.022 | 2.3524 | 1.87E-02 | 0.1213 | Decreasing |
| LIRI-JP::SP50171 | Liver | LIRI-JP | 724 | 0 | 104 | 39 | 12.5 | 0.010 | 0.038 | 0.041 | 0.046 | -2.3510 | 1.87E-02 | 0.1213 | Increasing |
| PACA-CA::SP125723 | Pancreas | PACA-CA | 485 | 0 | 128 | 22 | 9 | 0.008 | 0.014 | 0.011 | 0.035 | -2.3480 | 1.89E-02 | 0.1216 | Increasing |
| STAD-US::SP84086 | Stomach | STAD-US | 647 | 0 | 0 | 31 | 16 | 0.015 | 0.017 | 0.039 | 0.042 | -2.3408 | 1.92E-02 | 0.1235 | Increasing |
| LIRI-JP::SP112228 | Liver | LIRI-JP | 668 | 0 | 121 | 59 | 12 | 0.029 | 0.045 | 0.064 | 0.070 | -2.3320 | 1.97E-02 | 0.1242 | Increasing |
| LIRI-JP::SP112234 | Liver | LIRI-JP | 936 | 0 | 130 | 59 | 16 | 0.035 | 0.045 | 0.045 | 0.093 | -2.3320 | 1.97E-02 | 0.1242 | Increasing |
| PACA-CA::SP125768 | Pancreas | PACA-CA | 1467 | 0 | 0 | 15 | 12 | 0.000 | 0.011 | 0.011 | 0.022 | -2.3339 | 1.96E-02 | 0.1242 | Increasing |
| MELA-AU::SP124441 | Skin | SKCM-US | 551 | 0 | 64 | 25 | 11 | 0.007 | 0.016 | 0.038 | 0.028 | -2.3362 | 1.95E-02 | 0.1242 | Increasing |
| BTCA-SG::SP117031 | Biliary | BTCA-SG | 965 | 0 | 0 | 19 | 13 | 0.006 | 0.013 | 0.032 | 0.028 | -2.3261 | 2.00E-02 | 0.1256 | Increasing |
| LIHC-US::SP106743 | Liver | LIHC-US | 268 | 23 | 30 | 18 | 10.5 | 0.006 | 0.010 | 0.023 | 0.030 | -2.3234 | 2.02E-02 | 0.1260 | Increasing |
| LINC-JP::SP98893 | Liver | LIHC-US | 900 | 0 | 0 | 40 | 13 | 0.032 | 0.018 | 0.032 | 0.078 | -2.3172 | 2.05E-02 | 0.1275 | Increasing |
| PRAD-UK::SP114901 | Prostate | PRAD-US | 989 | 0 | 0 | 18 | 10 | 0.000 | 0.013 | 0.027 | 0.020 | -2.3142 | 2.07E-02 | 0.1280 | Increasing |
| LINC-JP::SP98877 | Liver | LIHC-US | 528 | 0 | 80 | 38 | 14 | 0.025 | 0.024 | 0.029 | 0.072 | -2.3067 | 2.11E-02 | 0.1290 | Increasing |
| PACA-CA::SP117760 | Pancreas | PACA-CA | 407 | 0 | 135 | 14 | 15 | 0.000 | 0.011 | 0.008 | 0.022 | -2.3047 | 2.12E-02 | 0.1290 | Increasing |
| PACA-CA::SP125811 | Pancreas | PACA-CA | 374 | 0 | 95 | 14 | 10.5 | 0.000 | 0.008 | 0.014 | 0.019 | -2.3047 | 2.12E-02 | 0.1290 | Increasing |
| GBM-US::SP24391 | Brain | GBM-US | 429 | 0 | 0 | 12 | 6 | 0.000 | 0.007 | 0.015 | 0.020 | -2.3082 | 2.10E-02 | 0.1290 | Increasing |
| OV-AU::SP101891 | Ovary | OV-AU | 1126 | 498 | 402 | 251 | 19 | 0.245 | 0.193 | 0.174 | 0.160 | 2.3031 | 2.13E-02 | 0.1291 | Decreasing |
| COAD-US::SP22750 | CRC | COAD-US | 1244 | 0 | 0 | 18 | 9 | 0.015 | 0.003 | 0.013 | 0.045 | -2.2898 | 2.20E-02 | 0.1331 | Increasing |
| PRAD-CA::SP112987 | Prostate | PRAD-US | 331 | 0 | 0 | 11 | 9.5 | 0.004 | 0.003 | 0.015 | 0.020 | -2.2870 | 2.22E-02 | 0.1335 | Increasing |
| PRAD-UK::SP114987 | Prostate | PRAD-US | 1410 | 525 | 581 | 266 | 17 | 0.291 | 0.224 | 0.189 | 0.206 | 2.2769 | 2.28E-02 | 0.1350 | Decreasing |
| BLCA-US::SP1114 | Bladder | BLCA-US | 583 | 0 | 0 | 14 | 12 | 0.021 | 0.013 | 0.007 | 0.000 | 2.2799 | 2.26E-02 | 0.1350 | Decreasing |
| UCEC-US::SP91746 | Endometrium | UCEC-US | 327 | 54 | 0 | 13 | 11 | 0.000 | 0.010 | 0.020 | 0.017 | -2.2796 | 2.26E-02 | 0.1350 | Increasing |
| LIRI-JP::SP50121 | Liver | LIRI-JP | 331 | 0 | 55 | 20 | 11 | 0.010 | 0.005 | 0.035 | 0.023 | -2.2751 | 2.29E-02 | 0.1350 | Increasing |
| LIRI-JP::SP50165 | Liver | LIRI-JP | 612 | 0 | 101 | 40 | 11 | 0.022 | 0.025 | 0.041 | 0.058 | -2.2764 | 2.28E-02 | 0.1350 | Increasing |

| | | | | | | | | | | | | | | | |
|---|---|---|---|---|---|---|---|---|---|---|---|---|---|---|---|
| LIRI-JP::SP98949 | Liver | LIRI-JP | 930 | 0 | 97 | 38 | 12 | 0.016 | 0.033 | 0.032 | 0.058 | -2.2670 | 2.34E-02 | 0.1373 | Increasing |
| UCEC-US::SP95126 | Endometrium | UCEC-US | 817 | 131 | 0 | 32 | 10 | 0.019 | 0.021 | 0.027 | 0.058 | -2.2641 | 2.36E-02 | 0.1374 | Increasing |
| PACA-CA::SP125706 | Pancreas | PACA-CA | 1429 | 658 | 329 | 255 | 13 | 0.271 | 0.193 | 0.149 | 0.194 | 2.2636 | 2.36E-02 | 0.1374 | Decreasing |
| PACA-AU::SP71404 | Pancreas | PACA-AU | 1305 | 0 | 0 | 21 | 10 | 0.007 | 0.008 | 0.023 | 0.027 | -2.2597 | 2.38E-02 | 0.1383 | Increasing |
| PRAD-UK::SP114957 | Prostate | PRAD-US | 1261 | 0 | 0 | 47 | 10 | 0.019 | 0.034 | 0.058 | 0.050 | -2.2486 | 2.45E-02 | 0.1418 | Increasing |
| RECA-EU::SP103428 | Kidney | RECA-EU | 1125 | 0 | 163 | 63 | 8 | 0.049 | 0.026 | 0.054 | 0.090 | -2.2443 | 2.48E-02 | 0.1428 | Increasing |
| LIRI-JP::SP107023 | Liver | LIRI-JP | 461 | 0 | 74 | 28 | 13 | 0.010 | 0.023 | 0.029 | 0.041 | -2.2420 | 2.50E-02 | 0.1430 | Increasing |
| BRCA-US::SP5017 | Breast | BRCA-US | 536 | 0 | 0 | 15 | 13 | 0.007 | 0.005 | 0.019 | 0.026 | -2.2364 | 2.53E-02 | 0.1446 | Increasing |
| LINC-JP::SP98839 | Liver | LIHC-US | 538 | 0 | 92 | 39 | 13 | 0.019 | 0.016 | 0.074 | 0.024 | -2.2327 | 2.56E-02 | 0.1454 | Increasing |
| COAD-US::SP19215 | CRC | COAD-US | 75239 | 0 | 0 | 59 | 9 | 0.040 | 0.041 | 0.050 | 0.091 | -2.2269 | 2.60E-02 | 0.1465 | Increasing |
| LUAD-US::SP50713 | Lung | LUAD-US | 442 | 0 | 63 | 23 | 11 | 0.016 | 0.011 | 0.016 | 0.049 | -2.2267 | 2.60E-02 | 0.1465 | Increasing |
| STAD-US::SP84998 | Stomach | STAD-US | 923 | 0 | 0 | 11 | 7.5 | 0.003 | 0.006 | 0.013 | 0.021 | -2.2162 | 2.67E-02 | 0.1482 | Increasing |
| HNSC-US::SP31190 | Head&Neck | HNSC-US | 710 | 0 | 70 | 37 | 12 | 0.021 | 0.025 | 0.036 | 0.058 | -2.2170 | 2.66E-02 | 0.1482 | Increasing |
| KIRP-US::SP97269 | Kidney | KIRP-US | 1303 | 0 | 199 | 82 | 9 | 0.056 | 0.059 | 0.069 | 0.113 | -2.2184 | 2.65E-02 | 0.1482 | Increasing |
| LINC-JP::SP98843 | Liver | LIHC-US | 431 | 0 | 63 | 17 | 12.5 | 0.009 | 0.008 | 0.016 | 0.036 | -2.2180 | 2.66E-02 | 0.1482 | Increasing |
| CESC-US::SP107575 | Cervix | CESC-US | 316 | 68 | 0 | 11 | 14.5 | 0.020 | 0.006 | 0.007 | 0.000 | 2.2076 | 2.73E-02 | 0.1509 | Decreasing |
| LIRI-JP::SP112268 | Liver | LIRI-JP | 778 | 0 | 133 | 59 | 12 | 0.029 | 0.048 | 0.060 | 0.070 | -2.2029 | 2.76E-02 | 0.1516 | Increasing |
| PACA-CA::SP125699 | Pancreas | PACA-CA | 638 | 0 | 0 | 18 | 14 | 0.004 | 0.014 | 0.008 | 0.029 | -2.2040 | 2.75E-02 | 0.1516 | Increasing |
| PRAD-UK::SP114953 | Prostate | PRAD-US | 1319 | 0 | 160 | 55 | 11 | 0.022 | 0.045 | 0.061 | 0.060 | -2.1954 | 2.81E-02 | 0.1540 | Increasing |
| LIRI-JP::SP99085 | Liver | LIRI-JP | 802 | 0 | 128 | 39 | 14 | 0.019 | 0.035 | 0.022 | 0.070 | -2.1922 | 2.84E-02 | 0.1541 | Increasing |
| LIRI-JP::SP99173 | Liver | LIRI-JP | 572 | 0 | 103 | 39 | 12 | 0.025 | 0.020 | 0.041 | 0.058 | -2.1922 | 2.84E-02 | 0.1541 | Increasing |
| RECA-EU::SP102827 | Kidney | RECA-EU | 1379 | 0 | 214 | 102 | 9 | 0.056 | 0.075 | 0.088 | 0.110 | -2.1907 | 2.85E-02 | 0.1541 | Increasing |
| LIHC-US::SP49328 | Liver | LIHC-US | 573 | 0 | 87 | 29 | 11 | 0.009 | 0.024 | 0.035 | 0.036 | -2.1842 | 2.90E-02 | 0.1545 | Increasing |
| HNSC-US::SP33544 | Head&Neck | HNSC-US | 1206 | 137 | 147 | 75 | 14 | 0.048 | 0.063 | 0.052 | 0.115 | -2.1847 | 2.89E-02 | 0.1545 | Increasing |
| KIRP-US::SP97124 | Kidney | KIRP-US | 1658 | 0 | 309 | 108 | 9 | 0.070 | 0.072 | 0.125 | 0.108 | -2.1838 | 2.90E-02 | 0.1545 | Increasing |
| PRAD-UK::SP111123 | Prostate | PRAD-US | 269 | 0 | 55 | 22 | 11.5 | 0.011 | 0.011 | 0.024 | 0.035 | -2.1855 | 2.89E-02 | 0.1545 | Increasing |
| RECA-EU::SP103197 | Kidney | RECA-EU | 883 | 0 | 158 | 56 | 10 | 0.033 | 0.038 | 0.040 | 0.080 | -2.1749 | 2.96E-02 | 0.1568 | Increasing |
| GBM-US::SP24565 | Brain | GBM-US | 239 | 0 | 41 | 14 | 6 | 0.005 | 0.007 | 0.015 | 0.026 | -2.1736 | 2.97E-02 | 0.1568 | Increasing |
| LIHC-US::SP98265 | Liver | LIHC-US | 857 | 0 | 135 | 42 | 15.5 | 0.019 | 0.031 | 0.052 | 0.048 | -2.1762 | 2.95E-02 | 0.1568 | Increasing |
| PRAD-CA::SP112815 | Prostate | PRAD-US | 354 | 0 | 57 | 16 | 12 | 0.007 | 0.011 | 0.009 | 0.035 | -2.1545 | 3.12E-02 | 0.1627 | Increasing |
| PRAD-UK::SP114934 | Prostate | PRAD-US | 507 | 0 | 0 | 16 | 12 | 0.004 | 0.011 | 0.018 | 0.025 | -2.1545 | 3.12E-02 | 0.1627 | Increasing |
| LIRI-JP::SP99209 | Liver | LIRI-JP | 538 | 0 | 92 | 31 | 13 | 0.016 | 0.023 | 0.025 | 0.052 | -2.1555 | 3.11E-02 | 0.1627 | Increasing |
| OV-AU::SP101552 | Ovary | OV-AU | 1440 | 690 | 511 | 336 | 16 | 0.347 | 0.234 | 0.216 | 0.269 | 2.1518 | 3.14E-02 | 0.1632 | Decreasing |
| PACA-AU::SP107786 | Pancreas | PACA-AU | 1042 | 0 | 341 | 54 | 9 | 0.026 | 0.038 | 0.041 | 0.063 | -2.1505 | 3.15E-02 | 0.1632 | Increasing |
| KIRP-US::SP97161 | Kidney | KIRP-US | 1287 | 0 | 185 | 73 | 10 | 0.038 | 0.062 | 0.069 | 0.088 | -2.1488 | 3.16E-02 | 0.1633 | Increasing |
| LIRI-JP::SP99117 | Liver | LIRI-JP | 586 | 0 | 91 | 25 | 11 | 0.010 | 0.018 | 0.029 | 0.035 | -2.1469 | 3.18E-02 | 0.1635 | Increasing |
| HNSC-US::SP31422 | Head&Neck | HNSC-US | 806 | 0 | 144 | 54 | 13 | 0.036 | 0.038 | 0.042 | 0.086 | -2.1451 | 3.19E-02 | 0.1637 | Increasing |
| MALY-DE::SP59368 | Blood | MALY-DE | 1462 | 0 | 0 | 35 | 11.5 | 0.006 | 0.029 | 0.014 | 0.048 | -2.1413 | 3.22E-02 | 0.1646 | Increasing |
| SKCM-US::SP82532 | Skin | SKCM-US | 1008 | 0 | 179 | 68 | 13 | 0.038 | 0.057 | 0.060 | 0.089 | -2.1271 | 3.34E-02 | 0.1699 | Increasing |
| OV-US::SP68348 | Ovary | OV-US | 1110 | 456 | 266 | 172 | 16 | 0.203 | 0.119 | 0.138 | 0.122 | 2.1246 | 3.36E-02 | 0.1699 | Decreasing |
| OV-US::SP63966 | Ovary | OV-US | 526 | 0 | 86 | 23 | 10 | 0.013 | 0.015 | 0.016 | 0.046 | -2.1254 | 3.36E-02 | 0.1699 | Increasing |
| BRCA-US::SP5052 | Breast | BRCA-US | 1519 | 585 | 743 | 325 | 16 | 0.351 | 0.253 | 0.249 | 0.254 | 2.1158 | 3.44E-02 | 0.1707 | Decreasing |
| LUSC-US::SP56541 | Lung | LUSC-US | 1483 | 0 | 254 | 59 | 15 | 0.038 | 0.045 | 0.046 | 0.091 | -2.1178 | 3.42E-02 | 0.1707 | Increasing |
| READ-US::SP80271 | CRC | READ-US | 1406 | 0 | 0 | 16 | 7 | 0.009 | 0.011 | 0.007 | 0.040 | -2.1166 | 3.43E-02 | 0.1707 | Increasing |
| LIRI-JP::SP98967 | Liver | LIRI-JP | 405 | 0 | 44 | 14 | 9.5 | 0.010 | 0.005 | 0.010 | 0.035 | -2.1156 | 3.44E-02 | 0.1707 | Increasing |
| BRCA-UK::SP116369 | Breast | BRCA-US | 1349 | 780 | 333 | 254 | 13 | 0.265 | 0.222 | 0.176 | 0.196 | 2.1198 | 3.40E-02 | 0.1707 | Decreasing |
| RECA-EU::SP103156 | Kidney | RECA-EU | 901 | 0 | 118 | 59 | 9 | 0.026 | 0.047 | 0.048 | 0.070 | -2.1123 | 3.47E-02 | 0.1714 | Increasing |
| LUAD-US::SP50518 | Lung | LUAD-US | 403 | 0 | 74 | 26 | 10 | 0.016 | 0.014 | 0.026 | 0.043 | -2.1111 | 3.48E-02 | 0.1714 | Increasing |
| LIRI-JP::SP98945 | Liver | LIRI-JP | 335 | 31 | 67 | 38 | 16 | 0.019 | 0.023 | 0.051 | 0.041 | -2.1061 | 3.52E-02 | 0.1719 | Increasing |
| MALY-DE::SP116718 | Blood | MALY-DE | 587 | 0 | 0 | 17 | 13 | 0.006 | 0.011 | 0.003 | 0.030 | -2.1061 | 3.52E-02 | 0.1719 | Increasing |

| | | | | | | | | | | | | | | |
|---|---|---|---|---|---|---|---|---|---|---|---|---|---|---|
| COAD-US::SP96110 | CRC | COAD-US | 1270 | 0 | 0 | 14 | 10 | 0.012 | 0.000 | 0.013 | 0.034 | -2.1060 | 3.52E-02 | 0.1719 | Increasing |
| OV-AU::SP101622 | Ovary | OV-AU | 948 | 482 | 350 | 230 | 21 | 0.220 | 0.186 | 0.144 | 0.160 | 2.0997 | 3.58E-02 | 0.1740 | Decreasing |
| PRAD-UK::SP114958 | Prostate | PRAD-US | 1319 | 0 | 0 | 49 | 10 | 0.019 | 0.040 | 0.058 | 0.050 | -2.0926 | 3.64E-02 | 0.1741 | Increasing |
| PRAD-UK::SP114994 | Prostate | PRAD-US | 1377 | 513 | 571 | 262 | 17 | 0.276 | 0.227 | 0.189 | 0.201 | 2.0778 | 3.77E-02 | 0.1741 | Decreasing |
| OV-US::SP62616 | Ovary | OV-US | 1442 | 966 | 315 | 329 | 17.5 | 0.330 | 0.290 | 0.227 | 0.261 | 2.0694 | 3.85E-02 | 0.1741 | Decreasing |
| LIRI-JP::SP107088 | Liver | LIRI-JP | 262 | 0 | 55 | 18 | 13 | 0.006 | 0.010 | 0.025 | 0.023 | -2.0649 | 3.89E-02 | 0.1741 | Increasing |
| PACA-AU::SP76955 | Pancreas | PACA-AU | 543 | 0 | 0 | 12 | 18 | 0.004 | 0.003 | 0.015 | 0.017 | -2.0940 | 3.63E-02 | 0.1741 | Increasing |
| HNSC-US::SP33496 | Head&Neck | HNSC-US | 1216 | 0 | 0 | 22 | 13 | 0.012 | 0.014 | 0.020 | 0.040 | -2.0684 | 3.86E-02 | 0.1741 | Increasing |
| LIRI-JP::SP99289 | Liver | LIRI-JP | 728 | 0 | 105 | 45 | 13 | 0.025 | 0.038 | 0.029 | 0.075 | -2.0817 | 3.74E-02 | 0.1741 | Increasing |
| LUSC-US::SP56460 | Lung | LUSC-US | 1404 | 0 | 260 | 51 | 12 | 0.061 | 0.048 | 0.029 | 0.028 | 2.0640 | 3.90E-02 | 0.1741 | Decreasing |
| PRAD-CA::SP112857 | Prostate | PRAD-US | 306 | 0 | 33 | 12 | 9.5 | 0.004 | 0.005 | 0.015 | 0.020 | -2.0789 | 3.76E-02 | 0.1741 | Increasing |
| LICA-FR::SP97775 | Liver | LICA-FR | 1011 | 0 | 199 | 84 | 11 | 0.056 | 0.042 | 0.090 | 0.084 | -2.0782 | 3.77E-02 | 0.1741 | Increasing |
| LIRI-JP::SP112269 | Liver | LIRI-JP | 687 | 0 | 84 | 43 | 10.5 | 0.019 | 0.040 | 0.032 | 0.064 | -2.0652 | 3.89E-02 | 0.1741 | Increasing |
| OV-AU::SP101532 | Ovary | OV-AU | 646 | 278 | 253 | 146 | 21 | 0.156 | 0.102 | 0.102 | 0.093 | 2.0872 | 3.69E-02 | 0.1741 | Decreasing |
| LIRI-JP::SP99025 | Liver | LIRI-JP | 651 | 0 | 101 | 43 | 16 | 0.022 | 0.033 | 0.041 | 0.058 | -2.0652 | 3.89E-02 | 0.1741 | Increasing |
| PACA-CA::SP125760 | Pancreas | PACA-CA | 412 | 0 | 92 | 12 | 12 | 0.011 | 0.022 | 0.000 | 0.003 | 2.0693 | 3.85E-02 | 0.1741 | Decreasing |
| PACA-CA::SP125770 | Pancreas | PACA-CA | 21973 | 0 | 0 | 397 | 14 | 0.347 | 0.319 | 0.321 | 0.245 | 2.0903 | 3.66E-02 | 0.1741 | Decreasing |
| LIRI-JP::SP99077 | Liver | LIRI-JP | 475 | 0 | 74 | 36 | 13 | 0.022 | 0.018 | 0.045 | 0.046 | -2.0934 | 3.63E-02 | 0.1741 | Increasing |
| MALY-DE::SP116674 | Blood | MALY-DE | 677 | 0 | 0 | 16 | 9 | 0.000 | 0.011 | 0.009 | 0.024 | -2.0876 | 3.68E-02 | 0.1741 | Increasing |
| GBM-US::SP25518 | Brain | GBM-US | 560 | 0 | 0 | 16 | 5 | 0.005 | 0.009 | 0.018 | 0.026 | -2.0676 | 3.87E-02 | 0.1741 | Increasing |
| LUSC-US::SP57251 | Lung | LUSC-US | 1885 | 0 | 0 | 44 | 10 | 0.051 | 0.037 | 0.042 | 0.006 | 2.0648 | 3.89E-02 | 0.1741 | Decreasing |
| BLCA-US::SP1419 | Bladder | BLCA-US | 783 | 0 | 104 | 36 | 9 | 0.015 | 0.029 | 0.043 | 0.042 | -2.0674 | 3.87E-02 | 0.1741 | Increasing |
| PRAD-CA::SP112945 | Prostate | PRAD-US | 348 | 0 | 49 | 18 | 8 | 0.004 | 0.008 | 0.033 | 0.015 | -2.0823 | 3.73E-02 | 0.1741 | Increasing |
| PRAD-CA::SP112863 | Prostate | PRAD-US | 437 | 0 | 0 | 15 | 8.5 | 0.007 | 0.008 | 0.012 | 0.030 | -2.0702 | 3.84E-02 | 0.1741 | Increasing |
| PACA-CA::SP125712 | Pancreas | PACA-CA | 557 | 0 | 167 | 20 | 14 | 0.015 | 0.003 | 0.014 | 0.032 | -2.0677 | 3.87E-02 | 0.1741 | Increasing |
| LIRI-JP::SP99205 | Liver | LIRI-JP | 321 | 0 | 68 | 34 | 16 | 0.019 | 0.023 | 0.032 | 0.052 | -2.0817 | 3.74E-02 | 0.1741 | Increasing |
| MALY-DE::SP116648 | Blood | MALY-DE | 2282 | 0 | 0 | 25 | 10 | 0.000 | 0.020 | 0.014 | 0.033 | -2.0685 | 3.86E-02 | 0.1741 | Increasing |
| PRAD-UK::SP111146 | Prostate | PRAD-US | 244 | 0 | 48 | 12 | 9 | 0.000 | 0.011 | 0.012 | 0.020 | -2.0789 | 3.76E-02 | 0.1741 | Increasing |
| BRCA-UK::SP116341 | Breast | BRCA-US | 591 | 363 | 103 | 103 | 15 | 0.086 | 0.129 | 0.060 | 0.053 | 2.0736 | 3.81E-02 | 0.1741 | Decreasing |
| CESC-US::SP107603 | Cervix | CESC-US | 279 | 0 | 0 | 16 | 10 | 0.008 | 0.011 | 0.010 | 0.037 | -2.0399 | 4.14E-02 | 0.1839 | Increasing |
| LUSC-US::SP57818 | Lung | LUSC-US | 2055 | 0 | 308 | 70 | 13 | 0.035 | 0.069 | 0.055 | 0.091 | -2.0328 | 4.21E-02 | 0.1843 | Increasing |
| BRCA-US::SP6766 | Breast | BRCA-US | 189 | 43 | 35 | 18 | 15 | 0.007 | 0.011 | 0.022 | 0.026 | -2.0340 | 4.20E-02 | 0.1843 | Increasing |
| STAD-US::SP84719 | Stomach | STAD-US | 864 | 0 | 0 | 12 | 11 | 0.006 | 0.006 | 0.010 | 0.027 | -2.0375 | 4.16E-02 | 0.1843 | Increasing |
| LIRI-JP::SP50105 | Liver | LIRI-JP | 584 | 0 | 100 | 39 | 12 | 0.029 | 0.020 | 0.035 | 0.064 | -2.0333 | 4.20E-02 | 0.1843 | Increasing |
| PACA-CA::SP125793 | Pancreas | PACA-CA | 313 | 104 | 60 | 25 | 12 | 0.041 | 0.014 | 0.011 | 0.016 | 2.0361 | 4.17E-02 | 0.1843 | Decreasing |
| MELA-AU::SP124462 | Skin | SKCM-US | 973 | 0 | 231 | 72 | 15 | 0.051 | 0.060 | 0.038 | 0.122 | -2.0311 | 4.22E-02 | 0.1845 | Increasing |
| OV-US::SP60362 | Ovary | OV-US | 261 | 0 | 32 | 11 | 10 | 0.000 | 0.008 | 0.020 | 0.012 | -2.0280 | 4.26E-02 | 0.1848 | Increasing |
| OV-AU::SP101740 | Ovary | OV-AU | 2130 | 1160 | 970 | 620 | 19 | 0.567 | 0.454 | 0.479 | 0.424 | 2.0282 | 4.25E-02 | 0.1848 | Decreasing |
| OV-US::SP65376 | Ovary | OV-US | 1333 | 721 | 612 | 317 | 20 | 0.311 | 0.292 | 0.207 | 0.255 | 2.0249 | 4.29E-02 | 0.1856 | Decreasing |
| LUAD-US::SP50827 | Lung | LUAD-US | 1928 | 0 | 0 | 33 | 9.5 | 0.019 | 0.022 | 0.033 | 0.049 | -2.0213 | 4.32E-02 | 0.1866 | Increasing |
| STAD-US::SP84056 | Stomach | STAD-US | 1337 | 260 | 0 | 82 | 12 | 0.105 | 0.056 | 0.052 | 0.064 | 2.0074 | 4.47E-02 | 0.1923 | Decreasing |
| BRCA-US::SP11878 | Breast | BRCA-US | 333 | 0 | 113 | 29 | 11 | 0.024 | 0.008 | 0.031 | 0.048 | -2.0052 | 4.49E-02 | 0.1928 | Increasing |
| LIRI-JP::SP99161 | Liver | LIRI-JP | 468 | 0 | 93 | 35 | 11 | 0.019 | 0.028 | 0.025 | 0.058 | -2.0036 | 4.51E-02 | 0.1930 | Increasing |
| LUSC-US::SP56941 | Lung | LUSC-US | 220 | 0 | 40 | 11 | 8 | 0.003 | 0.005 | 0.016 | 0.017 | -1.9961 | 4.59E-02 | 0.1953 | Increasing |
| RECA-EU::SP102755 | Kidney | RECA-EU | 1917 | 0 | 334 | 145 | 9 | 0.086 | 0.099 | 0.150 | 0.120 | -1.9973 | 4.58E-02 | 0.1953 | Increasing |
| PACA-AU::SP107781 | Pancreas | PACA-AU | 325 | 0 | 0 | 15 | 12 | 0.000 | 0.008 | 0.023 | 0.013 | -1.9788 | 4.78E-02 | 0.1965 | Increasing |
| PACA-AU::SP70288 | Pancreas | PACA-AU | 848 | 0 | 0 | 15 | 20 | 0.004 | 0.003 | 0.026 | 0.013 | -1.9788 | 4.78E-02 | 0.1965 | Increasing |
| UCEC-US::SP92707 | Endometrium | UCEC-US | 335 | 0 | 48 | 15 | 10 | 0.000 | 0.013 | 0.027 | 0.012 | -1.9792 | 4.78E-02 | 0.1965 | Increasing |
| LIHC-US::SP106631 | Liver | LIHC-US | 429 | 0 | 71 | 38 | 13.5 | 0.019 | 0.029 | 0.042 | 0.048 | -1.9859 | 4.70E-02 | 0.1965 | Increasing |
| OV-AU::SP102015 | Ovary | OV-AU | 956 | 373 | 422 | 227 | 21 | 0.207 | 0.186 | 0.159 | 0.140 | 1.9845 | 4.72E-02 | 0.1965 | Decreasing |

| | | | | | | | | | | | | | | | |
|---|---|---|---|---|---|---|---|---|---|---|---|---|---|---|---|
| PACA-CA::SP125775 | Pancreas | PACA-CA | 396 | 0 | 87 | 12 | 12 | 0.000 | 0.008 | 0.011 | 0.016 | -1.9795 | 4.78E-02 | 0.1965 | Increasing |
| PRAD-CA::SP112879 | Prostate | PRAD-US | 213 | 0 | 28 | 14 | 14.5 | 0.004 | 0.008 | 0.018 | 0.020 | -1.9825 | 4.74E-02 | 0.1965 | Increasing |
| COAD-US::SP19750 | CRC | COAD-US | 1265 | 0 | 0 | 13 | 11 | 0.009 | 0.005 | 0.007 | 0.034 | -1.9909 | 4.65E-02 | 0.1965 | Increasing |
| PACA-CA::SP117911 | Pancreas | PACA-CA | 624 | 0 | 0 | 12 | 11 | 0.004 | 0.003 | 0.014 | 0.016 | -1.9795 | 4.78E-02 | 0.1965 | Increasing |
| PACA-CA::SP117076 | Pancreas | PACA-CA | 301 | 0 | 79 | 12 | 15 | 0.008 | 0.003 | 0.006 | 0.022 | -1.9795 | 4.78E-02 | 0.1965 | Increasing |
| LICA-FR::SP97771 | Liver | LICA-FR | 679 | 0 | 126 | 51 | 12 | 0.026 | 0.033 | 0.050 | 0.056 | -1.9828 | 4.74E-02 | 0.1965 | Increasing |
| PAEN-AU::SP117758 | Pancreas | PAEN-AU | 263 | 0 | 47 | 13 | 16 | 0.010 | 0.003 | 0.005 | 0.024 | -1.9814 | 4.75E-02 | 0.1965 | Increasing |
| RECA-EU::SP103547 | Kidney | RECA-EU | 812 | 0 | 161 | 74 | 9 | 0.036 | 0.059 | 0.062 | 0.080 | -1.9723 | 4.86E-02 | 0.1990 | Increasing |
| LIHC-US::SP49591 | Liver | LIHC-US | 403 | 0 | 55 | 21 | 9 | 0.006 | 0.018 | 0.023 | 0.030 | -1.9702 | 4.88E-02 | 0.1994 | Increasing |
| OV-AU::SP102064 | Ovary | OV-AU | 713 | 320 | 311 | 161 | 21 | 0.159 | 0.120 | 0.120 | 0.093 | 1.9687 | 4.90E-02 | 0.1995 | Decreasing |
| LIHC-US::SP49286 | Liver | LIHC-US | 610 | 0 | 106 | 43 | 14 | 0.025 | 0.034 | 0.035 | 0.066 | -1.9578 | 0.0503 | 0.2041 | Increasing |
| PRAD-UK::SP114960 | Prostate | PRAD-US | 1300 | 0 | 0 | 49 | 11 | 0.015 | 0.045 | 0.058 | 0.045 | -1.9521 | 0.0509 | 0.2063 | Increasing |
| LUSC-US::SP56827 | Lung | LUSC-US | 1165 | 0 | 0 | 13 | 9 | 0.006 | 0.008 | 0.010 | 0.028 | -1.9477 | 0.0515 | 0.2070 | Increasing |
| STAD-US::SP85379 | Stomach | STAD-US | 1919 | 0 | 0 | 40 | 12 | 0.022 | 0.028 | 0.046 | 0.048 | -1.9489 | 0.0513 | 0.2070 | Increasing |
| KIRP-US::SP43532 | Kidney | KIRP-US | 1114 | 0 | 163 | 58 | 9 | 0.028 | 0.048 | 0.062 | 0.062 | -1.9470 | 0.0515 | 0.2070 | Increasing |
| PACA-CA::SP125784 | Pancreas | PACA-CA | 401 | 93 | 84 | 17 | 13 | 0.004 | 0.011 | 0.014 | 0.022 | -1.9404 | 0.0523 | 0.2074 | Increasing |
| BRCA-US::SP9481 | Breast | BRCA-US | 144 | 0 | 55 | 12 | 11 | 0.007 | 0.003 | 0.016 | 0.021 | -1.9437 | 0.0519 | 0.2074 | Increasing |
| MALY-DE::SP59456 | Blood | MALY-DE | 651 | 0 | 0 | 20 | 12 | 0.000 | 0.016 | 0.012 | 0.027 | -1.9402 | 0.0524 | 0.2074 | Increasing |
| PACA-CA::SP117440 | Pancreas | PACA-CA | 646 | 0 | 0 | 17 | 10.5 | 0.000 | 0.014 | 0.017 | 0.019 | -1.9404 | 0.0523 | 0.2074 | Increasing |
| LIRI-JP::SP99249 | Liver | LIRI-JP | 482 | 0 | 59 | 21 | 11 | 0.010 | 0.013 | 0.025 | 0.029 | -1.9417 | 0.0522 | 0.2074 | Increasing |
| LIRI-JP::SP50153 | Liver | LIRI-JP | 816 | 0 | 129 | 52 | 14 | 0.032 | 0.035 | 0.054 | 0.064 | -1.9352 | 0.0530 | 0.2093 | Increasing |
| PACA-AU::SP69762 | Pancreas | PACA-AU | 474 | 0 | 0 | 14 | 12 | 0.000 | 0.008 | 0.020 | 0.013 | -1.9284 | 0.0538 | 0.2120 | Increasing |
| SKCM-US::SP82636 | Skin | SKCM-US | 398 | 0 | 0 | 16 | 10.5 | 0.007 | 0.008 | 0.022 | 0.022 | -1.9184 | 0.0551 | 0.2152 | Increasing |
| MELA-AU::SP124336 | Skin | SKCM-US | 466 | 0 | 0 | 16 | 11.5 | 0.003 | 0.013 | 0.019 | 0.022 | -1.9184 | 0.0551 | 0.2152 | Increasing |
| ORCA-IN::SP117509 | Head&Neck | ORCA-IN | 751 | 0 | 0 | 28 | 14 | 0.013 | 0.021 | 0.025 | 0.040 | -1.9195 | 0.0549 | 0.2152 | Increasing |
| STAD-US::SP105006 | Stomach | STAD-US | 1001 | 116 | 0 | 32 | 14 | 0.015 | 0.025 | 0.033 | 0.042 | -1.9129 | 0.0558 | 0.2174 | Increasing |
| PRAD-UK::SP114920 | Prostate | PRAD-US | 867 | 0 | 0 | 16 | 14 | 0.004 | 0.011 | 0.021 | 0.020 | -1.9085 | 0.0563 | 0.2184 | Increasing |
| PRAD-CA::SP112907 | Prostate | PRAD-US | 1459 | 0 | 0 | 16 | 11 | 0.004 | 0.013 | 0.015 | 0.025 | -1.9085 | 0.0563 | 0.2184 | Increasing |
| BTCA-SG::SP117759 | Biliary | BTCA-SG | 1568 | 0 | 0 | 19 | 11 | 0.009 | 0.010 | 0.036 | 0.022 | -1.8991 | 0.0576 | 0.2221 | Increasing |
| HNSC-US::SP33094 | Head&Neck | HNSC-US | 649 | 0 | 85 | 28 | 9 | 0.018 | 0.019 | 0.020 | 0.052 | -1.8988 | 0.0576 | 0.2221 | Increasing |
| LIHC-US::SP98082 | Liver | LIHC-US | 650 | 0 | 105 | 37 | 12 | 0.022 | 0.026 | 0.035 | 0.054 | -1.8955 | 0.0580 | 0.2226 | Increasing |
| PACA-CA::SP78013 | Pancreas | PACA-CA | 797 | 0 | 0 | 16 | 11 | 0.008 | 0.008 | 0.008 | 0.025 | -1.8962 | 0.0579 | 0.2226 | Increasing |
| RECA-EU::SP102929 | Kidney | RECA-EU | 723 | 0 | 121 | 48 | 10 | 0.023 | 0.040 | 0.031 | 0.065 | -1.8907 | 0.0587 | 0.2244 | Increasing |
| HNSC-US::SP34005 | Head&Neck | HNSC-US | 263 | 0 | 39 | 12 | 11 | 0.006 | 0.008 | 0.007 | 0.029 | -1.8851 | 0.0594 | 0.2261 | Increasing |
| SKCM-US::SP82417 | Skin | SKCM-US | 793 | 0 | 95 | 34 | 11 | 0.034 | 0.039 | 0.025 | 0.006 | 1.8861 | 0.0593 | 0.2261 | Decreasing |
| READ-US::SP81494 | CRC | READ-US | 1192 | 0 | 0 | 14 | 9 | 0.003 | 0.013 | 0.013 | 0.023 | -1.8820 | 0.0598 | 0.2271 | Increasing |
| HNSC-US::SP30803 | Head&Neck | HNSC-US | 645 | 0 | 163 | 54 | 10 | 0.030 | 0.049 | 0.042 | 0.075 | -1.8802 | 0.0601 | 0.2274 | Increasing |
| MALY-DE::SP59400 | Blood | MALY-DE | 1440 | 172 | 0 | 48 | 10 | 0.038 | 0.025 | 0.032 | 0.060 | -1.8727 | 0.0611 | 0.2308 | Increasing |
| LIRI-JP::SP98953 | Liver | LIRI-JP | 1216 | 0 | 0 | 46 | 13 | 0.029 | 0.035 | 0.035 | 0.070 | -1.8707 | 0.0614 | 0.2312 | Increasing |
| LIHC-US::SP98078 | Liver | LIHC-US | 791 | 0 | 168 | 64 | 13 | 0.028 | 0.055 | 0.084 | 0.048 | -1.8684 | 0.0617 | 0.2318 | Increasing |
| OV-US::SP63796 | Ovary | OV-US | 185 | 0 | 39 | 12 | 13.5 | 0.003 | 0.010 | 0.010 | 0.023 | -1.8583 | 0.0631 | 0.2347 | Increasing |
| PACA-CA::SP117323 | Pancreas | PACA-CA | 921 | 0 | 214 | 20 | 11 | 0.004 | 0.020 | 0.008 | 0.029 | -1.8586 | 0.0631 | 0.2347 | Increasing |
| PRAD-CA::SP112969 | Prostate | PRAD-US | 242 | 0 | 33 | 18 | 10 | 0.019 | 0.026 | 0.006 | 0.005 | 1.8598 | 0.0629 | 0.2347 | Decreasing |
| UCEC-US::SP89909 | Endometrium | UCEC-US | 12731 | 0 | 0 | 14 | 7 | 0.010 | 0.005 | 0.013 | 0.029 | -1.8595 | 0.0630 | 0.2347 | Increasing |
| PACA-AU::SP70932 | Pancreas | PACA-AU | 538 | 0 | 133 | 17 | 12 | 0.004 | 0.016 | 0.006 | 0.027 | -1.8495 | 0.0644 | 0.2388 | Increasing |
| UCEC-US::SP89957 | Endometrium | UCEC-US | 502 | 0 | 79 | 16 | 10 | 0.006 | 0.010 | 0.020 | 0.023 | -1.8472 | 0.0647 | 0.2394 | Increasing |
| BTCA-SG::SP117712 | Biliary | BTCA-SG | 529 | 0 | 62 | 28 | 11 | 0.022 | 0.017 | 0.028 | 0.050 | -1.8425 | 0.0654 | 0.2413 | Increasing |
| COAD-US::SP20993 | CRC | COAD-US | 2072 | 0 | 0 | 16 | 10 | 0.003 | 0.016 | 0.017 | 0.023 | -1.8349 | 0.0665 | 0.2430 | Increasing |
| LIRI-JP::SP112288 | Liver | LIRI-JP | 610 | 0 | 71 | 35 | 13 | 0.019 | 0.025 | 0.035 | 0.046 | -1.8359 | 0.0664 | 0.2430 | Increasing |
| BRCA-UK::SP116365 | Breast | BRCA-US | 672 | 268 | 404 | 177 | 19 | 0.172 | 0.166 | 0.132 | 0.117 | 1.8383 | 0.0660 | 0.2430 | Decreasing |

| | | | | | | | | | | | | | | | |
|---|---|---|---|---|---|---|---|---|---|---|---|---|---|---|---|
| MALY-DE::SP116620 | Blood | MALY-DE | 2042 | 0 | 0 | 16 | 11 | 0.006 | 0.009 | 0.009 | 0.024 | -1.8360 | 0.0664 | 0.2430 | Increasing |
| LIRI-JP::SP99001 | Liver | LIRI-JP | 316 | 0 | 34 | 18 | 9.5 | 0.006 | 0.010 | 0.029 | 0.017 | -1.8310 | 0.0671 | 0.2444 | Increasing |
| LUAD-US::SP53073 | Lung | LUAD-US | 890 | 0 | 78 | 27 | 12 | 0.016 | 0.016 | 0.029 | 0.038 | -1.8283 | 0.0675 | 0.2453 | Increasing |
| OV-US::SP60243 | Ovary | OV-US | 514 | 240 | 274 | 116 | 24 | 0.128 | 0.094 | 0.089 | 0.075 | 1.8217 | 0.0685 | 0.2483 | Decreasing |
| PRAD-UK::SP114910 | Prostate | PRAD-US | 881 | 0 | 0 | 15 | 13 | 0.004 | 0.011 | 0.018 | 0.020 | -1.8162 | 0.0693 | 0.2501 | Increasing |
| PRAD-UK::SP114917 | Prostate | PRAD-US | 840 | 0 | 0 | 15 | 13 | 0.004 | 0.011 | 0.018 | 0.020 | -1.8162 | 0.0693 | 0.2501 | Increasing |
| PACA-CA::SP125780 | Pancreas | PACA-CA | 713 | 0 | 0 | 14 | 14 | 0.008 | 0.003 | 0.014 | 0.019 | -1.8049 | 0.0711 | 0.2558 | Increasing |
| RECA-EU::SP103213 | Kidney | RECA-EU | 569 | 0 | 104 | 35 | 8 | 0.016 | 0.026 | 0.028 | 0.045 | -1.8034 | 0.0713 | 0.2560 | Increasing |
| LIHC-US::SP49391 | Liver | LIHC-US | 334 | 0 | 66 | 27 | 11 | 0.016 | 0.016 | 0.032 | 0.036 | -1.7992 | 0.0720 | 0.2578 | Increasing |
| PRAD-CA::SP112819 | Prostate | PRAD-US | 368 | 0 | 0 | 12 | 8 | 0.007 | 0.005 | 0.009 | 0.025 | -1.7948 | 0.0727 | 0.2581 | Increasing |
| RECA-EU::SP102945 | Kidney | RECA-EU | 621 | 0 | 136 | 60 | 9 | 0.023 | 0.054 | 0.051 | 0.060 | -1.7933 | 0.0729 | 0.2581 | Increasing |
| THCA-US::SP85487 | Thyroid | THCA-US | 241 | 0 | 68 | 15 | 8 | 0.004 | 0.015 | 0.006 | 0.029 | -1.7949 | 0.0727 | 0.2581 | Increasing |
| PRAD-US::SP79968 | Prostate | PRAD-US | 435 | 0 | 0 | 12 | 7.5 | 0.007 | 0.003 | 0.015 | 0.020 | -1.7948 | 0.0727 | 0.2581 | Increasing |
| ORCA-IN::SP117003 | Head&Neck | ORCA-IN | 552 | 197 | 122 | 55 | 12 | 0.058 | 0.052 | 0.034 | 0.030 | 1.7929 | 0.0730 | 0.2581 | Decreasing |
| LIRI-JP::SP99279 | Liver | LIRI-JP | 590 | 0 | 83 | 38 | 13 | 0.032 | 0.015 | 0.038 | 0.058 | -1.7843 | 0.0744 | 0.2624 | Increasing |
| KIRP-US::SP109478 | Kidney | KIRP-US | 197 | 0 | 31 | 11 | 12 | 0.003 | 0.005 | 0.016 | 0.015 | -1.7819 | 0.0748 | 0.2631 | Increasing |
| SKCM-US::SP82451 | Skin | SKCM-US | 675 | 0 | 99 | 39 | 13 | 0.010 | 0.044 | 0.035 | 0.044 | -1.7784 | 0.0753 | 0.2638 | Increasing |
| MALY-DE::SP59448 | Blood | MALY-DE | 3383 | 0 | 0 | 22 | 9 | 0.013 | 0.013 | 0.009 | 0.033 | -1.7774 | 0.0755 | 0.2638 | Increasing |
| MELA-AU::SP124367 | Skin | SKCM-US | 758 | 0 | 171 | 61 | 13 | 0.038 | 0.049 | 0.054 | 0.078 | -1.7764 | 0.0757 | 0.2638 | Increasing |
| HNSC-US::SP31814 | Head&Neck | HNSC-US | 392 | 80 | 0 | 28 | 11 | 0.024 | 0.041 | 0.013 | 0.006 | 1.7790 | 0.0752 | 0.2638 | Decreasing |
| MELA-AU::SP124326 | Skin | SKCM-US | 506 | 0 | 119 | 37 | 14 | 0.027 | 0.026 | 0.022 | 0.067 | -1.7697 | 0.0768 | 0.2670 | Increasing |
| PACA-CA::SP117661 | Pancreas | PACA-CA | 705 | 0 | 0 | 18 | 11 | 0.008 | 0.008 | 0.017 | 0.022 | -1.7633 | 0.0779 | 0.2701 | Increasing |
| PACA-CA::SP77930 | Pancreas | PACA-CA | 236 | 0 | 64 | 13 | 10 | 0.008 | 0.006 | 0.006 | 0.022 | -1.7578 | 0.0788 | 0.2707 | Increasing |
| PRAD-UK::SP114900 | Prostate | PRAD-US | 969 | 0 | 0 | 17 | 10 | 0.007 | 0.011 | 0.018 | 0.025 | -1.7585 | 0.0787 | 0.2707 | Increasing |
| HNSC-US::SP32694 | Head&Neck | HNSC-US | 829 | 0 | 83 | 32 | 13.5 | 0.015 | 0.022 | 0.046 | 0.029 | -1.7596 | 0.0785 | 0.2707 | Increasing |
| PRAD-CA::SP112875 | Prostate | PRAD-US | 395 | 0 | 47 | 17 | 10 | 0.011 | 0.005 | 0.021 | 0.025 | -1.7585 | 0.0787 | 0.2707 | Increasing |
| MELA-AU::SP124391 | Skin | SKCM-US | 324 | 0 | 53 | 21 | 10 | 0.017 | 0.005 | 0.025 | 0.033 | -1.7530 | 0.0796 | 0.2726 | Increasing |
| PACA-CA::SP117165 | Pancreas | PACA-CA | 482 | 0 | 134 | 26 | 12 | 0.015 | 0.008 | 0.028 | 0.029 | -1.7524 | 0.0797 | 0.2726 | Increasing |
| MALY-DE::SP116701 | Blood | MALY-DE | 842 | 0 | 0 | 21 | 11 | 0.000 | 0.018 | 0.012 | 0.027 | -1.7465 | 0.0807 | 0.2735 | Increasing |
| MALY-DE::SP116683 | Blood | MALY-DE | 670 | 0 | 0 | 21 | 11.5 | 0.013 | 0.009 | 0.017 | 0.027 | -1.7465 | 0.0807 | 0.2735 | Increasing |
| RECA-EU::SP103444 | Kidney | RECA-EU | 565 | 0 | 90 | 48 | 9 | 0.016 | 0.049 | 0.028 | 0.060 | -1.7473 | 0.0806 | 0.2735 | Increasing |
| LIRI-JP::SP99225 | Liver | LIRI-JP | 197 | 0 | 26 | 11 | 14.5 | 0.003 | 0.008 | 0.013 | 0.017 | -1.7471 | 0.0806 | 0.2735 | Increasing |
| COAD-US::SP18121 | CRC | COAD-US | 101932 | 0 | 0 | 23 | 6 | 0.009 | 0.022 | 0.020 | 0.034 | -1.7441 | 0.0811 | 0.2743 | Increasing |
| PRAD-UK::SP114968 | Prostate | PRAD-US | 3091 | 1535 | 876 | 524 | 13 | 0.540 | 0.411 | 0.417 | 0.432 | 1.7351 | 0.0827 | 0.2777 | Decreasing |
| LIRI-JP::SP107155 | Liver | LIRI-JP | 436 | 0 | 56 | 23 | 13.5 | 0.003 | 0.025 | 0.025 | 0.023 | -1.7365 | 0.0825 | 0.2777 | Increasing |
| PAEN-AU::SP102537 | Pancreas | PAEN-AU | 194 | 0 | 32 | 14 | 12 | 0.005 | 0.003 | 0.019 | 0.015 | -1.7350 | 0.0827 | 0.2777 | Increasing |
| LIHC-US::SP98327 | Liver | LIHC-US | 615 | 0 | 82 | 37 | 12.5 | 0.019 | 0.031 | 0.035 | 0.048 | -1.7329 | 0.0831 | 0.2777 | Increasing |
| KIRC-US::SP38759 | Kidney | KIRC-US | 750 | 0 | 125 | 52 | 10 | 0.022 | 0.051 | 0.046 | 0.059 | -1.7337 | 0.0830 | 0.2777 | Increasing |
| RECA-EU::SP103483 | Kidney | RECA-EU | 421 | 0 | 54 | 28 | 10 | 0.016 | 0.014 | 0.028 | 0.035 | -1.7256 | 0.0844 | 0.2814 | Increasing |
| PRAD-CA::SP112973 | Prostate | PRAD-US | 467 | 0 | 54 | 14 | 10 | 0.000 | 0.013 | 0.018 | 0.015 | -1.7195 | 0.0855 | 0.2833 | Increasing |
| CESC-US::SP114020 | Cervix | CESC-US | 654 | 0 | 0 | 19 | 12 | 0.008 | 0.014 | 0.023 | 0.025 | -1.7192 | 0.0856 | 0.2833 | Increasing |
| PRAD-UK::SP111122 | Prostate | PRAD-US | 316 | 0 | 38 | 14 | 10 | 0.007 | 0.013 | 0.000 | 0.035 | -1.7195 | 0.0855 | 0.2833 | Increasing |
| PACA-CA::SP125753 | Pancreas | PACA-CA | 583 | 0 | 0 | 12 | 15 | 0.004 | 0.008 | 0.006 | 0.019 | -1.7096 | 0.0873 | 0.2833 | Increasing |
| GBM-US::SP24702 | Brain | GBM-US | 489 | 0 | 0 | 12 | 5 | 0.000 | 0.009 | 0.015 | 0.015 | -1.7166 | 0.0861 | 0.2833 | Increasing |
| MALY-DE::SP116709 | Blood | MALY-DE | 441 | 0 | 0 | 11 | 8 | 0.006 | 0.002 | 0.012 | 0.015 | -1.7120 | 0.0869 | 0.2833 | Increasing |
| LUSC-US::SP57084 | Lung | LUSC-US | 1992 | 0 | 0 | 45 | 11.5 | 0.032 | 0.032 | 0.036 | 0.068 | -1.7133 | 0.0867 | 0.2833 | Increasing |
| PAEN-AU::SP106797 | Pancreas | PAEN-AU | 140 | 0 | 21 | 13 | 10 | 0.000 | 0.008 | 0.013 | 0.015 | -1.7106 | 0.0872 | 0.2833 | Increasing |
| READ-US::SP81840 | CRC | READ-US | 2420 | 0 | 0 | 26 | 10 | 0.012 | 0.021 | 0.027 | 0.035 | -1.7169 | 0.0860 | 0.2833 | Increasing |
| PACA-CA::SP125731 | Pancreas | PACA-CA | 451 | 0 | 118 | 12 | 13.5 | 0.008 | 0.003 | 0.008 | 0.019 | -1.7096 | 0.0873 | 0.2833 | Increasing |
| PACA-CA::SP125761 | Pancreas | PACA-CA | 365 | 0 | 99 | 12 | 10 | 0.008 | 0.003 | 0.008 | 0.019 | -1.7096 | 0.0873 | 0.2833 | Increasing |

| | | | | | | | | | | | | | | | |
|---|---|---|---|---|---|---|---|---|---|---|---|---|---|---|---|
| PRAD-CA::SP112913 | Prostate | PRAD-US | 220 | 0 | 40 | 19 | 10 | 0.011 | 0.013 | 0.012 | 0.035 | -1.7130 | 0.0867 | 0.2833 | Increasing |
| KIRC-US::SP36036 | Kidney | KIRC-US | 424 | 0 | 81 | 19 | 7.5 | 0.007 | 0.013 | 0.022 | 0.025 | -1.7060 | 0.0880 | 0.2848 | Increasing |
| PACA-AU::SP110803 | Pancreas | PACA-AU | 381 | 0 | 234 | 47 | 9 | 0.059 | 0.030 | 0.035 | 0.027 | 1.7009 | 0.0890 | 0.2866 | Decreasing |
| STAD-US::SP85431 | Stomach | STAD-US | 523 | 64 | 0 | 22 | 11 | 0.015 | 0.014 | 0.013 | 0.042 | -1.7012 | 0.0889 | 0.2866 | Increasing |
| LIRI-JP::SP99185 | Liver | LIRI-JP | 435 | 0 | 64 | 30 | 12 | 0.016 | 0.025 | 0.022 | 0.046 | -1.6997 | 0.0892 | 0.2867 | Increasing |
| PRAD-CA::SP112797 | Prostate | PRAD-US | 387 | 0 | 0 | 11 | 11 | 0.004 | 0.008 | 0.009 | 0.020 | -1.6937 | 0.0903 | 0.2891 | Increasing |
| PRAD-US::SP80205 | Prostate | PRAD-US | 226 | 0 | 30 | 11 | 9 | 0.007 | 0.003 | 0.012 | 0.020 | -1.6937 | 0.0903 | 0.2891 | Increasing |
| MELA-AU::SP124380 | Skin | SKCM-US | 749 | 0 | 102 | 33 | 12.5 | 0.034 | 0.036 | 0.022 | 0.011 | 1.6861 | 0.0918 | 0.2926 | Decreasing |
| READ-US::SP80754 | CRC | READ-US | 1336 | 0 | 0 | 31 | 11 | 0.018 | 0.019 | 0.040 | 0.035 | -1.6859 | 0.0918 | 0.2926 | Increasing |
| KIRP-US::SP43696 | Kidney | KIRP-US | 1359 | 0 | 189 | 71 | 9 | 0.042 | 0.064 | 0.056 | 0.088 | -1.6826 | 0.0924 | 0.2927 | Increasing |
| MALY-DE::SP116663 | Blood | MALY-DE | 1943 | 0 | 0 | 19 | 10 | 0.006 | 0.011 | 0.014 | 0.024 | -1.6833 | 0.0923 | 0.2927 | Increasing |
| LIHC-US::SP98289 | Liver | LIHC-US | 763 | 0 | 132 | 45 | 11.5 | 0.035 | 0.021 | 0.055 | 0.054 | -1.6841 | 0.0922 | 0.2927 | Increasing |
| KIRP-US::SP43792 | Kidney | KIRP-US | 1295 | 0 | 216 | 82 | 9 | 0.059 | 0.056 | 0.081 | 0.093 | -1.6792 | 0.0931 | 0.2942 | Increasing |
| BRCA-US::SP5784 | Breast | BRCA-US | 456 | 0 | 110 | 19 | 11 | 0.017 | 0.008 | 0.009 | 0.042 | -1.6774 | 0.0935 | 0.2946 | Increasing |
| THCA-US::SP87534 | Thyroid | THCA-US | 189 | 0 | 37 | 11 | 9 | 0.000 | 0.008 | 0.019 | 0.010 | -1.6758 | 0.0938 | 0.2950 | Increasing |
| BRCA-US::SP8831 | Breast | BRCA-US | 2055 | 1320 | 546 | 473 | 15 | 0.369 | 0.375 | 0.444 | 0.440 | -1.6738 | 0.0942 | 0.2955 | Increasing |
| KIRP-US::SP109457 | Kidney | KIRP-US | 448 | 0 | 60 | 24 | 12 | 0.007 | 0.024 | 0.022 | 0.031 | -1.6717 | 0.0946 | 0.2962 | Increasing |
| LUSC-US::SP58882 | Lung | LUSC-US | 1035 | 0 | 211 | 41 | 11 | 0.022 | 0.034 | 0.039 | 0.051 | -1.6694 | 0.0950 | 0.2970 | Increasing |
| MELA-AU::SP124369 | Skin | SKCM-US | 905 | 0 | 191 | 63 | 13 | 0.044 | 0.044 | 0.060 | 0.078 | -1.6665 | 0.0956 | 0.2982 | Increasing |
| PRAD-UK::SP114926 | Prostate | PRAD-US | 569 | 0 | 0 | 16 | 13 | 0.004 | 0.013 | 0.018 | 0.020 | -1.6626 | 0.0964 | 0.2988 | Increasing |
| PACA-CA::SP125762 | Pancreas | PACA-CA | 575 | 0 | 166 | 16 | 12.5 | 0.004 | 0.011 | 0.014 | 0.019 | -1.6624 | 0.0964 | 0.2988 | Increasing |
| PRAD-CA::SP112893 | Prostate | PRAD-US | 236 | 0 | 37 | 16 | 11 | 0.007 | 0.008 | 0.021 | 0.020 | -1.6626 | 0.0964 | 0.2988 | Increasing |
| PACA-CA::SP125744 | Pancreas | PACA-CA | 492 | 0 | 0 | 11 | 14.5 | 0.000 | 0.011 | 0.006 | 0.016 | -1.6603 | 0.0969 | 0.2994 | Increasing |
| RECA-EU::SP102973 | Kidney | RECA-EU | 1218 | 0 | 115 | 43 | 9 | 0.013 | 0.040 | 0.040 | 0.040 | -1.6570 | 0.0975 | 0.3009 | Increasing |
| RECA-EU::SP103233 | Kidney | RECA-EU | 862 | 0 | 129 | 58 | 8 | 0.026 | 0.052 | 0.043 | 0.065 | -1.6545 | 0.0980 | 0.3018 | Increasing |
| PACA-CA::SP117656 | Pancreas | PACA-CA | 577 | 166 | 174 | 83 | 18.5 | 0.075 | 0.087 | 0.039 | 0.057 | 1.6533 | 0.0983 | 0.3019 | Decreasing |
| LIRI-JP::SP99261 | Liver | LIRI-JP | 604 | 0 | 108 | 40 | 16 | 0.038 | 0.018 | 0.025 | 0.075 | -1.6490 | 0.0992 | 0.3020 | Increasing |
| LIRI-JP::SP98898 | Liver | LIRI-JP | 708 | 0 | 101 | 40 | 13 | 0.019 | 0.033 | 0.045 | 0.041 | -1.6490 | 0.0992 | 0.3020 | Increasing |
| MELA-AU::SP124377 | Skin | SKCM-US | 562 | 0 | 180 | 75 | 16 | 0.048 | 0.060 | 0.072 | 0.083 | -1.6498 | 0.0990 | 0.3020 | Increasing |
| PACA-CA::SP125755 | Pancreas | PACA-CA | 590 | 0 | 203 | 20 | 12 | 0.011 | 0.011 | 0.011 | 0.029 | -1.6495 | 0.0990 | 0.3020 | Increasing |
| RECA-EU::SP102733 | Kidney | RECA-EU | 460 | 0 | 56 | 23 | 9 | 0.007 | 0.019 | 0.023 | 0.025 | -1.6453 | 0.0999 | 0.3037 | Increasing |
| BRCA-US::SP8795 | Breast | BRCA-US | 815 | 0 | 169 | 29 | 10 | 0.014 | 0.024 | 0.028 | 0.037 | -1.6411 | 0.1008 | 0.3051 | Increasing |
| KIRC-US::SP39248 | Kidney | KIRC-US | 424 | 0 | 57 | 23 | 9 | 0.015 | 0.013 | 0.022 | 0.034 | -1.6417 | 0.1007 | 0.3051 | Increasing |
| STAD-US::SP84858 | Stomach | STAD-US | 1939 | 290 | 0 | 99 | 12 | 0.071 | 0.076 | 0.092 | 0.111 | -1.6306 | 0.1030 | 0.3111 | Increasing |
| LUSC-US::SP56730 | Lung | LUSC-US | 1706 | 0 | 0 | 37 | 12.5 | 0.029 | 0.026 | 0.020 | 0.068 | -1.6252 | 0.1041 | 0.3135 | Increasing |
| UCEC-US::SP89443 | Endometrium | UCEC-US | 376 | 0 | 98 | 20 | 13 | 0.006 | 0.016 | 0.030 | 0.017 | -1.6248 | 0.1042 | 0.3135 | Increasing |
| PACA-AU::SP70677 | Pancreas | PACA-AU | 210 | 0 | 36 | 13 | 14 | 0.004 | 0.008 | 0.012 | 0.017 | -1.6173 | 0.1058 | 0.3157 | Increasing |
| LUAD-US::SP54745 | Lung | LUAD-US | 1101 | 0 | 0 | 23 | 12 | 0.013 | 0.014 | 0.029 | 0.027 | -1.6201 | 0.1052 | 0.3157 | Increasing |
| PACA-AU::SP108088 | Pancreas | PACA-AU | 503 | 0 | 0 | 13 | 12 | 0.004 | 0.005 | 0.018 | 0.013 | -1.6173 | 0.1058 | 0.3157 | Increasing |
| PACA-AU::SP71950 | Pancreas | PACA-AU | 596 | 0 | 145 | 13 | 14 | 0.000 | 0.016 | 0.003 | 0.020 | -1.6173 | 0.1058 | 0.3157 | Increasing |
| PRAD-CA::SP112785 | Prostate | PRAD-US | 668 | 185 | 76 | 68 | 11 | 0.078 | 0.058 | 0.049 | 0.045 | 1.6132 | 0.1067 | 0.3177 | Decreasing |
| LIRI-JP::SP99081 | Liver | LIRI-JP | 572 | 0 | 84 | 43 | 13 | 0.032 | 0.030 | 0.029 | 0.070 | -1.6114 | 0.1071 | 0.3182 | Increasing |
| MELA-AU::SP124273 | Skin | SKCM-US | 516 | 0 | 0 | 11 | 8 | 0.003 | 0.005 | 0.019 | 0.011 | -1.6093 | 0.1076 | 0.3184 | Increasing |
| OV-US::SP60322 | Ovary | OV-US | 403 | 202 | 96 | 86 | 14 | 0.101 | 0.058 | 0.076 | 0.052 | 1.6091 | 0.1076 | 0.3184 | Decreasing |
| PRAD-UK::SP114966 | Prostate | PRAD-US | 3007 | 1497 | 838 | 510 | 13 | 0.503 | 0.414 | 0.423 | 0.397 | 1.6080 | 0.1078 | 0.3185 | Decreasing |
| LIRI-JP::SP99013 | Liver | LIRI-JP | 832 | 121 | 217 | 103 | 11 | 0.083 | 0.068 | 0.089 | 0.127 | -1.6051 | 0.1085 | 0.3197 | Increasing |
| LIRI-JP::SP107068 | Liver | LIRI-JP | 555 | 0 | 0 | 18 | 12 | 0.013 | 0.005 | 0.025 | 0.023 | -1.5972 | 0.1102 | 0.3229 | Increasing |
| LIRI-JP::SP112165 | Liver | LIRI-JP | 493 | 0 | 64 | 29 | 9 | 0.019 | 0.015 | 0.035 | 0.035 | -1.5975 | 0.1102 | 0.3229 | Increasing |
| LIRI-JP::SP112261 | Liver | LIRI-JP | 310 | 0 | 70 | 29 | 11 | 0.019 | 0.018 | 0.029 | 0.041 | -1.5975 | 0.1102 | 0.3229 | Increasing |
| STAD-US::SP105425 | Stomach | STAD-US | 1650 | 0 | 0 | 21 | 11 | 0.009 | 0.020 | 0.016 | 0.032 | -1.5954 | 0.1106 | 0.3234 | Increasing |

| | | | | | | | | | | | | | | | |
|---|---|---|---|---|---|---|---|---|---|---|---|---|---|---|---|
| BRCA-UK::SP116343 | Breast | BRCA-US | 406 | 0 | 106 | 16 | 8 | 0.014 | 0.008 | 0.006 | 0.037 | -1.5908 | 0.1116 | 0.3257 | Increasing |
| COAD-US::SP16934 | CRC | COAD-US | 1582 | 0 | 0 | 12 | 12 | 0.006 | 0.008 | 0.010 | 0.023 | -1.5891 | 0.1120 | 0.3262 | Increasing |
| LIRI-JP::SP107109 | Liver | LIRI-JP | 293 | 0 | 43 | 14 | 7.5 | 0.000 | 0.020 | 0.006 | 0.023 | -1.5854 | 0.1129 | 0.3280 | Increasing |
| RECA-EU::SP103673 | Kidney | RECA-EU | 1191 | 0 | 166 | 57 | 9 | 0.036 | 0.031 | 0.065 | 0.050 | -1.5834 | 0.1133 | 0.3286 | Increasing |
| LIRI-JP::SP112198 | Liver | LIRI-JP | 459 | 0 | 66 | 27 | 13 | 0.010 | 0.025 | 0.029 | 0.029 | -1.5743 | 0.1154 | 0.3327 | Increasing |
| LIRI-JP::SP99157 | Liver | LIRI-JP | 495 | 0 | 76 | 27 | 13 | 0.016 | 0.018 | 0.029 | 0.035 | -1.5743 | 0.1154 | 0.3327 | Increasing |
| SKCM-US::SP83027 | Skin | SKCM-US | 232 | 0 | 43 | 13 | 12 | 0.010 | 0.000 | 0.022 | 0.017 | -1.5750 | 0.1152 | 0.3327 | Increasing |
| KIRP-US::SP43510 | Kidney | KIRP-US | 854 | 0 | 164 | 60 | 9 | 0.028 | 0.048 | 0.081 | 0.041 | -1.5717 | 0.1160 | 0.3338 | Increasing |
| OV-US::SP60001 | Ovary | OV-US | 1030 | 594 | 180 | 230 | 17 | 0.209 | 0.221 | 0.168 | 0.162 | 1.5701 | 0.1164 | 0.3342 | Decreasing |
| PACA-AU::SP69565 | Pancreas | PACA-AU | 1509 | 683 | 0 | 175 | 15 | 0.179 | 0.122 | 0.129 | 0.124 | 1.5623 | 0.1182 | 0.3362 | Decreasing |
| PRAD-UK::SP114916 | Prostate | PRAD-US | 794 | 0 | 0 | 15 | 13 | 0.004 | 0.011 | 0.021 | 0.015 | -1.5622 | 0.1182 | 0.3362 | Increasing |
| PRAD-US::SP102690 | Prostate | PRAD-US | 232 | 0 | 32 | 15 | 7 | 0.007 | 0.008 | 0.018 | 0.020 | -1.5622 | 0.1182 | 0.3362 | Increasing |
| MALY-DE::SP59440 | Blood | MALY-DE | 773 | 0 | 0 | 33 | 13 | 0.006 | 0.031 | 0.012 | 0.042 | -1.5637 | 0.1179 | 0.3362 | Increasing |
| PACA-AU::SP72262 | Pancreas | PACA-AU | 378 | 0 | 101 | 23 | 12 | 0.007 | 0.022 | 0.012 | 0.030 | -1.5660 | 0.1173 | 0.3362 | Increasing |
| RECA-EU::SP102795 | Kidney | RECA-EU | 1496 | 0 | 264 | 110 | 10 | 0.086 | 0.071 | 0.077 | 0.135 | -1.5600 | 0.1187 | 0.3370 | Increasing |
| LIRI-JP::SP99003 | Liver | LIRI-JP | 856 | 0 | 106 | 39 | 13 | 0.019 | 0.035 | 0.035 | 0.046 | -1.5568 | 0.1195 | 0.3372 | Increasing |
| PACA-CA::SP125721 | Pancreas | PACA-CA | 506 | 0 | 138 | 14 | 9 | 0.000 | 0.014 | 0.011 | 0.016 | -1.5550 | 0.1199 | 0.3372 | Increasing |
| RECA-EU::SP102881 | Kidney | RECA-EU | 518 | 0 | 86 | 36 | 8 | 0.020 | 0.024 | 0.034 | 0.040 | -1.5546 | 0.1200 | 0.3372 | Increasing |
| PACA-AU::SP76017 | Pancreas | PACA-AU | 406 | 0 | 0 | 12 | 12.5 | 0.004 | 0.008 | 0.009 | 0.017 | -1.5539 | 0.1202 | 0.3372 | Increasing |
| PACA-AU::SP110798 | Pancreas | PACA-AU | 427 | 0 | 111 | 12 | 13 | 0.007 | 0.005 | 0.006 | 0.020 | -1.5539 | 0.1202 | 0.3372 | Increasing |
| PACA-AU::SP76843 | Pancreas | PACA-AU | 859 | 0 | 0 | 12 | 11 | 0.004 | 0.008 | 0.009 | 0.017 | -1.5539 | 0.1202 | 0.3372 | Increasing |
| PACA-CA::SP125693 | Pancreas | PACA-CA | 1346 | 0 | 0 | 22 | 12 | 0.011 | 0.017 | 0.008 | 0.032 | -1.5506 | 0.1210 | 0.3387 | Increasing |
| KIRC-US::SP41453 | Kidney | KIRC-US | 578 | 0 | 91 | 39 | 9 | 0.025 | 0.024 | 0.043 | 0.044 | -1.5405 | 0.1234 | 0.3422 | Increasing |
| PAEN-AU::SP102597 | Pancreas | PAEN-AU | 199 | 0 | 24 | 13 | 7 | 0.010 | 0.021 | 0.003 | 0.006 | 1.5396 | 0.1237 | 0.3422 | Decreasing |
| MELA-AU::SP124359 | Skin | SKCM-US | 423 | 0 | 56 | 17 | 12 | 0.010 | 0.005 | 0.028 | 0.017 | -1.5430 | 0.1228 | 0.3422 | Increasing |
| BRCA-US::SP9433 | Breast | BRCA-US | 217 | 0 | 45 | 11 | 12 | 0.007 | 0.005 | 0.009 | 0.021 | -1.5408 | 0.1234 | 0.3422 | Increasing |
| RECA-EU::SP102716 | Kidney | RECA-EU | 336 | 0 | 50 | 11 | 8 | 0.007 | 0.005 | 0.009 | 0.020 | -1.5415 | 0.1232 | 0.3422 | Increasing |
| RECA-EU::SP103856 | Kidney | RECA-EU | 169 | 0 | 26 | 11 | 9 | 0.007 | 0.002 | 0.014 | 0.015 | -1.5415 | 0.1232 | 0.3422 | Increasing |
| BRCA-US::SP2826 | Breast | BRCA-US | 327 | 0 | 249 | 66 | 13 | 0.086 | 0.040 | 0.053 | 0.048 | 1.5357 | 0.1246 | 0.3442 | Decreasing |
| LIRI-JP::SP112147 | Liver | LIRI-JP | 547 | 0 | 84 | 37 | 11 | 0.019 | 0.033 | 0.032 | 0.046 | -1.5289 | 0.1263 | 0.3468 | Increasing |
| PACA-CA::SP125758 | Pancreas | PACA-CA | 421 | 0 | 94 | 12 | 15.5 | 0.015 | 0.011 | 0.008 | 0.003 | 1.5294 | 0.1262 | 0.3468 | Decreasing |
| LIRI-JP::SP99193 | Liver | LIRI-JP | 404 | 0 | 0 | 23 | 12.5 | 0.013 | 0.018 | 0.019 | 0.035 | -1.5297 | 0.1261 | 0.3468 | Increasing |
| OV-US::SP59803 | Ovary | OV-US | 572 | 62 | 108 | 39 | 12 | 0.026 | 0.025 | 0.043 | 0.046 | -1.5269 | 0.1268 | 0.3475 | Increasing |
| PACA-CA::SP125737 | Pancreas | PACA-CA | 524 | 71 | 73 | 32 | 11 | 0.019 | 0.014 | 0.034 | 0.032 | -1.5245 | 0.1274 | 0.3478 | Increasing |
| PACA-AU::SP75322 | Pancreas | PACA-AU | 1788 | 0 | 0 | 46 | 12 | 0.033 | 0.022 | 0.041 | 0.050 | -1.5250 | 0.1273 | 0.3478 | Increasing |
| PRAD-UK::SP114932 | Prostate | PRAD-US | 555 | 0 | 0 | 17 | 13 | 0.004 | 0.016 | 0.018 | 0.020 | -1.5199 | 0.1285 | 0.3503 | Increasing |
| MALY-DE::SP116645 | Blood | MALY-DE | 1338 | 0 | 0 | 14 | 10 | 0.006 | 0.007 | 0.012 | 0.018 | -1.5157 | 0.1296 | 0.3526 | Increasing |
| SKCM-US::SP82445 | Skin | SKCM-US | 1740 | 0 | 0 | 52 | 14 | 0.065 | 0.034 | 0.047 | 0.028 | 1.5101 | 0.1310 | 0.3557 | Decreasing |
| LIRI-JP::SP112138 | Liver | LIRI-JP | 466 | 0 | 60 | 21 | 11 | 0.010 | 0.015 | 0.025 | 0.023 | -1.5087 | 0.1314 | 0.3561 | Increasing |
| LIHC-US::SP98359 | Liver | LIHC-US | 707 | 0 | 112 | 43 | 11 | 0.022 | 0.042 | 0.035 | 0.054 | -1.5055 | 0.1322 | 0.3576 | Increasing |
| PRAD-CA::SP112859 | Prostate | PRAD-US | 465 | 0 | 58 | 20 | 10 | 0.026 | 0.021 | 0.006 | 0.015 | 1.4960 | 0.1346 | 0.3578 | Decreasing |
| PACA-AU::SP107956 | Pancreas | PACA-AU | 865 | 0 | 0 | 15 | 13 | 0.007 | 0.008 | 0.012 | 0.020 | -1.4957 | 0.1347 | 0.3578 | Increasing |
| GACA-CN::SP135270 | Stomach | STAD-US | 878 | 216 | 183 | 69 | 13 | 0.074 | 0.059 | 0.052 | 0.042 | 1.5003 | 0.1335 | 0.3578 | Decreasing |
| PACA-AU::SP107915 | Pancreas | PACA-AU | 435 | 0 | 0 | 13 | 11 | 0.015 | 0.014 | 0.009 | 0.003 | 1.4963 | 0.1346 | 0.3578 | Decreasing |
| PACA-CA::SP117643 | Pancreas | PACA-CA | 1771 | 894 | 388 | 351 | 13 | 0.264 | 0.341 | 0.234 | 0.242 | 1.4967 | 0.1345 | 0.3578 | Decreasing |
| KIRC-US::SP35989 | Kidney | KIRC-US | 404 | 0 | 69 | 28 | 8 | 0.025 | 0.011 | 0.025 | 0.044 | -1.4990 | 0.1339 | 0.3578 | Increasing |
| HNSC-US::SP32742 | Head&Neck | HNSC-US | 545 | 65 | 0 | 17 | 11.5 | 0.009 | 0.016 | 0.007 | 0.035 | -1.4964 | 0.1346 | 0.3578 | Increasing |
| MELA-AU::SP124386 | Skin | SKCM-US | 875 | 0 | 312 | 83 | 17 | 0.055 | 0.073 | 0.066 | 0.100 | -1.5012 | 0.1333 | 0.3578 | Increasing |
| OV-AU::SP101670 | Ovary | OV-AU | 1219 | 632 | 362 | 263 | 16 | 0.236 | 0.218 | 0.159 | 0.207 | 1.5010 | 0.1333 | 0.3578 | Decreasing |
| LINC-JP::SP98885 | Liver | LIHC-US | 715 | 0 | 121 | 56 | 13 | 0.025 | 0.055 | 0.061 | 0.048 | -1.5000 | 0.1336 | 0.3578 | Increasing |

| | | | | | | | | | | | | | | | |
|---|---|---|---|---|---|---|---|---|---|---|---|---|---|---|---|
| LIRI-JP::SP98907 | Liver | LIRI-JP | 918 | 0 | 118 | 45 | 13 | 0.029 | 0.033 | 0.045 | 0.052 | -1.4902 | 0.1362 | 0.3610 | Increasing |
| KIRP-US::SP97278 | Kidney | KIRP-US | 946 | 0 | 127 | 55 | 8 | 0.028 | 0.051 | 0.053 | 0.057 | -1.4828 | 0.1381 | 0.3655 | Increasing |
| BRCA-US::SP11292 | Breast | BRCA-US | 157 | 67 | 54 | 39 | 14 | 0.045 | 0.032 | 0.035 | 0.016 | 1.4802 | 0.1388 | 0.3666 | Decreasing |
| BLCA-US::SP1059 | Bladder | BLCA-US | 1105 | 0 | 116 | 47 | 10 | 0.037 | 0.058 | 0.039 | 0.006 | 1.4778 | 0.1394 | 0.3670 | Decreasing |
| OV-AU::SP102143 | Ovary | OV-AU | 906 | 609 | 297 | 232 | 18 | 0.245 | 0.150 | 0.150 | 0.202 | 1.4782 | 0.1394 | 0.3670 | Decreasing |
| LUAD-US::SP50263 | Lung | LUAD-US | 522 | 0 | 61 | 29 | 12 | 0.013 | 0.019 | 0.049 | 0.016 | -1.4747 | 0.1403 | 0.3683 | Increasing |
| BRCA-UK::SP116353 | Breast | BRCA-US | 862 | 0 | 162 | 33 | 11.5 | 0.017 | 0.032 | 0.022 | 0.048 | -1.4740 | 0.1405 | 0.3683 | Increasing |
| LIRI-JP::SP99271 | Liver | LIRI-JP | 293 | 0 | 32 | 17 | 12 | 0.003 | 0.015 | 0.025 | 0.012 | -1.4720 | 0.1410 | 0.3684 | Increasing |
| BLCA-US::SP1144 | Bladder | BLCA-US | 391 | 0 | 71 | 25 | 17 | 0.018 | 0.011 | 0.033 | 0.030 | -1.4711 | 0.1413 | 0.3684 | Increasing |
| GACA-CN::SP135186 | Stomach | STAD-US | 434 | 0 | 0 | 16 | 12 | 0.012 | 0.006 | 0.016 | 0.027 | -1.4726 | 0.1409 | 0.3684 | Increasing |
| OV-US::SP63356 | Ovary | OV-US | 376 | 0 | 98 | 28 | 11 | 0.013 | 0.025 | 0.026 | 0.035 | -1.4665 | 0.1425 | 0.3697 | Increasing |
| LGG-US::SP48504 | Brain | LGG-US | 393 | 0 | 0 | 14 | 5 | 0.000 | 0.012 | 0.014 | 0.017 | -1.4655 | 0.1428 | 0.3697 | Increasing |
| LIRI-JP::SP50173 | Liver | LIRI-JP | 347 | 0 | 45 | 26 | 12 | 0.010 | 0.025 | 0.025 | 0.029 | -1.4656 | 0.1427 | 0.3697 | Increasing |
| HNSC-US::SP32894 | Head&Neck | HNSC-US | 233 | 0 | 53 | 13 | 12 | 0.006 | 0.011 | 0.010 | 0.023 | -1.4673 | 0.1423 | 0.3697 | Increasing |
| PRAD-CA::SP112837 | Prostate | PRAD-US | 421 | 0 | 74 | 21 | 9 | 0.011 | 0.016 | 0.018 | 0.030 | -1.4619 | 0.1438 | 0.3702 | Increasing |
| PACA-AU::SP70465 | Pancreas | PACA-AU | 422 | 0 | 119 | 18 | 11 | 0.015 | 0.005 | 0.012 | 0.027 | -1.4621 | 0.1437 | 0.3702 | Increasing |
| OV-AU::SP101662 | Ovary | OV-AU | 587 | 364 | 96 | 128 | 17 | 0.127 | 0.102 | 0.066 | 0.109 | 1.4623 | 0.1437 | 0.3702 | Decreasing |
| PRAD-UK::SP114935 | Prostate | PRAD-US | 532 | 0 | 0 | 14 | 12 | 0.004 | 0.011 | 0.018 | 0.015 | -1.4566 | 0.1452 | 0.3707 | Increasing |
| PRAD-UK::SP114918 | Prostate | PRAD-US | 652 | 0 | 0 | 14 | 11.5 | 0.004 | 0.011 | 0.018 | 0.015 | -1.4566 | 0.1452 | 0.3707 | Increasing |
| UCEC-US::SP92195 | Endometrium | UCEC-US | 522 | 0 | 88 | 24 | 10 | 0.022 | 0.008 | 0.023 | 0.041 | -1.4582 | 0.1448 | 0.3707 | Increasing |
| KIRC-US::SP39594 | Kidney | KIRC-US | 728 | 0 | 129 | 48 | 8 | 0.040 | 0.032 | 0.031 | 0.074 | -1.4601 | 0.1443 | 0.3707 | Increasing |
| PRAD-CA::SP112869 | Prostate | PRAD-US | 380 | 0 | 36 | 14 | 13 | 0.011 | 0.008 | 0.006 | 0.030 | -1.4566 | 0.1452 | 0.3707 | Increasing |
| MELA-AU::SP124334 | Skin | SKCM-US | 552 | 0 | 91 | 26 | 11 | 0.017 | 0.021 | 0.016 | 0.044 | -1.4521 | 0.1465 | 0.3728 | Increasing |
| COAD-US::SP19295 | CRC | COAD-US | 4361 | 0 | 0 | 11 | 10.5 | 0.006 | 0.005 | 0.013 | 0.017 | -1.4482 | 0.1476 | 0.3728 | Increasing |
| LIRI-JP::SP98904 | Liver | LIRI-JP | 320 | 0 | 30 | 13 | 14 | 0.003 | 0.010 | 0.019 | 0.012 | -1.4491 | 0.1473 | 0.3728 | Increasing |
| BRCA-US::SP5448 | Breast | BRCA-US | 441 | 0 | 103 | 21 | 13 | 0.010 | 0.018 | 0.016 | 0.032 | -1.4482 | 0.1476 | 0.3728 | Increasing |
| LIRI-JP::SP112274 | Liver | LIRI-JP | 162 | 0 | 28 | 13 | 13 | 0.006 | 0.010 | 0.010 | 0.023 | -1.4491 | 0.1473 | 0.3728 | Increasing |
| LUSC-US::SP58101 | Lung | LUSC-US | 1681 | 0 | 0 | 47 | 10 | 0.019 | 0.045 | 0.059 | 0.034 | -1.4492 | 0.1473 | 0.3728 | Increasing |
| HNSC-US::SP31334 | Head&Neck | HNSC-US | 531 | 74 | 65 | 36 | 11 | 0.048 | 0.019 | 0.029 | 0.023 | 1.4380 | 0.1504 | 0.3757 | Decreasing |
| LIRI-JP::SP107063 | Liver | LIRI-JP | 358 | 0 | 62 | 24 | 15 | 0.010 | 0.020 | 0.029 | 0.023 | -1.4393 | 0.1501 | 0.3757 | Increasing |
| PAEN-AU::SP102529 | Pancreas | PAEN-AU | 171 | 0 | 40 | 13 | 14.5 | 0.000 | 0.008 | 0.016 | 0.012 | -1.4397 | 0.1499 | 0.3757 | Increasing |
| THCA-US::SP120767 | Thyroid | THCA-US | 138 | 0 | 29 | 14 | 7.5 | 0.008 | 0.005 | 0.022 | 0.015 | -1.4355 | 0.1511 | 0.3757 | Increasing |
| KIRP-US::SP43808 | Kidney | KIRP-US | 683 | 0 | 119 | 48 | 8 | 0.049 | 0.048 | 0.034 | 0.026 | 1.4413 | 0.1495 | 0.3757 | Decreasing |
| LUSC-US::SP57735 | Lung | LUSC-US | 1547 | 0 | 0 | 35 | 11 | 0.019 | 0.024 | 0.049 | 0.028 | -1.4374 | 0.1506 | 0.3757 | Increasing |
| STAD-US::SP105086 | Stomach | STAD-US | 1588 | 221 | 0 | 68 | 13.5 | 0.046 | 0.062 | 0.046 | 0.090 | -1.4349 | 0.1513 | 0.3757 | Increasing |
| SKCM-US::SP82399 | Skin | SKCM-US | 811 | 0 | 211 | 71 | 13 | 0.085 | 0.054 | 0.044 | 0.061 | 1.4352 | 0.1512 | 0.3757 | Decreasing |
| LIHC-US::SP49114 | Liver | LIHC-US | 1104 | 0 | 242 | 93 | 13 | 0.073 | 0.073 | 0.068 | 0.126 | -1.4369 | 0.1507 | 0.3757 | Increasing |
| RECA-EU::SP103037 | Kidney | RECA-EU | 302 | 0 | 51 | 21 | 8 | 0.007 | 0.016 | 0.023 | 0.020 | -1.4402 | 0.1498 | 0.3757 | Increasing |
| PACA-AU::SP70178 | Pancreas | PACA-AU | 841 | 0 | 0 | 27 | 11 | 0.007 | 0.027 | 0.018 | 0.030 | -1.4306 | 0.1526 | 0.3768 | Increasing |
| SKCM-US::SP82644 | Skin | SKCM-US | 1145 | 0 | 195 | 102 | 17 | 0.072 | 0.085 | 0.085 | 0.117 | -1.4307 | 0.1525 | 0.3768 | Increasing |
| OV-US::SP63716 | Ovary | OV-US | 251 | 0 | 38 | 11 | 13 | 0.007 | 0.008 | 0.007 | 0.023 | -1.4309 | 0.1525 | 0.3768 | Increasing |
| PACA-AU::SP71846 | Pancreas | PACA-AU | 641 | 0 | 0 | 14 | 10 | 0.000 | 0.014 | 0.015 | 0.013 | -1.4283 | 0.1532 | 0.3778 | Increasing |
| PACA-CA::SP125778 | Pancreas | PACA-CA | 2242 | 1325 | 706 | 493 | 13 | 0.392 | 0.447 | 0.321 | 0.366 | 1.4264 | 0.1537 | 0.3785 | Decreasing |
| MELA-AU::SP124346 | Skin | SKCM-US | 558 | 0 | 0 | 16 | 8.5 | 0.007 | 0.008 | 0.028 | 0.011 | -1.4242 | 0.1544 | 0.3794 | Increasing |
| GBM-US::SP27825 | Brain | GBM-US | 520 | 0 | 0 | 12 | 5 | 0.000 | 0.009 | 0.018 | 0.010 | -1.4208 | 0.1554 | 0.3812 | Increasing |
| PRAD-CA::SP112981 | Prostate | PRAD-US | 254 | 0 | 33 | 16 | 9 | 0.000 | 0.021 | 0.012 | 0.020 | -1.4166 | 0.1566 | 0.3829 | Increasing |
| PACA-AU::SP71734 | Pancreas | PACA-AU | 468 | 0 | 0 | 12 | 12 | 0.018 | 0.005 | 0.012 | 0.003 | 1.4168 | 0.1565 | 0.3829 | Decreasing |
| LIRI-JP::SP99089 | Liver | LIRI-JP | 576 | 0 | 67 | 29 | 13.5 | 0.022 | 0.013 | 0.035 | 0.035 | -1.4133 | 0.1576 | 0.3847 | Increasing |
| MELA-AU::SP124458 | Skin | SKCM-US | 1137 | 0 | 367 | 133 | 15 | 0.082 | 0.114 | 0.145 | 0.106 | -1.4119 | 0.1580 | 0.3850 | Increasing |
| MELA-AU::SP124319 | Skin | SKCM-US | 657 | 0 | 173 | 49 | 13 | 0.034 | 0.039 | 0.038 | 0.067 | -1.4069 | 0.1595 | 0.3879 | Increasing |

| | | | | | | | | | | | | | | | |
|---|---|---|---|---|---|---|---|---|---|---|---|---|---|---|---|
| OV-US::SP59860 | Ovary | OV-US | 606 | 83 | 105 | 36 | 13 | 0.020 | 0.025 | 0.049 | 0.029 | -1.4035 | 0.1605 | 0.3897 | Increasing |
| LIRI-JP::SP50101 | Liver | LIRI-JP | 519 | 0 | 95 | 39 | 13 | 0.019 | 0.038 | 0.032 | 0.046 | -1.3979 | 0.1621 | 0.3924 | Increasing |
| PRAD-CA::SP112809 | Prostate | PRAD-US | 468 | 0 | 0 | 11 | 12 | 0.000 | 0.011 | 0.015 | 0.010 | -1.3971 | 0.1624 | 0.3924 | Increasing |
| PRAD-CA::SP112923 | Prostate | PRAD-US | 300 | 0 | 0 | 11 | 10 | 0.004 | 0.008 | 0.012 | 0.015 | -1.3971 | 0.1624 | 0.3924 | Increasing |
| RECA-EU::SP103300 | Kidney | RECA-EU | 1097 | 0 | 182 | 81 | 9 | 0.053 | 0.061 | 0.060 | 0.090 | -1.3939 | 0.1634 | 0.3941 | Increasing |
| BRCA-US::SP8564 | Breast | BRCA-US | 436 | 0 | 87 | 32 | 10 | 0.017 | 0.021 | 0.044 | 0.026 | -1.3833 | 0.1666 | 0.3975 | Increasing |
| PACA-CA::SP125742 | Pancreas | PACA-CA | 602 | 0 | 346 | 58 | 10.5 | 0.057 | 0.042 | 0.056 | 0.025 | 1.3776 | 0.1683 | 0.3975 | Decreasing |
| PACA-AU::SP76136 | Pancreas | PACA-AU | 1095 | 0 | 435 | 72 | 11 | 0.044 | 0.046 | 0.070 | 0.063 | -1.3806 | 0.1674 | 0.3975 | Increasing |
| LIRI-JP::SP107007 | Liver | LIRI-JP | 821 | 0 | 124 | 52 | 12 | 0.048 | 0.020 | 0.057 | 0.064 | -1.3849 | 0.1661 | 0.3975 | Increasing |
| LUAD-US::SP56079 | Lung | LUAD-US | 1503 | 0 | 0 | 19 | 11 | 0.009 | 0.014 | 0.023 | 0.022 | -1.3854 | 0.1659 | 0.3975 | Increasing |
| RECA-EU::SP102839 | Kidney | RECA-EU | 687 | 0 | 118 | 47 | 9 | 0.023 | 0.042 | 0.031 | 0.055 | -1.3819 | 0.1670 | 0.3975 | Increasing |
| THCA-US::SP86929 | Thyroid | THCA-US | 159 | 0 | 49 | 11 | 12.5 | 0.004 | 0.010 | 0.006 | 0.020 | -1.3786 | 0.1680 | 0.3975 | Increasing |
| SKCM-US::SP104056 | Skin | SKCM-US | 2471 | 0 | 602 | 183 | 14 | 0.191 | 0.153 | 0.123 | 0.161 | 1.3801 | 0.1676 | 0.3975 | Decreasing |
| LIRI-JP::SP50137 | Liver | LIRI-JP | 276 | 0 | 40 | 16 | 9.5 | 0.019 | 0.015 | 0.010 | 0.006 | 1.3876 | 0.1653 | 0.3975 | Decreasing |
| KIRP-US::SP106560 | Kidney | KIRP-US | 647 | 0 | 138 | 48 | 10 | 0.035 | 0.032 | 0.047 | 0.057 | -1.3776 | 0.1683 | 0.3975 | Increasing |
| THCA-US::SP86989 | Thyroid | THCA-US | 155 | 0 | 50 | 11 | 18 | 0.004 | 0.005 | 0.019 | 0.010 | -1.3786 | 0.1680 | 0.3975 | Increasing |
| UCEC-US::SP92460 | Endometrium | UCEC-US | 14442 | 0 | 0 | 18 | 10 | 0.006 | 0.018 | 0.017 | 0.023 | -1.3789 | 0.1679 | 0.3975 | Increasing |
| MELA-AU::SP124420 | Skin | SKCM-US | 625 | 0 | 151 | 61 | 15 | 0.065 | 0.057 | 0.038 | 0.044 | 1.3874 | 0.1653 | 0.3975 | Decreasing |
| LIHC-US::SP109384 | Liver | LIHC-US | 348 | 0 | 66 | 30 | 11 | 0.016 | 0.026 | 0.029 | 0.036 | -1.3751 | 0.1691 | 0.3987 | Increasing |
| MALY-DE::SP116630 | Blood | MALY-DE | 1545 | 0 | 0 | 17 | 13 | 0.013 | 0.009 | 0.009 | 0.024 | -1.3738 | 0.1695 | 0.3990 | Increasing |
| SKCM-US::SP82988 | Skin | SKCM-US | 482 | 0 | 0 | 11 | 12 | 0.021 | 0.005 | 0.003 | 0.011 | 1.3708 | 0.1704 | 0.3995 | Decreasing |
| MELA-AU::SP124264 | Skin | SKCM-US | 553 | 0 | 0 | 11 | 13.5 | 0.007 | 0.021 | 0.003 | 0.000 | 1.3708 | 0.1704 | 0.3995 | Decreasing |
| COAD-US::SP18787 | CRC | COAD-US | 2296 | 0 | 0 | 25 | 10.5 | 0.018 | 0.011 | 0.033 | 0.028 | -1.3705 | 0.1705 | 0.3995 | Increasing |
| PACA-CA::SP117441 | Pancreas | PACA-CA | 503 | 0 | 157 | 15 | 10 | 0.008 | 0.011 | 0.006 | 0.022 | -1.3682 | 0.1713 | 0.4005 | Increasing |
| BRCA-US::SP7785 | Breast | BRCA-US | 1252 | 526 | 204 | 161 | 12 | 0.138 | 0.161 | 0.132 | 0.095 | 1.3473 | 0.1779 | 0.4089 | Decreasing |
| ORCA-IN::SP117932 | Head&Neck | ORCA-IN | 617 | 0 | 0 | 16 | 9 | 0.006 | 0.016 | 0.009 | 0.025 | -1.3471 | 0.1779 | 0.4089 | Increasing |
| BRCA-US::SP6730 | Breast | BRCA-US | 513 | 0 | 120 | 26 | 15.5 | 0.014 | 0.018 | 0.031 | 0.026 | -1.3550 | 0.1754 | 0.4089 | Increasing |
| STAD-US::SP84062 | Stomach | STAD-US | 213 | 0 | 35 | 13 | 12 | 0.003 | 0.014 | 0.013 | 0.016 | -1.3440 | 0.1789 | 0.4089 | Increasing |
| GACA-CN::SP135222 | Stomach | STAD-US | 474 | 0 | 0 | 11 | 11 | 0.006 | 0.006 | 0.013 | 0.016 | -1.3477 | 0.1778 | 0.4089 | Increasing |
| OV-AU::SP101648 | Ovary | OV-AU | 517 | 233 | 143 | 111 | 19 | 0.099 | 0.093 | 0.078 | 0.067 | 1.3440 | 0.1790 | 0.4089 | Decreasing |
| PRAD-CA::SP112779 | Prostate | PRAD-US | 426 | 0 | 0 | 13 | 9 | 0.007 | 0.008 | 0.012 | 0.020 | -1.3451 | 0.1786 | 0.4089 | Increasing |
| BRCA-US::SP8157 | Breast | BRCA-US | 215 | 47 | 34 | 22 | 13 | 0.007 | 0.021 | 0.025 | 0.021 | -1.3429 | 0.1793 | 0.4089 | Increasing |
| PRAD-US::SP79971 | Prostate | PRAD-US | 189 | 0 | 23 | 13 | 10 | 0.007 | 0.008 | 0.012 | 0.020 | -1.3451 | 0.1786 | 0.4089 | Increasing |
| KIRP-US::SP109470 | Kidney | KIRP-US | 1533 | 0 | 204 | 64 | 9 | 0.042 | 0.051 | 0.062 | 0.067 | -1.3465 | 0.1781 | 0.4089 | Increasing |
| RECA-EU::SP103288 | Kidney | RECA-EU | 412 | 0 | 55 | 26 | 9 | 0.013 | 0.026 | 0.006 | 0.045 | -1.3428 | 0.1793 | 0.4089 | Increasing |
| MELA-AU::SP124401 | Skin | SKCM-US | 2733 | 0 | 497 | 174 | 15 | 0.178 | 0.153 | 0.110 | 0.156 | 1.3445 | 0.1788 | 0.4089 | Decreasing |
| BRCA-US::SP4557 | Breast | BRCA-US | 311 | 0 | 102 | 24 | 14 | 0.007 | 0.026 | 0.022 | 0.026 | -1.3480 | 0.1777 | 0.4089 | Increasing |
| OV-AU::SP101674 | Ovary | OV-AU | 442 | 225 | 125 | 107 | 17 | 0.111 | 0.075 | 0.069 | 0.083 | 1.3430 | 0.1793 | 0.4089 | Decreasing |
| GACA-CN::SP135240 | Stomach | STAD-US | 699 | 0 | 0 | 11 | 13 | 0.003 | 0.011 | 0.010 | 0.016 | -1.3477 | 0.1778 | 0.4089 | Increasing |
| PACA-CA::SP117290 | Pancreas | PACA-CA | 1255 | 0 | 0 | 22 | 12 | 0.011 | 0.020 | 0.006 | 0.032 | -1.3513 | 0.1766 | 0.4089 | Increasing |
| BRCA-US::SP4535 | Breast | BRCA-US | 180 | 0 | 40 | 18 | 15 | 0.007 | 0.013 | 0.025 | 0.016 | -1.3407 | 0.1800 | 0.4089 | Increasing |
| BRCA-US::SP10150 | Breast | BRCA-US | 463 | 0 | 90 | 20 | 11.5 | 0.010 | 0.011 | 0.031 | 0.016 | -1.3402 | 0.1802 | 0.4089 | Increasing |
| MELA-AU::SP124364 | Skin | SKCM-US | 1791 | 0 | 0 | 51 | 11 | 0.051 | 0.049 | 0.038 | 0.028 | 1.3412 | 0.1799 | 0.4089 | Decreasing |
| PRAD-UK::SP114962 | Prostate | PRAD-US | 2755 | 1436 | 721 | 470 | 13 | 0.499 | 0.327 | 0.414 | 0.382 | 1.3301 | 0.1835 | 0.4151 | Decreasing |
| OV-AU::SP101616 | Ovary | OV-AU | 607 | 160 | 0 | 34 | 12.5 | 0.048 | 0.014 | 0.024 | 0.026 | 1.3302 | 0.1835 | 0.4151 | Decreasing |
| KIRC-US::SP38271 | Kidney | KIRC-US | 791 | 0 | 151 | 52 | 10 | 0.044 | 0.024 | 0.062 | 0.054 | -1.3282 | 0.1841 | 0.4159 | Increasing |
| HNSC-US::SP30071 | Head&Neck | HNSC-US | 812 | 0 | 0 | 21 | 10 | 0.009 | 0.027 | 0.003 | 0.040 | -1.3259 | 0.1849 | 0.4163 | Increasing |
| PACA-AU::SP108049 | Pancreas | PACA-AU | 262 | 0 | 96 | 16 | 12 | 0.000 | 0.016 | 0.018 | 0.013 | -1.3265 | 0.1847 | 0.4163 | Increasing |
| CESC-US::SP13242 | Cervix | CESC-US | 272 | 0 | 0 | 11 | 11 | 0.006 | 0.006 | 0.017 | 0.012 | -1.3236 | 0.1856 | 0.4167 | Increasing |
| UCEC-US::SP91265 | Endometrium | UCEC-US | 765 | 0 | 105 | 33 | 11 | 0.025 | 0.021 | 0.030 | 0.046 | -1.3242 | 0.1855 | 0.4167 | Increasing |

| | | | | | | | | | | | | | | | |
|---|---|---|---|---|---|---|---|---|---|---|---|---|---|---|---|
| LUSC-US::SP57450 | Lung | LUSC-US | 859 | 0 | 141 | 33 | 13 | 0.041 | 0.026 | 0.016 | 0.028 | 1.3191 | 0.1871 | 0.4194 | Decreasing |
| PRAD-CA::SP112789 | Prostate | PRAD-US | 618 | 177 | 94 | 76 | 15 | 0.101 | 0.045 | 0.058 | 0.065 | 1.3138 | 0.1889 | 0.4221 | Decreasing |
| LGG-US::SP48189 | Brain | LGG-US | 475 | 0 | 0 | 12 | 6 | 0.006 | 0.005 | 0.017 | 0.013 | -1.3143 | 0.1887 | 0.4221 | Increasing |
| LUSC-US::SP58245 | Lung | LUSC-US | 1995 | 0 | 0 | 32 | 13 | 0.026 | 0.021 | 0.023 | 0.051 | -1.3101 | 0.1902 | 0.4242 | Increasing |
| PRAD-CA::SP102620 | Prostate | PRAD-US | 305 | 0 | 32 | 15 | 11 | 0.007 | 0.005 | 0.027 | 0.010 | -1.3081 | 0.1908 | 0.4250 | Increasing |
| LIRI-JP::SP98959 | Liver | LIRI-JP | 295 | 0 | 39 | 12 | 10 | 0.006 | 0.008 | 0.013 | 0.017 | -1.3041 | 0.1922 | 0.4269 | Increasing |
| PACA-CA::SP125689 | Pancreas | PACA-CA | 1124 | 0 | 254 | 46 | 12.5 | 0.026 | 0.025 | 0.051 | 0.038 | -1.3023 | 0.1928 | 0.4269 | Increasing |
| MELA-AU::SP124329 | Skin | SKCM-US | 223 | 0 | 52 | 17 | 12.5 | 0.010 | 0.010 | 0.019 | 0.022 | -1.3032 | 0.1925 | 0.4269 | Increasing |
| BTCA-SG::SP117017 | Biliary | BTCA-SG | 387 | 0 | 74 | 14 | 10 | 0.009 | 0.007 | 0.024 | 0.017 | -1.3028 | 0.1926 | 0.4269 | Increasing |
| LIRI-JP::SP107034 | Liver | LIRI-JP | 321 | 0 | 47 | 21 | 14 | 0.010 | 0.015 | 0.029 | 0.017 | -1.2922 | 0.1963 | 0.4339 | Increasing |
| UCEC-US::SP95550 | Endometrium | UCEC-US | 647 | 111 | 0 | 31 | 10 | 0.019 | 0.026 | 0.027 | 0.041 | -1.2888 | 0.1975 | 0.4358 | Increasing |
| PACA-AU::SP69573 | Pancreas | PACA-AU | 310 | 0 | 84 | 12 | 14 | 0.004 | 0.008 | 0.012 | 0.013 | -1.2838 | 0.1992 | 0.4383 | Increasing |
| HNSC-US::SP30083 | Head&Neck | HNSC-US | 340 | 0 | 49 | 15 | 11 | 0.021 | 0.011 | 0.007 | 0.012 | 1.2841 | 0.1991 | 0.4383 | Decreasing |
| LIRI-JP::SP50159 | Liver | LIRI-JP | 738 | 0 | 149 | 54 | 13 | 0.035 | 0.050 | 0.032 | 0.075 | -1.2814 | 0.2000 | 0.4395 | Increasing |
| OV-AU::SP101580 | Ovary | OV-AU | 482 | 148 | 145 | 68 | 20 | 0.061 | 0.059 | 0.048 | 0.036 | 1.2772 | 0.2015 | 0.4421 | Decreasing |
| RECA-EU::SP103685 | Kidney | RECA-EU | 689 | 0 | 87 | 42 | 8 | 0.030 | 0.031 | 0.023 | 0.060 | -1.2705 | 0.2039 | 0.4460 | Increasing |
| RECA-EU::SP103595 | Kidney | RECA-EU | 552 | 0 | 94 | 42 | 10 | 0.023 | 0.031 | 0.040 | 0.040 | -1.2705 | 0.2039 | 0.4460 | Increasing |
| LIRI-JP::SP107097 | Liver | LIRI-JP | 535 | 0 | 78 | 31 | 10 | 0.022 | 0.023 | 0.022 | 0.046 | -1.2646 | 0.2060 | 0.4492 | Increasing |
| LIRI-JP::SP99253 | Liver | LIRI-JP | 685 | 0 | 91 | 31 | 11 | 0.022 | 0.020 | 0.029 | 0.041 | -1.2646 | 0.2060 | 0.4492 | Increasing |
| RECA-EU::SP103575 | Kidney | RECA-EU | 651 | 0 | 86 | 29 | 11 | 0.010 | 0.031 | 0.017 | 0.035 | -1.2621 | 0.2069 | 0.4505 | Increasing |
| PACA-AU::SP76675 | Pancreas | PACA-AU | 1380 | 688 | 0 | 187 | 16 | 0.194 | 0.131 | 0.117 | 0.154 | 1.2545 | 0.2096 | 0.4554 | Decreasing |
| PACA-AU::SP70780 | Pancreas | PACA-AU | 714 | 0 | 0 | 15 | 9 | 0.011 | 0.008 | 0.006 | 0.023 | -1.2542 | 0.2098 | 0.4554 | Increasing |
| LINC-JP::SP98861 | Liver | LIHC-US | 773 | 83 | 102 | 53 | 11 | 0.038 | 0.044 | 0.039 | 0.072 | -1.2483 | 0.2119 | 0.4586 | Increasing |
| LUSC-US::SP56607 | Lung | LUSC-US | 921 | 0 | 0 | 28 | 12 | 0.016 | 0.018 | 0.039 | 0.023 | -1.2486 | 0.2118 | 0.4586 | Increasing |
| BRCA-US::SP8229 | Breast | BRCA-US | 249 | 0 | 73 | 11 | 16 | 0.000 | 0.013 | 0.013 | 0.011 | -1.2452 | 0.2131 | 0.4604 | Increasing |
| LIHC-US::SP98192 | Liver | LIHC-US | 452 | 0 | 0 | 12 | 19 | 0.019 | 0.008 | 0.003 | 0.012 | 1.2421 | 0.2142 | 0.4607 | Decreasing |
| PACA-AU::SP108020 | Pancreas | PACA-AU | 542 | 158 | 0 | 18 | 17.5 | 0.007 | 0.011 | 0.020 | 0.017 | -1.2416 | 0.2144 | 0.4607 | Increasing |
| RECA-EU::SP102747 | Kidney | RECA-EU | 210 | 0 | 35 | 11 | 9.5 | 0.000 | 0.009 | 0.017 | 0.005 | -1.2420 | 0.2142 | 0.4607 | Increasing |
| BRCA-UK::SP2155 | Breast | BRCA-US | 793 | 544 | 0 | 132 | 14 | 0.090 | 0.100 | 0.157 | 0.095 | -1.2414 | 0.2144 | 0.4607 | Increasing |
| LUSC-US::SP56537 | Lung | LUSC-US | 1741 | 0 | 0 | 38 | 11 | 0.022 | 0.029 | 0.046 | 0.034 | -1.2388 | 0.2154 | 0.4607 | Increasing |
| PACA-CA::SP125745 | Pancreas | PACA-CA | 401 | 0 | 0 | 13 | 12.5 | 0.000 | 0.011 | 0.017 | 0.010 | -1.2391 | 0.2153 | 0.4607 | Increasing |
| PACA-AU::SP107784 | Pancreas | PACA-AU | 1339 | 0 | 0 | 21 | 12 | 0.011 | 0.014 | 0.018 | 0.023 | -1.2390 | 0.2154 | 0.4607 | Increasing |
| LIRI-JP::SP98925 | Liver | LIRI-JP | 343 | 0 | 65 | 24 | 13 | 0.016 | 0.013 | 0.032 | 0.023 | -1.2368 | 0.2162 | 0.4609 | Increasing |
| RECA-EU::SP103045 | Kidney | RECA-EU | 778 | 0 | 119 | 47 | 9 | 0.026 | 0.038 | 0.037 | 0.050 | -1.2371 | 0.2161 | 0.4609 | Increasing |
| RECA-EU::SP103826 | Kidney | RECA-EU | 605 | 0 | 107 | 54 | 9 | 0.026 | 0.045 | 0.051 | 0.045 | -1.2282 | 0.2194 | 0.4657 | Increasing |
| LIRI-JP::SP99007 | Liver | LIRI-JP | 386 | 0 | 61 | 29 | 11 | 0.016 | 0.020 | 0.038 | 0.023 | -1.2291 | 0.2191 | 0.4657 | Increasing |
| BLCA-US::SP1009 | Bladder | BLCA-US | 509 | 0 | 61 | 18 | 11 | 0.006 | 0.018 | 0.020 | 0.018 | -1.2297 | 0.2188 | 0.4657 | Increasing |
| PRAD-CA::SP112961 | Prostate | PRAD-US | 539 | 127 | 0 | 34 | 10 | 0.045 | 0.029 | 0.012 | 0.035 | 1.2251 | 0.2206 | 0.4669 | Decreasing |
| BRCA-UK::SP116367 | Breast | BRCA-US | 396 | 0 | 134 | 43 | 15 | 0.031 | 0.032 | 0.038 | 0.053 | -1.2250 | 0.2206 | 0.4669 | Increasing |
| BRCA-US::SP4523 | Breast | BRCA-US | 315 | 0 | 77 | 15 | 17 | 0.000 | 0.021 | 0.013 | 0.016 | -1.2239 | 0.2210 | 0.4671 | Increasing |
| KIRP-US::SP109544 | Kidney | KIRP-US | 1047 | 0 | 127 | 46 | 10 | 0.035 | 0.029 | 0.047 | 0.052 | -1.2226 | 0.2215 | 0.4675 | Increasing |
| KIRC-US::SP34191 | Kidney | KIRC-US | 480 | 0 | 66 | 31 | 9 | 0.022 | 0.016 | 0.040 | 0.029 | -1.2208 | 0.2222 | 0.4682 | Increasing |
| UCEC-US::SP92787 | Endometrium | UCEC-US | 295 | 0 | 68 | 27 | 9 | 0.016 | 0.023 | 0.023 | 0.035 | -1.2150 | 0.2244 | 0.4716 | Increasing |
| COAD-US::SP21057 | CRC | COAD-US | 2157 | 0 | 0 | 27 | 9.5 | 0.018 | 0.019 | 0.026 | 0.034 | -1.2148 | 0.2244 | 0.4716 | Increasing |
| LIRI-JP::SP99153 | Liver | LIRI-JP | 270 | 0 | 24 | 15 | 9 | 0.010 | 0.010 | 0.013 | 0.023 | -1.2019 | 0.2294 | 0.4814 | Increasing |
| PACA-CA::SP125726 | Pancreas | PACA-CA | 661 | 0 | 0 | 16 | 11.5 | 0.004 | 0.014 | 0.014 | 0.016 | -1.1949 | 0.2321 | 0.4864 | Increasing |
| RECA-EU::SP103100 | Kidney | RECA-EU | 509 | 0 | 75 | 32 | 11 | 0.016 | 0.024 | 0.031 | 0.030 | -1.1926 | 0.2330 | 0.4869 | Increasing |
| RECA-EU::SP103416 | Kidney | RECA-EU | 487 | 0 | 75 | 32 | 10.5 | 0.023 | 0.014 | 0.037 | 0.030 | -1.1926 | 0.2330 | 0.4869 | Increasing |
| KIRC-US::SP38044 | Kidney | KIRC-US | 893 | 0 | 166 | 67 | 9 | 0.073 | 0.054 | 0.056 | 0.044 | 1.1904 | 0.2339 | 0.4879 | Decreasing |
| COAD-US::SP19606 | CRC | COAD-US | 1014 | 0 | 0 | 11 | 11 | 0.009 | 0.016 | 0.007 | 0.000 | 1.1887 | 0.2346 | 0.4880 | Decreasing |

| | | | | | | | | | | | | | | | |
|---|---|---|---|---|---|---|---|---|---|---|---|---|---|---|---|
| OV-AU::SP101523 | Ovary | OV-AU | 687 | 383 | 276 | 176 | 21 | 0.175 | 0.123 | 0.117 | 0.145 | 1.1891 | 0.2344 | 0.4880 | Decreasing |
| PACA-AU::SP73319 | Pancreas | PACA-AU | 379 | 0 | 111 | 18 | 12 | 0.022 | 0.014 | 0.012 | 0.010 | 1.1840 | 0.2364 | 0.4911 | Decreasing |
| COAD-US::SP96112 | CRC | COAD-US | 773 | 0 | 0 | 13 | 8 | 0.006 | 0.005 | 0.026 | 0.006 | -1.1823 | 0.2371 | 0.4912 | Increasing |
| LIRI-JP::SP50155 | Liver | LIRI-JP | 992 | 0 | 145 | 56 | 14 | 0.032 | 0.053 | 0.048 | 0.058 | -1.1822 | 0.2371 | 0.4912 | Increasing |
| LUSC-US::SP58326 | Lung | LUSC-US | 3142 | 0 | 0 | 34 | 12 | 0.038 | 0.029 | 0.023 | 0.023 | 1.1810 | 0.2376 | 0.4914 | Decreasing |
| OV-AU::SP102096 | Ovary | OV-AU | 390 | 0 | 98 | 30 | 11 | 0.016 | 0.025 | 0.021 | 0.036 | -1.1788 | 0.2385 | 0.4917 | Increasing |
| BRCA-US::SP2801 | Breast | BRCA-US | 476 | 0 | 155 | 28 | 12 | 0.014 | 0.029 | 0.019 | 0.037 | -1.1781 | 0.2388 | 0.4917 | Increasing |
| KIRP-US::SP43664 | Kidney | KIRP-US | 1731 | 0 | 332 | 129 | 10 | 0.101 | 0.094 | 0.128 | 0.124 | -1.1782 | 0.2387 | 0.4917 | Increasing |
| SKCM-US::SP103894 | Skin | SKCM-US | 4737 | 0 | 0 | 114 | 14 | 0.079 | 0.106 | 0.085 | 0.128 | -1.1749 | 0.2400 | 0.4937 | Increasing |
| LIRI-JP::SP99269 | Liver | LIRI-JP | 484 | 0 | 72 | 35 | 12 | 0.019 | 0.028 | 0.041 | 0.029 | -1.1652 | 0.2439 | 0.4942 | Increasing |
| PACA-AU::SP70266 | Pancreas | PACA-AU | 409 | 0 | 94 | 17 | 10 | 0.015 | 0.005 | 0.012 | 0.023 | -1.1688 | 0.2425 | 0.4942 | Increasing |
| LIHC-US::SP49247 | Liver | LIHC-US | 908 | 0 | 113 | 41 | 12 | 0.025 | 0.037 | 0.035 | 0.048 | -1.1649 | 0.2440 | 0.4942 | Increasing |
| PACA-CA::SP117309 | Pancreas | PACA-CA | 796 | 0 | 0 | 19 | 10.5 | 0.015 | 0.008 | 0.011 | 0.025 | -1.1680 | 0.2428 | 0.4942 | Increasing |
| LIHC-US::SP49541 | Liver | LIHC-US | 841 | 90 | 94 | 62 | 14 | 0.054 | 0.060 | 0.061 | 0.018 | 1.1703 | 0.2419 | 0.4942 | Decreasing |
| LIRI-JP::SP112245 | Liver | LIRI-JP | 296 | 0 | 48 | 20 | 9 | 0.010 | 0.015 | 0.025 | 0.017 | -1.1660 | 0.2436 | 0.4942 | Increasing |
| HNSC-US::SP29940 | Head&Neck | HNSC-US | 384 | 0 | 0 | 11 | 10 | 0.006 | 0.005 | 0.016 | 0.012 | -1.1692 | 0.2423 | 0.4942 | Increasing |
| MELA-AU::SP124418 | Skin | SKCM-US | 630 | 0 | 128 | 34 | 11 | 0.024 | 0.026 | 0.028 | 0.044 | -1.1650 | 0.2440 | 0.4942 | Increasing |
| RECA-EU::SP103535 | Kidney | RECA-EU | 1387 | 0 | 172 | 74 | 10 | 0.046 | 0.054 | 0.068 | 0.065 | -1.1641 | 0.2444 | 0.4942 | Increasing |
| LIRI-JP::SP99201 | Liver | LIRI-JP | 368 | 0 | 61 | 20 | 13 | 0.010 | 0.018 | 0.019 | 0.023 | -1.1660 | 0.2436 | 0.4942 | Increasing |
| OV-AU::SP101700 | Ovary | OV-AU | 607 | 272 | 293 | 164 | 21 | 0.153 | 0.111 | 0.147 | 0.093 | 1.1691 | 0.2424 | 0.4942 | Decreasing |
| RECA-EU::SP103455 | Kidney | RECA-EU | 994 | 0 | 156 | 62 | 10 | 0.036 | 0.049 | 0.051 | 0.060 | -1.1712 | 0.2415 | 0.4942 | Increasing |
| OV-AU::SP102084 | Ovary | OV-AU | 538 | 138 | 69 | 43 | 14 | 0.038 | 0.039 | 0.030 | 0.021 | 1.1608 | 0.2457 | 0.4945 | Decreasing |
| SKCM-US::SP82471 | Skin | SKCM-US | 466 | 0 | 0 | 12 | 10 | 0.007 | 0.008 | 0.013 | 0.017 | -1.1621 | 0.2452 | 0.4945 | Increasing |
| UCEC-US::SP94661 | Endometrium | UCEC-US | 501 | 0 | 61 | 11 | 10 | 0.003 | 0.010 | 0.013 | 0.012 | -1.1604 | 0.2459 | 0.4945 | Increasing |
| HNSC-US::SP30077 | Head&Neck | HNSC-US | 1099 | 155 | 0 | 64 | 13.5 | 0.076 | 0.044 | 0.042 | 0.058 | 1.1605 | 0.2459 | 0.4945 | Decreasing |
| MALY-DE::SP116686 | Blood | MALY-DE | 370 | 64 | 0 | 12 | 9 | 0.000 | 0.011 | 0.006 | 0.015 | -1.1542 | 0.2484 | 0.4975 | Increasing |
| LIRI-JP::SP99041 | Liver | LIRI-JP | 555 | 0 | 76 | 25 | 13 | 0.013 | 0.020 | 0.029 | 0.023 | -1.1548 | 0.2482 | 0.4975 | Increasing |
| HNSC-US::SP32014 | Head&Neck | HNSC-US | 434 | 63 | 0 | 23 | 15 | 0.015 | 0.011 | 0.036 | 0.017 | -1.1557 | 0.2478 | 0.4975 | Increasing |
| MELA-AU::SP124399 | Skin | SKCM-US | 998 | 0 | 226 | 81 | 11 | 0.065 | 0.060 | 0.069 | 0.094 | -1.1522 | 0.2492 | 0.4980 | Increasing |
| PACA-CA::SP125776 | Pancreas | PACA-CA | 577 | 0 | 124 | 22 | 12.5 | 0.011 | 0.014 | 0.020 | 0.022 | -1.1519 | 0.2493 | 0.4980 | Increasing |
| COAD-US::SP96122 | CRC | COAD-US | 2485 | 0 | 0 | 18 | 12 | 0.012 | 0.014 | 0.013 | 0.028 | -1.1446 | 0.2524 | 0.5034 | Increasing |
| LIRI-JP::SP112213 | Liver | LIRI-JP | 678 | 0 | 104 | 38 | 11 | 0.022 | 0.035 | 0.029 | 0.046 | -1.1405 | 0.2541 | 0.5060 | Increasing |
| KIRC-US::SP34431 | Kidney | KIRC-US | 471 | 0 | 59 | 24 | 9 | 0.011 | 0.024 | 0.019 | 0.029 | -1.1319 | 0.2577 | 0.5111 | Increasing |
| LIRI-JP::SP99237 | Liver | LIRI-JP | 448 | 0 | 88 | 28 | 12 | 0.029 | 0.025 | 0.022 | 0.012 | 1.1326 | 0.2574 | 0.5111 | Decreasing |
| BRCA-US::SP10944 | Breast | BRCA-US | 511 | 0 | 139 | 33 | 11 | 0.028 | 0.021 | 0.025 | 0.048 | -1.1327 | 0.2573 | 0.5111 | Increasing |
| LUAD-US::SP55142 | Lung | LUAD-US | 5834 | 716 | 0 | 168 | 13 | 0.180 | 0.117 | 0.140 | 0.136 | 1.1255 | 0.2604 | 0.5132 | Decreasing |
| LIRI-JP::SP112186 | Liver | LIRI-JP | 574 | 0 | 87 | 33 | 16 | 0.029 | 0.018 | 0.029 | 0.046 | -1.1265 | 0.2600 | 0.5132 | Increasing |
| PACA-CA::SP125686 | Pancreas | PACA-CA | 571 | 0 | 158 | 23 | 12 | 0.023 | 0.014 | 0.031 | 0.003 | 1.1263 | 0.2600 | 0.5132 | Decreasing |
| KIRP-US::SP113926 | Kidney | KIRP-US | 318 | 0 | 76 | 43 | 9 | 0.021 | 0.048 | 0.028 | 0.052 | -1.1270 | 0.2597 | 0.5132 | Increasing |
| BRCA-UK::SP116373 | Breast | BRCA-US | 586 | 283 | 183 | 138 | 19 | 0.103 | 0.116 | 0.116 | 0.143 | -1.1252 | 0.2605 | 0.5132 | Increasing |
| PACA-CA::SP125756 | Pancreas | PACA-CA | 1191 | 424 | 383 | 192 | 18 | 0.177 | 0.151 | 0.127 | 0.146 | 1.1242 | 0.2609 | 0.5133 | Decreasing |
| PAEN-AU::SP117043 | Pancreas | PAEN-AU | 154 | 0 | 27 | 12 | 13 | 0.000 | 0.008 | 0.016 | 0.009 | -1.1230 | 0.2614 | 0.5137 | Increasing |
| BRCA-US::SP5980 | Breast | BRCA-US | 1116 | 525 | 591 | 237 | 18 | 0.234 | 0.200 | 0.170 | 0.207 | 1.1212 | 0.2622 | 0.5145 | Decreasing |
| LINC-JP::SP98871 | Liver | LIHC-US | 407 | 0 | 59 | 26 | 15 | 0.016 | 0.018 | 0.032 | 0.024 | -1.1121 | 0.2661 | 0.5179 | Increasing |
| KIRC-US::SP40736 | Kidney | KIRC-US | 850 | 0 | 143 | 49 | 9 | 0.036 | 0.038 | 0.040 | 0.059 | -1.1125 | 0.2659 | 0.5179 | Increasing |
| OV-US::SP59938 | Ovary | OV-US | 527 | 291 | 152 | 113 | 20 | 0.098 | 0.109 | 0.095 | 0.064 | 1.1123 | 0.2660 | 0.5179 | Decreasing |
| HNSC-US::SP30113 | Head&Neck | HNSC-US | 3593 | 0 | 0 | 52 | 12 | 0.048 | 0.052 | 0.039 | 0.029 | 1.1135 | 0.2655 | 0.5179 | Decreasing |
| HNSC-US::SP30143 | Head&Neck | HNSC-US | 329 | 0 | 54 | 21 | 9 | 0.006 | 0.027 | 0.016 | 0.023 | -1.1136 | 0.2655 | 0.5179 | Increasing |
| COAD-US::SP96116 | CRC | COAD-US | 1355 | 0 | 0 | 17 | 11.5 | 0.021 | 0.016 | 0.003 | 0.017 | 1.1135 | 0.2655 | 0.5179 | Decreasing |
| MELA-AU::SP124415 | Skin | SKCM-US | 2246 | 0 | 313 | 87 | 13 | 0.089 | 0.075 | 0.063 | 0.067 | 1.1096 | 0.2672 | 0.5193 | Decreasing |

| | | | | | | | | | | | | | | | |
|---|---|---|---|---|---|---|---|---|---|---|---|---|---|---|---|
| BRCA-US::SP11948 | Breast | BRCA-US | 909 | 365 | 486 | 220 | 18.5 | 0.220 | 0.179 | 0.167 | 0.185 | 1.1073 | 0.2682 | 0.5198 | Decreasing |
| OV-AU::SP101678 | Ovary | OV-AU | 906 | 309 | 334 | 201 | 20 | 0.159 | 0.179 | 0.138 | 0.135 | 1.1079 | 0.2679 | 0.5198 | Decreasing |
| RECA-EU::SP103245 | Kidney | RECA-EU | 892 | 0 | 163 | 68 | 10 | 0.040 | 0.054 | 0.060 | 0.060 | -1.1061 | 0.2687 | 0.5201 | Increasing |
| OV-AU::SP101881 | Ovary | OV-AU | 758 | 339 | 279 | 150 | 19 | 0.150 | 0.102 | 0.105 | 0.119 | 1.1051 | 0.2691 | 0.5203 | Decreasing |
| LIRI-JP::SP99305 | Liver | LIRI-JP | 268 | 0 | 42 | 15 | 17 | 0.016 | 0.013 | 0.016 | 0.000 | 1.1034 | 0.2699 | 0.5207 | Decreasing |
| PRAD-CA::SP112903 | Prostate | PRAD-US | 254 | 0 | 32 | 11 | 9 | 0.011 | 0.003 | 0.009 | 0.020 | -1.1004 | 0.2711 | 0.5207 | Increasing |
| OV-AU::SP101584 | Ovary | OV-AU | 424 | 50 | 68 | 31 | 11.5 | 0.022 | 0.014 | 0.039 | 0.026 | -1.1029 | 0.2701 | 0.5207 | Increasing |
| PACA-CA::SP125741 | Pancreas | PACA-CA | 819 | 0 | 0 | 18 | 16.5 | 0.008 | 0.008 | 0.025 | 0.013 | -1.1021 | 0.2704 | 0.5207 | Increasing |
| STAD-US::SP104984 | Stomach | STAD-US | 754 | 0 | 0 | 15 | 11 | 0.009 | 0.008 | 0.020 | 0.016 | -1.1005 | 0.2711 | 0.5207 | Increasing |
| BLCA-US::SP1724 | Bladder | BLCA-US | 295 | 0 | 31 | 12 | 17 | 0.006 | 0.013 | 0.003 | 0.024 | -1.0988 | 0.2718 | 0.5214 | Increasing |
| PACA-CA::SP125749 | Pancreas | PACA-CA | 306 | 0 | 46 | 11 | 13.5 | 0.008 | 0.006 | 0.006 | 0.016 | -1.0965 | 0.2729 | 0.5219 | Increasing |
| OV-AU::SP102074 | Ovary | OV-AU | 234 | 34 | 51 | 24 | 11 | 0.006 | 0.027 | 0.015 | 0.026 | -1.0950 | 0.2735 | 0.5219 | Increasing |
| LUSC-US::SP57619 | Lung | LUSC-US | 1475 | 0 | 209 | 46 | 15 | 0.048 | 0.045 | 0.023 | 0.040 | 1.0933 | 0.2743 | 0.5219 | Decreasing |
| PACA-CA::SP117215 | Pancreas | PACA-CA | 456 | 0 | 148 | 13 | 14 | 0.015 | 0.014 | 0.003 | 0.010 | 1.0948 | 0.2736 | 0.5219 | Decreasing |
| MELA-AU::SP124291 | Skin | SKCM-US | 306 | 0 | 55 | 17 | 13 | 0.017 | 0.018 | 0.013 | 0.006 | 1.0939 | 0.2740 | 0.5219 | Decreasing |
| BRCA-US::SP5636 | Breast | BRCA-US | 271 | 0 | 63 | 14 | 9 | 0.003 | 0.016 | 0.013 | 0.016 | -1.0951 | 0.2735 | 0.5219 | Increasing |
| PACA-CA::SP125696 | Pancreas | PACA-CA | 619 | 0 | 0 | 21 | 9.5 | 0.004 | 0.017 | 0.028 | 0.013 | -1.0884 | 0.2764 | 0.5240 | Increasing |
| PACA-CA::SP117908 | Pancreas | PACA-CA | 498 | 0 | 156 | 21 | 12 | 0.011 | 0.020 | 0.006 | 0.029 | -1.0884 | 0.2764 | 0.5240 | Increasing |
| BLCA-US::SP1377 | Bladder | BLCA-US | 385 | 0 | 0 | 17 | 14 | 0.009 | 0.013 | 0.020 | 0.018 | -1.0888 | 0.2762 | 0.5240 | Increasing |
| GACA-CN::SP135431 | Stomach | STAD-US | 1645 | 0 | 0 | 22 | 9.5 | 0.009 | 0.022 | 0.023 | 0.021 | -1.0871 | 0.2770 | 0.5243 | Increasing |
| BRCA-UK::SP2145 | Breast | BRCA-US | 783 | 542 | 179 | 163 | 14.5 | 0.155 | 0.145 | 0.123 | 0.127 | 1.0847 | 0.2781 | 0.5257 | Decreasing |
| MALY-DE::SP59312 | Blood | MALY-DE | 1298 | 0 | 0 | 16 | 11 | 0.000 | 0.011 | 0.020 | 0.012 | -1.0812 | 0.2796 | 0.5279 | Increasing |
| KIRP-US::SP106577 | Kidney | KIRP-US | 1507 | 0 | 196 | 81 | 9 | 0.059 | 0.062 | 0.081 | 0.077 | -1.0775 | 0.2813 | 0.5304 | Increasing |
| LIRI-JP::SP50145 | Liver | LIRI-JP | 426 | 0 | 67 | 21 | 11 | 0.016 | 0.015 | 0.013 | 0.035 | -1.0757 | 0.2820 | 0.5311 | Increasing |
| PRAD-CA::SP112901 | Prostate | PRAD-US | 217 | 0 | 31 | 13 | 11 | 0.011 | 0.005 | 0.012 | 0.020 | -1.0723 | 0.2836 | 0.5322 | Increasing |
| PACA-AU::SP108033 | Pancreas | PACA-AU | 555 | 0 | 0 | 14 | 15 | 0.018 | 0.005 | 0.018 | 0.003 | 1.0720 | 0.2837 | 0.5322 | Decreasing |
| PRAD-US::SP79958 | Prostate | PRAD-US | 361 | 0 | 40 | 13 | 19 | 0.004 | 0.016 | 0.006 | 0.020 | -1.0723 | 0.2836 | 0.5322 | Increasing |
| MELA-AU::SP124439 | Skin | SKCM-US | 1230 | 0 | 285 | 59 | 13 | 0.072 | 0.039 | 0.044 | 0.050 | 1.0691 | 0.2850 | 0.5333 | Decreasing |
| BRCA-US::SP9648 | Breast | BRCA-US | 1037 | 390 | 376 | 192 | 19 | 0.196 | 0.156 | 0.138 | 0.170 | 1.0692 | 0.2850 | 0.5333 | Decreasing |
| OV-AU::SP101592 | Ovary | OV-AU | 330 | 44 | 45 | 22 | 12 | 0.022 | 0.007 | 0.015 | 0.036 | -1.0661 | 0.2864 | 0.5344 | Increasing |
| OV-AU::SP102133 | Ovary | OV-AU | 20838 | 0 | 0 | 22 | 10 | 0.006 | 0.020 | 0.024 | 0.016 | -1.0661 | 0.2864 | 0.5344 | Increasing |
| CESC-US::SP107624 | Cervix | CESC-US | 169 | 45 | 0 | 16 | 16 | 0.011 | 0.011 | 0.013 | 0.025 | -1.0640 | 0.2873 | 0.5355 | Increasing |
| PACA-AU::SP71758 | Pancreas | PACA-AU | 331 | 0 | 224 | 40 | 9 | 0.022 | 0.027 | 0.041 | 0.033 | -1.0619 | 0.2883 | 0.5366 | Increasing |
| RECA-EU::SP103844 | Kidney | RECA-EU | 440 | 0 | 63 | 36 | 8 | 0.023 | 0.028 | 0.023 | 0.045 | -1.0580 | 0.2901 | 0.5385 | Increasing |
| READ-US::SP82103 | CRC | READ-US | 1505 | 0 | 0 | 12 | 9 | 0.003 | 0.013 | 0.013 | 0.012 | -1.0580 | 0.2901 | 0.5385 | Increasing |
| PRAD-UK::SP114904 | Prostate | PRAD-US | 846 | 0 | 0 | 15 | 13 | 0.007 | 0.011 | 0.018 | 0.015 | -1.0541 | 0.2918 | 0.5404 | Increasing |
| PRAD-US::SP79907 | Prostate | PRAD-US | 266 | 0 | 41 | 15 | 12 | 0.004 | 0.016 | 0.015 | 0.015 | -1.0541 | 0.2918 | 0.5404 | Increasing |
| COAD-US::SP119755 | CRC | COAD-US | 2461 | 0 | 0 | 19 | 12 | 0.018 | 0.003 | 0.026 | 0.023 | -1.0521 | 0.2928 | 0.5408 | Increasing |
| UCEC-US::SP93652 | Endometrium | UCEC-US | 454 | 0 | 87 | 19 | 13 | 0.019 | 0.005 | 0.020 | 0.029 | -1.0527 | 0.2925 | 0.5408 | Increasing |
| PACA-AU::SP110779 | Pancreas | PACA-AU | 644 | 0 | 134 | 11 | 11 | 0.011 | 0.011 | 0.009 | 0.003 | 1.0509 | 0.2933 | 0.5411 | Decreasing |
| BTCA-SG::SP117655 | Biliary | BTCA-SG | 1499 | 0 | 0 | 19 | 11 | 0.009 | 0.023 | 0.020 | 0.022 | -1.0449 | 0.2961 | 0.5454 | Increasing |
| LUSC-US::SP57933 | Lung | LUSC-US | 3434 | 1088 | 454 | 405 | 12 | 0.386 | 0.322 | 0.342 | 0.323 | 1.0442 | 0.2964 | 0.5454 | Decreasing |
| HNSC-US::SP30093 | Head&Neck | HNSC-US | 445 | 0 | 0 | 12 | 10 | 0.009 | 0.008 | 0.007 | 0.023 | -1.0424 | 0.2972 | 0.5455 | Increasing |
| HNSC-US::SP31606 | Head&Neck | HNSC-US | 323 | 0 | 0 | 12 | 10 | 0.009 | 0.008 | 0.007 | 0.023 | -1.0424 | 0.2972 | 0.5455 | Increasing |
| OV-AU::SP102168 | Ovary | OV-AU | 266 | 0 | 48 | 20 | 13 | 0.013 | 0.014 | 0.015 | 0.026 | -1.0367 | 0.2999 | 0.5484 | Increasing |
| MALY-DE::SP116657 | Blood | MALY-DE | 1994 | 0 | 0 | 33 | 12 | 0.032 | 0.022 | 0.009 | 0.045 | -1.0381 | 0.2992 | 0.5484 | Decreasing |
| PRAD-CA::SP112983 | Prostate | PRAD-US | 456 | 0 | 65 | 19 | 8 | 0.011 | 0.016 | 0.015 | 0.025 | -1.0359 | 0.3002 | 0.5484 | Increasing |
| UCEC-US::SP94540 | Endometrium | UCEC-US | 583 | 0 | 164 | 14 | 11 | 0.016 | 0.013 | 0.010 | 0.006 | 1.0358 | 0.3003 | 0.5484 | Decreasing |
| PACA-CA::SP125695 | Pancreas | PACA-CA | 379 | 0 | 101 | 17 | 11 | 0.000 | 0.020 | 0.017 | 0.013 | -1.0332 | 0.3015 | 0.5499 | Increasing |
| BTCA-SG::SP117425 | Biliary | BTCA-SG | 597 | 0 | 0 | 17 | 11 | 0.019 | 0.020 | 0.016 | 0.006 | 1.0314 | 0.3023 | 0.5508 | Decreasing |

| | | | | | | | | | | | | | | | |
|---|---|---|---|---|---|---|---|---|---|---|---|---|---|---|---|
| LIRI-JP::SP99217 | Liver | LIRI-JP | 279 | 0 | 43 | 13 | 11 | 0.016 | 0.010 | 0.010 | 0.006 | 1.0272 | 0.3043 | 0.5537 | Decreasing |
| PACA-AU::SP71228 | Pancreas | PACA-AU | 1074 | 0 | 0 | 29 | 11 | 0.018 | 0.016 | 0.029 | 0.027 | -1.0258 | 0.3050 | 0.5542 | Increasing |
| RECA-EU::SP102804 | Kidney | RECA-EU | 1031 | 0 | 129 | 60 | 8 | 0.033 | 0.052 | 0.048 | 0.055 | -1.0239 | 0.3059 | 0.5544 | Increasing |
| BRCA-US::SP8660 | Breast | BRCA-US | 812 | 523 | 240 | 167 | 13 | 0.141 | 0.137 | 0.113 | 0.201 | -1.0239 | 0.3059 | 0.5544 | Increasing |
| KIRC-US::SP37970 | Kidney | KIRC-US | 1018 | 0 | 153 | 57 | 9 | 0.033 | 0.059 | 0.040 | 0.064 | -1.0181 | 0.3086 | 0.5587 | Increasing |
| SKCM-US::SP82900 | Skin | SKCM-US | 954 | 0 | 252 | 79 | 14 | 0.055 | 0.080 | 0.041 | 0.106 | -1.0171 | 0.3091 | 0.5589 | Increasing |
| SKCM-US::SP82796 | Skin | SKCM-US | 418 | 0 | 0 | 11 | 13 | 0.007 | 0.005 | 0.016 | 0.011 | -1.0133 | 0.3109 | 0.5608 | Increasing |
| PACA-AU::SP73263 | Pancreas | PACA-AU | 615 | 0 | 0 | 12 | 10 | 0.004 | 0.011 | 0.009 | 0.013 | -1.0137 | 0.3107 | 0.5608 | Increasing |
| HNSC-US::SP32662 | Head&Neck | HNSC-US | 745 | 0 | 123 | 33 | 11 | 0.030 | 0.014 | 0.036 | 0.040 | -1.0087 | 0.3131 | 0.5634 | Increasing |
| KIRP-US::SP106656 | Kidney | KIRP-US | 1311 | 0 | 204 | 65 | 9 | 0.080 | 0.043 | 0.044 | 0.062 | 1.0086 | 0.3132 | 0.5634 | Decreasing |
| LIRI-JP::SP107150 | Liver | LIRI-JP | 495 | 0 | 73 | 27 | 11 | 0.029 | 0.008 | 0.025 | 0.041 | -1.0016 | 0.3165 | 0.5674 | Increasing |
| OV-AU::SP101690 | Ovary | OV-AU | 567 | 110 | 81 | 63 | 13 | 0.057 | 0.057 | 0.030 | 0.052 | 1.0023 | 0.3162 | 0.5674 | Decreasing |
| PAEN-AU::SP102611 | Pancreas | PAEN-AU | 883 | 0 | 135 | 24 | 12 | 0.021 | 0.021 | 0.024 | 0.009 | 1.0032 | 0.3158 | 0.5674 | Decreasing |
| GBM-US::SP25380 | Brain | GBM-US | 505 | 0 | 0 | 11 | 8 | 0.005 | 0.007 | 0.015 | 0.010 | -0.9999 | 0.3174 | 0.5681 | Increasing |
| READ-US::SP81137 | CRC | READ-US | 2673 | 0 | 0 | 45 | 16 | 0.034 | 0.029 | 0.054 | 0.040 | -0.9936 | 0.3204 | 0.5729 | Increasing |
| LUSC-US::SP56533 | Lung | LUSC-US | 1112 | 0 | 159 | 38 | 13 | 0.035 | 0.037 | 0.033 | 0.017 | 0.9867 | 0.3238 | 0.5782 | Decreasing |
| LIHC-US::SP98164 | Liver | LIHC-US | 888 | 0 | 112 | 50 | 12 | 0.044 | 0.029 | 0.048 | 0.060 | -0.9830 | 0.3256 | 0.5808 | Increasing |
| GBM-US::SP25494 | Brain | GBM-US | 1962 | 0 | 493 | 129 | 9 | 0.114 | 0.116 | 0.119 | 0.077 | 0.9802 | 0.3270 | 0.5824 | Decreasing |
| MELA-AU::SP124333 | Skin | SKCM-US | 393 | 0 | 91 | 24 | 12.5 | 0.024 | 0.026 | 0.013 | 0.017 | 0.9794 | 0.3274 | 0.5825 | Decreasing |
| LIRI-JP::SP112174 | Liver | LIRI-JP | 364 | 0 | 112 | 30 | 11 | 0.019 | 0.025 | 0.025 | 0.035 | -0.9753 | 0.3294 | 0.5840 | Increasing |
| LIHC-US::SP49205 | Liver | LIHC-US | 376 | 46 | 48 | 20 | 13 | 0.016 | 0.013 | 0.016 | 0.030 | -0.9754 | 0.3294 | 0.5840 | Increasing |
| PRAD-CA::SP112995 | Prostate | PRAD-US | 333 | 0 | 35 | 11 | 8.5 | 0.007 | 0.016 | 0.009 | 0.000 | 0.9760 | 0.3291 | 0.5840 | Decreasing |
| KIRP-US::SP43651 | Kidney | KIRP-US | 1363 | 0 | 173 | 69 | 9 | 0.049 | 0.062 | 0.053 | 0.077 | -0.9683 | 0.3329 | 0.5879 | Increasing |
| LUSC-US::SP58612 | Lung | LUSC-US | 1494 | 0 | 225 | 59 | 12 | 0.032 | 0.053 | 0.075 | 0.034 | -0.9695 | 0.3323 | 0.5879 | Increasing |
| OV-AU::SP102187 | Ovary | OV-AU | 586 | 222 | 154 | 100 | 18 | 0.086 | 0.086 | 0.063 | 0.072 | 0.9688 | 0.3327 | 0.5879 | Decreasing |
| GACA-CN::SP135213 | Stomach | STAD-US | 352 | 86 | 0 | 31 | 13 | 0.018 | 0.028 | 0.029 | 0.032 | -0.9612 | 0.3364 | 0.5923 | Increasing |
| PACA-AU::SP110822 | Pancreas | PACA-AU | 662 | 0 | 0 | 20 | 12 | 0.015 | 0.014 | 0.009 | 0.027 | -0.9601 | 0.3370 | 0.5923 | Increasing |
| OV-AU::SP101519 | Ovary | OV-AU | 409 | 68 | 64 | 38 | 11 | 0.029 | 0.016 | 0.051 | 0.026 | -0.9605 | 0.3368 | 0.5923 | Increasing |
| MELA-AU::SP124365 | Skin | SKCM-US | 742 | 0 | 165 | 57 | 13 | 0.041 | 0.041 | 0.063 | 0.050 | -0.9617 | 0.3362 | 0.5923 | Increasing |
| BRCA-US::SP8532 | Breast | BRCA-US | 242 | 0 | 82 | 13 | 16 | 0.003 | 0.018 | 0.003 | 0.021 | -0.9581 | 0.3380 | 0.5926 | Increasing |
| BRCA-US::SP10470 | Breast | BRCA-US | 796 | 384 | 308 | 185 | 18 | 0.152 | 0.150 | 0.148 | 0.196 | -0.9586 | 0.3378 | 0.5926 | Increasing |
| PRAD-UK::SP114930 | Prostate | PRAD-US | 759 | 0 | 129 | 32 | 9 | 0.037 | 0.026 | 0.021 | 0.025 | 0.9532 | 0.3405 | 0.5963 | Decreasing |
| OV-AU::SP101548 | Ovary | OV-AU | 618 | 404 | 103 | 125 | 15 | 0.111 | 0.095 | 0.096 | 0.083 | 0.9495 | 0.3424 | 0.5981 | Decreasing |
| BRCA-UK::SP2160 | Breast | BRCA-US | 176 | 34 | 33 | 11 | 15 | 0.003 | 0.013 | 0.006 | 0.016 | -0.9496 | 0.3423 | 0.5981 | Increasing |
| OV-AU::SP101596 | Ovary | OV-AU | 397 | 0 | 47 | 14 | 9 | 0.013 | 0.007 | 0.006 | 0.026 | -0.9473 | 0.3435 | 0.5994 | Increasing |
| PRAD-UK::SP114949 | Prostate | PRAD-US | 688 | 0 | 0 | 12 | 9.5 | 0.007 | 0.005 | 0.018 | 0.010 | -0.9428 | 0.3458 | 0.6026 | Increasing |
| PACA-AU::SP107770 | Pancreas | PACA-AU | 396 | 0 | 129 | 17 | 13 | 0.004 | 0.016 | 0.018 | 0.013 | -0.9419 | 0.3463 | 0.6027 | Increasing |
| MALY-DE::SP116668 | Blood | MALY-DE | 1782 | 0 | 0 | 18 | 12 | 0.006 | 0.016 | 0.009 | 0.021 | -0.9392 | 0.3476 | 0.6044 | Increasing |
| PRAD-CA::SP112997 | Prostate | PRAD-US | 316 | 0 | 63 | 26 | 10 | 0.015 | 0.024 | 0.021 | 0.030 | -0.9376 | 0.3484 | 0.6044 | Increasing |
| UCEC-US::SP94332 | Endometrium | UCEC-US | 1084 | 512 | 464 | 273 | 22 | 0.247 | 0.243 | 0.209 | 0.221 | 0.9381 | 0.3482 | 0.6044 | Decreasing |
| PRAD-UK::SP114912 | Prostate | PRAD-US | 798 | 0 | 0 | 14 | 13 | 0.004 | 0.013 | 0.018 | 0.010 | -0.9307 | 0.3520 | 0.6072 | Increasing |
| BRCA-US::SP3415 | Breast | BRCA-US | 461 | 180 | 153 | 90 | 15 | 0.059 | 0.079 | 0.091 | 0.074 | -0.9311 | 0.3518 | 0.6072 | Increasing |
| SKCM-US::SP104330 | Skin | SKCM-US | 82 | 0 | 42 | 16 | 14 | 0.010 | 0.008 | 0.025 | 0.011 | -0.9300 | 0.3524 | 0.6072 | Increasing |
| PRAD-CA::SP112991 | Prostate | PRAD-US | 502 | 0 | 0 | 14 | 10 | 0.011 | 0.008 | 0.012 | 0.020 | -0.9307 | 0.3520 | 0.6072 | Increasing |
| OV-AU::SP101795 | Ovary | OV-AU | 732 | 330 | 361 | 185 | 21 | 0.150 | 0.159 | 0.129 | 0.129 | 0.9295 | 0.3526 | 0.6072 | Decreasing |
| BRCA-UK::SP2148 | Breast | BRCA-US | 331 | 146 | 155 | 66 | 18 | 0.055 | 0.074 | 0.041 | 0.048 | 0.9323 | 0.3512 | 0.6072 | Decreasing |
| STAD-US::SP84743 | Stomach | STAD-US | 3209 | 0 | 0 | 50 | 14.5 | 0.034 | 0.048 | 0.036 | 0.058 | -0.9229 | 0.3560 | 0.6073 | Increasing |
| PACA-AU::SP70956 | Pancreas | PACA-AU | 385 | 0 | 0 | 11 | 12.5 | 0.007 | 0.003 | 0.015 | 0.010 | -0.9236 | 0.3557 | 0.6073 | Increasing |
| PACA-AU::SP107874 | Pancreas | PACA-AU | 827 | 0 | 0 | 20 | 13.5 | 0.022 | 0.016 | 0.012 | 0.013 | 0.9226 | 0.3562 | 0.6073 | Decreasing |
| KIRC-US::SP35849 | Kidney | KIRC-US | 283 | 0 | 50 | 20 | 9 | 0.015 | 0.011 | 0.025 | 0.020 | -0.9243 | 0.3553 | 0.6073 | Increasing |

| | | | | | | | | | | | | | | | |
|---|---|---|---|---|---|---|---|---|---|---|---|---|---|---|---|
| HNSC-US::SP115162 | Head&Neck | HNSC-US | 205 | 0 | 34 | 13 | 12.5 | 0.006 | 0.011 | 0.016 | 0.012 | -0.9275 | 0.3536 | 0.6073 | Increasing |
| PACA-AU::SP75977 | Pancreas | PACA-AU | 357 | 0 | 80 | 14 | 9 | 0.015 | 0.000 | 0.015 | 0.017 | -0.9283 | 0.3533 | 0.6073 | Increasing |
| MALY-DE::SP116706 | Blood | MALY-DE | 706 | 0 | 0 | 13 | 11 | 0.006 | 0.009 | 0.009 | 0.015 | -0.9222 | 0.3564 | 0.6073 | Increasing |
| BRCA-US::SP6223 | Breast | BRCA-US | 555 | 189 | 200 | 80 | 18 | 0.052 | 0.066 | 0.091 | 0.058 | -0.9266 | 0.3541 | 0.6073 | Increasing |
| PRAD-UK::SP111131 | Prostate | PRAD-US | 314 | 0 | 45 | 16 | 9 | 0.011 | 0.011 | 0.015 | 0.020 | -0.9247 | 0.3551 | 0.6073 | Increasing |
| BRCA-US::SP9930 | Breast | BRCA-US | 827 | 368 | 352 | 182 | 19 | 0.152 | 0.174 | 0.157 | 0.117 | 0.9206 | 0.3573 | 0.6080 | Decreasing |
| MELA-AU::SP124271 | Skin | SKCM-US | 894 | 0 | 134 | 35 | 11 | 0.031 | 0.034 | 0.035 | 0.011 | 0.9112 | 0.3622 | 0.6157 | Decreasing |
| PACA-AU::SP70860 | Pancreas | PACA-AU | 672 | 0 | 0 | 22 | 10 | 0.018 | 0.011 | 0.015 | 0.027 | -0.9072 | 0.3643 | 0.6185 | Increasing |
| SKCM-US::SP82433 | Skin | SKCM-US | 406 | 0 | 56 | 14 | 15 | 0.010 | 0.013 | 0.003 | 0.028 | -0.9030 | 0.3665 | 0.6216 | Increasing |
| PACA-CA::SP77888 | Pancreas | PACA-CA | 164 | 0 | 27 | 12 | 16 | 0.004 | 0.011 | 0.008 | 0.013 | -0.8998 | 0.3682 | 0.6227 | Increasing |
| LUSC-US::SP56474 | Lung | LUSC-US | 776 | 0 | 143 | 33 | 11 | 0.035 | 0.011 | 0.033 | 0.045 | -0.8986 | 0.3689 | 0.6227 | Increasing |
| LIHC-US::SP98053 | Liver | LIHC-US | 774 | 0 | 104 | 29 | 9 | 0.025 | 0.016 | 0.029 | 0.036 | -0.8991 | 0.3686 | 0.6227 | Increasing |
| BRCA-UK::SP116363 | Breast | BRCA-US | 311 | 157 | 154 | 72 | 16 | 0.076 | 0.055 | 0.060 | 0.053 | 0.9002 | 0.3680 | 0.6227 | Decreasing |
| LIHC-US::SP115830 | Liver | LIHC-US | 584 | 0 | 0 | 21 | 11 | 0.013 | 0.018 | 0.019 | 0.024 | -0.8916 | 0.3726 | 0.6282 | Increasing |
| LIRI-JP::SP112217 | Liver | LIRI-JP | 924 | 88 | 167 | 76 | 14 | 0.076 | 0.060 | 0.057 | 0.058 | 0.8905 | 0.3732 | 0.6285 | Decreasing |
| BRCA-US::SP5279 | Breast | BRCA-US | 871 | 0 | 172 | 27 | 16 | 0.017 | 0.029 | 0.009 | 0.042 | -0.8874 | 0.3749 | 0.6306 | Increasing |
| ORCA-IN::SP117482 | Head&Neck | ORCA-IN | 521 | 0 | 0 | 18 | 10.5 | 0.019 | 0.005 | 0.012 | 0.030 | -0.8860 | 0.3756 | 0.6310 | Increasing |
| PACA-CA::SP125704 | Pancreas | PACA-CA | 399 | 0 | 0 | 15 | 9 | 0.019 | 0.003 | 0.003 | 0.025 | -0.8853 | 0.3760 | 0.6310 | Increasing |
| PACA-CA::SP117981 | Pancreas | PACA-CA | 364 | 0 | 117 | 18 | 11 | 0.019 | 0.017 | 0.008 | 0.013 | 0.8814 | 0.3781 | 0.6316 | Decreasing |
| MELA-AU::SP124394 | Skin | SKCM-US | 609 | 0 | 160 | 46 | 15 | 0.044 | 0.036 | 0.050 | 0.017 | 0.8823 | 0.3776 | 0.6316 | Decreasing |
| GACA-CN::SP135405 | Stomach | STAD-US | 812 | 0 | 0 | 25 | 12 | 0.028 | 0.022 | 0.013 | 0.021 | 0.8836 | 0.3769 | 0.6316 | Decreasing |
| KIRC-US::SP39349 | Kidney | KIRC-US | 577 | 0 | 105 | 33 | 10 | 0.022 | 0.024 | 0.037 | 0.029 | -0.8818 | 0.3779 | 0.6316 | Increasing |
| BRCA-US::SP9816 | Breast | BRCA-US | 738 | 222 | 366 | 125 | 18 | 0.117 | 0.124 | 0.066 | 0.122 | 0.8805 | 0.3786 | 0.6317 | Decreasing |
| PACA-AU::SP71974 | Pancreas | PACA-AU | 2125 | 966 | 0 | 284 | 15 | 0.264 | 0.201 | 0.211 | 0.221 | 0.8787 | 0.3796 | 0.6326 | Decreasing |
| OV-AU::SP102174 | Ovary | OV-AU | 447 | 69 | 59 | 42 | 13 | 0.032 | 0.029 | 0.027 | 0.052 | -0.8722 | 0.3831 | 0.6329 | Increasing |
| OV-AU::SP101708 | Ovary | OV-AU | 335 | 56 | 0 | 18 | 9 | 0.013 | 0.023 | 0.006 | 0.010 | 0.8712 | 0.3837 | 0.6329 | Decreasing |
| LIHC-US::SP98090 | Liver | LIHC-US | 1889 | 0 | 0 | 35 | 11 | 0.025 | 0.021 | 0.048 | 0.024 | -0.8725 | 0.3829 | 0.6329 | Increasing |
| OV-US::SP69077 | Ovary | OV-US | 1429 | 722 | 707 | 374 | 20 | 0.376 | 0.292 | 0.267 | 0.365 | 0.8719 | 0.3833 | 0.6329 | Decreasing |
| LIHC-US::SP49322 | Liver | LIHC-US | 652 | 0 | 0 | 16 | 11.5 | 0.016 | 0.005 | 0.016 | 0.024 | -0.8724 | 0.3830 | 0.6329 | Increasing |
| STAD-US::SP84439 | Stomach | STAD-US | 96781 | 0 | 0 | 51 | 8 | 0.049 | 0.028 | 0.039 | 0.069 | -0.8730 | 0.3827 | 0.6329 | Increasing |
| BLCA-US::SP1174 | Bladder | BLCA-US | 1037 | 0 | 0 | 28 | 13 | 0.015 | 0.029 | 0.023 | 0.030 | -0.8717 | 0.3834 | 0.6329 | Increasing |
| LIRI-JP::SP107030 | Liver | LIRI-JP | 470 | 0 | 53 | 21 | 9 | 0.022 | 0.020 | 0.010 | 0.017 | 0.8726 | 0.3829 | 0.6329 | Decreasing |
| BRCA-UK::SP2357 | Breast | BRCA-US | 528 | 191 | 121 | 78 | 18 | 0.079 | 0.066 | 0.057 | 0.064 | 0.8723 | 0.3831 | 0.6329 | Decreasing |
| SKCM-US::SP83242 | Skin | SKCM-US | 516 | 0 | 89 | 30 | 13 | 0.031 | 0.028 | 0.019 | 0.022 | 0.8694 | 0.3846 | 0.6338 | Decreasing |
| HNSC-US::SP29987 | Head&Neck | HNSC-US | 1136 | 0 | 148 | 47 | 14 | 0.039 | 0.025 | 0.059 | 0.040 | -0.8684 | 0.3852 | 0.6340 | Increasing |
| UCEC-US::SP90725 | Endometrium | UCEC-US | 421 | 0 | 0 | 16 | 12 | 0.010 | 0.016 | 0.010 | 0.023 | -0.8623 | 0.3885 | 0.6347 | Increasing |
| LINC-JP::SP98859 | Liver | LIHC-US | 443 | 0 | 73 | 27 | 11 | 0.028 | 0.024 | 0.019 | 0.018 | 0.8644 | 0.3874 | 0.6347 | Decreasing |
| COAD-US::SP17905 | CRC | COAD-US | 25734 | 0 | 0 | 11 | 7 | 0.009 | 0.005 | 0.010 | 0.017 | -0.8622 | 0.3886 | 0.6347 | Increasing |
| LUAD-US::SP52232 | Lung | LUAD-US | 936 | 0 | 0 | 15 | 13.5 | 0.013 | 0.011 | 0.007 | 0.027 | -0.8620 | 0.3887 | 0.6347 | Increasing |
| READ-US::SP81172 | CRC | READ-US | 1326 | 0 | 0 | 19 | 9 | 0.006 | 0.021 | 0.023 | 0.012 | -0.8646 | 0.3872 | 0.6347 | Increasing |
| RECA-EU::SP103057 | Kidney | RECA-EU | 939 | 0 | 110 | 46 | 9 | 0.016 | 0.047 | 0.043 | 0.030 | -0.8624 | 0.3885 | 0.6347 | Increasing |
| HNSC-US::SP33774 | Head&Neck | HNSC-US | 299 | 53 | 0 | 17 | 9 | 0.021 | 0.005 | 0.026 | 0.000 | 0.8636 | 0.3878 | 0.6347 | Decreasing |
| LUSC-US::SP56566 | Lung | LUSC-US | 852 | 0 | 144 | 31 | 13 | 0.016 | 0.034 | 0.023 | 0.034 | -0.8549 | 0.3926 | 0.6387 | Increasing |
| PACA-CA::SP117340 | Pancreas | PACA-CA | 832 | 0 | 196 | 40 | 12 | 0.026 | 0.028 | 0.031 | 0.038 | -0.8544 | 0.3929 | 0.6387 | Increasing |
| STAD-US::SP105577 | Stomach | STAD-US | 828 | 0 | 101 | 30 | 11.5 | 0.022 | 0.025 | 0.023 | 0.037 | -0.8551 | 0.3925 | 0.6387 | Increasing |
| BRCA-US::SP96147 | Breast | BRCA-US | 258 | 0 | 36 | 16 | 12.5 | 0.010 | 0.016 | 0.006 | 0.026 | -0.8556 | 0.3922 | 0.6387 | Increasing |
| BRCA-UK::SP2147 | Breast | BRCA-US | 1201 | 702 | 236 | 230 | 14 | 0.207 | 0.211 | 0.170 | 0.191 | 0.8530 | 0.3936 | 0.6392 | Decreasing |
| LIRI-JP::SP99309 | Liver | LIRI-JP | 495 | 0 | 0 | 11 | 10 | 0.010 | 0.003 | 0.016 | 0.012 | -0.8498 | 0.3954 | 0.6392 | Increasing |
| LIRI-JP::SP99341 | Liver | LIRI-JP | 335 | 0 | 0 | 11 | 8 | 0.006 | 0.008 | 0.013 | 0.012 | -0.8498 | 0.3954 | 0.6392 | Increasing |
| LIRI-JP::SP99213 | Liver | LIRI-JP | 542 | 0 | 0 | 11 | 7 | 0.006 | 0.010 | 0.006 | 0.017 | -0.8498 | 0.3954 | 0.6392 | Increasing |

| | | | | | | | | | | | | | | | |
|---|---|---|---|---|---|---|---|---|---|---|---|---|---|---|---|
| PACA-AU::SP71456 | Pancreas | PACA-AU | 708 | 0 | 157 | 34 | 13 | 0.022 | 0.024 | 0.026 | 0.033 | -0.8507 | 0.3949 | 0.6392 | Increasing |
| LIRI-JP::SP112188 | Liver | LIRI-JP | 326 | 0 | 39 | 16 | 11.5 | 0.010 | 0.013 | 0.016 | 0.017 | -0.8445 | 0.3984 | 0.6426 | Increasing |
| LIRI-JP::SP112237 | Liver | LIRI-JP | 437 | 0 | 49 | 16 | 12.5 | 0.010 | 0.013 | 0.016 | 0.017 | -0.8445 | 0.3984 | 0.6426 | Increasing |
| LIHC-US::SP49223 | Liver | LIHC-US | 287 | 46 | 47 | 19 | 10 | 0.013 | 0.010 | 0.029 | 0.012 | -0.8373 | 0.4024 | 0.6464 | Increasing |
| KIRC-US::SP42829 | Kidney | KIRC-US | 793 | 0 | 94 | 49 | 8 | 0.040 | 0.038 | 0.037 | 0.059 | -0.8340 | 0.4043 | 0.6464 | Increasing |
| KIRP-US::SP97194 | Kidney | KIRP-US | 2180 | 0 | 324 | 129 | 9 | 0.091 | 0.115 | 0.118 | 0.113 | -0.8343 | 0.4041 | 0.6464 | Increasing |
| LUSC-US::SP57629 | Lung | LUSC-US | 2063 | 0 | 0 | 34 | 11 | 0.022 | 0.032 | 0.026 | 0.040 | -0.8357 | 0.4033 | 0.6464 | Increasing |
| MALY-DE::SP59300 | Blood | MALY-DE | 795 | 190 | 112 | 75 | 15 | 0.070 | 0.063 | 0.052 | 0.054 | 0.8332 | 0.4048 | 0.6464 | Decreasing |
| OV-AU::SP102035 | Ovary | OV-AU | 999 | 675 | 324 | 260 | 17 | 0.217 | 0.216 | 0.171 | 0.207 | 0.8331 | 0.4048 | 0.6464 | Decreasing |
| PACA-AU::SP69898 | Pancreas | PACA-AU | 1321 | 0 | 0 | 19 | 11 | 0.026 | 0.011 | 0.009 | 0.017 | 0.8349 | 0.4038 | 0.6464 | Decreasing |
| BRCA-UK::SP116375 | Breast | BRCA-US | 498 | 0 | 97 | 23 | 9.5 | 0.017 | 0.018 | 0.016 | 0.032 | -0.8341 | 0.4042 | 0.6464 | Increasing |
| BRCA-UK::SP116359 | Breast | BRCA-US | 632 | 257 | 257 | 120 | 18 | 0.090 | 0.132 | 0.094 | 0.074 | 0.8341 | 0.4042 | 0.6464 | Decreasing |
| MALY-DE::SP116618 | Blood | MALY-DE | 385 | 68 | 0 | 16 | 10 | 0.000 | 0.018 | 0.006 | 0.018 | -0.8296 | 0.4068 | 0.6489 | Increasing |
| KIRP-US::SP43770 | Kidney | KIRP-US | 613 | 0 | 104 | 40 | 10 | 0.049 | 0.024 | 0.034 | 0.031 | 0.8268 | 0.4083 | 0.6506 | Decreasing |
| MELA-AU::SP124305 | Skin | SKCM-US | 458 | 0 | 84 | 29 | 12 | 0.027 | 0.028 | 0.022 | 0.017 | 0.8242 | 0.4098 | 0.6523 | Decreasing |
| LIRI-JP::SP50099 | Liver | LIRI-JP | 759 | 0 | 102 | 38 | 13 | 0.045 | 0.015 | 0.019 | 0.070 | -0.8187 | 0.4130 | 0.6537 | Increasing |
| BRCA-US::SP2881 | Breast | BRCA-US | 795 | 0 | 162 | 35 | 11 | 0.024 | 0.032 | 0.025 | 0.042 | -0.8200 | 0.4122 | 0.6537 | Increasing |
| BRCA-US::SP2793 | Breast | BRCA-US | 1933 | 889 | 666 | 403 | 17 | 0.369 | 0.338 | 0.337 | 0.323 | 0.8187 | 0.4129 | 0.6537 | Decreasing |
| KIRC-US::SP36586 | Kidney | KIRC-US | 349 | 0 | 49 | 21 | 12 | 0.015 | 0.016 | 0.019 | 0.025 | -0.8195 | 0.4125 | 0.6537 | Increasing |
| HNSC-US::SP30675 | Head&Neck | HNSC-US | 2521 | 0 | 0 | 32 | 13 | 0.042 | 0.016 | 0.020 | 0.035 | 0.8206 | 0.4119 | 0.6537 | Decreasing |
| RECA-EU::SP103128 | Kidney | RECA-EU | 707 | 0 | 136 | 64 | 9 | 0.036 | 0.049 | 0.068 | 0.040 | -0.8175 | 0.4136 | 0.6541 | Increasing |
| OV-AU::SP102161 | Ovary | OV-AU | 401 | 104 | 65 | 43 | 12 | 0.029 | 0.032 | 0.036 | 0.041 | -0.8138 | 0.4157 | 0.6551 | Increasing |
| CLLE-ES::SP13367 | Blood | CLLE-ES | 249 | 0 | 0 | 11 | 10 | 0.013 | 0.006 | 0.006 | 0.008 | 0.8149 | 0.4152 | 0.6551 | Decreasing |
| PACA-CA::SP125785 | Pancreas | PACA-CA | 476 | 0 | 0 | 20 | 14 | 0.008 | 0.020 | 0.014 | 0.019 | -0.8132 | 0.4161 | 0.6551 | Increasing |
| STAD-US::SP84962 | Stomach | STAD-US | 718 | 0 | 0 | 28 | 12 | 0.015 | 0.022 | 0.039 | 0.016 | -0.8140 | 0.4156 | 0.6551 | Increasing |
| BRCA-US::SP6519 | Breast | BRCA-US | 193 | 37 | 42 | 12 | 15 | 0.003 | 0.013 | 0.013 | 0.011 | -0.8117 | 0.4170 | 0.6558 | Increasing |
| PRAD-UK::SP111154 | Prostate | PRAD-US | 461 | 0 | 60 | 19 | 10 | 0.011 | 0.016 | 0.018 | 0.020 | -0.8102 | 0.4178 | 0.6564 | Increasing |
| PRAD-CA::SP112953 | Prostate | PRAD-US | 451 | 0 | 49 | 17 | 12.5 | 0.007 | 0.016 | 0.018 | 0.015 | -0.8040 | 0.4214 | 0.6598 | Increasing |
| PACA-CA::SP125791 | Pancreas | PACA-CA | 528 | 0 | 0 | 14 | 8 | 0.015 | 0.006 | 0.003 | 0.022 | -0.8053 | 0.4206 | 0.6598 | Increasing |
| PRAD-UK::SP111125 | Prostate | PRAD-US | 514 | 0 | 63 | 17 | 9 | 0.007 | 0.013 | 0.024 | 0.010 | -0.8040 | 0.4214 | 0.6598 | Increasing |
| PRAD-UK::SP114908 | Prostate | PRAD-US | 680 | 0 | 0 | 15 | 11.5 | 0.007 | 0.013 | 0.015 | 0.015 | -0.8001 | 0.4237 | 0.6598 | Increasing |
| UCEC-US::SP93772 | Endometrium | UCEC-US | 387 | 0 | 76 | 22 | 11 | 0.019 | 0.016 | 0.013 | 0.035 | -0.8012 | 0.4230 | 0.6598 | Increasing |
| PACA-AU::SP76515 | Pancreas | PACA-AU | 570 | 0 | 180 | 21 | 11 | 0.022 | 0.014 | 0.020 | 0.010 | 0.8025 | 0.4223 | 0.6598 | Decreasing |
| PACA-AU::SP108059 | Pancreas | PACA-AU | 392 | 0 | 78 | 18 | 9 | 0.007 | 0.019 | 0.009 | 0.020 | -0.8006 | 0.4234 | 0.6598 | Increasing |
| OV-US::SP67346 | Ovary | OV-US | 656 | 96 | 63 | 42 | 16 | 0.049 | 0.033 | 0.020 | 0.046 | 0.8012 | 0.4230 | 0.6598 | Decreasing |
| OV-AU::SP101528 | Ovary | OV-AU | 293 | 54 | 44 | 21 | 15 | 0.019 | 0.014 | 0.027 | 0.000 | 0.7961 | 0.4260 | 0.6628 | Decreasing |
| MELA-AU::SP124431 | Skin | SKCM-US | 699 | 0 | 208 | 71 | 16 | 0.065 | 0.044 | 0.066 | 0.078 | -0.7935 | 0.4275 | 0.6630 | Increasing |
| LIRI-JP::SP99165 | Liver | LIRI-JP | 514 | 0 | 82 | 30 | 11 | 0.025 | 0.018 | 0.029 | 0.035 | -0.7941 | 0.4271 | 0.6630 | Increasing |
| LIRI-JP::SP50151 | Liver | LIRI-JP | 412 | 0 | 66 | 30 | 13 | 0.013 | 0.030 | 0.035 | 0.017 | -0.7941 | 0.4271 | 0.6630 | Increasing |
| LUSC-US::SP56502 | Lung | LUSC-US | 1882 | 0 | 0 | 37 | 11 | 0.029 | 0.048 | 0.016 | 0.028 | 0.7914 | 0.4287 | 0.6642 | Decreasing |
| LIRI-JP::SP99137 | Liver | LIRI-JP | 837 | 0 | 155 | 53 | 12 | 0.045 | 0.038 | 0.041 | 0.064 | -0.7875 | 0.4310 | 0.6669 | Increasing |
| GACA-CN::SP135425 | Stomach | STAD-US | 842 | 0 | 0 | 18 | 11 | 0.018 | 0.020 | 0.007 | 0.016 | 0.7860 | 0.4319 | 0.6676 | Decreasing |
| OV-AU::SP101716 | Ovary | OV-AU | 755 | 103 | 91 | 45 | 12 | 0.041 | 0.032 | 0.042 | 0.021 | 0.7835 | 0.4333 | 0.6692 | Decreasing |
| PAEN-AU::SP102517 | Pancreas | PAEN-AU | 174 | 0 | 29 | 11 | 15.5 | 0.010 | 0.005 | 0.005 | 0.015 | -0.7808 | 0.4349 | 0.6699 | Increasing |
| LIRI-JP::SP98991 | Liver | LIRI-JP | 289 | 0 | 75 | 33 | 10 | 0.032 | 0.020 | 0.019 | 0.052 | -0.7811 | 0.4348 | 0.6699 | Increasing |
| OV-US::SP68725 | Ovary | OV-US | 421 | 219 | 115 | 70 | 19 | 0.056 | 0.061 | 0.046 | 0.087 | -0.7803 | 0.4352 | 0.6699 | Increasing |
| RECA-EU::SP102897 | Kidney | RECA-EU | 13768 | 0 | 0 | 85 | 8 | 0.069 | 0.042 | 0.096 | 0.060 | -0.7788 | 0.4361 | 0.6705 | Increasing |
| UCEC-US::SP90503 | Endometrium | UCEC-US | 346 | 0 | 71 | 17 | 9.5 | 0.003 | 0.021 | 0.023 | 0.006 | -0.7694 | 0.4416 | 0.6783 | Increasing |
| LUSC-US::SP56821 | Lung | LUSC-US | 1524 | 0 | 0 | 19 | 10.5 | 0.013 | 0.013 | 0.023 | 0.017 | -0.7636 | 0.4451 | 0.6830 | Increasing |
| KIRC-US::SP37516 | Kidney | KIRC-US | 557 | 0 | 83 | 28 | 9 | 0.022 | 0.019 | 0.028 | 0.029 | -0.7620 | 0.4460 | 0.6837 | Increasing |

| | | | | | | | | | | | | | | | |
|---|---|---|---|---|---|---|---|---|---|---|---|---|---|---|---|
| PRAD-CA::SP112883 | Prostate | PRAD-US | 234 | 0 | 25 | 12 | 9 | 0.015 | 0.008 | 0.012 | 0.005 | 0.7612 | 0.4465 | 0.6837 | Decreasing |
| LINC-JP::SP98841 | Liver | LIHC-US | 378 | 53 | 0 | 12 | 7 | 0.009 | 0.008 | 0.010 | 0.018 | -0.7555 | 0.4499 | 0.6882 | Increasing |
| MALY-DE::SP59412 | Blood | MALY-DE | 1089 | 0 | 0 | 13 | 7.5 | 0.026 | 0.007 | 0.006 | 0.012 | 0.7525 | 0.4517 | 0.6895 | Decreasing |
| MALY-DE::SP59352 | Blood | MALY-DE | 657 | 0 | 0 | 13 | 8 | 0.019 | 0.009 | 0.009 | 0.009 | 0.7525 | 0.4517 | 0.6895 | Decreasing |
| COAD-US::SP16886 | CRC | COAD-US | 7736 | 0 | 0 | 12 | 13 | 0.003 | 0.014 | 0.017 | 0.006 | -0.7475 | 0.4547 | 0.6897 | Increasing |
| OV-US::SP66159 | Ovary | OV-US | 1262 | 649 | 247 | 224 | 12 | 0.219 | 0.173 | 0.188 | 0.186 | 0.7476 | 0.4547 | 0.6897 | Decreasing |
| KIRP-US::SP97113 | Kidney | KIRP-US | 830 | 0 | 143 | 54 | 9 | 0.038 | 0.051 | 0.037 | 0.062 | -0.7469 | 0.4551 | 0.6897 | Increasing |
| GBM-US::SP24815 | Brain | GBM-US | 581 | 0 | 0 | 16 | 6 | 0.018 | 0.012 | 0.018 | 0.005 | 0.7503 | 0.4531 | 0.6897 | Decreasing |
| LIRI-JP::SP99129 | Liver | LIRI-JP | 762 | 0 | 163 | 48 | 12.5 | 0.041 | 0.033 | 0.038 | 0.058 | -0.7468 | 0.4552 | 0.6897 | Increasing |
| STAD-US::SP105375 | Stomach | STAD-US | 3099 | 0 | 0 | 48 | 11 | 0.031 | 0.042 | 0.052 | 0.037 | -0.7491 | 0.4538 | 0.6897 | Increasing |
| GACA-CN::SP135182 | Stomach | STAD-US | 2858 | 0 | 0 | 37 | 10 | 0.028 | 0.025 | 0.043 | 0.032 | -0.7497 | 0.4534 | 0.6897 | Increasing |
| READ-US::SP80615 | CRC | READ-US | 19514 | 0 | 0 | 54 | 14 | 0.055 | 0.042 | 0.044 | 0.040 | 0.7450 | 0.4563 | 0.6906 | Decreasing |
| LIRI-JP::SP99325 | Liver | LIRI-JP | 129378 | 0 | 0 | 66 | 6 | 0.048 | 0.048 | 0.076 | 0.046 | -0.7383 | 0.4603 | 0.6960 | Increasing |
| BRCA-US::SP96511 | Breast | BRCA-US | 1216 | 753 | 279 | 243 | 16 | 0.231 | 0.203 | 0.186 | 0.212 | 0.7341 | 0.4629 | 0.6992 | Decreasing |
| KIRP-US::SP97249 | Kidney | KIRP-US | 1168 | 0 | 207 | 74 | 9 | 0.049 | 0.067 | 0.072 | 0.062 | -0.7314 | 0.4646 | 0.7009 | Increasing |
| READ-US::SP80367 | CRC | READ-US | 1105 | 0 | 0 | 14 | 12 | 0.018 | 0.011 | 0.003 | 0.017 | 0.7272 | 0.4671 | 0.7026 | Decreasing |
| PACA-CA::SP118048 | Pancreas | PACA-CA | 1437 | 0 | 0 | 16 | 12 | 0.015 | 0.008 | 0.006 | 0.022 | -0.7274 | 0.4670 | 0.7026 | Increasing |
| PACA-CA::SP125757 | Pancreas | PACA-CA | 521 | 0 | 154 | 16 | 13 | 0.008 | 0.017 | 0.006 | 0.019 | -0.7274 | 0.4670 | 0.7026 | Increasing |
| STAD-US::SP105213 | Stomach | STAD-US | 1001 | 0 | 0 | 24 | 13 | 0.018 | 0.017 | 0.023 | 0.027 | -0.7257 | 0.4680 | 0.7033 | Increasing |
| ORCA-IN::SP117779 | Head&Neck | ORCA-IN | 338 | 48 | 39 | 22 | 10 | 0.026 | 0.013 | 0.019 | 0.015 | 0.7204 | 0.4713 | 0.7073 | Decreasing |
| RECA-EU::SP102965 | Kidney | RECA-EU | 579 | 0 | 97 | 41 | 9 | 0.016 | 0.047 | 0.023 | 0.040 | -0.7198 | 0.4717 | 0.7073 | Decreasing |
| MALY-DE::SP59304 | Blood | MALY-DE | 3432 | 473 | 0 | 124 | 10 | 0.089 | 0.115 | 0.084 | 0.090 | 0.7182 | 0.4726 | 0.7080 | Decreasing |
| PACA-CA::SP125764 | Pancreas | PACA-CA | 1892 | 866 | 293 | 303 | 14 | 0.256 | 0.226 | 0.240 | 0.220 | 0.7157 | 0.4742 | 0.7096 | Decreasing |
| LIRI-JP::SP50125 | Liver | LIRI-JP | 614 | 0 | 97 | 37 | 11 | 0.025 | 0.030 | 0.035 | 0.035 | -0.7134 | 0.4756 | 0.7109 | Increasing |
| ORCA-IN::SP117376 | Head&Neck | ORCA-IN | 1276 | 0 | 0 | 27 | 12 | 0.013 | 0.023 | 0.034 | 0.015 | -0.7119 | 0.4765 | 0.7116 | Increasing |
| MELA-AU::SP124382 | Skin | SKCM-US | 3166 | 0 | 902 | 307 | 13 | 0.274 | 0.225 | 0.274 | 0.295 | -0.7069 | 0.4797 | 0.7148 | Increasing |
| SKCM-US::SP83844 | Skin | SKCM-US | 537 | 0 | 153 | 41 | 14 | 0.034 | 0.028 | 0.038 | 0.044 | -0.7073 | 0.4794 | 0.7148 | Increasing |
| LIRI-JP::SP112156 | Liver | LIRI-JP | 346 | 0 | 28 | 12 | 14 | 0.013 | 0.013 | 0.003 | 0.012 | 0.7005 | 0.4836 | 0.7199 | Decreasing |
| BRCA-UK::SP116377 | Breast | BRCA-US | 574 | 0 | 140 | 27 | 11 | 0.014 | 0.026 | 0.028 | 0.021 | -0.6987 | 0.4848 | 0.7209 | Increasing |
| LIRI-JP::SP98896 | Liver | LIRI-JP | 986 | 0 | 140 | 52 | 12 | 0.051 | 0.033 | 0.032 | 0.075 | -0.6970 | 0.4858 | 0.7210 | Increasing |
| LUSC-US::SP58668 | Lung | LUSC-US | 2298 | 0 | 0 | 42 | 12.5 | 0.035 | 0.029 | 0.039 | 0.045 | -0.6976 | 0.4854 | 0.7210 | Increasing |
| MELA-AU::SP124454 | Skin | SKCM-US | 467 | 0 | 87 | 30 | 12 | 0.038 | 0.018 | 0.022 | 0.028 | 0.6889 | 0.4909 | 0.7248 | Decreasing |
| BRCA-US::SP7421 | Breast | BRCA-US | 446 | 78 | 86 | 32 | 13 | 0.021 | 0.032 | 0.022 | 0.037 | -0.6900 | 0.4902 | 0.7248 | Increasing |
| RECA-EU::SP103706 | Kidney | RECA-EU | 1048 | 0 | 173 | 59 | 9 | 0.033 | 0.052 | 0.051 | 0.045 | -0.6900 | 0.4902 | 0.7248 | Increasing |
| HNSC-US::SP31030 | Head&Neck | HNSC-US | 994 | 0 | 0 | 21 | 11 | 0.018 | 0.011 | 0.023 | 0.023 | -0.6889 | 0.4909 | 0.7248 | Increasing |
| PRAD-CA::SP112775 | Prostate | PRAD-US | 375 | 0 | 55 | 18 | 10 | 0.007 | 0.016 | 0.024 | 0.010 | -0.6909 | 0.4896 | 0.7248 | Increasing |
| PRAD-UK::SP114924 | Prostate | PRAD-US | 965 | 0 | 216 | 83 | 10 | 0.089 | 0.061 | 0.067 | 0.070 | 0.6881 | 0.4914 | 0.7249 | Decreasing |
| OV-AU::SP101588 | Ovary | OV-AU | 410 | 53 | 47 | 27 | 13 | 0.022 | 0.020 | 0.030 | 0.005 | 0.6836 | 0.4942 | 0.7283 | Decreasing |
| SKCM-US::SP83099 | Skin | SKCM-US | 255 | 0 | 50 | 14 | 12 | 0.014 | 0.016 | 0.006 | 0.011 | 0.6820 | 0.4953 | 0.7291 | Decreasing |
| STAD-US::SP84982 | Stomach | STAD-US | 53700 | 0 | 0 | 30 | 6 | 0.028 | 0.020 | 0.020 | 0.042 | -0.6798 | 0.4966 | 0.7303 | Increasing |
| RECA-EU::SP103619 | Kidney | RECA-EU | 218 | 0 | 36 | 13 | 6 | 0.010 | 0.007 | 0.011 | 0.015 | -0.6740 | 0.5003 | 0.7350 | Increasing |
| MALY-DE::SP116606 | Blood | MALY-DE | 433 | 111 | 59 | 44 | 11 | 0.045 | 0.038 | 0.023 | 0.036 | 0.6725 | 0.5012 | 0.7356 | Decreasing |
| SKCM-US::SP83382 | Skin | SKCM-US | 614 | 0 | 179 | 63 | 17 | 0.044 | 0.057 | 0.054 | 0.061 | -0.6703 | 0.5027 | 0.7363 | Increasing |
| BLCA-US::SP953 | Bladder | BLCA-US | 895 | 0 | 0 | 19 | 13 | 0.024 | 0.005 | 0.026 | 0.006 | 0.6703 | 0.5026 | 0.7363 | Increasing |
| PACA-AU::SP107805 | Pancreas | PACA-AU | 539 | 0 | 122 | 19 | 10 | 0.011 | 0.014 | 0.018 | 0.017 | -0.6675 | 0.5045 | 0.7381 | Increasing |
| BRCA-US::SP5393 | Breast | BRCA-US | 590 | 0 | 96 | 25 | 8 | 0.017 | 0.021 | 0.022 | 0.026 | -0.6650 | 0.5060 | 0.7391 | Increasing |
| LINC-JP::SP98869 | Liver | LIHC-US | 501 | 0 | 64 | 24 | 12 | 0.028 | 0.005 | 0.023 | 0.036 | -0.6649 | 0.5061 | 0.7391 | Increasing |
| LINC-JP::SP98855 | Liver | LIHC-US | 489 | 0 | 68 | 22 | 12 | 0.025 | 0.018 | 0.010 | 0.024 | 0.6632 | 0.5072 | 0.7399 | Decreasing |
| LIRI-JP::SP99133 | Liver | LIRI-JP | 478 | 0 | 89 | 35 | 10 | 0.019 | 0.038 | 0.025 | 0.035 | -0.6621 | 0.5079 | 0.7401 | Increasing |
| KIRC-US::SP37369 | Kidney | KIRC-US | 476 | 0 | 75 | 27 | 10 | 0.018 | 0.021 | 0.028 | 0.025 | -0.6612 | 0.5085 | 0.7403 | Increasing |

| | | | | | | | | | | | | | | | |
|---|---|---|---|---|---|---|---|---|---|---|---|---|---|---|---|
| PRAD-UK::SP114946 | Prostate | PRAD-US | 649 | 0 | 0 | 12 | 9 | 0.011 | 0.005 | 0.012 | 0.015 | -0.6588 | 0.5100 | 0.7406 | Increasing |
| BRCA-US::SP10084 | Breast | BRCA-US | 686 | 0 | 190 | 45 | 11 | 0.034 | 0.034 | 0.044 | 0.042 | -0.6584 | 0.5103 | 0.7406 | Increasing |
| PRAD-UK::SP111152 | Prostate | PRAD-US | 286 | 0 | 43 | 12 | 10.5 | 0.007 | 0.011 | 0.009 | 0.015 | -0.6588 | 0.5100 | 0.7406 | Increasing |
| PRAD-UK::SP111126 | Prostate | PRAD-US | 262 | 0 | 58 | 27 | 11 | 0.026 | 0.016 | 0.021 | 0.035 | -0.6569 | 0.5112 | 0.7413 | Increasing |
| LINC-JP::SP98847 | Liver | LIHC-US | 479 | 0 | 72 | 30 | 11 | 0.022 | 0.021 | 0.035 | 0.024 | -0.6531 | 0.5137 | 0.7441 | Increasing |
| OV-AU::SP101604 | Ovary | OV-AU | 447 | 177 | 108 | 85 | 21 | 0.067 | 0.064 | 0.054 | 0.093 | -0.6518 | 0.5145 | 0.7446 | Increasing |
| LIRI-JP::SP99293 | Liver | LIRI-JP | 317 | 0 | 34 | 11 | 12.5 | 0.006 | 0.018 | 0.003 | 0.006 | 0.6458 | 0.5184 | 0.7478 | Decreasing |
| COAD-US::SP19983 | CRC | COAD-US | 2329 | 0 | 0 | 22 | 7.5 | 0.024 | 0.016 | 0.017 | 0.017 | 0.6452 | 0.5188 | 0.7478 | Decreasing |
| BRCA-US::SP3016 | Breast | BRCA-US | 718 | 226 | 132 | 71 | 11 | 0.055 | 0.055 | 0.069 | 0.064 | -0.6460 | 0.5183 | 0.7478 | Increasing |
| UCEC-US::SP90893 | Endometrium | UCEC-US | 456 | 0 | 82 | 27 | 9.5 | 0.016 | 0.026 | 0.027 | 0.023 | -0.6464 | 0.5180 | 0.7478 | Increasing |
| LIRI-JP::SP107078 | Liver | LIRI-JP | 326 | 0 | 64 | 21 | 11 | 0.013 | 0.020 | 0.016 | 0.023 | -0.6428 | 0.5204 | 0.7491 | Increasing |
| PACA-CA::SP117476 | Pancreas | PACA-CA | 508 | 0 | 0 | 22 | 8.5 | 0.038 | 0.000 | 0.014 | 0.022 | 0.6422 | 0.5207 | 0.7491 | Decreasing |
| OV-AU::SP101654 | Ovary | OV-AU | 619 | 93 | 0 | 30 | 11 | 0.016 | 0.039 | 0.015 | 0.016 | 0.6397 | 0.5224 | 0.7507 | Decreasing |
| SKCM-US::SP82836 | Skin | SKCM-US | 426 | 0 | 54 | 14 | 12.5 | 0.007 | 0.010 | 0.022 | 0.006 | -0.6388 | 0.5229 | 0.7508 | Increasing |
| PACA-CA::SP117745 | Pancreas | PACA-CA | 1088 | 0 | 0 | 12 | 11 | 0.004 | 0.014 | 0.006 | 0.013 | -0.6299 | 0.5287 | 0.7515 | Increasing |
| LIRI-JP::SP98933 | Liver | LIRI-JP | 496 | 0 | 58 | 24 | 11 | 0.013 | 0.025 | 0.019 | 0.023 | -0.6293 | 0.5292 | 0.7515 | Increasing |
| RECA-EU::SP103436 | Kidney | RECA-EU | 517 | 0 | 61 | 18 | 10 | 0.013 | 0.012 | 0.014 | 0.020 | -0.6311 | 0.5280 | 0.7515 | Increasing |
| OV-AU::SP101600 | Ovary | OV-AU | 210 | 0 | 36 | 17 | 14 | 0.010 | 0.016 | 0.009 | 0.021 | -0.6297 | 0.5289 | 0.7515 | Increasing |
| RECA-EU::SP103515 | Kidney | RECA-EU | 845 | 0 | 127 | 55 | 9.5 | 0.030 | 0.049 | 0.048 | 0.040 | -0.6344 | 0.5258 | 0.7515 | Increasing |
| LUAD-US::SP55509 | Lung | LUAD-US | 2181 | 0 | 0 | 20 | 12 | 0.019 | 0.011 | 0.016 | 0.027 | -0.6339 | 0.5261 | 0.7515 | Increasing |
| BRCA-US::SP5473 | Breast | BRCA-US | 485 | 0 | 142 | 23 | 10 | 0.024 | 0.011 | 0.019 | 0.032 | -0.6297 | 0.5289 | 0.7515 | Increasing |
| STAD-US::SP85130 | Stomach | STAD-US | 1616 | 0 | 0 | 36 | 9 | 0.031 | 0.036 | 0.029 | 0.021 | 0.6315 | 0.5277 | 0.7515 | Decreasing |
| PACA-CA::SP125714 | Pancreas | PACA-CA | 605 | 0 | 0 | 12 | 10 | 0.011 | 0.006 | 0.006 | 0.016 | -0.6299 | 0.5287 | 0.7515 | Increasing |
| UCEC-US::SP90629 | Endometrium | UCEC-US | 934 | 0 | 133 | 39 | 12 | 0.029 | 0.034 | 0.033 | 0.041 | -0.6367 | 0.5243 | 0.7515 | Increasing |
| BRCA-UK::SP116331 | Breast | BRCA-US | 428 | 0 | 89 | 20 | 8.5 | 0.014 | 0.026 | 0.013 | 0.011 | 0.6328 | 0.5269 | 0.7515 | Decreasing |
| PACA-AU::SP76057 | Pancreas | PACA-AU | 579 | 0 | 154 | 16 | 13 | 0.011 | 0.011 | 0.012 | 0.017 | -0.6248 | 0.5321 | 0.7520 | Increasing |
| LIHC-US::SP49433 | Liver | LIHC-US | 381 | 0 | 52 | 16 | 12 | 0.009 | 0.016 | 0.013 | 0.018 | -0.6252 | 0.5318 | 0.7520 | Increasing |
| STAD-US::SP85230 | Stomach | STAD-US | 1054 | 412 | 287 | 180 | 16 | 0.175 | 0.143 | 0.134 | 0.165 | 0.6248 | 0.5321 | 0.7520 | Decreasing |
| PACA-AU::SP69705 | Pancreas | PACA-AU | 504 | 0 | 0 | 16 | 12.5 | 0.011 | 0.011 | 0.012 | 0.017 | -0.6248 | 0.5321 | 0.7520 | Increasing |
| BRCA-US::SP6673 | Breast | BRCA-US | 2410 | 1174 | 666 | 464 | 12 | 0.417 | 0.388 | 0.393 | 0.376 | 0.6267 | 0.5309 | 0.7520 | Decreasing |
| PACA-AU::SP76915 | Pancreas | PACA-AU | 492 | 132 | 0 | 19 | 12 | 0.018 | 0.016 | 0.012 | 0.013 | 0.6202 | 0.5351 | 0.7553 | Decreasing |
| OV-US::SP67428 | Ovary | OV-US | 823 | 424 | 160 | 146 | 12 | 0.128 | 0.142 | 0.092 | 0.133 | 0.6197 | 0.5355 | 0.7553 | Decreasing |
| STAD-US::SP84922 | Stomach | STAD-US | 2336 | 0 | 0 | 45 | 13 | 0.034 | 0.042 | 0.029 | 0.053 | -0.6179 | 0.5366 | 0.7554 | Increasing |
| MELA-AU::SP124444 | Skin | SKCM-US | 407 | 0 | 55 | 13 | 10 | 0.007 | 0.021 | 0.006 | 0.006 | 0.6180 | 0.5366 | 0.7554 | Decreasing |
| BRCA-US::SP2766 | Breast | BRCA-US | 790 | 350 | 354 | 168 | 18 | 0.131 | 0.145 | 0.145 | 0.154 | -0.6166 | 0.5375 | 0.7559 | Increasing |
| MELA-AU::SP124403 | Skin | SKCM-US | 155 | 0 | 55 | 17 | 14 | 0.017 | 0.016 | 0.013 | 0.011 | 0.6145 | 0.5389 | 0.7562 | Decreasing |
| KIRP-US::SP97243 | Kidney | KIRP-US | 730 | 0 | 146 | 54 | 10 | 0.042 | 0.045 | 0.044 | 0.057 | -0.6140 | 0.5392 | 0.7562 | Increasing |
| RECA-EU::SP103523 | Kidney | RECA-EU | 417 | 0 | 80 | 33 | 9 | 0.036 | 0.019 | 0.026 | 0.025 | 0.6152 | 0.5384 | 0.7562 | Decreasing |
| LIHC-US::SP115498 | Liver | LIHC-US | 934 | 0 | 0 | 19 | 12 | 0.016 | 0.010 | 0.023 | 0.018 | -0.6105 | 0.5415 | 0.7587 | Increasing |
| LUAD-US::SP52284 | Lung | LUAD-US | 437 | 0 | 58 | 23 | 12 | 0.016 | 0.019 | 0.023 | 0.022 | -0.6090 | 0.5425 | 0.7593 | Increasing |
| OV-US::SP59761 | Ovary | OV-US | 371 | 61 | 50 | 24 | 13 | 0.029 | 0.013 | 0.023 | 0.017 | 0.6056 | 0.5448 | 0.7612 | Decreasing |
| LIRI-JP::SP112182 | Liver | LIRI-JP | 465 | 0 | 64 | 36 | 13 | 0.019 | 0.038 | 0.032 | 0.029 | -0.6054 | 0.5449 | 0.7612 | Increasing |
| PRAD-CA::SP102647 | Prostate | PRAD-US | 415 | 0 | 56 | 21 | 12.5 | 0.011 | 0.021 | 0.018 | 0.020 | -0.6032 | 0.5464 | 0.7625 | Increasing |
| GACA-CN::SP135234 | Stomach | STAD-US | 1152 | 0 | 0 | 23 | 13 | 0.018 | 0.017 | 0.020 | 0.027 | -0.6019 | 0.5472 | 0.7629 | Increasing |
| LIRI-JP::SP107146 | Liver | LIRI-JP | 599 | 0 | 87 | 33 | 12.5 | 0.035 | 0.020 | 0.035 | 0.017 | 0.6004 | 0.5482 | 0.7636 | Decreasing |
| RECA-EU::SP103005 | Kidney | RECA-EU | 460 | 0 | 81 | 28 | 10 | 0.013 | 0.031 | 0.014 | 0.030 | -0.5994 | 0.5489 | 0.7639 | Increasing |
| RECA-EU::SP102873 | Kidney | RECA-EU | 498 | 0 | 81 | 33 | 9.5 | 0.010 | 0.033 | 0.040 | 0.010 | -0.5951 | 0.5518 | 0.7670 | Increasing |
| BRCA-US::SP4472 | Breast | BRCA-US | 827 | 412 | 343 | 196 | 18 | 0.179 | 0.169 | 0.154 | 0.164 | 0.5944 | 0.5522 | 0.7670 | Decreasing |
| KIRC-US::SP39997 | Kidney | KIRC-US | 740 | 0 | 105 | 38 | 9 | 0.044 | 0.027 | 0.028 | 0.034 | 0.5920 | 0.5538 | 0.7673 | Decreasing |
| MELA-AU::SP124456 | Skin | SKCM-US | 449 | 0 | 227 | 62 | 18 | 0.044 | 0.052 | 0.063 | 0.050 | -0.5912 | 0.5544 | 0.7673 | Increasing |

| | | | | | | | | | | | | | | | |
|---|---|---|---|---|---|---|---|---|---|---|---|---|---|---|---|
| LIRI-JP::SP50111 | Liver | LIRI-JP | 446 | 0 | 0 | 15 | 12 | 0.019 | 0.010 | 0.006 | 0.017 | 0.5911 | 0.5544 | 0.7673 | Decreasing |
| HNSC-US::SP30332 | Head&Neck | HNSC-US | 2387 | 0 | 0 | 35 | 10 | 0.027 | 0.033 | 0.042 | 0.006 | 0.5909 | 0.5546 | 0.7673 | Decreasing |
| BLCA-US::SP963 | Bladder | BLCA-US | 534 | 0 | 117 | 36 | 10.5 | 0.034 | 0.021 | 0.033 | 0.042 | -0.5896 | 0.5554 | 0.7678 | Increasing |
| BRCA-US::SP4820 | Breast | BRCA-US | 1359 | 801 | 356 | 245 | 11 | 0.241 | 0.198 | 0.173 | 0.238 | 0.5863 | 0.5577 | 0.7690 | Decreasing |
| PACA-CA::SP125786 | Pancreas | PACA-CA | 573 | 356 | 148 | 109 | 13 | 0.094 | 0.078 | 0.093 | 0.073 | 0.5868 | 0.5573 | 0.7690 | Decreasing |
| BRCA-UK::SP2153 | Breast | BRCA-US | 709 | 121 | 0 | 26 | 12 | 0.017 | 0.024 | 0.022 | 0.026 | -0.5859 | 0.5579 | 0.7690 | Increasing |
| HNSC-US::SP31046 | Head&Neck | HNSC-US | 186 | 0 | 32 | 11 | 9 | 0.015 | 0.000 | 0.007 | 0.023 | -0.5824 | 0.5603 | 0.7697 | Increasing |
| PRAD-UK::SP114945 | Prostate | PRAD-US | 1264 | 0 | 325 | 105 | 9 | 0.093 | 0.082 | 0.079 | 0.115 | -0.5807 | 0.5615 | 0.7697 | Increasing |
| PRAD-UK::SP114943 | Prostate | PRAD-US | 1303 | 0 | 331 | 105 | 9 | 0.093 | 0.082 | 0.079 | 0.115 | -0.5807 | 0.5615 | 0.7697 | Increasing |
| PACA-CA::SP125802 | Pancreas | PACA-CA | 487 | 0 | 0 | 17 | 11 | 0.015 | 0.011 | 0.006 | 0.022 | -0.5797 | 0.5621 | 0.7697 | Increasing |
| UCEC-US::SP92364 | Endometrium | UCEC-US | 62255 | 0 | 0 | 34 | 6 | 0.025 | 0.028 | 0.030 | 0.035 | -0.5815 | 0.5609 | 0.7697 | Increasing |
| LIRI-JP::SP50133 | Liver | LIRI-JP | 364 | 0 | 46 | 19 | 10 | 0.019 | 0.008 | 0.019 | 0.023 | -0.5789 | 0.5627 | 0.7697 | Increasing |
| HNSC-US::SP30843 | Head&Neck | HNSC-US | 174 | 0 | 32 | 11 | 10.5 | 0.003 | 0.016 | 0.007 | 0.012 | -0.5824 | 0.5603 | 0.7697 | Decreasing |
| STAD-US::SP105159 | Stomach | STAD-US | 675 | 0 | 0 | 14 | 11 | 0.015 | 0.011 | 0.010 | 0.011 | 0.5791 | 0.5625 | 0.7697 | Decreasing |
| UCEC-US::SP94917 | Endometrium | UCEC-US | 18292 | 0 | 0 | 20 | 8 | 0.013 | 0.028 | 0.010 | 0.012 | 0.5774 | 0.5637 | 0.7703 | Decreasing |
| LUSC-US::SP57846 | Lung | LUSC-US | 1638 | 0 | 0 | 27 | 10 | 0.019 | 0.026 | 0.016 | 0.034 | -0.5728 | 0.5668 | 0.7739 | Increasing |
| MELA-AU::SP124412 | Skin | SKCM-US | 472 | 0 | 167 | 42 | 13 | 0.041 | 0.034 | 0.038 | 0.028 | 0.5712 | 0.5679 | 0.7746 | Decreasing |
| BLCA-US::SP1677 | Bladder | BLCA-US | 634 | 0 | 80 | 30 | 13 | 0.028 | 0.018 | 0.026 | 0.036 | -0.5682 | 0.5699 | 0.7766 | Increasing |
| OV-AU::SP101642 | Ovary | OV-AU | 1496 | 712 | 428 | 315 | 17 | 0.264 | 0.204 | 0.272 | 0.264 | -0.5648 | 0.5722 | 0.7791 | Increasing |
| PACA-CA::SP117158 | Pancreas | PACA-CA | 311 | 0 | 92 | 15 | 11 | 0.011 | 0.017 | 0.008 | 0.010 | 0.5632 | 0.5733 | 0.7798 | Decreasing |
| OV-AU::SP101628 | Ovary | OV-AU | 306 | 0 | 57 | 23 | 11.5 | 0.013 | 0.018 | 0.024 | 0.016 | -0.5614 | 0.5745 | 0.7807 | Increasing |
| LIHC-US::SP49175 | Liver | LIHC-US | 649 | 0 | 0 | 15 | 13.5 | 0.009 | 0.021 | 0.010 | 0.006 | 0.5592 | 0.5760 | 0.7821 | Decreasing |
| MELA-AU::SP124447 | Skin | SKCM-US | 3909 | 0 | 1215 | 449 | 17 | 0.393 | 0.391 | 0.365 | 0.372 | 0.5570 | 0.5776 | 0.7834 | Decreasing |
| OV-US::SP65807 | Ovary | OV-US | 1122 | 725 | 317 | 239 | 15 | 0.242 | 0.188 | 0.155 | 0.255 | 0.5555 | 0.5785 | 0.7840 | Decreasing |
| PACA-CA::SP125809 | Pancreas | PACA-CA | 810 | 0 | 0 | 22 | 10 | 0.019 | 0.008 | 0.023 | 0.019 | -0.5539 | 0.5796 | 0.7848 | Increasing |
| LIRI-JP::SP107091 | Liver | LIRI-JP | 660 | 0 | 81 | 37 | 10 | 0.029 | 0.030 | 0.029 | 0.041 | -0.5503 | 0.5821 | 0.7852 | Increasing |
| MELA-AU::SP124452 | Skin | SKCM-US | 656 | 0 | 278 | 76 | 15 | 0.065 | 0.070 | 0.066 | 0.050 | 0.5524 | 0.5807 | 0.7852 | Decreasing |
| SKCM-US::SP83019 | Skin | SKCM-US | 623 | 0 | 97 | 33 | 13 | 0.031 | 0.021 | 0.028 | 0.039 | -0.5506 | 0.5819 | 0.7852 | Increasing |
| LIHC-US::SP98313 | Liver | LIHC-US | 635 | 0 | 0 | 14 | 11 | 0.009 | 0.010 | 0.016 | 0.012 | -0.5518 | 0.5811 | 0.7852 | Increasing |
| LUSC-US::SP57901 | Lung | LUSC-US | 1359 | 0 | 0 | 39 | 11 | 0.019 | 0.045 | 0.036 | 0.028 | -0.5489 | 0.5831 | 0.7857 | Increasing |
| STAD-US::SP83967 | Stomach | STAD-US | 482 | 0 | 55 | 13 | 8 | 0.006 | 0.014 | 0.013 | 0.011 | -0.5452 | 0.5856 | 0.7876 | Increasing |
| OV-AU::SP101732 | Ovary | OV-AU | 895 | 373 | 279 | 155 | 16 | 0.111 | 0.125 | 0.117 | 0.135 | -0.5460 | 0.5850 | 0.7876 | Increasing |
| LUSC-US::SP57267 | Lung | LUSC-US | 2245 | 0 | 0 | 57 | 12 | 0.051 | 0.032 | 0.068 | 0.045 | -0.5438 | 0.5866 | 0.7876 | Increasing |
| PRAD-UK::SP111156 | Prostate | PRAD-US | 214 | 0 | 41 | 19 | 11 | 0.019 | 0.016 | 0.018 | 0.010 | 0.5440 | 0.5864 | 0.7876 | Decreasing |
| GBM-US::SP24135 | Brain | GBM-US | 1077 | 0 | 289 | 90 | 9 | 0.086 | 0.061 | 0.080 | 0.092 | -0.5427 | 0.5874 | 0.7878 | Increasing |
| MELA-AU::SP124389 | Skin | SKCM-US | 431 | 0 | 0 | 15 | 12.5 | 0.014 | 0.008 | 0.016 | 0.017 | -0.5336 | 0.5936 | 0.7897 | Increasing |
| KIRC-US::SP37636 | Kidney | KIRC-US | 313 | 0 | 56 | 24 | 11 | 0.018 | 0.021 | 0.015 | 0.029 | -0.5349 | 0.5927 | 0.7897 | Increasing |
| SKCM-US::SP82614 | Skin | SKCM-US | 549 | 0 | 141 | 44 | 16 | 0.027 | 0.044 | 0.038 | 0.039 | -0.5365 | 0.5916 | 0.7897 | Increasing |
| LIRI-JP::SP112125 | Liver | LIRI-JP | 401 | 0 | 67 | 23 | 13 | 0.022 | 0.023 | 0.010 | 0.023 | 0.5389 | 0.5900 | 0.7897 | Decreasing |
| PACA-AU::SP70313 | Pancreas | PACA-AU | 397 | 0 | 104 | 15 | 10 | 0.011 | 0.011 | 0.009 | 0.017 | -0.5295 | 0.5964 | 0.7897 | Increasing |
| COAD-US::SP96126 | CRC | COAD-US | 3401 | 0 | 0 | 20 | 11 | 0.015 | 0.014 | 0.023 | 0.017 | -0.5305 | 0.5958 | 0.7897 | Increasing |
| PRAD-CA::SP112975 | Prostate | PRAD-US | 374 | 0 | 47 | 13 | 12 | 0.007 | 0.013 | 0.009 | 0.015 | -0.5266 | 0.5985 | 0.7897 | Increasing |
| PRAD-CA::SP112867 | Prostate | PRAD-US | 268 | 0 | 34 | 13 | 7 | 0.011 | 0.005 | 0.018 | 0.010 | -0.5266 | 0.5985 | 0.7897 | Increasing |
| LIHC-US::SP49551 | Liver | LIHC-US | 484 | 0 | 48 | 17 | 8.5 | 0.016 | 0.013 | 0.006 | 0.030 | -0.5396 | 0.5895 | 0.7897 | Increasing |
| PACA-CA::SP125705 | Pancreas | PACA-CA | 517 | 0 | 0 | 11 | 12 | 0.008 | 0.008 | 0.006 | 0.013 | -0.5326 | 0.5943 | 0.7897 | Increasing |
| UCEC-US::SP91666 | Endometrium | UCEC-US | 553 | 0 | 0 | 17 | 15 | 0.013 | 0.013 | 0.017 | 0.017 | -0.5306 | 0.5957 | 0.7897 | Increasing |
| PACA-AU::SP76001 | Pancreas | PACA-AU | 269 | 0 | 72 | 15 | 15 | 0.007 | 0.014 | 0.012 | 0.013 | -0.5295 | 0.5964 | 0.7897 | Increasing |
| GBM-US::SP27339 | Brain | GBM-US | 313 | 0 | 0 | 12 | 5 | 0.000 | 0.014 | 0.015 | 0.005 | -0.5334 | 0.5938 | 0.7897 | Increasing |
| LIHC-US::SP98065 | Liver | LIHC-US | 617 | 0 | 97 | 41 | 12 | 0.038 | 0.039 | 0.026 | 0.036 | 0.5334 | 0.5938 | 0.7897 | Decreasing |
| RECA-EU::SP102989 | Kidney | RECA-EU | 876 | 0 | 147 | 57 | 10 | 0.046 | 0.038 | 0.045 | 0.055 | -0.5310 | 0.5954 | 0.7897 | Increasing |

| | | | | | | | | | | | | | | |
|---|---|---|---|---|---|---|---|---|---|---|---|---|---|---|
| BRCA-UK::SP2349 | Breast | BRCA-US | 317 | 0 | 101 | 12 | 10 | 0.007 | 0.008 | 0.019 | 0.005 | -0.5287 | 0.5970 | 0.7897 | Increasing |
| PRAD-UK::SP111129 | Prostate | PRAD-US | 399 | 0 | 41 | 13 | 7 | 0.007 | 0.013 | 0.009 | 0.015 | -0.5266 | 0.5985 | 0.7897 | Increasing |
| BRCA-UK::SP116349 | Breast | BRCA-US | 201 | 0 | 46 | 12 | 9 | 0.007 | 0.011 | 0.013 | 0.011 | -0.5287 | 0.5970 | 0.7897 | Increasing |
| RECA-EU::SP103065 | Kidney | RECA-EU | 1167 | 0 | 151 | 65 | 9 | 0.046 | 0.049 | 0.054 | 0.055 | -0.5216 | 0.6019 | 0.7934 | Increasing |
| UCEC-US::SP89389 | Endometrium | UCEC-US | 554 | 0 | 79 | 23 | 12 | 0.016 | 0.023 | 0.013 | 0.029 | -0.5205 | 0.6027 | 0.7937 | Increasing |
| OV-US::SP61713 | Ovary | OV-US | 756 | 398 | 358 | 171 | 20 | 0.141 | 0.175 | 0.105 | 0.157 | 0.5187 | 0.6040 | 0.7945 | Decreasing |
| OV-AU::SP101521 | Ovary | OV-AU | 352 | 0 | 45 | 13 | 12.5 | 0.006 | 0.014 | 0.006 | 0.016 | -0.5182 | 0.6043 | 0.7945 | Increasing |
| PRAD-UK::SP114937 | Prostate | PRAD-US | 1200 | 0 | 320 | 102 | 9 | 0.093 | 0.079 | 0.073 | 0.115 | -0.5083 | 0.6112 | 0.7974 | Increasing |
| LIHC-US::SP98092 | Liver | LIHC-US | 558 | 0 | 87 | 32 | 10.5 | 0.041 | 0.018 | 0.016 | 0.042 | 0.5138 | 0.6074 | 0.7974 | Decreasing |
| PRAD-CA::SP112791 | Prostate | PRAD-US | 317 | 0 | 42 | 11 | 9 | 0.000 | 0.016 | 0.012 | 0.005 | -0.5072 | 0.6120 | 0.7974 | Increasing |
| LIRI-JP::SP112201 | Liver | LIRI-JP | 213 | 0 | 31 | 17 | 10 | 0.010 | 0.018 | 0.013 | 0.017 | -0.5096 | 0.6103 | 0.7974 | Increasing |
| RECA-EU::SP103340 | Kidney | RECA-EU | 739 | 0 | 107 | 47 | 8 | 0.033 | 0.040 | 0.028 | 0.050 | -0.5127 | 0.6082 | 0.7974 | Increasing |
| LIRI-JP::SP112283 | Liver | LIRI-JP | 212 | 0 | 42 | 18 | 9.5 | 0.016 | 0.020 | 0.006 | 0.017 | 0.5072 | 0.6120 | 0.7974 | Decreasing |
| PACA-AU::SP77203 | Pancreas | PACA-AU | 2502 | 1166 | 0 | 342 | 16 | 0.330 | 0.242 | 0.199 | 0.317 | 0.5068 | 0.6123 | 0.7974 | Decreasing |
| LIRI-JP::SP99101 | Liver | LIRI-JP | 532 | 0 | 0 | 17 | 12 | 0.016 | 0.013 | 0.006 | 0.029 | -0.5096 | 0.6103 | 0.7974 | Increasing |
| LIRI-JP::SP50131 | Liver | LIRI-JP | 1341 | 0 | 246 | 91 | 12 | 0.083 | 0.058 | 0.086 | 0.087 | -0.5061 | 0.6128 | 0.7974 | Increasing |
| LUAD-US::SP53810 | Lung | LUAD-US | 5445 | 0 | 0 | 33 | 13 | 0.028 | 0.033 | 0.026 | 0.022 | 0.5109 | 0.6094 | 0.7974 | Decreasing |
| KIRP-US::SP43688 | Kidney | KIRP-US | 902 | 0 | 142 | 51 | 9 | 0.035 | 0.048 | 0.044 | 0.046 | -0.5055 | 0.6132 | 0.7974 | Increasing |
| OV-AU::SP101526 | Ovary | OV-AU | 543 | 96 | 120 | 42 | 11 | 0.041 | 0.027 | 0.033 | 0.031 | 0.5110 | 0.6093 | 0.7974 | Decreasing |
| OV-US::SP62110 | Ovary | OV-US | 535 | 0 | 119 | 26 | 13 | 0.033 | 0.015 | 0.016 | 0.029 | 0.5009 | 0.6164 | 0.8006 | Decreasing |
| LIRI-JP::SP107044 | Liver | LIRI-JP | 241 | 0 | 33 | 20 | 11 | 0.019 | 0.008 | 0.025 | 0.017 | -0.5005 | 0.6167 | 0.8006 | Increasing |
| SKCM-US::SP83146 | Skin | SKCM-US | 1707 | 0 | 451 | 155 | 15 | 0.120 | 0.137 | 0.132 | 0.139 | -0.4981 | 0.6184 | 0.8020 | Increasing |
| LIRI-JP::SP99141 | Liver | LIRI-JP | 615 | 0 | 67 | 23 | 12 | 0.016 | 0.023 | 0.013 | 0.029 | -0.4954 | 0.6203 | 0.8038 | Increasing |
| LIRI-JP::SP99121 | Liver | LIRI-JP | 784 | 0 | 93 | 35 | 11 | 0.022 | 0.033 | 0.032 | 0.029 | -0.4944 | 0.6210 | 0.8040 | Increasing |
| KIRC-US::SP36498 | Kidney | KIRC-US | 689 | 0 | 93 | 43 | 9 | 0.036 | 0.032 | 0.037 | 0.044 | -0.4930 | 0.6220 | 0.8045 | Increasing |
| LIRI-JP::SP98921 | Liver | LIRI-JP | 391 | 0 | 65 | 29 | 13.5 | 0.029 | 0.015 | 0.025 | 0.035 | -0.4922 | 0.6226 | 0.8046 | Increasing |
| LINC-JP::SP98879 | Liver | LIHC-US | 548 | 0 | 106 | 39 | 12 | 0.025 | 0.037 | 0.039 | 0.030 | -0.4914 | 0.6232 | 0.8046 | Increasing |
| MALY-DE::SP124971 | Blood | MALY-DE | 1554 | 0 | 0 | 22 | 11 | 0.006 | 0.022 | 0.012 | 0.021 | -0.4900 | 0.6241 | 0.8048 | Increasing |
| READ-US::SP80423 | CRC | READ-US | 1163 | 0 | 0 | 15 | 7 | 0.009 | 0.011 | 0.023 | 0.006 | -0.4896 | 0.6244 | 0.8048 | Increasing |
| STAD-US::SP105253 | Stomach | STAD-US | 2113 | 0 | 0 | 36 | 11 | 0.025 | 0.028 | 0.046 | 0.021 | -0.4887 | 0.6251 | 0.8049 | Increasing |
| SKCM-US::SP82756 | Skin | SKCM-US | 229 | 0 | 40 | 15 | 14 | 0.014 | 0.013 | 0.016 | 0.006 | 0.4872 | 0.6261 | 0.8056 | Decreasing |
| COAD-US::SP96120 | CRC | COAD-US | 2073 | 0 | 0 | 23 | 10 | 0.018 | 0.024 | 0.020 | 0.011 | 0.4847 | 0.6279 | 0.8064 | Decreasing |
| LUAD-US::SP50611 | Lung | LUAD-US | 1895 | 0 | 0 | 11 | 10.5 | 0.003 | 0.016 | 0.007 | 0.011 | -0.4848 | 0.6278 | 0.8064 | Decreasing |
| MELA-AU::SP124376 | Skin | SKCM-US | 527 | 0 | 123 | 40 | 16 | 0.044 | 0.026 | 0.035 | 0.033 | 0.4830 | 0.6291 | 0.8066 | Decreasing |
| SKCM-US::SP82780 | Skin | SKCM-US | 394 | 0 | 82 | 26 | 13.5 | 0.010 | 0.039 | 0.006 | 0.033 | -0.4829 | 0.6292 | 0.8066 | Decreasing |
| UCEC-US::SP94236 | Endometrium | UCEC-US | 514 | 0 | 95 | 18 | 10.5 | 0.010 | 0.028 | 0.007 | 0.012 | 0.4781 | 0.6326 | 0.8082 | Decreasing |
| PRAD-CA::SP112801 | Prostate | PRAD-US | 468 | 0 | 0 | 12 | 10 | 0.011 | 0.013 | 0.006 | 0.010 | 0.4772 | 0.6332 | 0.8082 | Decreasing |
| MELA-AU::SP124295 | Skin | SKCM-US | 585 | 0 | 0 | 13 | 14 | 0.014 | 0.010 | 0.000 | 0.028 | -0.4785 | 0.6323 | 0.8082 | Increasing |
| OV-AU::SP102090 | Ovary | OV-AU | 378 | 0 | 46 | 16 | 12 | 0.010 | 0.014 | 0.012 | 0.016 | -0.4791 | 0.6319 | 0.8082 | Increasing |
| PRAD-CA::SP112849 | Prostate | PRAD-US | 306 | 0 | 33 | 12 | 10 | 0.015 | 0.011 | 0.003 | 0.015 | 0.4772 | 0.6332 | 0.8082 | Decreasing |
| KIRC-US::SP43201 | Kidney | KIRC-US | 561 | 0 | 128 | 48 | 11 | 0.022 | 0.054 | 0.049 | 0.029 | -0.4751 | 0.6347 | 0.8095 | Increasing |
| PRAD-UK::SP114971 | Prostate | PRAD-US | 745 | 0 | 0 | 21 | 9 | 0.022 | 0.016 | 0.018 | 0.015 | 0.4703 | 0.6382 | 0.8131 | Decreasing |
| PACA-CA::SP118083 | Pancreas | PACA-CA | 586 | 119 | 112 | 36 | 11.5 | 0.038 | 0.025 | 0.020 | 0.032 | 0.4673 | 0.6403 | 0.8136 | Decreasing |
| LIHC-US::SP106677 | Liver | LIHC-US | 502 | 0 | 70 | 27 | 15 | 0.025 | 0.013 | 0.032 | 0.024 | -0.4674 | 0.6402 | 0.8136 | Increasing |
| COAD-US::SP96129 | CRC | COAD-US | 1754 | 0 | 0 | 12 | 9 | 0.009 | 0.008 | 0.013 | 0.011 | -0.4670 | 0.6405 | 0.8136 | Increasing |
| OV-AU::SP101540 | Ovary | OV-AU | 411 | 0 | 72 | 27 | 13 | 0.013 | 0.025 | 0.027 | 0.016 | -0.4666 | 0.6408 | 0.8136 | Increasing |
| ORCA-IN::SP117260 | Head&Neck | ORCA-IN | 382 | 0 | 60 | 28 | 12 | 0.032 | 0.018 | 0.016 | 0.030 | 0.4629 | 0.6434 | 0.8163 | Decreasing |
| BLCA-US::SP1086 | Bladder | BLCA-US | 460 | 0 | 68 | 19 | 12.5 | 0.009 | 0.024 | 0.013 | 0.018 | -0.4598 | 0.6457 | 0.8183 | Decreasing |
| PRAD-CA::SP112787 | Prostate | PRAD-US | 385 | 0 | 39 | 18 | 12 | 0.015 | 0.013 | 0.015 | 0.020 | -0.4591 | 0.6462 | 0.8183 | Increasing |
| GACA-CN::SP135393 | Stomach | STAD-US | 597 | 0 | 0 | 12 | 9 | 0.015 | 0.008 | 0.003 | 0.016 | 0.4570 | 0.6477 | 0.8195 | Decreasing |

| | | | | | | | | | | | | | | | |
|---|---|---|---|---|---|---|---|---|---|---|---|---|---|---|---|
| PACA-AU::SP71552 | Pancreas | PACA-AU | 747 | 0 | 226 | 28 | 11 | 0.022 | 0.027 | 0.018 | 0.020 | 0.4552 | 0.6490 | 0.8204 | Decreasing |
| LUSC-US::SP56754 | Lung | LUSC-US | 1094 | 0 | 0 | 23 | 13 | 0.010 | 0.029 | 0.020 | 0.017 | -0.4530 | 0.6506 | 0.8210 | Decreasing |
| BRCA-US::SP2997 | Breast | BRCA-US | 248 | 0 | 83 | 24 | 10 | 0.021 | 0.029 | 0.006 | 0.026 | 0.4531 | 0.6505 | 0.8210 | Decreasing |
| UCEC-US::SP90269 | Endometrium | UCEC-US | 492 | 0 | 0 | 13 | 11 | 0.010 | 0.016 | 0.010 | 0.006 | 0.4519 | 0.6514 | 0.8212 | Decreasing |
| UCEC-US::SP89651 | Endometrium | UCEC-US | 287 | 42 | 37 | 18 | 11 | 0.016 | 0.010 | 0.020 | 0.017 | -0.4504 | 0.6524 | 0.8212 | Increasing |
| OV-AU::SP102113 | Ovary | OV-AU | 539 | 218 | 215 | 123 | 18 | 0.108 | 0.102 | 0.060 | 0.124 | 0.4511 | 0.6519 | 0.8212 | Decreasing |
| OV-AU::SP101694 | Ovary | OV-AU | 496 | 90 | 74 | 37 | 13 | 0.038 | 0.018 | 0.039 | 0.021 | 0.4484 | 0.6538 | 0.8223 | Increasing |
| LIRI-JP::SP112221 | Liver | LIRI-JP | 236 | 0 | 0 | 12 | 10 | 0.003 | 0.018 | 0.006 | 0.012 | -0.4450 | 0.6563 | 0.8233 | Decreasing |
| LIRI-JP::SP99337 | Liver | LIRI-JP | 444 | 0 | 0 | 12 | 9 | 0.003 | 0.013 | 0.019 | 0.000 | -0.4450 | 0.6563 | 0.8233 | Increasing |
| LUSC-US::SP58991 | Lung | LUSC-US | 1483 | 0 | 187 | 29 | 10 | 0.026 | 0.018 | 0.029 | 0.028 | -0.4453 | 0.6561 | 0.8233 | Increasing |
| PACA-CA::SP125691 | Pancreas | PACA-CA | 1476 | 0 | 0 | 22 | 9 | 0.008 | 0.034 | 0.011 | 0.013 | 0.4429 | 0.6579 | 0.8245 | Decreasing |
| PACA-CA::SP117949 | Pancreas | PACA-CA | 925 | 0 | 0 | 18 | 16 | 0.015 | 0.017 | 0.011 | 0.013 | 0.4406 | 0.6595 | 0.8250 | Decreasing |
| PACA-CA::SP117291 | Pancreas | PACA-CA | 710 | 0 | 0 | 18 | 12 | 0.015 | 0.014 | 0.017 | 0.010 | 0.4406 | 0.6595 | 0.8250 | Decreasing |
| CESC-US::SP13084 | Cervix | CESC-US | 423 | 74 | 0 | 18 | 12.5 | 0.011 | 0.019 | 0.013 | 0.019 | -0.4384 | 0.6611 | 0.8256 | Increasing |
| KIRC-US::SP35412 | Kidney | KIRC-US | 710 | 39 | 111 | 55 | 10 | 0.062 | 0.035 | 0.046 | 0.049 | 0.4390 | 0.6607 | 0.8256 | Decreasing |
| PRAD-UK::SP114906 | Prostate | PRAD-US | 617 | 0 | 0 | 16 | 13 | 0.007 | 0.016 | 0.018 | 0.010 | -0.4328 | 0.6652 | 0.8299 | Increasing |
| UCEC-US::SP92332 | Endometrium | UCEC-US | 720 | 150 | 0 | 34 | 10 | 0.038 | 0.026 | 0.017 | 0.041 | 0.4319 | 0.6658 | 0.8299 | Decreasing |
| BRCA-UK::SP116361 | Breast | BRCA-US | 387 | 172 | 174 | 95 | 19 | 0.090 | 0.053 | 0.110 | 0.074 | -0.4314 | 0.6662 | 0.8299 | Increasing |
| PACA-AU::SP107774 | Pancreas | PACA-AU | 561 | 0 | 0 | 14 | 14 | 0.007 | 0.014 | 0.009 | 0.013 | -0.4282 | 0.6685 | 0.8299 | Increasing |
| MELA-AU::SP124409 | Skin | SKCM-US | 1158 | 0 | 150 | 46 | 14 | 0.048 | 0.034 | 0.019 | 0.072 | -0.4293 | 0.6677 | 0.8299 | Increasing |
| RECA-EU::SP103221 | Kidney | RECA-EU | 1323 | 0 | 181 | 83 | 10 | 0.069 | 0.045 | 0.091 | 0.055 | -0.4285 | 0.6683 | 0.8299 | Increasing |
| OV-AU::SP101544 | Ovary | OV-AU | 417 | 70 | 0 | 22 | 12 | 0.016 | 0.014 | 0.024 | 0.016 | -0.4290 | 0.6679 | 0.8299 | Increasing |
| BRCA-US::SP7692 | Breast | BRCA-US | 852 | 0 | 206 | 36 | 12.5 | 0.024 | 0.032 | 0.038 | 0.026 | -0.4255 | 0.6705 | 0.8310 | Increasing |
| PRAD-UK::SP114921 | Prostate | PRAD-US | 1190 | 0 | 239 | 90 | 10 | 0.097 | 0.058 | 0.082 | 0.075 | 0.4254 | 0.6706 | 0.8310 | Increasing |
| PRAD-UK::SP114973 | Prostate | PRAD-US | 737 | 0 | 0 | 23 | 10 | 0.019 | 0.018 | 0.018 | 0.025 | -0.4163 | 0.6772 | 0.8356 | Increasing |
| BRCA-US::SP9979 | Breast | BRCA-US | 159 | 0 | 46 | 18 | 14.5 | 0.007 | 0.021 | 0.019 | 0.011 | -0.4164 | 0.6771 | 0.8356 | Increasing |
| BRCA-US::SP2714 | Breast | BRCA-US | 1120 | 503 | 486 | 231 | 16 | 0.203 | 0.187 | 0.179 | 0.233 | -0.4167 | 0.6769 | 0.8356 | Increasing |
| GBM-US::SP23775 | Brain | GBM-US | 1025 | 0 | 312 | 97 | 10 | 0.096 | 0.066 | 0.110 | 0.056 | 0.4169 | 0.6767 | 0.8356 | Increasing |
| SKCM-US::SP83083 | Skin | SKCM-US | 596 | 0 | 111 | 42 | 15 | 0.055 | 0.026 | 0.016 | 0.061 | 0.4186 | 0.6755 | 0.8356 | Decreasing |
| LIHC-US::SP49187 | Liver | LIHC-US | 420 | 0 | 68 | 31 | 10 | 0.022 | 0.029 | 0.026 | 0.030 | -0.4153 | 0.6779 | 0.8359 | Increasing |
| BRCA-US::SP6813 | Breast | BRCA-US | 443 | 0 | 123 | 20 | 12 | 0.021 | 0.016 | 0.016 | 0.016 | 0.4136 | 0.6792 | 0.8367 | Decreasing |
| BRCA-UK::SP116371 | Breast | BRCA-US | 502 | 0 | 119 | 31 | 11 | 0.024 | 0.021 | 0.038 | 0.021 | -0.4095 | 0.6822 | 0.8397 | Increasing |
| OV-US::SP61453 | Ovary | OV-US | 270 | 0 | 58 | 21 | 10.5 | 0.010 | 0.025 | 0.016 | 0.017 | -0.4058 | 0.6849 | 0.8423 | Decreasing |
| PACA-CA::SP117216 | Pancreas | PACA-CA | 555 | 0 | 0 | 15 | 11 | 0.015 | 0.006 | 0.011 | 0.016 | -0.4025 | 0.6873 | 0.8439 | Increasing |
| PACA-CA::SP116968 | Pancreas | PACA-CA | 728 | 0 | 0 | 15 | 10 | 0.008 | 0.020 | 0.000 | 0.019 | -0.4025 | 0.6873 | 0.8439 | Decreasing |
| MALY-DE::SP116659 | Blood | MALY-DE | 878 | 0 | 0 | 17 | 10 | 0.006 | 0.018 | 0.006 | 0.018 | -0.3974 | 0.6910 | 0.8477 | Decreasing |
| UCEC-US::SP90245 | Endometrium | UCEC-US | 539 | 0 | 105 | 28 | 10 | 0.025 | 0.013 | 0.040 | 0.017 | -0.3963 | 0.6919 | 0.8480 | Increasing |
| BRCA-US::SP5808 | Breast | BRCA-US | 881 | 263 | 231 | 118 | 15 | 0.072 | 0.121 | 0.113 | 0.079 | -0.3943 | 0.6933 | 0.8484 | Decreasing |
| LUAD-US::SP51446 | Lung | LUAD-US | 412 | 0 | 64 | 19 | 12 | 0.022 | 0.014 | 0.010 | 0.022 | 0.3944 | 0.6933 | 0.8484 | Decreasing |
| KIRP-US::SP106638 | Kidney | KIRP-US | 1546 | 0 | 260 | 95 | 9 | 0.066 | 0.096 | 0.072 | 0.088 | -0.3914 | 0.6955 | 0.8503 | Decreasing |
| LIRI-JP::SP107123 | Liver | LIRI-JP | 606 | 0 | 94 | 40 | 13 | 0.035 | 0.033 | 0.038 | 0.023 | 0.3901 | 0.6964 | 0.8507 | Decreasing |
| KIRC-US::SP35617 | Kidney | KIRC-US | 675 | 0 | 97 | 40 | 9.5 | 0.033 | 0.040 | 0.031 | 0.029 | 0.3884 | 0.6977 | 0.8509 | Decreasing |
| UCEC-US::SP89687 | Endometrium | UCEC-US | 507 | 0 | 77 | 25 | 14 | 0.022 | 0.018 | 0.020 | 0.029 | -0.3885 | 0.6976 | 0.8509 | Increasing |
| LIRI-JP::SP98900 | Liver | LIRI-JP | 564 | 0 | 79 | 28 | 13 | 0.029 | 0.015 | 0.035 | 0.012 | 0.3827 | 0.7020 | 0.8511 | Increasing |
| LUSC-US::SP56771 | Lung | LUSC-US | 792 | 0 | 156 | 33 | 11 | 0.022 | 0.032 | 0.029 | 0.028 | -0.3868 | 0.6989 | 0.8511 | Increasing |
| BRCA-US::SP8891 | Breast | BRCA-US | 668 | 339 | 219 | 120 | 15 | 0.127 | 0.082 | 0.094 | 0.117 | 0.3866 | 0.6991 | 0.8511 | Increasing |
| PRAD-CA::SP112873 | Prostate | PRAD-US | 294 | 0 | 0 | 11 | 7 | 0.011 | 0.008 | 0.012 | 0.005 | 0.3827 | 0.7019 | 0.8511 | Decreasing |
| KIRC-US::SP39102 | Kidney | KIRC-US | 453 | 0 | 56 | 20 | 7 | 0.022 | 0.013 | 0.019 | 0.015 | 0.3836 | 0.7013 | 0.8511 | Decreasing |
| LIRI-JP::SP107019 | Liver | LIRI-JP | 666 | 0 | 77 | 24 | 12 | 0.022 | 0.023 | 0.013 | 0.023 | 0.3832 | 0.7016 | 0.8511 | Decreasing |
| LIHC-US::SP49229 | Liver | LIHC-US | 921 | 0 | 0 | 19 | 9 | 0.013 | 0.018 | 0.016 | 0.018 | -0.3837 | 0.7012 | 0.8511 | Increasing |

| | | | | | | | | | | | | | | | |
|---|---|---|---|---|---|---|---|---|---|---|---|---|---|---|---|
| STAD-US::SP84392 | Stomach | STAD-US | 103366 | 0 | 0 | 56 | 8 | 0.046 | 0.036 | 0.069 | 0.037 | -0.3814 | 0.7029 | 0.8515 | Increasing |
| COAD-US::SP17294 | CRC | COAD-US | 2365 | 0 | 0 | 16 | 11 | 0.012 | 0.016 | 0.007 | 0.023 | -0.3773 | 0.7059 | 0.8544 | Increasing |
| LIRI-JP::SP107017 | Liver | LIRI-JP | 434 | 0 | 72 | 31 | 12 | 0.022 | 0.028 | 0.025 | 0.029 | -0.3737 | 0.7086 | 0.8570 | Increasing |
| BLCA-US::SP1003 | Bladder | BLCA-US | 650 | 0 | 82 | 27 | 11 | 0.028 | 0.016 | 0.020 | 0.036 | -0.3684 | 0.7125 | 0.8603 | Increasing |
| UCEC-US::SP92947 | Endometrium | UCEC-US | 527 | 0 | 0 | 16 | 10.5 | 0.019 | 0.008 | 0.017 | 0.012 | 0.3687 | 0.7123 | 0.8603 | Increasing |
| UCEC-US::SP90125 | Endometrium | UCEC-US | 457 | 0 | 75 | 13 | 10 | 0.016 | 0.008 | 0.000 | 0.029 | -0.3676 | 0.7132 | 0.8603 | Increasing |
| LUAD-US::SP54113 | Lung | LUAD-US | 3876 | 0 | 0 | 78 | 12 | 0.060 | 0.073 | 0.059 | 0.076 | -0.3625 | 0.7170 | 0.8610 | Increasing |
| PRAD-UK::SP114944 | Prostate | PRAD-US | 1222 | 0 | 321 | 101 | 10 | 0.093 | 0.079 | 0.073 | 0.110 | -0.3532 | 0.7239 | 0.8610 | Increasing |
| PRAD-UK::SP114982 | Prostate | PRAD-US | 722 | 0 | 0 | 26 | 10 | 0.015 | 0.029 | 0.018 | 0.025 | -0.3588 | 0.7198 | 0.8610 | Decreasing |
| PRAD-UK::SP114980 | Prostate | PRAD-US | 809 | 0 | 0 | 26 | 10 | 0.019 | 0.024 | 0.021 | 0.025 | -0.3588 | 0.7198 | 0.8610 | Increasing |
| THCA-US::SP85511 | Thyroid | THCA-US | 157 | 0 | 39 | 12 | 10 | 0.008 | 0.013 | 0.006 | 0.015 | -0.3535 | 0.7237 | 0.8610 | Increasing |
| HNSC-US::SP30907 | Head&Neck | HNSC-US | 745 | 0 | 115 | 49 | 14 | 0.048 | 0.025 | 0.052 | 0.046 | -0.3573 | 0.7209 | 0.8610 | Increasing |
| PACA-CA::SP125719 | Pancreas | PACA-CA | 322 | 0 | 79 | 12 | 10 | 0.004 | 0.011 | 0.014 | 0.006 | -0.3600 | 0.7188 | 0.8610 | Increasing |
| HNSC-US::SP31790 | Head&Neck | HNSC-US | 483 | 0 | 84 | 25 | 14.5 | 0.030 | 0.008 | 0.029 | 0.017 | 0.3604 | 0.7185 | 0.8610 | Increasing |
| PACA-CA::SP125781 | Pancreas | PACA-CA | 396 | 0 | 0 | 17 | 12 | 0.015 | 0.008 | 0.014 | 0.016 | -0.3529 | 0.7242 | 0.8610 | Increasing |
| GBM-US::SP28659 | Brain | GBM-US | 1198 | 0 | 327 | 98 | 9 | 0.086 | 0.092 | 0.065 | 0.092 | 0.3561 | 0.7218 | 0.8610 | Decreasing |
| MALY-DE::SP116642 | Blood | MALY-DE | 1239 | 176 | 0 | 30 | 11 | 0.006 | 0.031 | 0.020 | 0.024 | -0.3550 | 0.7226 | 0.8610 | Increasing |
| RECA-EU::SP103140 | Kidney | RECA-EU | 1254 | 0 | 150 | 66 | 9 | 0.043 | 0.057 | 0.054 | 0.050 | -0.3526 | 0.7244 | 0.8610 | Increasing |
| PACA-CA::SP116949 | Pancreas | PACA-CA | 713 | 0 | 166 | 22 | 10 | 0.015 | 0.020 | 0.011 | 0.022 | -0.3545 | 0.7229 | 0.8610 | Decreasing |
| PACA-CA::SP117040 | Pancreas | PACA-CA | 668 | 0 | 0 | 17 | 12 | 0.015 | 0.008 | 0.014 | 0.016 | -0.3529 | 0.7242 | 0.8610 | Increasing |
| KIRC-US::SP35951 | Kidney | KIRC-US | 827 | 0 | 113 | 38 | 9 | 0.018 | 0.048 | 0.025 | 0.034 | -0.3568 | 0.7212 | 0.8610 | Decreasing |
| PACA-CA::SP117412 | Pancreas | PACA-CA | 1080 | 341 | 230 | 108 | 15 | 0.106 | 0.070 | 0.073 | 0.092 | 0.3596 | 0.7192 | 0.8610 | Decreasing |
| RECA-EU::SP103694 | Kidney | RECA-EU | 1556 | 0 | 251 | 120 | 9 | 0.096 | 0.087 | 0.094 | 0.105 | -0.3601 | 0.7188 | 0.8610 | Increasing |
| BRCA-US::SP3631 | Breast | BRCA-US | 323 | 0 | 69 | 16 | 13.5 | 0.010 | 0.018 | 0.006 | 0.021 | -0.3654 | 0.7148 | 0.8610 | Decreasing |
| OV-US::SP66960 | Ovary | OV-US | 374 | 0 | 62 | 15 | 14 | 0.020 | 0.008 | 0.010 | 0.017 | 0.3510 | 0.7256 | 0.8610 | Increasing |
| LIRI-JP::SP50139 | Liver | LIRI-JP | 272 | 20 | 43 | 19 | 14.5 | 0.019 | 0.010 | 0.016 | 0.023 | -0.3513 | 0.7254 | 0.8610 | Increasing |
| CESC-US::SP109957 | Cervix | CESC-US | 552 | 0 | 0 | 27 | 11 | 0.023 | 0.019 | 0.027 | 0.025 | -0.3491 | 0.7270 | 0.8618 | Increasing |
| LIRI-JP::SP99057 | Liver | LIRI-JP | 129 | 19 | 17 | 13 | 12 | 0.010 | 0.013 | 0.006 | 0.017 | -0.3485 | 0.7275 | 0.8618 | Increasing |
| KIRP-US::SP97104 | Kidney | KIRP-US | 507 | 187 | 104 | 75 | 11.5 | 0.080 | 0.048 | 0.047 | 0.098 | -0.3477 | 0.7280 | 0.8618 | Increasing |
| LUSC-US::SP58342 | Lung | LUSC-US | 2560 | 0 | 0 | 43 | 12 | 0.032 | 0.042 | 0.029 | 0.045 | -0.3464 | 0.7290 | 0.8622 | Increasing |
| MELA-AU::SP124449 | Skin | SKCM-US | 789 | 0 | 98 | 28 | 11 | 0.031 | 0.013 | 0.025 | 0.033 | -0.3431 | 0.7315 | 0.8645 | Increasing |
| BRCA-US::SP12049 | Breast | BRCA-US | 1395 | 677 | 718 | 320 | 19 | 0.276 | 0.282 | 0.258 | 0.270 | 0.3390 | 0.7346 | 0.8666 | Decreasing |
| OV-US::SP63546 | Ovary | OV-US | 428 | 61 | 0 | 25 | 11 | 0.016 | 0.025 | 0.020 | 0.023 | -0.3390 | 0.7346 | 0.8666 | Increasing |
| GACA-CN::SP135299 | Stomach | STAD-US | 825 | 0 | 0 | 21 | 13 | 0.015 | 0.020 | 0.016 | 0.021 | -0.3383 | 0.7351 | 0.8666 | Increasing |
| PRAD-UK::SP114941 | Prostate | PRAD-US | 1222 | 0 | 328 | 99 | 9 | 0.093 | 0.076 | 0.070 | 0.110 | -0.3350 | 0.7376 | 0.8667 | Increasing |
| HNSC-US::SP31126 | Head&Neck | HNSC-US | 1560 | 0 | 435 | 131 | 9 | 0.106 | 0.123 | 0.114 | 0.092 | 0.3319 | 0.7399 | 0.8667 | Decreasing |
| KIRC-US::SP39907 | Kidney | KIRC-US | 489 | 0 | 56 | 24 | 10.5 | 0.015 | 0.030 | 0.009 | 0.029 | -0.3359 | 0.7369 | 0.8667 | Decreasing |
| LUAD-US::SP53387 | Lung | LUAD-US | 1304 | 0 | 0 | 42 | 11 | 0.019 | 0.054 | 0.036 | 0.027 | -0.3351 | 0.7376 | 0.8667 | Decreasing |
| LIRI-JP::SP99181 | Liver | LIRI-JP | 407 | 0 | 47 | 23 | 14 | 0.022 | 0.020 | 0.013 | 0.023 | 0.3320 | 0.7399 | 0.8667 | Decreasing |
| RECA-EU::SP103730 | Kidney | RECA-EU | 716 | 0 | 150 | 61 | 9.5 | 0.036 | 0.061 | 0.037 | 0.055 | -0.3351 | 0.7375 | 0.8667 | Decreasing |
| PACA-AU::SP108068 | Pancreas | PACA-AU | 540 | 0 | 119 | 20 | 14 | 0.007 | 0.024 | 0.012 | 0.017 | -0.3325 | 0.7395 | 0.8667 | Decreasing |
| LIRI-JP::SP99229 | Liver | LIRI-JP | 524 | 0 | 77 | 23 | 15 | 0.019 | 0.025 | 0.010 | 0.023 | 0.3320 | 0.7399 | 0.8667 | Decreasing |
| LUSC-US::SP57669 | Lung | LUSC-US | 1952 | 0 | 0 | 31 | 13 | 0.019 | 0.034 | 0.023 | 0.028 | -0.3269 | 0.7437 | 0.8704 | Decreasing |
| READ-US::SP81711 | CRC | READ-US | 2483 | 0 | 0 | 37 | 11 | 0.031 | 0.035 | 0.034 | 0.023 | 0.3224 | 0.7471 | 0.8734 | Decreasing |
| BRCA-US::SP8394 | Breast | BRCA-US | 300 | 0 | 54 | 12 | 8 | 0.010 | 0.013 | 0.006 | 0.011 | 0.3204 | 0.7487 | 0.8734 | Decreasing |
| BRCA-UK::SP2158 | Breast | BRCA-US | 249 | 0 | 54 | 12 | 12 | 0.014 | 0.008 | 0.009 | 0.011 | 0.3204 | 0.7487 | 0.8734 | Decreasing |
| BRCA-UK::SP116345 | Breast | BRCA-US | 314 | 0 | 78 | 12 | 8 | 0.017 | 0.005 | 0.006 | 0.016 | 0.3204 | 0.7487 | 0.8734 | Decreasing |
| OV-AU::SP101536 | Ovary | OV-AU | 250 | 0 | 39 | 15 | 11 | 0.013 | 0.009 | 0.012 | 0.016 | -0.3192 | 0.7496 | 0.8737 | Increasing |
| PRAD-UK::SP114984 | Prostate | PRAD-US | 656 | 0 | 0 | 19 | 10 | 0.015 | 0.018 | 0.018 | 0.010 | 0.3183 | 0.7502 | 0.8738 | Decreasing |
| SKCM-US::SP82461 | Skin | SKCM-US | 750 | 0 | 238 | 79 | 14 | 0.075 | 0.062 | 0.066 | 0.067 | 0.3173 | 0.7510 | 0.8739 | Decreasing |

| | | | | | | | | | | | | | | | |
|---|---|---|---|---|---|---|---|---|---|---|---|---|---|---|---|
| READ-US::SP80657 | CRC | READ-US | 690 | 0 | 0 | 14 | 9.5 | 0.006 | 0.013 | 0.023 | 0.000 | -0.3165 | 0.7516 | 0.8740 | Increasing |
| OV-AU::SP101724 | Ovary | OV-AU | 670 | 119 | 106 | 58 | 12 | 0.051 | 0.045 | 0.036 | 0.052 | 0.3140 | 0.7535 | 0.8748 | Decreasing |
| BRCA-US::SP8987 | Breast | BRCA-US | 1122 | 633 | 305 | 217 | 12 | 0.214 | 0.150 | 0.201 | 0.180 | 0.3141 | 0.7534 | 0.8748 | Increasing |
| LIHC-US::SP49481 | Liver | LIHC-US | 182 | 0 | 53 | 20 | 13 | 0.016 | 0.013 | 0.026 | 0.012 | -0.3122 | 0.7549 | 0.8757 | Increasing |
| PRAD-CA::SP112965 | Prostate | PRAD-US | 276 | 0 | 41 | 16 | 12 | 0.011 | 0.018 | 0.012 | 0.010 | 0.3051 | 0.7603 | 0.8785 | Decreasing |
| KIRC-US::SP34493 | Kidney | KIRC-US | 350 | 0 | 54 | 19 | 9.5 | 0.018 | 0.013 | 0.022 | 0.010 | 0.3068 | 0.7590 | 0.8785 | Decreasing |
| RECA-EU::SP103679 | Kidney | RECA-EU | 409 | 41 | 0 | 18 | 8 | 0.007 | 0.026 | 0.009 | 0.010 | 0.3054 | 0.7601 | 0.8785 | Decreasing |
| PRAD-CA::SP112887 | Prostate | PRAD-US | 249 | 0 | 43 | 16 | 10 | 0.015 | 0.013 | 0.015 | 0.010 | 0.3051 | 0.7603 | 0.8785 | Decreasing |
| KIRC-US::SP40047 | Kidney | KIRC-US | 1016 | 0 | 181 | 73 | 9 | 0.062 | 0.064 | 0.049 | 0.079 | -0.3054 | 0.7601 | 0.8785 | Increasing |
| BRCA-UK::SP116333 | Breast | BRCA-US | 365 | 112 | 0 | 28 | 9 | 0.028 | 0.024 | 0.019 | 0.026 | 0.3041 | 0.7611 | 0.8786 | Decreasing |
| LIHC-US::SP115501 | Liver | LIHC-US | 605 | 0 | 0 | 17 | 16 | 0.013 | 0.016 | 0.013 | 0.018 | -0.2998 | 0.7643 | 0.8817 | Increasing |
| PRAD-UK::SP114986 | Prostate | PRAD-US | 734 | 0 | 0 | 22 | 10 | 0.015 | 0.021 | 0.018 | 0.020 | -0.2978 | 0.7659 | 0.8821 | Increasing |
| BRCA-US::SP5844 | Breast | BRCA-US | 588 | 0 | 111 | 22 | 7 | 0.007 | 0.029 | 0.022 | 0.011 | -0.2978 | 0.7659 | 0.8821 | Decreasing |
| PRAD-UK::SP114914 | Prostate | PRAD-US | 760 | 0 | 0 | 15 | 12 | 0.011 | 0.011 | 0.018 | 0.010 | -0.2920 | 0.7702 | 0.8837 | Increasing |
| PRAD-UK::SP114951 | Prostate | PRAD-US | 624 | 0 | 0 | 13 | 11 | 0.015 | 0.008 | 0.012 | 0.010 | 0.2920 | 0.7703 | 0.8837 | Decreasing |
| BRCA-US::SP3368 | Breast | BRCA-US | 279 | 45 | 44 | 15 | 10.5 | 0.014 | 0.016 | 0.006 | 0.016 | 0.2949 | 0.7681 | 0.8837 | Decreasing |
| PRAD-CA::SP102626 | Prostate | PRAD-US | 364 | 0 | 47 | 15 | 9.5 | 0.004 | 0.018 | 0.018 | 0.005 | -0.2920 | 0.7702 | 0.8837 | Increasing |
| OV-US::SP62450 | Ovary | OV-US | 194 | 31 | 36 | 18 | 15 | 0.013 | 0.023 | 0.007 | 0.017 | 0.2911 | 0.7710 | 0.8837 | Decreasing |
| UCEC-US::SP92659 | Endometrium | UCEC-US | 3775 | 0 | 0 | 17 | 10.5 | 0.010 | 0.016 | 0.023 | 0.006 | -0.2917 | 0.7705 | 0.8837 | Increasing |
| PACA-CA::SP116965 | Pancreas | PACA-CA | 2352 | 1519 | 0 | 339 | 13 | 0.256 | 0.260 | 0.265 | 0.267 | -0.2884 | 0.7731 | 0.8844 | Increasing |
| PACA-AU::SP70756 | Pancreas | PACA-AU | 465 | 0 | 0 | 15 | 9.5 | 0.011 | 0.011 | 0.012 | 0.013 | -0.2880 | 0.7734 | 0.8844 | Increasing |
| PACA-AU::SP110770 | Pancreas | PACA-AU | 626 | 0 | 0 | 15 | 12 | 0.007 | 0.011 | 0.020 | 0.007 | -0.2880 | 0.7734 | 0.8844 | Increasing |
| COAD-US::SP21193 | CRC | COAD-US | 1066 | 0 | 0 | 19 | 10.5 | 0.015 | 0.016 | 0.023 | 0.006 | 0.2855 | 0.7753 | 0.8859 | Decreasing |
| LIRI-JP::SP98913 | Liver | LIRI-JP | 714 | 0 | 120 | 52 | 14 | 0.048 | 0.035 | 0.045 | 0.052 | -0.2843 | 0.7762 | 0.8862 | Increasing |
| MELA-AU::SP124311 | Skin | SKCM-US | 1018 | 0 | 261 | 78 | 13 | 0.065 | 0.075 | 0.057 | 0.067 | 0.2827 | 0.7774 | 0.8869 | Decreasing |
| KIRC-US::SP37903 | Kidney | KIRC-US | 1186 | 0 | 190 | 68 | 10 | 0.058 | 0.059 | 0.062 | 0.049 | 0.2818 | 0.7781 | 0.8870 | Decreasing |
| PRAD-UK::SP114976 | Prostate | PRAD-US | 731 | 0 | 0 | 27 | 10 | 0.015 | 0.032 | 0.018 | 0.025 | -0.2782 | 0.7808 | 0.8880 | Decreasing |
| SKCM-US::SP83482 | Skin | SKCM-US | 423 | 0 | 57 | 15 | 11 | 0.014 | 0.008 | 0.019 | 0.011 | -0.2784 | 0.7807 | 0.8880 | Increasing |
| LIRI-JP::SP112300 | Liver | LIRI-JP | 382 | 0 | 54 | 22 | 11 | 0.025 | 0.013 | 0.016 | 0.023 | 0.2788 | 0.7804 | 0.8880 | Increasing |
| PACA-CA::SP125725 | Pancreas | PACA-CA | 716 | 0 | 0 | 28 | 12 | 0.030 | 0.017 | 0.017 | 0.025 | 0.2747 | 0.7835 | 0.8887 | Decreasing |
| LUAD-US::SP51824 | Lung | LUAD-US | 910 | 0 | 86 | 28 | 12 | 0.035 | 0.014 | 0.010 | 0.049 | -0.2736 | 0.7844 | 0.8887 | Increasing |
| BRCA-US::SP4875 | Breast | BRCA-US | 1143 | 441 | 462 | 239 | 19 | 0.203 | 0.195 | 0.236 | 0.164 | 0.2724 | 0.7853 | 0.8887 | Increasing |
| STAD-US::SP84790 | Stomach | STAD-US | 1224 | 0 | 0 | 19 | 11 | 0.015 | 0.017 | 0.013 | 0.021 | -0.2694 | 0.7876 | 0.8887 | Increasing |
| UCEC-US::SP94933 | Endometrium | UCEC-US | 66330 | 0 | 0 | 38 | 8 | 0.022 | 0.041 | 0.046 | 0.006 | 0.2687 | 0.7882 | 0.8887 | Decreasing |
| MELA-AU::SP124436 | Skin | SKCM-US | 2288 | 0 | 404 | 134 | 12 | 0.106 | 0.122 | 0.107 | 0.122 | -0.2687 | 0.7882 | 0.8887 | Increasing |
| LUAD-US::SP53548 | Lung | LUAD-US | 261 | 0 | 38 | 16 | 10 | 0.013 | 0.014 | 0.013 | 0.016 | -0.2761 | 0.7825 | 0.8887 | Increasing |
| LIRI-JP::SP99033 | Liver | LIRI-JP | 668 | 0 | 57 | 18 | 14 | 0.019 | 0.013 | 0.013 | 0.017 | 0.2734 | 0.7845 | 0.8887 | Decreasing |
| LUSC-US::SP56569 | Lung | LUSC-US | 1218 | 0 | 0 | 25 | 10 | 0.029 | 0.013 | 0.023 | 0.023 | 0.2691 | 0.7878 | 0.8887 | Increasing |
| GACA-CN::SP135420 | Stomach | STAD-US | 784 | 0 | 0 | 17 | 12.5 | 0.015 | 0.011 | 0.023 | 0.005 | 0.2722 | 0.7855 | 0.8887 | Increasing |
| BRCA-US::SP4265 | Breast | BRCA-US | 274 | 61 | 116 | 43 | 15 | 0.045 | 0.032 | 0.031 | 0.042 | 0.2700 | 0.7871 | 0.8887 | Decreasing |
| BRCA-US::SP6429 | Breast | BRCA-US | 822 | 321 | 222 | 131 | 13 | 0.124 | 0.106 | 0.101 | 0.122 | 0.2662 | 0.7901 | 0.8902 | Decreasing |
| OV-AU::SP101515 | Ovary | OV-AU | 416 | 173 | 151 | 80 | 21 | 0.064 | 0.068 | 0.051 | 0.067 | 0.2651 | 0.7909 | 0.8904 | Decreasing |
| OV-US::SP68659 | Ovary | OV-US | 486 | 205 | 243 | 127 | 22 | 0.114 | 0.099 | 0.102 | 0.128 | -0.2615 | 0.7937 | 0.8928 | Increasing |
| MELA-AU::SP124384 | Skin | SKCM-US | 22934 | 0 | 0 | 96 | 15 | 0.092 | 0.067 | 0.079 | 0.100 | -0.2606 | 0.7944 | 0.8930 | Increasing |
| PACA-CA::SP117002 | Pancreas | PACA-CA | 769 | 0 | 0 | 24 | 11 | 0.023 | 0.017 | 0.017 | 0.019 | 0.2543 | 0.7992 | 0.8975 | Decreasing |
| PRAD-CA::SP112971 | Prostate | PRAD-US | 344 | 0 | 41 | 13 | 8 | 0.015 | 0.005 | 0.012 | 0.015 | -0.2537 | 0.7997 | 0.8975 | Increasing |
| PACA-CA::SP117463 | Pancreas | PACA-CA | 606 | 0 | 0 | 11 | 12 | 0.008 | 0.008 | 0.008 | 0.010 | -0.2507 | 0.8020 | 0.8994 | Increasing |
| COAD-US::SP18946 | CRC | COAD-US | 16986 | 0 | 0 | 21 | 7 | 0.015 | 0.019 | 0.020 | 0.017 | -0.2467 | 0.8051 | 0.9010 | Increasing |
| LUAD-US::SP54363 | Lung | LUAD-US | 1513 | 0 | 0 | 31 | 16 | 0.028 | 0.022 | 0.036 | 0.016 | 0.2471 | 0.8048 | 0.9010 | Increasing |
| UCEC-US::SP95222 | Endometrium | UCEC-US | 748 | 104 | 0 | 14 | 13 | 0.016 | 0.008 | 0.013 | 0.012 | 0.2462 | 0.8055 | 0.9010 | Increasing |

| | | | | | | | | | | | | | | | |
|---|---|---|---|---|---|---|---|---|---|---|---|---|---|---|---|
| BRCA-UK::SP116339 | Breast | BRCA-US | 400 | 0 | 75 | 12 | 13 | 0.010 | 0.008 | 0.013 | 0.011 | -0.2457 | 0.8059 | 0.9010 | Increasing |
| LUAD-US::SP50317 | Lung | LUAD-US | 531 | 0 | 52 | 20 | 8 | 0.009 | 0.027 | 0.020 | 0.005 | 0.2334 | 0.8154 | 0.9017 | Decreasing |
| MELA-AU::SP124428 | Skin | SKCM-US | 1741 | 0 | 355 | 140 | 16 | 0.137 | 0.111 | 0.098 | 0.145 | 0.2353 | 0.8140 | 0.9017 | Decreasing |
| LUSC-US::SP57941 | Lung | LUSC-US | 2833 | 0 | 0 | 42 | 9 | 0.026 | 0.042 | 0.046 | 0.023 | -0.2439 | 0.8073 | 0.9017 | Increasing |
| BRCA-US::SP9251 | Breast | BRCA-US | 233 | 43 | 45 | 11 | 15 | 0.007 | 0.013 | 0.009 | 0.005 | 0.2328 | 0.8159 | 0.9017 | Decreasing |
| PACA-CA::SP117201 | Pancreas | PACA-CA | 741 | 0 | 0 | 23 | 13 | 0.015 | 0.020 | 0.017 | 0.019 | -0.2384 | 0.8115 | 0.9017 | Increasing |
| UCEC-US::SP95646 | Endometrium | UCEC-US | 507 | 70 | 0 | 21 | 14 | 0.010 | 0.023 | 0.027 | 0.006 | -0.2357 | 0.8136 | 0.9017 | Decreasing |
| BTCA-SG::SP117627 | Biliary | BTCA-SG | 457 | 0 | 78 | 19 | 10 | 0.025 | 0.003 | 0.032 | 0.011 | 0.2363 | 0.8132 | 0.9017 | Increasing |
| PACA-CA::SP125804 | Pancreas | PACA-CA | 1536 | 566 | 626 | 314 | 20 | 0.230 | 0.260 | 0.225 | 0.255 | -0.2409 | 0.8096 | 0.9017 | Decreasing |
| PACA-AU::SP107952 | Pancreas | PACA-AU | 638 | 0 | 123 | 19 | 11.5 | 0.007 | 0.022 | 0.015 | 0.013 | -0.2382 | 0.8117 | 0.9017 | Decreasing |
| PACA-AU::SP70106 | Pancreas | PACA-AU | 670 | 300 | 160 | 82 | 13 | 0.059 | 0.068 | 0.061 | 0.067 | -0.2394 | 0.8108 | 0.9017 | Increasing |
| RECA-EU::SP103261 | Kidney | RECA-EU | 1602 | 0 | 240 | 110 | 9 | 0.079 | 0.092 | 0.082 | 0.090 | -0.2342 | 0.8148 | 0.9017 | Increasing |
| LIRI-JP::SP99093 | Liver | LIRI-JP | 626 | 0 | 61 | 27 | 13 | 0.022 | 0.023 | 0.019 | 0.029 | -0.2379 | 0.8120 | 0.9017 | Increasing |
| LIRI-JP::SP50157 | Liver | LIRI-JP | 517 | 0 | 102 | 40 | 10 | 0.025 | 0.045 | 0.022 | 0.041 | -0.2373 | 0.8124 | 0.9017 | Decreasing |
| BRCA-UK::SP2156 | Breast | BRCA-US | 92 | 0 | 27 | 11 | 14 | 0.010 | 0.011 | 0.006 | 0.011 | 0.2328 | 0.8159 | 0.9017 | Decreasing |
| PRAD-UK::SP106825 | Prostate | PRAD-US | 549 | 0 | 0 | 18 | 11 | 0.019 | 0.008 | 0.027 | 0.005 | 0.2366 | 0.8130 | 0.9017 | Increasing |
| PRAD-UK::SP114974 | Prostate | PRAD-US | 733 | 0 | 0 | 18 | 9 | 0.011 | 0.018 | 0.015 | 0.015 | -0.2272 | 0.8203 | 0.9059 | Increasing |
| COAD-US::SP23078 | CRC | COAD-US | 1078 | 0 | 0 | 28 | 12 | 0.027 | 0.022 | 0.013 | 0.040 | -0.2237 | 0.8230 | 0.9073 | Increasing |
| LIHC-US::SP98096 | Liver | LIHC-US | 743 | 0 | 99 | 41 | 11.5 | 0.032 | 0.047 | 0.019 | 0.042 | 0.2246 | 0.8223 | 0.9073 | Decreasing |
| LIRI-JP::SP99105 | Liver | LIRI-JP | 329 | 0 | 51 | 21 | 11 | 0.016 | 0.018 | 0.025 | 0.006 | 0.2232 | 0.8234 | 0.9073 | Decreasing |
| BLCA-US::SP1365 | Bladder | BLCA-US | 767 | 0 | 181 | 48 | 9 | 0.031 | 0.063 | 0.023 | 0.042 | 0.2197 | 0.8261 | 0.9095 | Decreasing |
| MELA-AU::SP124425 | Skin | SKCM-US | 415 | 0 | 109 | 41 | 12 | 0.048 | 0.026 | 0.025 | 0.050 | 0.2188 | 0.8268 | 0.9096 | Increasing |
| LUAD-US::SP50321 | Lung | LUAD-US | 599 | 0 | 87 | 29 | 13 | 0.013 | 0.038 | 0.026 | 0.016 | -0.2141 | 0.8305 | 0.9122 | Decreasing |
| LUSC-US::SP56644 | Lung | LUSC-US | 2153 | 0 | 0 | 62 | 14 | 0.064 | 0.037 | 0.052 | 0.068 | -0.2134 | 0.8310 | 0.9122 | Increasing |
| BRCA-UK::SP116355 | Breast | BRCA-US | 373 | 118 | 172 | 87 | 16 | 0.065 | 0.084 | 0.066 | 0.079 | -0.2148 | 0.8299 | 0.9122 | Decreasing |
| PRAD-CA::SP112825 | Prostate | PRAD-US | 256 | 0 | 36 | 11 | 14 | 0.011 | 0.005 | 0.012 | 0.010 | -0.2105 | 0.8332 | 0.9125 | Increasing |
| PRAD-CA::SP112905 | Prostate | PRAD-US | 297 | 0 | 44 | 11 | 10 | 0.007 | 0.008 | 0.015 | 0.005 | -0.2105 | 0.8332 | 0.9125 | Increasing |
| PRAD-UK::SP106827 | Prostate | PRAD-US | 349 | 0 | 0 | 11 | 10.5 | 0.007 | 0.011 | 0.009 | 0.010 | -0.2105 | 0.8332 | 0.9125 | Increasing |
| LIRI-JP::SP50183 | Liver | LIRI-JP | 715 | 0 | 112 | 32 | 10 | 0.035 | 0.018 | 0.029 | 0.029 | 0.2087 | 0.8347 | 0.9135 | Increasing |
| PACA-CA::SP125694 | Pancreas | PACA-CA | 533 | 0 | 151 | 16 | 14 | 0.019 | 0.008 | 0.008 | 0.016 | 0.2077 | 0.8355 | 0.9136 | Decreasing |
| PACA-AU::SP75202 | Pancreas | PACA-AU | 375 | 0 | 130 | 12 | 13 | 0.011 | 0.008 | 0.006 | 0.013 | -0.2035 | 0.8387 | 0.9136 | Increasing |
| CESC-US::SP13078 | Cervix | CESC-US | 429 | 76 | 0 | 17 | 13 | 0.014 | 0.017 | 0.013 | 0.012 | 0.2051 | 0.8375 | 0.9136 | Decreasing |
| PACA-AU::SP70122 | Pancreas | PACA-AU | 631 | 0 | 0 | 12 | 13.5 | 0.011 | 0.008 | 0.006 | 0.013 | -0.2035 | 0.8387 | 0.9136 | Increasing |
| OV-US::SP64976 | Ovary | OV-US | 377 | 77 | 0 | 24 | 12 | 0.013 | 0.030 | 0.013 | 0.023 | -0.2028 | 0.8393 | 0.9136 | Decreasing |
| PACA-AU::SP75681 | Pancreas | PACA-AU | 481 | 0 | 143 | 11 | 8.5 | 0.007 | 0.011 | 0.009 | 0.007 | 0.2047 | 0.8378 | 0.9136 | Decreasing |
| BRCA-US::SP96163 | Breast | BRCA-US | 747 | 363 | 345 | 175 | 17 | 0.158 | 0.121 | 0.179 | 0.138 | -0.2032 | 0.8390 | 0.9136 | Increasing |
| MELA-AU::SP124434 | Skin | SKCM-US | 648 | 0 | 105 | 47 | 15 | 0.038 | 0.044 | 0.041 | 0.033 | 0.1991 | 0.8422 | 0.9161 | Decreasing |
| MELA-AU::SP124406 | Skin | SKCM-US | 736 | 0 | 109 | 39 | 12 | 0.038 | 0.031 | 0.022 | 0.050 | -0.1957 | 0.8448 | 0.9183 | Increasing |
| MALY-DE::SP116649 | Blood | MALY-DE | 1023 | 0 | 0 | 11 | 10 | 0.000 | 0.013 | 0.006 | 0.009 | -0.1948 | 0.8456 | 0.9184 | Increasing |
| HNSC-US::SP30011 | Head&Neck | HNSC-US | 710 | 71 | 207 | 87 | 11 | 0.085 | 0.057 | 0.085 | 0.069 | 0.1925 | 0.8474 | 0.9188 | Increasing |
| PAEN-AU::SP118043 | Pancreas | PAEN-AU | 196 | 0 | 51 | 11 | 16 | 0.010 | 0.005 | 0.011 | 0.009 | -0.1919 | 0.8478 | 0.9188 | Increasing |
| LIRI-JP::SP99329 | Liver | LIRI-JP | 521 | 304 | 100 | 126 | 13 | 0.124 | 0.090 | 0.092 | 0.127 | 0.1931 | 0.8469 | 0.9188 | Increasing |
| THCA-US::SP88098 | Thyroid | THCA-US | 122 | 0 | 46 | 11 | 8 | 0.004 | 0.013 | 0.013 | 0.005 | -0.1898 | 0.8494 | 0.9199 | Increasing |
| LGG-US::SP48850 | Brain | LGG-US | 377 | 0 | 0 | 14 | 13.5 | 0.006 | 0.015 | 0.017 | 0.004 | 0.1874 | 0.8513 | 0.9212 | Increasing |
| KIRP-US::SP97258 | Kidney | KIRP-US | 964 | 0 | 176 | 60 | 9 | 0.049 | 0.040 | 0.078 | 0.031 | -0.1850 | 0.8533 | 0.9226 | Increasing |
| MELA-AU::SP124298 | Skin | SKCM-US | 2119 | 0 | 584 | 188 | 13 | 0.171 | 0.161 | 0.135 | 0.183 | 0.1820 | 0.8556 | 0.9244 | Decreasing |
| SKCM-US::SP82431 | Skin | SKCM-US | 2068 | 0 | 525 | 179 | 13 | 0.164 | 0.132 | 0.161 | 0.161 | -0.1788 | 0.8581 | 0.9264 | Increasing |
| KIRP-US::SP43514 | Kidney | KIRP-US | 651 | 0 | 120 | 44 | 9 | 0.024 | 0.051 | 0.037 | 0.031 | -0.1780 | 0.8587 | 0.9264 | Decreasing |
| BRCA-US::SP11171 | Breast | BRCA-US | 532 | 181 | 235 | 116 | 17 | 0.114 | 0.074 | 0.123 | 0.085 | 0.1768 | 0.8597 | 0.9268 | Increasing |
| LUAD-US::SP52779 | Lung | LUAD-US | 1613 | 0 | 0 | 33 | 10.5 | 0.038 | 0.016 | 0.029 | 0.033 | 0.1733 | 0.8625 | 0.9291 | Increasing |

| | | | | | | | | | | | | | | | |
|---|---|---|---|---|---|---|---|---|---|---|---|---|---|---|---|
| UCEC-US::SP91730 | Endometrium | UCEC-US | 998 | 411 | 194 | 156 | 15 | 0.124 | 0.145 | 0.123 | 0.139 | -0.1694 | 0.8654 | 0.9310 | Decreasing |
| PAEN-AU::SP102541 | Pancreas | PAEN-AU | 175 | 0 | 35 | 16 | 9 | 0.010 | 0.013 | 0.016 | 0.009 | 0.1681 | 0.8665 | 0.9310 | Increasing |
| KIRC-US::SP34246 | Kidney | KIRC-US | 765 | 0 | 120 | 36 | 8 | 0.025 | 0.032 | 0.037 | 0.025 | -0.1677 | 0.8668 | 0.9310 | Increasing |
| BRCA-UK::SP2146 | Breast | BRCA-US | 316 | 144 | 113 | 62 | 16 | 0.055 | 0.053 | 0.050 | 0.053 | 0.1679 | 0.8667 | 0.9310 | Decreasing |
| ORCA-IN::SP117124 | Head&Neck | ORCA-IN | 580 | 0 | 71 | 23 | 11 | 0.023 | 0.016 | 0.019 | 0.020 | 0.1668 | 0.8675 | 0.9311 | Increasing |
| RECA-EU::SP103603 | Kidney | RECA-EU | 705 | 0 | 141 | 49 | 8.5 | 0.033 | 0.042 | 0.045 | 0.025 | 0.1648 | 0.8691 | 0.9321 | Decreasing |
| PACA-CA::SP117690 | Pancreas | PACA-CA | 388 | 0 | 0 | 15 | 13 | 0.011 | 0.011 | 0.011 | 0.013 | -0.1611 | 0.8720 | 0.9346 | Increasing |
| LUSC-US::SP58349 | Lung | LUSC-US | 1840 | 0 | 344 | 97 | 13 | 0.077 | 0.085 | 0.101 | 0.057 | 0.1600 | 0.8729 | 0.9348 | Decreasing |
| CESC-US::SP107607 | Cervix | CESC-US | 320 | 74 | 0 | 16 | 18 | 0.020 | 0.003 | 0.020 | 0.012 | 0.1560 | 0.8761 | 0.9354 | Increasing |
| PACA-AU::SP71536 | Pancreas | PACA-AU | 374 | 114 | 0 | 16 | 10 | 0.015 | 0.011 | 0.009 | 0.017 | -0.1571 | 0.8752 | 0.9354 | Decreasing |
| PACA-CA::SP117648 | Pancreas | PACA-CA | 300 | 0 | 95 | 22 | 11 | 0.011 | 0.020 | 0.023 | 0.013 | -0.1552 | 0.8767 | 0.9354 | Increasing |
| RECA-EU::SP103021 | Kidney | RECA-EU | 684 | 0 | 111 | 37 | 9 | 0.033 | 0.021 | 0.040 | 0.020 | 0.1566 | 0.8756 | 0.9354 | Increasing |
| PACA-CA::SP117650 | Pancreas | PACA-CA | 269 | 0 | 72 | 22 | 9 | 0.011 | 0.025 | 0.011 | 0.019 | -0.1552 | 0.8767 | 0.9354 | Decreasing |
| MELA-AU::SP124362 | Skin | SKCM-US | 1029 | 0 | 120 | 49 | 11.5 | 0.034 | 0.052 | 0.041 | 0.033 | 0.1463 | 0.8837 | 0.9367 | Decreasing |
| PRAD-UK::SP114978 | Prostate | PRAD-US | 689 | 0 | 0 | 25 | 10 | 0.019 | 0.026 | 0.018 | 0.020 | 0.1477 | 0.8826 | 0.9367 | Decreasing |
| PRAD-CA::SP112843 | Prostate | PRAD-US | 395 | 0 | 38 | 17 | 12 | 0.022 | 0.008 | 0.012 | 0.020 | 0.1504 | 0.8804 | 0.9367 | Increasing |
| OV-AU::SP101921 | Ovary | OV-AU | 361 | 0 | 45 | 14 | 12.5 | 0.006 | 0.014 | 0.015 | 0.005 | -0.1486 | 0.8818 | 0.9367 | Increasing |
| RECA-EU::SP103080 | Kidney | RECA-EU | 1139 | 0 | 174 | 65 | 10 | 0.040 | 0.057 | 0.062 | 0.035 | -0.1521 | 0.8791 | 0.9367 | Increasing |
| PRAD-CA::SP102630 | Prostate | PRAD-US | 451 | 0 | 62 | 25 | 11 | 0.015 | 0.026 | 0.027 | 0.010 | 0.1477 | 0.8826 | 0.9367 | Decreasing |
| OV-AU::SP102123 | Ovary | OV-AU | 338 | 0 | 65 | 26 | 11 | 0.013 | 0.032 | 0.009 | 0.026 | -0.1468 | 0.8833 | 0.9367 | Decreasing |
| HNSC-US::SP31854 | Head&Neck | HNSC-US | 1477 | 215 | 0 | 59 | 14 | 0.061 | 0.041 | 0.042 | 0.063 | 0.1483 | 0.8821 | 0.9367 | Increasing |
| SKCM-US::SP103866 | Skin | SKCM-US | 673 | 0 | 98 | 30 | 11 | 0.034 | 0.021 | 0.016 | 0.039 | 0.1476 | 0.8827 | 0.9367 | Decreasing |
| RECA-EU::SP102957 | Kidney | RECA-EU | 1295 | 0 | 81 | 43 | 9 | 0.026 | 0.038 | 0.040 | 0.025 | -0.1423 | 0.8868 | 0.9390 | Increasing |
| PRAD-CA::SP112935 | Prostate | PRAD-US | 263 | 0 | 42 | 14 | 10.5 | 0.011 | 0.013 | 0.009 | 0.015 | -0.1419 | 0.8871 | 0.9390 | Decreasing |
| LUAD-US::SP55711 | Lung | LUAD-US | 427 | 0 | 78 | 15 | 9.5 | 0.009 | 0.016 | 0.016 | 0.005 | 0.1396 | 0.8890 | 0.9402 | Decreasing |
| GACA-CN::SP135436 | Stomach | STAD-US | 1315 | 0 | 0 | 24 | 12 | 0.015 | 0.028 | 0.016 | 0.021 | -0.1377 | 0.8905 | 0.9411 | Decreasing |
| SKCM-US::SP104530 | Skin | SKCM-US | 1290 | 0 | 183 | 52 | 13 | 0.051 | 0.036 | 0.041 | 0.056 | -0.1347 | 0.8929 | 0.9430 | Increasing |
| LUSC-US::SP56553 | Lung | LUSC-US | 2833 | 0 | 0 | 40 | 11 | 0.029 | 0.040 | 0.039 | 0.023 | 0.1235 | 0.9017 | 0.9462 | Decreasing |
| PACA-CA::SP117980 | Pancreas | PACA-CA | 516 | 0 | 142 | 17 | 9 | 0.011 | 0.017 | 0.008 | 0.016 | -0.1261 | 0.8996 | 0.9462 | Decreasing |
| RECA-EU::SP102913 | Kidney | RECA-EU | 776 | 0 | 141 | 54 | 9 | 0.043 | 0.045 | 0.037 | 0.045 | 0.1234 | 0.9018 | 0.9462 | Decreasing |
| PACA-CA::SP117214 | Pancreas | PACA-CA | 609 | 0 | 153 | 17 | 10 | 0.011 | 0.014 | 0.014 | 0.013 | -0.1261 | 0.8996 | 0.9462 | Increasing |
| LIRI-JP::SP99177 | Liver | LIRI-JP | 604 | 0 | 0 | 19 | 10 | 0.010 | 0.023 | 0.016 | 0.012 | -0.1237 | 0.9015 | 0.9462 | Decreasing |
| KIRP-US::SP97145 | Kidney | KIRP-US | 363 | 0 | 47 | 18 | 8 | 0.010 | 0.024 | 0.006 | 0.021 | -0.1243 | 0.9011 | 0.9462 | Decreasing |
| BRCA-US::SP10563 | Breast | BRCA-US | 302 | 0 | 77 | 16 | 15.5 | 0.017 | 0.013 | 0.006 | 0.021 | 0.1248 | 0.9007 | 0.9462 | Decreasing |
| BRCA-US::SP12186 | Breast | BRCA-US | 277 | 0 | 58 | 16 | 13 | 0.010 | 0.016 | 0.019 | 0.005 | 0.1248 | 0.9007 | 0.9462 | Decreasing |
| OV-US::SP60842 | Ovary | OV-US | 585 | 199 | 180 | 90 | 19 | 0.082 | 0.079 | 0.059 | 0.093 | 0.1291 | 0.8973 | 0.9462 | Decreasing |
| UCEC-US::SP94588 | Endometrium | UCEC-US | 538 | 0 | 88 | 27 | 13 | 0.022 | 0.028 | 0.013 | 0.029 | 0.1117 | 0.9110 | 0.9552 | Decreasing |
| BRCA-UK::SP2154 | Breast | BRCA-US | 1221 | 743 | 242 | 243 | 13 | 0.203 | 0.224 | 0.179 | 0.223 | 0.1052 | 0.9162 | 0.9599 | Decreasing |
| LINC-JP::SP98833 | Liver | LIHC-US | 357 | 42 | 46 | 27 | 14 | 0.022 | 0.026 | 0.019 | 0.024 | 0.1034 | 0.9177 | 0.9607 | Decreasing |
| LUSC-US::SP57024 | Lung | LUSC-US | 943 | 0 | 126 | 29 | 13 | 0.029 | 0.021 | 0.023 | 0.028 | 0.1006 | 0.9199 | 0.9610 | Increasing |
| LIRI-JP::SP106993 | Liver | LIRI-JP | 1021 | 0 | 106 | 37 | 14 | 0.032 | 0.033 | 0.025 | 0.035 | 0.1020 | 0.9187 | 0.9610 | Decreasing |
| PACA-CA::SP125783 | Pancreas | PACA-CA | 361 | 0 | 86 | 17 | 13 | 0.011 | 0.020 | 0.006 | 0.016 | 0.1007 | 0.9198 | 0.9610 | Decreasing |
| MALY-DE::SP116694 | Blood | MALY-DE | 943 | 0 | 0 | 14 | 13 | 0.013 | 0.013 | 0.003 | 0.015 | 0.0982 | 0.9218 | 0.9616 | Decreasing |
| SKCM-US::SP82429 | Skin | SKCM-US | 353 | 0 | 71 | 26 | 12.5 | 0.021 | 0.026 | 0.019 | 0.022 | 0.0986 | 0.9214 | 0.9616 | Decreasing |
| RECA-EU::SP102816 | Kidney | RECA-EU | 713 | 0 | 135 | 62 | 9 | 0.063 | 0.028 | 0.062 | 0.045 | 0.0902 | 0.9281 | 0.9620 | Increasing |
| MELA-AU::SP124396 | Skin | SKCM-US | 673 | 0 | 88 | 32 | 9 | 0.027 | 0.031 | 0.016 | 0.039 | -0.0922 | 0.9265 | 0.9620 | Decreasing |
| LIRI-JP::SP107152 | Liver | LIRI-JP | 680 | 0 | 70 | 26 | 11.5 | 0.022 | 0.020 | 0.025 | 0.017 | 0.0908 | 0.9276 | 0.9620 | Increasing |
| OV-AU::SP101564 | Ovary | OV-AU | 266 | 43 | 32 | 17 | 13.5 | 0.013 | 0.014 | 0.015 | 0.010 | 0.0950 | 0.9243 | 0.9620 | Decreasing |
| LUAD-US::SP53618 | Lung | LUAD-US | 2331 | 0 | 0 | 44 | 14 | 0.032 | 0.041 | 0.046 | 0.027 | -0.0923 | 0.9264 | 0.9620 | Increasing |
| PACA-CA::SP117522 | Pancreas | PACA-CA | 503 | 0 | 136 | 12 | 12 | 0.011 | 0.008 | 0.006 | 0.013 | -0.0901 | 0.9282 | 0.9620 | Decreasing |

| | | | | | | | | | | | | | | | |
|---|---|---|---|---|---|---|---|---|---|---|---|---|---|---|---|
| OV-AU::SP101576 | Ovary | OV-AU | 506 | 79 | 65 | 36 | 9 | 0.035 | 0.020 | 0.027 | 0.036 | -0.0961 | 0.9235 | 0.9620 | Increasing |
| LUSC-US::SP57066 | Lung | LUSC-US | 2333 | 0 | 0 | 52 | 11 | 0.029 | 0.063 | 0.042 | 0.034 | -0.0902 | 0.9281 | 0.9620 | Decreasing |
| PRAD-CA::SP112929 | Prostate | PRAD-US | 381 | 0 | 0 | 12 | 12.5 | 0.007 | 0.013 | 0.009 | 0.010 | -0.0908 | 0.9276 | 0.9620 | Decreasing |
| PACA-CA::SP125777 | Pancreas | PACA-CA | 1437 | 790 | 405 | 262 | 12 | 0.230 | 0.157 | 0.240 | 0.191 | 0.0894 | 0.9288 | 0.9620 | Increasing |
| LIRI-JP::SP112215 | Liver | LIRI-JP | 676 | 0 | 76 | 33 | 14 | 0.032 | 0.025 | 0.022 | 0.035 | 0.0824 | 0.9344 | 0.9652 | Increasing |
| HNSC-US::SP30493 | Head&Neck | HNSC-US | 187 | 0 | 32 | 12 | 11 | 0.012 | 0.005 | 0.016 | 0.006 | 0.0812 | 0.9353 | 0.9652 | Increasing |
| PACA-CA::SP116950 | Pancreas | PACA-CA | 888 | 0 | 0 | 15 | 10 | 0.008 | 0.017 | 0.011 | 0.010 | 0.0804 | 0.9360 | 0.9652 | Decreasing |
| LUSC-US::SP57651 | Lung | LUSC-US | 719 | 0 | 131 | 29 | 12 | 0.029 | 0.018 | 0.026 | 0.028 | -0.0814 | 0.9351 | 0.9652 | Increasing |
| KIRP-US::SP106602 | Kidney | KIRP-US | 736 | 0 | 110 | 49 | 9 | 0.045 | 0.037 | 0.044 | 0.041 | 0.0816 | 0.9349 | 0.9652 | Increasing |
| PACA-CA::SP125738 | Pancreas | PACA-CA | 284 | 0 | 74 | 15 | 11 | 0.015 | 0.008 | 0.011 | 0.013 | 0.0804 | 0.9360 | 0.9652 | Increasing |
| OV-AU::SP102103 | Ovary | OV-AU | 298 | 0 | 46 | 20 | 10 | 0.019 | 0.014 | 0.012 | 0.021 | 0.0769 | 0.9387 | 0.9661 | Decreasing |
| PACA-AU::SP107920 | Pancreas | PACA-AU | 528 | 0 | 0 | 16 | 11 | 0.015 | 0.011 | 0.012 | 0.013 | 0.0768 | 0.9388 | 0.9661 | Decreasing |
| KIRC-US::SP39298 | Kidney | KIRC-US | 1649 | 0 | 271 | 113 | 8.5 | 0.098 | 0.102 | 0.080 | 0.108 | 0.0773 | 0.9384 | 0.9661 | Decreasing |
| OV-US::SP61703 | Ovary | OV-US | 566 | 0 | 126 | 53 | 11 | 0.046 | 0.041 | 0.053 | 0.041 | -0.0747 | 0.9404 | 0.9671 | Increasing |
| LIRI-JP::SP99145 | Liver | LIRI-JP | 454 | 0 | 0 | 13 | 10 | 0.013 | 0.008 | 0.013 | 0.012 | -0.0734 | 0.9415 | 0.9675 | Increasing |
| LIRI-JP::SP107050 | Liver | LIRI-JP | 536 | 0 | 55 | 22 | 10 | 0.029 | 0.010 | 0.010 | 0.035 | 0.0673 | 0.9464 | 0.9704 | Increasing |
| PACA-AU::SP71132 | Pancreas | PACA-AU | 850 | 0 | 0 | 12 | 11.5 | 0.011 | 0.005 | 0.015 | 0.007 | 0.0665 | 0.9470 | 0.9704 | Increasing |
| PACA-AU::SP110816 | Pancreas | PACA-AU | 318 | 0 | 100 | 12 | 10 | 0.015 | 0.005 | 0.006 | 0.013 | 0.0665 | 0.9470 | 0.9704 | Decreasing |
| PACA-AU::SP77131 | Pancreas | PACA-AU | 734 | 0 | 0 | 12 | 11 | 0.011 | 0.011 | 0.003 | 0.013 | 0.0665 | 0.9470 | 0.9704 | Decreasing |
| RECA-EU::SP103742 | Kidney | RECA-EU | 969 | 0 | 136 | 67 | 9 | 0.059 | 0.047 | 0.045 | 0.065 | -0.0648 | 0.9484 | 0.9711 | Increasing |
| PACA-AU::SP107813 | Pancreas | PACA-AU | 656 | 0 | 0 | 13 | 15 | 0.011 | 0.005 | 0.018 | 0.007 | -0.0605 | 0.9518 | 0.9739 | Increasing |
| PACA-CA::SP117660 | Pancreas | PACA-CA | 541 | 0 | 155 | 13 | 12 | 0.008 | 0.014 | 0.008 | 0.010 | 0.0575 | 0.9541 | 0.9757 | Decreasing |
| PACA-CA::SP117001 | Pancreas | PACA-CA | 482 | 0 | 0 | 14 | 14.5 | 0.011 | 0.011 | 0.008 | 0.013 | -0.0557 | 0.9556 | 0.9765 | Decreasing |
| LUSC-US::SP57586 | Lung | LUSC-US | 1310 | 0 | 189 | 35 | 11.5 | 0.038 | 0.021 | 0.026 | 0.040 | 0.0534 | 0.9574 | 0.9774 | Increasing |
| LIRI-JP::SP50161 | Liver | LIRI-JP | 737 | 83 | 140 | 75 | 12 | 0.067 | 0.055 | 0.067 | 0.064 | -0.0529 | 0.9578 | 0.9774 | Increasing |
| PACA-AU::SP110789 | Pancreas | PACA-AU | 1173 | 663 | 213 | 209 | 14 | 0.161 | 0.171 | 0.152 | 0.167 | 0.0511 | 0.9593 | 0.9777 | Decreasing |
| MELA-AU::SP124307 | Skin | SKCM-US | 751 | 0 | 114 | 50 | 17 | 0.034 | 0.054 | 0.038 | 0.039 | 0.0507 | 0.9595 | 0.9777 | Decreasing |
| RECA-EU::SP103396 | Kidney | RECA-EU | 316 | 0 | 56 | 25 | 9 | 0.010 | 0.033 | 0.011 | 0.020 | -0.0485 | 0.9613 | 0.9789 | Decreasing |
| LUSC-US::SP56704 | Lung | LUSC-US | 2182 | 0 | 0 | 21 | 9 | 0.022 | 0.011 | 0.023 | 0.017 | 0.0414 | 0.9670 | 0.9826 | Increasing |
| COAD-US::SP21528 | CRC | COAD-US | 1317 | 0 | 0 | 15 | 12 | 0.012 | 0.016 | 0.007 | 0.017 | 0.0424 | 0.9662 | 0.9826 | Decreasing |
| BRCA-UK::SP116335 | Breast | BRCA-US | 511 | 81 | 67 | 37 | 10 | 0.031 | 0.034 | 0.025 | 0.037 | -0.0419 | 0.9666 | 0.9826 | Decreasing |
| BRCA-US::SP11808 | Breast | BRCA-US | 343 | 0 | 78 | 12 | 11 | 0.010 | 0.011 | 0.009 | 0.011 | 0.0373 | 0.9702 | 0.9852 | Decreasing |
| PACA-CA::SP117164 | Pancreas | PACA-CA | 388 | 0 | 0 | 11 | 11 | 0.008 | 0.006 | 0.017 | 0.003 | 0.0312 | 0.9751 | 0.9867 | Increasing |
| HNSC-US::SP33688 | Head&Neck | HNSC-US | 1352 | 639 | 390 | 261 | 15 | 0.230 | 0.227 | 0.186 | 0.259 | 0.0322 | 0.9743 | 0.9867 | Decreasing |
| MALY-DE::SP116610 | Blood | MALY-DE | 1134 | 165 | 0 | 49 | 14 | 0.032 | 0.045 | 0.029 | 0.042 | -0.0320 | 0.9745 | 0.9867 | Decreasing |
| PACA-AU::SP107945 | Pancreas | PACA-AU | 609 | 0 | 0 | 17 | 11.5 | 0.011 | 0.016 | 0.012 | 0.013 | -0.0343 | 0.9727 | 0.9867 | Decreasing |
| PACA-CA::SP125722 | Pancreas | PACA-CA | 269 | 0 | 66 | 11 | 11.5 | 0.011 | 0.008 | 0.003 | 0.013 | 0.0312 | 0.9751 | 0.9867 | Decreasing |
| SKCM-US::SP82459 | Skin | SKCM-US | 447 | 0 | 67 | 21 | 16 | 0.024 | 0.010 | 0.019 | 0.022 | -0.0275 | 0.9781 | 0.9876 | Increasing |
| ORCA-IN::SP118015 | Head&Neck | ORCA-IN | 373 | 0 | 0 | 18 | 13 | 0.019 | 0.010 | 0.012 | 0.020 | 0.0283 | 0.9774 | 0.9876 | Increasing |
| BRCA-UK::SP116357 | Breast | BRCA-US | 332 | 136 | 142 | 69 | 16 | 0.059 | 0.058 | 0.060 | 0.058 | -0.0285 | 0.9773 | 0.9876 | Increasing |
| LIHC-US::SP98305 | Liver | LIHC-US | 1171 | 0 | 0 | 46 | 12 | 0.038 | 0.042 | 0.035 | 0.042 | -0.0216 | 0.9827 | 0.9883 | Decreasing |
| LIHC-US::SP120755 | Liver | LIHC-US | 139 | 0 | 30 | 14 | 13.5 | 0.009 | 0.013 | 0.016 | 0.006 | -0.0234 | 0.9813 | 0.9883 | Decreasing |
| PACA-CA::SP125763 | Pancreas | PACA-CA | 1240 | 0 | 0 | 25 | 11 | 0.011 | 0.025 | 0.025 | 0.013 | -0.0209 | 0.9833 | 0.9883 | Increasing |
| BTCA-SG::SP117775 | Biliary | BTCA-SG | 1011 | 0 | 140 | 33 | 12.5 | 0.022 | 0.050 | 0.024 | 0.028 | -0.0212 | 0.9831 | 0.9883 | Decreasing |
| KIRC-US::SP34186 | Kidney | KIRC-US | 1033 | 0 | 164 | 64 | 9 | 0.051 | 0.059 | 0.052 | 0.054 | -0.0205 | 0.9836 | 0.9883 | Decreasing |
| LIRI-JP::SP106998 | Liver | LIRI-JP | 803 | 0 | 0 | 24 | 11 | 0.016 | 0.023 | 0.025 | 0.012 | -0.0218 | 0.9826 | 0.9883 | Decreasing |
| GBM-US::SP29559 | Brain | GBM-US | 819 | 0 | 323 | 109 | 11 | 0.114 | 0.083 | 0.077 | 0.117 | 0.0200 | 0.9840 | 0.9883 | Decreasing |
| LUSC-US::SP57189 | Lung | LUSC-US | 4718 | 0 | 0 | 47 | 12 | 0.038 | 0.045 | 0.033 | 0.045 | -0.0198 | 0.9842 | 0.9883 | Decreasing |
| SKCM-US::SP83312 | Skin | SKCM-US | 485 | 0 | 102 | 37 | 15 | 0.027 | 0.034 | 0.038 | 0.022 | 0.0177 | 0.9859 | 0.9888 | Decreasing |
| STAD-US::SP85339 | Stomach | STAD-US | 1697 | 0 | 0 | 20 | 12 | 0.018 | 0.017 | 0.013 | 0.021 | 0.0174 | 0.9861 | 0.9888 | Decreasing |

| | | | | | | | | | | | | | | | |
|---|---|---|---|---|---|---|---|---|---|---|---|---|---|---|---|
| MELA-AU::SP124460 | Skin | SKCM-US | 1031 | 0 | 113 | 40 | 13.5 | 0.027 | 0.041 | 0.035 | 0.028 | 0.0141 | 0.9887 | 0.9908 | Decreasing |
| RECA-EU::SP102921 | Kidney | RECA-EU | 587 | 0 | 100 | 43 | 9 | 0.036 | 0.031 | 0.034 | 0.035 | 0.0091 | 0.9927 | 0.9941 | Increasing |
| PACA-CA::SP125698 | Pancreas | PACA-CA | 1261 | 592 | 261 | 223 | 13 | 0.162 | 0.187 | 0.166 | 0.172 | -0.0074 | 0.9941 | 0.9948 | Decreasing |
| COAD-US::SP19670 | CRC | COAD-US | 1181 | 0 | 0 | 18 | 11 | 0.018 | 0.005 | 0.030 | 0.006 | 0.0006 | 0.9995 | 0.9995 | Increasing |
| LGG-US::SP47708 | Brain | LGG-US | 241 | 0 | 0 | 7 | 5.5 | Samples with <= 10 >= 5 bp deletions weren't analyzed | | | | | | | |
| THCA-US::SP85623 | Thyroid | THCA-US | 69 | 0 | 14 | 5 | 10.5 | Samples with <= 10 >= 5 bp deletions weren't analyzed | | | | | | | |
| PRAD-CA::SP112799 | Prostate | PRAD-US | 157 | 0 | 19 | 9 | 12 | Samples with <= 10 >= 5 bp deletions weren't analyzed | | | | | | | |
| COAD-US::SP22031 | CRC | COAD-US | 3532 | 0 | 0 | 3 | 17 | Samples with <= 10 >= 5 bp deletions weren't analyzed | | | | | | | |
| MALY-DE::SP59388 | Blood | MALY-DE | 382 | 0 | 0 | 1 | 5.5 | Samples with <= 10 >= 5 bp deletions weren't analyzed | | | | | | | |
| PRAD-US::SP80037 | Prostate | PRAD-US | 313 | 0 | 45 | 9 | 17 | Samples with <= 10 >= 5 bp deletions weren't analyzed | | | | | | | |
| PAEN-AU::SP102617 | Pancreas | PAEN-AU | 168 | 0 | 21 | 5 | 10.5 | Samples with <= 10 >= 5 bp deletions weren't analyzed | | | | | | | |
| GBM-US::SP27201 | Brain | GBM-US | 501 | 0 | 0 | 8 | 5 | Samples with <= 10 >= 5 bp deletions weren't analyzed | | | | | | | |
| PAEN-AU::SP102605 | Pancreas | PAEN-AU | 49 | 0 | 0 | 3 | 7 | Samples with <= 10 >= 5 bp deletions weren't analyzed | | | | | | | |
| MALY-DE::SP59416 | Blood | MALY-DE | 503 | 0 | 0 | 8 | 11 | Samples with <= 10 >= 5 bp deletions weren't analyzed | | | | | | | |
| GBM-US::SP28791 | Brain | GBM-US | 167 | 0 | 0 | 3 | 5 | Samples with <= 10 >= 5 bp deletions weren't analyzed | | | | | | | |
| CESC-US::SP13206 | Cervix | CESC-US | 362 | 62 | 0 | 8 | 10.5 | Samples with <= 10 >= 5 bp deletions weren't analyzed | | | | | | | |
| CLLE-ES::SP13291 | Blood | CLLE-ES | 137 | 0 | 0 | 5 | 16.5 | Samples with <= 10 >= 5 bp deletions weren't analyzed | | | | | | | |
| PACA-AU::SP110736 | Pancreas | PACA-AU | 540 | 0 | 0 | 9 | 13.5 | Samples with <= 10 >= 5 bp deletions weren't analyzed | | | | | | | |
| PAEN-AU::SP117086 | Pancreas | PAEN-AU | 15 | 0 | 4 | 2 | 11 | Samples with <= 10 >= 5 bp deletions weren't analyzed | | | | | | | |
| MALY-DE::SP124977 | Blood | MALY-DE | 811 | 0 | 0 | 6 | 11 | Samples with <= 10 >= 5 bp deletions weren't analyzed | | | | | | | |
| THCA-US::SP85491 | Thyroid | THCA-US | 99 | 0 | 19 | 6 | 7 | Samples with <= 10 >= 5 bp deletions weren't analyzed | | | | | | | |
| PAEN-AU::SP117700 | Pancreas | PAEN-AU | 104 | 0 | 24 | 5 | 10 | Samples with <= 10 >= 5 bp deletions weren't analyzed | | | | | | | |
| CLLE-ES::SP13514 | Blood | CLLE-ES | 174 | 0 | 0 | 4 | 10 | Samples with <= 10 >= 5 bp deletions weren't analyzed | | | | | | | |
| MALY-DE::SP59356 | Blood | MALY-DE | 417 | 0 | 0 | 5 | 12.5 | Samples with <= 10 >= 5 bp deletions weren't analyzed | | | | | | | |
| PACA-CA::SP125807 | Pancreas | PACA-CA | 885 | 0 | 0 | 10 | 12 | Samples with <= 10 >= 5 bp deletions weren't analyzed | | | | | | | |
| THCA-US::SP87582 | Thyroid | THCA-US | 66 | 0 | 9 | 5 | 7 | Samples with <= 10 >= 5 bp deletions weren't analyzed | | | | | | | |
| PRAD-CA::SP112835 | Prostate | PRAD-US | 239 | 0 | 0 | 8 | 7 | Samples with <= 10 >= 5 bp deletions weren't analyzed | | | | | | | |
| GBM-US::SP23739 | Brain | GBM-US | 254 | 0 | 58 | 9 | 6 | Samples with <= 10 >= 5 bp deletions weren't analyzed | | | | | | | |
| GACA-CN::SP135184 | Stomach | STAD-US | 7 | 0 | 0 | 1 | 5 | Samples with <= 10 >= 5 bp deletions weren't analyzed | | | | | | | |
| PRAD-CA::SP112955 | Prostate | PRAD-US | 210 | 0 | 17 | 7 | 9 | Samples with <= 10 >= 5 bp deletions weren't analyzed | | | | | | | |
| CLLE-ES::SP115117 | Blood | CLLE-ES | 64 | 0 | 0 | 2 | 8 | Samples with <= 10 >= 5 bp deletions weren't analyzed | | | | | | | |
| BRCA-UK::SP2293 | Breast | BRCA-US | 346 | 0 | 0 | 10 | 10.5 | Samples with <= 10 >= 5 bp deletions weren't analyzed | | | | | | | |
| PRAD-CA::SP112807 | Prostate | PRAD-US | 311 | 0 | 0 | 4 | 9 | Samples with <= 10 >= 5 bp deletions weren't analyzed | | | | | | | |
| OV-US::SP59707 | Ovary | OV-US | 189 | 0 | 27 | 9 | 10 | Samples with <= 10 >= 5 bp deletions weren't analyzed | | | | | | | |
| CLLE-ES::SP115114 | Blood | CLLE-ES | 222 | 0 | 0 | 5 | 9 | Samples with <= 10 >= 5 bp deletions weren't analyzed | | | | | | | |
| THCA-US::SP89090 | Thyroid | THCA-US | 73 | 0 | 7 | 4 | 8 | Samples with <= 10 >= 5 bp deletions weren't analyzed | | | | | | | |
| STAD-US::SP85251 | Stomach | STAD-US | 416 | 0 | 0 | 9 | 12 | Samples with <= 10 >= 5 bp deletions weren't analyzed | | | | | | | |
| LUAD-US::SP50406 | Lung | LUAD-US | 13 | 0 | 0 | 0 | NA | Samples with <= 10 >= 5 bp deletions weren't analyzed | | | | | | | |
| CESC-US::SP109941 | Cervix | CESC-US | 214 | 48 | 0 | 7 | 8 | Samples with <= 10 >= 5 bp deletions weren't analyzed | | | | | | | |
| GACA-CN::SP135226 | Stomach | STAD-US | 229 | 0 | 0 | 5 | 5.5 | Samples with <= 10 >= 5 bp deletions weren't analyzed | | | | | | | |
| PRAD-CA::SP112911 | Prostate | PRAD-US | 165 | 0 | 30 | 8 | 12.5 | Samples with <= 10 >= 5 bp deletions weren't analyzed | | | | | | | |
| CESC-US::SP109801 | Cervix | CESC-US | 398 | 54 | 0 | 8 | 12 | Samples with <= 10 >= 5 bp deletions weren't analyzed | | | | | | | |
| STAD-US::SP84408 | Stomach | STAD-US | 284 | 0 | 38 | 3 | 11 | Samples with <= 10 >= 5 bp deletions weren't analyzed | | | | | | | |
| LIRI-JP::SP107073 | Liver | LIRI-JP | 50 | 0 | 9 | 1 | 8 | Samples with <= 10 >= 5 bp deletions weren't analyzed | | | | | | | |
| PRAD-CA::SP112847 | Prostate | PRAD-US | 191 | 0 | 24 | 6 | 15 | Samples with <= 10 >= 5 bp deletions weren't analyzed | | | | | | | |
| COAD-US::SP96124 | CRC | COAD-US | 1699 | 0 | 0 | 8 | 12.5 | Samples with <= 10 >= 5 bp deletions weren't analyzed | | | | | | | |
| CLLE-ES::SP13279 | Blood | CLLE-ES | 100 | 0 | 0 | 3 | 12 | Samples with <= 10 >= 5 bp deletions weren't analyzed | | | | | | | |
| CLLE-ES::SP13487 | Blood | CLLE-ES | 89 | 0 | 0 | 4 | 18.5 | Samples with <= 10 >= 5 bp deletions weren't analyzed | | | | | | | |
| PRAD-CA::SP112919 | Prostate | PRAD-US | 209 | 0 | 0 | 2 | 7 | Samples with <= 10 >= 5 bp deletions weren't analyzed | | | | | | | |
| CLLE-ES::SP13361 | Blood | CLLE-ES | 184 | 0 | 0 | 8 | 10 | Samples with <= 10 >= 5 bp deletions weren't analyzed | | | | | | | |

| | | | | | | | | |
|---|---|---|---|---|---|---|---|---|
| BRCA-US::SP2799 | Breast | BRCA-US | 196 | 0 | 59 | 9 | 7 | Samples with <= 10 >= 5 bp deletions weren't analyzed |
| MELA-AU::SP124339 | Skin | SKCM-US | 130 | 0 | 37 | 10 | 10 | Samples with <= 10 >= 5 bp deletions weren't analyzed |
| CLLE-ES::SP13421 | Blood | CLLE-ES | 180 | 0 | 0 | 5 | 16.5 | Samples with <= 10 >= 5 bp deletions weren't analyzed |
| READ-US::SP82087 | CRC | READ-US | 1074 | 0 | 0 | 9 | 14 | Samples with <= 10 >= 5 bp deletions weren't analyzed |
| PRAD-CA::SP102652 | Prostate | PRAD-US | 223 | 0 | 26 | 10 | 17 | Samples with <= 10 >= 5 bp deletions weren't analyzed |
| CLLE-ES::SP116738 | Blood | CLLE-ES | 105 | 0 | 0 | 1 | 9 | Samples with <= 10 >= 5 bp deletions weren't analyzed |
| CLLE-ES::SP13293 | Blood | CLLE-ES | 104 | 0 | 0 | 4 | 6.5 | Samples with <= 10 >= 5 bp deletions weren't analyzed |
| THCA-US::SP85952 | Thyroid | THCA-US | 57 | 0 | 19 | 8 | 11 | Samples with <= 10 >= 5 bp deletions weren't analyzed |
| MALY-DE::SP116712 | Blood | MALY-DE | 117 | 0 | 0 | 5 | 21 | Samples with <= 10 >= 5 bp deletions weren't analyzed |
| LUAD-US::SP55387 | Lung | LUAD-US | 269 | 0 | 0 | 5 | 15 | Samples with <= 10 >= 5 bp deletions weren't analyzed |
| PAEN-AU::SP102507 | Pancreas | PAEN-AU | 174 | 0 | 23 | 7 | 9 | Samples with <= 10 >= 5 bp deletions weren't analyzed |
| THCA-US::SP85840 | Thyroid | THCA-US | 95 | 0 | 10 | 2 | 9.5 | Samples with <= 10 >= 5 bp deletions weren't analyzed |
| THCA-US::SP87675 | Thyroid | THCA-US | 70 | 0 | 11 | 5 | 10.5 | Samples with <= 10 >= 5 bp deletions weren't analyzed |
| THCA-US::SP85836 | Thyroid | THCA-US | 156 | 0 | 27 | 10 | 15 | Samples with <= 10 >= 5 bp deletions weren't analyzed |
| CLLE-ES::SP13281 | Blood | CLLE-ES | 94 | 0 | 0 | 2 | 16 | Samples with <= 10 >= 5 bp deletions weren't analyzed |
| MALY-DE::SP116726 | Blood | MALY-DE | 1071 | 0 | 0 | 9 | 9 | Samples with <= 10 >= 5 bp deletions weren't analyzed |
| PAEN-AU::SP102485 | Pancreas | PAEN-AU | 46 | 0 | 10 | 4 | 7 | Samples with <= 10 >= 5 bp deletions weren't analyzed |
| LIRI-JP::SP107094 | Liver | LIRI-JP | 205 | 0 | 0 | 9 | 9 | Samples with <= 10 >= 5 bp deletions weren't analyzed |
| GACA-CN::SP135262 | Stomach | STAD-US | 133 | 0 | 0 | 2 | 8 | Samples with <= 10 >= 5 bp deletions weren't analyzed |
| THCA-US::SP105708 | Thyroid | THCA-US | 97 | 0 | 19 | 6 | 12 | Samples with <= 10 >= 5 bp deletions weren't analyzed |
| CLLE-ES::SP13677 | Blood | CLLE-ES | 93 | 0 | 0 | 2 | 7.5 | Samples with <= 10 >= 5 bp deletions weren't analyzed |
| OV-US::SP69155 | Ovary | OV-US | 130 | 0 | 21 | 6 | 12 | Samples with <= 10 >= 5 bp deletions weren't analyzed |
| PACA-CA::SP117972 | Pancreas | PACA-CA | 459 | 0 | 0 | 9 | 14.5 | Samples with <= 10 >= 5 bp deletions weren't analyzed |
| LAML-KR::SP109002 | Blood | LAML-US | 53 | 0 | 0 | 0 | 6 | Samples with <= 10 >= 5 bp deletions weren't analyzed |
| MALY-DE::SP116723 | Blood | MALY-DE | 252 | 0 | 0 | 4 | 11 | Samples with <= 10 >= 5 bp deletions weren't analyzed |
| THCA-US::SP85818 | Thyroid | THCA-US | 2 | 0 | 0 | 0 | NA | Samples with <= 10 >= 5 bp deletions weren't analyzed |
| HNSC-US::SP32958 | Head&Neck | HNSC-US | 562 | 0 | 0 | 7 | 13 | Samples with <= 10 >= 5 bp deletions weren't analyzed |
| PRAD-CA::SP112967 | Prostate | PRAD-US | 24 | 0 | 0 | 0 | NA | Samples with <= 10 >= 5 bp deletions weren't analyzed |
| CLLE-ES::SP13399 | Blood | CLLE-ES | 109 | 0 | 0 | 3 | 8.5 | Samples with <= 10 >= 5 bp deletions weren't analyzed |
| PRAD-CA::SP102622 | Prostate | PRAD-US | 366 | 0 | 41 | 10 | 9 | Samples with <= 10 >= 5 bp deletions weren't analyzed |
| PACA-AU::SP70082 | Pancreas | PACA-AU | 218 | 0 | 56 | 10 | 11 | Samples with <= 10 >= 5 bp deletions weren't analyzed |
| MELA-AU::SP124355 | Skin | SKCM-US | 228 | 0 | 43 | 10 | 13.5 | Samples with <= 10 >= 5 bp deletions weren't analyzed |
| LIRI-JP::SP99287 | Liver | LIRI-JP | 167 | 0 | 0 | 7 | 11 | Samples with <= 10 >= 5 bp deletions weren't analyzed |
| CLLE-ES::SP116732 | Blood | CLLE-ES | 144 | 0 | 0 | 8 | 13.5 | Samples with <= 10 >= 5 bp deletions weren't analyzed |
| MALY-DE::SP59276 | Blood | MALY-DE | 185 | 0 | 0 | 4 | 6 | Samples with <= 10 >= 5 bp deletions weren't analyzed |
| CLLE-ES::SP115073 | Blood | CLLE-ES | 69 | 0 | 0 | 3 | 10 | Samples with <= 10 >= 5 bp deletions weren't analyzed |
| PACA-CA::SP125800 | Pancreas | PACA-CA | 56 | 8 | 0 | 3 | 21 | Samples with <= 10 >= 5 bp deletions weren't analyzed |
| LIRI-JP::SP107115 | Liver | LIRI-JP | 17 | 0 | 0 | 0 | NA | Samples with <= 10 >= 5 bp deletions weren't analyzed |
| MALY-DE::SP116703 | Blood | MALY-DE | 229 | 0 | 0 | 5 | 8 | Samples with <= 10 >= 5 bp deletions weren't analyzed |
| THCA-US::SP105922 | Thyroid | THCA-US | 61 | 0 | 15 | 3 | 8 | Samples with <= 10 >= 5 bp deletions weren't analyzed |
| GBM-US::SP25332 | Brain | GBM-US | 157 | 0 | 23 | 6 | 7.5 | Samples with <= 10 >= 5 bp deletions weren't analyzed |
| PRAD-US::SP80213 | Prostate | PRAD-US | 11 | 0 | 0 | 0 | 39 | Samples with <= 10 >= 5 bp deletions weren't analyzed |
| PACA-CA::SP117351 | Pancreas | PACA-CA | 341 | 0 | 0 | 4 | 10 | Samples with <= 10 >= 5 bp deletions weren't analyzed |
| BRCA-US::SP4593 | Breast | BRCA-US | 164 | 0 | 57 | 6 | 13 | Samples with <= 10 >= 5 bp deletions weren't analyzed |
| PAEN-AU::SP117228 | Pancreas | PAEN-AU | 114 | 0 | 12 | 7 | 6 | Samples with <= 10 >= 5 bp deletions weren't analyzed |
| CLLE-ES::SP13307 | Blood | CLLE-ES | 136 | 0 | 0 | 1 | 12 | Samples with <= 10 >= 5 bp deletions weren't analyzed |
| PAEN-AU::SP118044 | Pancreas | PAEN-AU | 11 | 0 | 2 | 1 | 9 | Samples with <= 10 >= 5 bp deletions weren't analyzed |
| CLLE-ES::SP13299 | Blood | CLLE-ES | 219 | 0 | 0 | 4 | 10 | Samples with <= 10 >= 5 bp deletions weren't analyzed |
| BRCA-US::SP8085 | Breast | BRCA-US | 323 | 0 | 51 | 10 | 9.5 | Samples with <= 10 >= 5 bp deletions weren't analyzed |
| PRAD-CA::SP112793 | Prostate | PRAD-US | 275 | 0 | 40 | 9 | 11 | Samples with <= 10 >= 5 bp deletions weren't analyzed |

| | | | | | | | | | |
|---|---|---|---|---|---|---|---|---|---|
| MALY-DE::SP59296 | Blood | MALY-DE | 270 | 0 | 0 | 0 | 32 | | Samples with <= 10 >= 5 bp deletions weren't analyzed |
| PAEN-AU::SP117984 | Pancreas | PAEN-AU | 120 | 0 | 11 | 7 | 12 | | Samples with <= 10 >= 5 bp deletions weren't analyzed |
| THCA-US::SP86118 | Thyroid | THCA-US | 94 | 0 | 32 | 8 | 14 | | Samples with <= 10 >= 5 bp deletions weren't analyzed |
| PACA-CA::SP117878 | Pancreas | PACA-CA | 533 | 0 | 0 | 9 | 10.5 | | Samples with <= 10 >= 5 bp deletions weren't analyzed |
| CLLE-ES::SP13481 | Blood | CLLE-ES | 82 | 0 | 0 | 4 | 9 | | Samples with <= 10 >= 5 bp deletions weren't analyzed |
| LGG-US::SP47990 | Brain | LGG-US | 145 | 0 | 54 | 8 | 10 | | Samples with <= 10 >= 5 bp deletions weren't analyzed |
| BRCA-US::SP11235 | Breast | BRCA-US | 242 | 65 | 0 | 9 | 9 | | Samples with <= 10 >= 5 bp deletions weren't analyzed |
| MALY-DE::SP59280 | Blood | MALY-DE | 493 | 0 | 0 | 3 | 17 | | Samples with <= 10 >= 5 bp deletions weren't analyzed |
| MALY-DE::SP59344 | Blood | MALY-DE | 221 | 0 | 0 | 3 | 9 | | Samples with <= 10 >= 5 bp deletions weren't analyzed |
| PACA-AU::SP108017 | Pancreas | PACA-AU | 290 | 0 | 0 | 4 | 11 | | Samples with <= 10 >= 5 bp deletions weren't analyzed |
| LIRI-JP::SP107132 | Liver | LIRI-JP | 100 | 0 | 12 | 5 | 9 | | Samples with <= 10 >= 5 bp deletions weren't analyzed |
| MALY-DE::SP59460 | Blood | MALY-DE | 829 | 0 | 0 | 10 | 12 | | Samples with <= 10 >= 5 bp deletions weren't analyzed |
| PACA-CA::SP125772 | Pancreas | PACA-CA | 244 | 0 | 67 | 4 | 13 | | Samples with <= 10 >= 5 bp deletions weren't analyzed |
| GBM-US::SP28335 | Brain | GBM-US | 315 | 0 | 78 | 10 | 6 | | Samples with <= 10 >= 5 bp deletions weren't analyzed |
| PRAD-CA::SP112963 | Prostate | PRAD-US | 295 | 0 | 0 | 7 | 15 | | Samples with <= 10 >= 5 bp deletions weren't analyzed |
| BRCA-US::SP10635 | Breast | BRCA-US | 150 | 0 | 43 | 8 | 11 | | Samples with <= 10 >= 5 bp deletions weren't analyzed |
| THCA-US::SP86306 | Thyroid | THCA-US | 159 | 0 | 42 | 3 | 8 | | Samples with <= 10 >= 5 bp deletions weren't analyzed |
| CLLE-ES::SP116780 | Blood | CLLE-ES | 119 | 0 | 0 | 4 | 12 | | Samples with <= 10 >= 5 bp deletions weren't analyzed |
| THCA-US::SP87099 | Thyroid | THCA-US | 104 | 0 | 25 | 6 | 11 | | Samples with <= 10 >= 5 bp deletions weren't analyzed |
| MALY-DE::SP59404 | Blood | MALY-DE | 279 | 0 | 0 | 1 | 25 | | Samples with <= 10 >= 5 bp deletions weren't analyzed |
| GBM-US::SP26709 | Brain | GBM-US | 234 | 0 | 28 | 6 | 6 | | Samples with <= 10 >= 5 bp deletions weren't analyzed |
| LUAD-US::SP50485 | Lung | LUAD-US | 156 | 19 | 0 | 8 | 10.5 | | Samples with <= 10 >= 5 bp deletions weren't analyzed |
| BRCA-US::SP5381 | Breast | BRCA-US | 109 | 13 | 14 | 5 | 12 | | Samples with <= 10 >= 5 bp deletions weren't analyzed |
| PACA-CA::SP118046 | Pancreas | PACA-CA | 790 | 0 | 0 | 10 | 12 | | Samples with <= 10 >= 5 bp deletions weren't analyzed |
| LAML-KR::SP109001 | Blood | LAML-US | 11 | 0 | 0 | 2 | 7.5 | | Samples with <= 10 >= 5 bp deletions weren't analyzed |
| GACA-CN::SP135230 | Stomach | STAD-US | 314 | 0 | 0 | 6 | 10 | | Samples with <= 10 >= 5 bp deletions weren't analyzed |
| ORCA-IN::SP117189 | Head&Neck | ORCA-IN | 111 | 0 | 23 | 4 | 12.5 | | Samples with <= 10 >= 5 bp deletions weren't analyzed |
| MELA-AU::SP124353 | Skin | SKCM-US | 141 | 0 | 33 | 10 | 14.5 | | Samples with <= 10 >= 5 bp deletions weren't analyzed |
| MELA-AU::SP124278 | Skin | SKCM-US | 350 | 0 | 0 | 10 | 17 | | Samples with <= 10 >= 5 bp deletions weren't analyzed |
| CLLE-ES::SP13433 | Blood | CLLE-ES | 121 | 0 | 0 | 3 | 7 | | Samples with <= 10 >= 5 bp deletions weren't analyzed |
| PACA-CA::SP125766 | Pancreas | PACA-CA | 510 | 0 | 106 | 8 | 9 | | Samples with <= 10 >= 5 bp deletions weren't analyzed |
| PAEN-AU::SP102573 | Pancreas | PAEN-AU | 54 | 10 | 7 | 7 | 17.5 | | Samples with <= 10 >= 5 bp deletions weren't analyzed |
| GBM-US::SP25350 | Brain | GBM-US | 445 | 0 | 0 | 5 | 6 | | Samples with <= 10 >= 5 bp deletions weren't analyzed |
| LUAD-US::SP51037 | Lung | LUAD-US | 231 | 0 | 30 | 10 | 9 | | Samples with <= 10 >= 5 bp deletions weren't analyzed |
| GACA-CN::SP135205 | Stomach | STAD-US | 99 | 0 | 0 | 2 | 10 | | Samples with <= 10 >= 5 bp deletions weren't analyzed |
| LIRI-JP::SP99297 | Liver | LIRI-JP | 23 | 0 | 4 | 1 | 5.5 | | Samples with <= 10 >= 5 bp deletions weren't analyzed |
| CLLE-ES::SP115064 | Blood | CLLE-ES | 169 | 0 | 0 | 2 | 7.5 | | Samples with <= 10 >= 5 bp deletions weren't analyzed |
| LGG-US::SP48414 | Brain | LGG-US | 152 | 0 | 0 | 4 | 8 | | Samples with <= 10 >= 5 bp deletions weren't analyzed |
| PAEN-AU::SP117667 | Pancreas | PAEN-AU | 28 | 4 | 6 | 3 | 15 | | Samples with <= 10 >= 5 bp deletions weren't analyzed |
| PAEN-AU::SP106815 | Pancreas | PAEN-AU | 201 | 27 | 0 | 9 | 13.5 | | Samples with <= 10 >= 5 bp deletions weren't analyzed |
| LGG-US::SP48135 | Brain | LGG-US | 363 | 0 | 0 | 5 | 10.5 | | Samples with <= 10 >= 5 bp deletions weren't analyzed |
| CLLE-ES::SP13359 | Blood | CLLE-ES | 97 | 0 | 0 | 8 | 13 | | Samples with <= 10 >= 5 bp deletions weren't analyzed |
| LIRI-JP::SP98937 | Liver | LIRI-JP | 152 | 0 | 17 | 7 | 7 | | Samples with <= 10 >= 5 bp deletions weren't analyzed |
| MALY-DE::SP116614 | Blood | MALY-DE | 1293 | 0 | 0 | 9 | 7 | | Samples with <= 10 >= 5 bp deletions weren't analyzed |
| PRAD-CA::SP112949 | Prostate | PRAD-US | 89 | 0 | 11 | 5 | 14 | | Samples with <= 10 >= 5 bp deletions weren't analyzed |
| CLLE-ES::SP116734 | Blood | CLLE-ES | 195 | 0 | 0 | 1 | 10 | | Samples with <= 10 >= 5 bp deletions weren't analyzed |
| MALY-DE::SP59464 | Blood | MALY-DE | 286 | 0 | 0 | 2 | 10 | | Samples with <= 10 >= 5 bp deletions weren't analyzed |
| CLLE-ES::SP116770 | Blood | CLLE-ES | 80 | 0 | 0 | 7 | 10 | | Samples with <= 10 >= 5 bp deletions weren't analyzed |
| THCA-US::SP87434 | Thyroid | THCA-US | 72 | 0 | 9 | 2 | 8 | | Samples with <= 10 >= 5 bp deletions weren't analyzed |
| CLLE-ES::SP13323 | Blood | CLLE-ES | 98 | 0 | 0 | 7 | 8.5 | | Samples with <= 10 >= 5 bp deletions weren't analyzed |

| | | | | | | | | |
|---|---|---|---|---|---|---|---|---|
| SKCM-US::SP113197 | Skin | SKCM-US | 178 | 0 | 20 | 2 | 12 | Samples with <= 10 >= 5 bp deletions weren't analyzed |
| PACA-CA::SP77872 | Pancreas | PACA-CA | 134 | 0 | 37 | 5 | 10.5 | Samples with <= 10 >= 5 bp deletions weren't analyzed |
| GBM-US::SP25833 | Brain | GBM-US | 413 | 0 | 0 | 9 | 5 | Samples with <= 10 >= 5 bp deletions weren't analyzed |
| MALY-DE::SP59364 | Blood | MALY-DE | 603 | 0 | 0 | 5 | 10 | Samples with <= 10 >= 5 bp deletions weren't analyzed |
| BRCA-US::SP11045 | Breast | BRCA-US | 119 | 0 | 37 | 7 | 14 | Samples with <= 10 >= 5 bp deletions weren't analyzed |
| PACA-CA::SP125685 | Pancreas | PACA-CA | 446 | 0 | 92 | 8 | 15 | Samples with <= 10 >= 5 bp deletions weren't analyzed |
| PRAD-CA::SP112915 | Prostate | PRAD-US | 277 | 0 | 34 | 8 | 11 | Samples with <= 10 >= 5 bp deletions weren't analyzed |
| CESC-US::SP107640 | Cervix | CESC-US | 304 | 0 | 0 | 6 | 11 | Samples with <= 10 >= 5 bp deletions weren't analyzed |
| PAEN-AU::SP106734 | Pancreas | PAEN-AU | 50 | 0 | 8 | 4 | 8 | Samples with <= 10 >= 5 bp deletions weren't analyzed |
| PRAD-CA::SP112823 | Prostate | PRAD-US | 192 | 0 | 29 | 8 | 13 | Samples with <= 10 >= 5 bp deletions weren't analyzed |
| ORCA-IN::SP117217 | Head&Neck | ORCA-IN | 27 | 0 | 7 | 0 | 6 | Samples with <= 10 >= 5 bp deletions weren't analyzed |
| PACA-CA::SP125717 | Pancreas | PACA-CA | 467 | 0 | 0 | 6 | 12 | Samples with <= 10 >= 5 bp deletions weren't analyzed |
| THCA-US::SP85864 | Thyroid | THCA-US | 26 | 0 | 0 | 0 | 35 | Samples with <= 10 >= 5 bp deletions weren't analyzed |
| PRAD-CA::SP102633 | Prostate | PRAD-US | 118 | 0 | 14 | 4 | 7.5 | Samples with <= 10 >= 5 bp deletions weren't analyzed |
| LGG-US::SP47652 | Brain | LGG-US | 118 | 0 | 0 | 3 | 10 | Samples with <= 10 >= 5 bp deletions weren't analyzed |
| PAEN-AU::SP117796 | Pancreas | PAEN-AU | 143 | 0 | 21 | 10 | 9 | Samples with <= 10 >= 5 bp deletions weren't analyzed |
| PRAD-CA::SP112899 | Prostate | PRAD-US | 200 | 0 | 21 | 3 | 13 | Samples with <= 10 >= 5 bp deletions weren't analyzed |
| GACA-CN::SP135180 | Stomach | STAD-US | 5 | 0 | 0 | 0 | NA | Samples with <= 10 >= 5 bp deletions weren't analyzed |
| GBM-US::SP25905 | Brain | GBM-US | 209 | 0 | 0 | 9 | 8 | Samples with <= 10 >= 5 bp deletions weren't analyzed |
| MALY-DE::SP59444 | Blood | MALY-DE | 277 | 0 | 0 | 7 | 6 | Samples with <= 10 >= 5 bp deletions weren't analyzed |
| PACA-CA::SP125707 | Pancreas | PACA-CA | 30 | 5 | 0 | 1 | 34 | Samples with <= 10 >= 5 bp deletions weren't analyzed |
| PACA-CA::SP125794 | Pancreas | PACA-CA | 595 | 0 | 0 | 8 | 10.5 | Samples with <= 10 >= 5 bp deletions weren't analyzed |
| CLLE-ES::SP13283 | Blood | CLLE-ES | 64 | 0 | 0 | 1 | 17 | Samples with <= 10 >= 5 bp deletions weren't analyzed |
| CLLE-ES::SP116728 | Blood | CLLE-ES | 97 | 0 | 0 | 6 | 7 | Samples with <= 10 >= 5 bp deletions weren't analyzed |
| CLLE-ES::SP13478 | Blood | CLLE-ES | 62 | 0 | 0 | 5 | 13 | Samples with <= 10 >= 5 bp deletions weren't analyzed |
| THCA-US::SP105807 | Thyroid | THCA-US | 104 | 0 | 17 | 2 | 10.5 | Samples with <= 10 >= 5 bp deletions weren't analyzed |
| LAML-KR::SP109006 | Blood | LAML-US | 66 | 0 | 0 | 1 | 7 | Samples with <= 10 >= 5 bp deletions weren't analyzed |
| MALY-DE::SP59270 | Blood | MALY-DE | 171 | 0 | 0 | 1 | 10 | Samples with <= 10 >= 5 bp deletions weren't analyzed |
| GBM-US::SP26439 | Brain | GBM-US | 161 | 0 | 18 | 5 | 9 | Samples with <= 10 >= 5 bp deletions weren't analyzed |
| PACA-AU::SP77339 | Pancreas | PACA-AU | 487 | 0 | 127 | 10 | 12 | Samples with <= 10 >= 5 bp deletions weren't analyzed |
| MALY-DE::SP116652 | Blood | MALY-DE | 191 | 0 | 0 | 3 | 13 | Samples with <= 10 >= 5 bp deletions weren't analyzed |
| PRAD-CA::SP112895 | Prostate | PRAD-US | 294 | 0 | 0 | 10 | 10.5 | Samples with <= 10 >= 5 bp deletions weren't analyzed |
| CLLE-ES::SP116744 | Blood | CLLE-ES | 120 | 0 | 0 | 8 | 13 | Samples with <= 10 >= 5 bp deletions weren't analyzed |
| CLLE-ES::SP115093 | Blood | CLLE-ES | 230 | 0 | 0 | 6 | 6.5 | Samples with <= 10 >= 5 bp deletions weren't analyzed |
| MELA-AU::SP124351 | Skin | SKCM-US | 137 | 0 | 34 | 9 | 10.5 | Samples with <= 10 >= 5 bp deletions weren't analyzed |
| CLLE-ES::SP13315 | Blood | CLLE-ES | 34 | 0 | 0 | 0 | 5 | Samples with <= 10 >= 5 bp deletions weren't analyzed |
| GBM-US::SP24236 | Brain | GBM-US | 604 | 0 | 0 | 6 | 5 | Samples with <= 10 >= 5 bp deletions weren't analyzed |
| THCA-US::SP89291 | Thyroid | THCA-US | 53 | 0 | 11 | 4 | 8 | Samples with <= 10 >= 5 bp deletions weren't analyzed |
| THCA-US::SP88050 | Thyroid | THCA-US | 44 | 0 | 7 | 2 | 11 | Samples with <= 10 >= 5 bp deletions weren't analyzed |
| CLLE-ES::SP13331 | Blood | CLLE-ES | 165 | 0 | 0 | 7 | 8 | Samples with <= 10 >= 5 bp deletions weren't analyzed |
| GACA-CN::SP135217 | Stomach | STAD-US | 1 | 0 | 1 | 0 | NA | Samples with <= 10 >= 5 bp deletions weren't analyzed |
| PAEN-AU::SP117493 | Pancreas | PAEN-AU | 20 | 0 | 0 | 0 | NA | Samples with <= 10 >= 5 bp deletions weren't analyzed |
| THCA-US::SP88593 | Thyroid | THCA-US | 74 | 0 | 9 | 1 | 10 | Samples with <= 10 >= 5 bp deletions weren't analyzed |
| PACA-CA::SP77986 | Pancreas | PACA-CA | 109 | 0 | 24 | 5 | 10 | Samples with <= 10 >= 5 bp deletions weren't analyzed |
| MELA-AU::SP124323 | Skin | SKCM-US | 225 | 0 | 33 | 9 | 11 | Samples with <= 10 >= 5 bp deletions weren't analyzed |
| MALY-DE::SP116720 | Blood | MALY-DE | 252 | 0 | 0 | 3 | 16 | Samples with <= 10 >= 5 bp deletions weren't analyzed |
| PAEN-AU::SP106722 | Pancreas | PAEN-AU | 97 | 11 | 19 | 7 | 20 | Samples with <= 10 >= 5 bp deletions weren't analyzed |
| MALY-DE::SP59284 | Blood | MALY-DE | 478 | 0 | 0 | 4 | 7 | Samples with <= 10 >= 5 bp deletions weren't analyzed |
| PRAD-US::SP79939 | Prostate | PRAD-US | 23 | 0 | 4 | 3 | 14.5 | Samples with <= 10 >= 5 bp deletions weren't analyzed |
| LAML-KR::SP108998 | Blood | LAML-US | 138 | 0 | 0 | 3 | 8 | Samples with <= 10 >= 5 bp deletions weren't analyzed |

| | | | | | | | | |
|---|---|---|---|---|---|---|---|---|
| MELA-AU::SP124423 | Skin | SKCM-US | 79 | 0 | 19 | 6 | 16 | Samples with <= 10 >= 5 bp deletions weren't analyzed |
| PAEN-AU::SP106753 | Pancreas | PAEN-AU | 30 | 7 | 0 | 3 | 7.5 | Samples with <= 10 >= 5 bp deletions weren't analyzed |
| BTCA-SG::SP117332 | Biliary | BTCA-SG | 155 | 0 | 25 | 10 | 10 | Samples with <= 10 >= 5 bp deletions weren't analyzed |
| PRAD-US::SP109649 | Prostate | PRAD-US | 84 | 0 | 14 | 5 | 9 | Samples with <= 10 >= 5 bp deletions weren't analyzed |
| PRAD-CA::SP112795 | Prostate | PRAD-US | 319 | 0 | 0 | 3 | 12 | Samples with <= 10 >= 5 bp deletions weren't analyzed |
| PRAD-CA::SP112927 | Prostate | PRAD-US | 282 | 0 | 0 | 8 | 11.5 | Samples with <= 10 >= 5 bp deletions weren't analyzed |
| PAEN-AU::SP102523 | Pancreas | PAEN-AU | 138 | 0 | 14 | 3 | 13 | Samples with <= 10 >= 5 bp deletions weren't analyzed |
| LIRI-JP::SP99301 | Liver | LIRI-JP | 1282 | 0 | 0 | 1 | 5 | Samples with <= 10 >= 5 bp deletions weren't analyzed |
| LUAD-US::SP50412 | Lung | LUAD-US | 119 | 0 | 13 | 6 | 8 | Samples with <= 10 >= 5 bp deletions weren't analyzed |
| CLLE-ES::SP115108 | Blood | CLLE-ES | 206 | 0 | 0 | 4 | 14.5 | Samples with <= 10 >= 5 bp deletions weren't analyzed |
| THCA-US::SP88158 | Thyroid | THCA-US | 29 | 0 | 7 | 3 | 10.5 | Samples with <= 10 >= 5 bp deletions weren't analyzed |
| OV-AU::SP102055 | Ovary | OV-AU | 209 | 29 | 0 | 4 | 12 | Samples with <= 10 >= 5 bp deletions weren't analyzed |
| THCA-US::SP88776 | Thyroid | THCA-US | 103 | 0 | 24 | 6 | 9 | Samples with <= 10 >= 5 bp deletions weren't analyzed |
| LIRI-JP::SP98989 | Liver | LIRI-JP | 338 | 0 | 0 | 10 | 12 | Samples with <= 10 >= 5 bp deletions weren't analyzed |
| PAEN-AU::SP102591 | Pancreas | PAEN-AU | 78 | 9 | 9 | 6 | 21 | Samples with <= 10 >= 5 bp deletions weren't analyzed |
| MALY-DE::SP59436 | Blood | MALY-DE | 61 | 0 | 0 | 3 | 7 | Samples with <= 10 >= 5 bp deletions weren't analyzed |
| CLLE-ES::SP116752 | Blood | CLLE-ES | 124 | 0 | 0 | 2 | 5 | Samples with <= 10 >= 5 bp deletions weren't analyzed |
| PACA-CA::SP117523 | Pancreas | PACA-CA | 468 | 0 | 137 | 10 | 12 | Samples with <= 10 >= 5 bp deletions weren't analyzed |
| CLLE-ES::SP115085 | Blood | CLLE-ES | 126 | 0 | 0 | 5 | 8 | Samples with <= 10 >= 5 bp deletions weren't analyzed |
| PRAD-CA::SP112933 | Prostate | PRAD-US | 156 | 0 | 14 | 6 | 11.5 | Samples with <= 10 >= 5 bp deletions weren't analyzed |
| THCA-US::SP85725 | Thyroid | THCA-US | 84 | 0 | 10 | 2 | 8 | Samples with <= 10 >= 5 bp deletions weren't analyzed |
| CLLE-ES::SP115102 | Blood | CLLE-ES | 115 | 0 | 0 | 5 | 9.5 | Samples with <= 10 >= 5 bp deletions weren't analyzed |
| COAD-US::SP96118 | CRC | COAD-US | 9479 | 0 | 0 | 7 | 6 | Samples with <= 10 >= 5 bp deletions weren't analyzed |
| THCA-US::SP86660 | Thyroid | THCA-US | 162 | 0 | 42 | 6 | 9 | Samples with <= 10 >= 5 bp deletions weren't analyzed |
| GACA-CN::SP135238 | Stomach | STAD-US | 8 | 0 | 0 | 0 | NA | Samples with <= 10 >= 5 bp deletions weren't analyzed |
| KIRP-US::SP43822 | Kidney | KIRP-US | 462 | 0 | 0 | 9 | 11 | Samples with <= 10 >= 5 bp deletions weren't analyzed |
| THCA-US::SP87903 | Thyroid | THCA-US | 48 | 0 | 9 | 2 | 14 | Samples with <= 10 >= 5 bp deletions weren't analyzed |
| MALY-DE::SP59320 | Blood | MALY-DE | 332 | 0 | 0 | 5 | 6 | Samples with <= 10 >= 5 bp deletions weren't analyzed |
| GBM-US::SP27957 | Brain | GBM-US | 378 | 0 | 0 | 6 | 7.5 | Samples with <= 10 >= 5 bp deletions weren't analyzed |
| LIRI-JP::SP112257 | Liver | LIRI-JP | 49 | 0 | 5 | 3 | 7 | Samples with <= 10 >= 5 bp deletions weren't analyzed |
| UCEC-US::SP94060 | Endometrium | UCEC-US | 379 | 0 | 51 | 4 | 9.5 | Samples with <= 10 >= 5 bp deletions weren't analyzed |
| PACA-CA::SP117653 | Pancreas | PACA-CA | 261 | 0 | 57 | 8 | 11.5 | Samples with <= 10 >= 5 bp deletions weren't analyzed |
| CLLE-ES::SP13379 | Blood | CLLE-ES | 111 | 0 | 0 | 5 | 9 | Samples with <= 10 >= 5 bp deletions weren't analyzed |
| LGG-US::SP48888 | Brain | LGG-US | 75 | 0 | 0 | 1 | 16 | Samples with <= 10 >= 5 bp deletions weren't analyzed |
| LGG-US::SP48426 | Brain | LGG-US | 23 | 0 | 0 | 0 | NA | Samples with <= 10 >= 5 bp deletions weren't analyzed |
| GACA-CN::SP135411 | Stomach | STAD-US | 286 | 0 | 0 | 3 | 8.5 | Samples with <= 10 >= 5 bp deletions weren't analyzed |
| MALY-DE::SP116672 | Blood | MALY-DE | 124 | 0 | 0 | 1 | 8 | Samples with <= 10 >= 5 bp deletions weren't analyzed |
| PRAD-CA::SP112999 | Prostate | PRAD-US | 86 | 0 | 12 | 4 | 10 | Samples with <= 10 >= 5 bp deletions weren't analyzed |
| PRAD-CA::SP112939 | Prostate | PRAD-US | 177 | 0 | 28 | 8 | 10.5 | Samples with <= 10 >= 5 bp deletions weren't analyzed |
| PAEN-AU::SP102561 | Pancreas | PAEN-AU | 28 | 6 | 5 | 2 | 9 | Samples with <= 10 >= 5 bp deletions weren't analyzed |
| PACA-CA::SP125710 | Pancreas | PACA-CA | 511 | 0 | 95 | 10 | 16 | Samples with <= 10 >= 5 bp deletions weren't analyzed |
| HNSC-US::SP30185 | Head&Neck | HNSC-US | 103 | 20 | 0 | 5 | 11 | Samples with <= 10 >= 5 bp deletions weren't analyzed |
| COAD-US::SP21400 | CRC | COAD-US | 10604 | 0 | 0 | 9 | 8 | Samples with <= 10 >= 5 bp deletions weren't analyzed |
| OV-AU::SP101686 | Ovary | OV-AU | 251 | 25 | 0 | 8 | 10.5 | Samples with <= 10 >= 5 bp deletions weren't analyzed |
| CLLE-ES::SP116750 | Blood | CLLE-ES | 188 | 0 | 0 | 4 | 10.5 | Samples with <= 10 >= 5 bp deletions weren't analyzed |
| PRAD-CA::SP112921 | Prostate | PRAD-US | 185 | 0 | 34 | 10 | 11.5 | Samples with <= 10 >= 5 bp deletions weren't analyzed |
| GACA-CN::SP135188 | Stomach | STAD-US | 202 | 0 | 22 | 4 | 9 | Samples with <= 10 >= 5 bp deletions weren't analyzed |
| BRCA-US::SP5666 | Breast | BRCA-US | 213 | 0 | 58 | 6 | 12 | Samples with <= 10 >= 5 bp deletions weren't analyzed |
| GBM-US::SP27603 | Brain | GBM-US | 381 | 0 | 0 | 9 | 6 | Samples with <= 10 >= 5 bp deletions weren't analyzed |
| LIRI-JP::SP112276 | Liver | LIRI-JP | 205 | 0 | 17 | 6 | 12 | Samples with <= 10 >= 5 bp deletions weren't analyzed |

| | | | | | | | | |
|---|---|---|---|---|---|---|---|---|
| CLLE-ES::SP115125 | Blood | CLLE-ES | 66 | 0 | 0 | 1 | 5.5 | Samples with <= 10 >= 5 bp deletions weren't analyzed |
| LIRI-JP::SP99317 | Liver | LIRI-JP | 39 | 0 | 6 | 4 | 7.5 | Samples with <= 10 >= 5 bp deletions weren't analyzed |
| PACA-CA::SP117722 | Pancreas | PACA-CA | 496 | 0 | 0 | 10 | 14 | Samples with <= 10 >= 5 bp deletions weren't analyzed |
| UCEC-US::SP92723 | Endometrium | UCEC-US | 448 | 0 | 0 | 7 | 10 | Samples with <= 10 >= 5 bp deletions weren't analyzed |
| HNSC-US::SP33837 | Head&Neck | HNSC-US | 239 | 0 | 28 | 6 | 11 | Samples with <= 10 >= 5 bp deletions weren't analyzed |
| PRAD-CA::SP112831 | Prostate | PRAD-US | 214 | 0 | 0 | 9 | 10 | Samples with <= 10 >= 5 bp deletions weren't analyzed |
| MALY-DE::SP116635 | Blood | MALY-DE | 417 | 0 | 0 | 10 | 11 | Samples with <= 10 >= 5 bp deletions weren't analyzed |
| PRAD-US::SP79988 | Prostate | PRAD-US | 64 | 0 | 13 | 5 | 11 | Samples with <= 10 >= 5 bp deletions weren't analyzed |
| PRAD-CA::SP112891 | Prostate | PRAD-US | 260 | 0 | 41 | 10 | 14 | Samples with <= 10 >= 5 bp deletions weren't analyzed |
| MELA-AU::SP124281 | Skin | SKCM-US | 186 | 0 | 0 | 3 | 10 | Samples with <= 10 >= 5 bp deletions weren't analyzed |
| LIRI-JP::SP99321 | Liver | LIRI-JP | 248 | 0 | 0 | 6 | 15 | Samples with <= 10 >= 5 bp deletions weren't analyzed |
| CLLE-ES::SP115089 | Blood | CLLE-ES | 173 | 0 | 0 | 5 | 8.5 | Samples with <= 10 >= 5 bp deletions weren't analyzed |
| MALY-DE::SP59328 | Blood | MALY-DE | 286 | 0 | 0 | 5 | 10 | Samples with <= 10 >= 5 bp deletions weren't analyzed |
| PRAD-US::SP80183 | Prostate | PRAD-US | 185 | 0 | 17 | 8 | 11 | Samples with <= 10 >= 5 bp deletions weren't analyzed |
| THCA-US::SP105759 | Thyroid | THCA-US | 62 | 0 | 9 | 1 | 6.5 | Samples with <= 10 >= 5 bp deletions weren't analyzed |
| LIRI-JP::SP99333 | Liver | LIRI-JP | 253 | 0 | 28 | 10 | 14 | Samples with <= 10 >= 5 bp deletions weren't analyzed |
| LIRI-JP::SP99037 | Liver | LIRI-JP | 142 | 0 | 15 | 3 | 14 | Samples with <= 10 >= 5 bp deletions weren't analyzed |
| MALY-DE::SP116627 | Blood | MALY-DE | 271 | 0 | 0 | 4 | 7 | Samples with <= 10 >= 5 bp deletions weren't analyzed |
| THCA-US::SP86775 | Thyroid | THCA-US | 20 | 0 | 7 | 2 | 7 | Samples with <= 10 >= 5 bp deletions weren't analyzed |
| THCA-US::SP88322 | Thyroid | THCA-US | 93 | 0 | 17 | 5 | 13 | Samples with <= 10 >= 5 bp deletions weren't analyzed |
| PACA-CA::SP77956 | Pancreas | PACA-CA | 5474 | 0 | 0 | 10 | 10 | Samples with <= 10 >= 5 bp deletions weren't analyzed |
| PRAD-CA::SP112897 | Prostate | PRAD-US | 335 | 0 | 29 | 9 | 12 | Samples with <= 10 >= 5 bp deletions weren't analyzed |
| MALY-DE::SP116725 | Blood | MALY-DE | 160 | 0 | 0 | 2 | 8 | Samples with <= 10 >= 5 bp deletions weren't analyzed |
| PRAD-CA::SP112885 | Prostate | PRAD-US | 209 | 0 | 0 | 6 | 11.5 | Samples with <= 10 >= 5 bp deletions weren't analyzed |
| BRCA-US::SP2781 | Breast | BRCA-US | 177 | 19 | 12 | 9 | 12.5 | Samples with <= 10 >= 5 bp deletions weren't analyzed |
| CLLE-ES::SP13297 | Blood | CLLE-ES | 123 | 0 | 0 | 4 | 16 | Samples with <= 10 >= 5 bp deletions weren't analyzed |
| GACA-CN::SP135194 | Stomach | STAD-US | 367 | 0 | 0 | 7 | 8 | Samples with <= 10 >= 5 bp deletions weren't analyzed |
| LIRI-JP::SP99345 | Liver | LIRI-JP | 56 | 0 | 14 | 5 | 13.5 | Samples with <= 10 >= 5 bp deletions weren't analyzed |
| PAEN-AU::SP102489 | Pancreas | PAEN-AU | 53 | 0 | 20 | 8 | 12 | Samples with <= 10 >= 5 bp deletions weren't analyzed |
| BLCA-US::SP1712 | Bladder | BLCA-US | 310 | 0 | 0 | 5 | 12 | Samples with <= 10 >= 5 bp deletions weren't analyzed |
| CLLE-ES::SP13339 | Blood | CLLE-ES | 61 | 0 | 0 | 4 | 7 | Samples with <= 10 >= 5 bp deletions weren't analyzed |
| PACA-CA::SP125715 | Pancreas | PACA-CA | 129 | 0 | 27 | 10 | 17 | Samples with <= 10 >= 5 bp deletions weren't analyzed |
| THCA-US::SP85582 | Thyroid | THCA-US | 87 | 0 | 10 | 3 | 8 | Samples with <= 10 >= 5 bp deletions weren't analyzed |
| PACA-CA::SP116962 | Pancreas | PACA-CA | 295 | 0 | 69 | 8 | 11 | Samples with <= 10 >= 5 bp deletions weren't analyzed |
| MELA-AU::SP124286 | Skin | SKCM-US | 114 | 0 | 19 | 7 | 13 | Samples with <= 10 >= 5 bp deletions weren't analyzed |
| LIHC-US::SP98060 | Liver | LIHC-US | 90 | 0 | 11 | 4 | 7.5 | Samples with <= 10 >= 5 bp deletions weren't analyzed |
| CLLE-ES::SP13671 | Blood | CLLE-ES | 138 | 0 | 0 | 4 | 6 | Samples with <= 10 >= 5 bp deletions weren't analyzed |
| PACA-AU::SP69812 | Pancreas | PACA-AU | 268 | 0 | 0 | 9 | 13 | Samples with <= 10 >= 5 bp deletions weren't analyzed |
| PRAD-CA::SP112821 | Prostate | PRAD-US | 212 | 0 | 0 | 3 | 10 | Samples with <= 10 >= 5 bp deletions weren't analyzed |
| PRAD-CA::SP112813 | Prostate | PRAD-US | 107 | 0 | 11 | 5 | 12 | Samples with <= 10 >= 5 bp deletions weren't analyzed |
| MALY-DE::SP59292 | Blood | MALY-DE | 283 | 0 | 0 | 5 | 10 | Samples with <= 10 >= 5 bp deletions weren't analyzed |
| GACA-CN::SP135258 | Stomach | STAD-US | 71 | 0 | 0 | 2 | 8 | Samples with <= 10 >= 5 bp deletions weren't analyzed |
| MALY-DE::SP59372 | Blood | MALY-DE | 144 | 28 | 20 | 10 | 11 | Samples with <= 10 >= 5 bp deletions weren't analyzed |
| GBM-US::SP26499 | Brain | GBM-US | 252 | 0 | 0 | 7 | 7 | Samples with <= 10 >= 5 bp deletions weren't analyzed |
| CESC-US::SP109953 | Cervix | CESC-US | 409 | 0 | 0 | 6 | 10.5 | Samples with <= 10 >= 5 bp deletions weren't analyzed |
| PRAD-CA::SP112977 | Prostate | PRAD-US | 227 | 0 | 24 | 10 | 11 | Samples with <= 10 >= 5 bp deletions weren't analyzed |
| RECA-EU::SP103467 | Kidney | RECA-EU | 195 | 0 | 21 | 9 | 12 | Samples with <= 10 >= 5 bp deletions weren't analyzed |
| LGG-US::SP48480 | Brain | LGG-US | 313 | 0 | 0 | 5 | 6 | Samples with <= 10 >= 5 bp deletions weren't analyzed |
| COAD-US::SP17430 | CRC | COAD-US | 635 | 0 | 0 | 10 | 13.5 | Samples with <= 10 >= 5 bp deletions weren't analyzed |
| LGG-US::SP48263 | Brain | LGG-US | 83 | 0 | 0 | 4 | 9.5 | Samples with <= 10 >= 5 bp deletions weren't analyzed |

| | | | | | | | | |
|---|---|---|---|---|---|---|---|---|
| MALY-DE::SP116638 | Blood | MALY-DE | 329 | 0 | 0 | 8 | 10.5 | Samples with <= 10 >= 5 bp deletions weren't analyzed |
| CLLE-ES::SP116776 | Blood | CLLE-ES | 88 | 0 | 0 | 0 | 16 | Samples with <= 10 >= 5 bp deletions weren't analyzed |
| CLLE-ES::SP13683 | Blood | CLLE-ES | 95 | 0 | 0 | 0 | 13 | Samples with <= 10 >= 5 bp deletions weren't analyzed |
| MALY-DE::SP59336 | Blood | MALY-DE | 199 | 0 | 0 | 5 | 13 | Samples with <= 10 >= 5 bp deletions weren't analyzed |
| LGG-US::SP47628 | Brain | LGG-US | 61 | 0 | 0 | 3 | 10 | Samples with <= 10 >= 5 bp deletions weren't analyzed |
| THCA-US::SP87446 | Thyroid | THCA-US | 203 | 0 | 56 | 8 | 15 | Samples with <= 10 >= 5 bp deletions weren't analyzed |
| PRAD-CA::SP112959 | Prostate | PRAD-US | 58 | 0 | 0 | 1 | 10 | Samples with <= 10 >= 5 bp deletions weren't analyzed |
| PRAD-CA::SP112931 | Prostate | PRAD-US | 209 | 0 | 21 | 9 | 9 | Samples with <= 10 >= 5 bp deletions weren't analyzed |
| PRAD-CA::SP112865 | Prostate | PRAD-US | 275 | 0 | 0 | 6 | 9 | Samples with <= 10 >= 5 bp deletions weren't analyzed |
| CLLE-ES::SP116758 | Blood | CLLE-ES | 166 | 0 | 0 | 2 | 10 | Samples with <= 10 >= 5 bp deletions weren't analyzed |
| LGG-US::SP47808 | Brain | LGG-US | 213 | 0 | 0 | 3 | 5.5 | Samples with <= 10 >= 5 bp deletions weren't analyzed |
| PAEN-AU::SP117223 | Pancreas | PAEN-AU | 85 | 0 | 12 | 2 | 9 | Samples with <= 10 >= 5 bp deletions weren't analyzed |
| BLCA-US::SP1431 | Bladder | BLCA-US | 398 | 0 | 0 | 7 | 9 | Samples with <= 10 >= 5 bp deletions weren't analyzed |
| PRAD-CA::SP113003 | Prostate | PRAD-US | 185 | 0 | 36 | 8 | 9 | Samples with <= 10 >= 5 bp deletions weren't analyzed |
| CLLE-ES::SP116782 | Blood | CLLE-ES | 227 | 0 | 0 | 1 | 12 | Samples with <= 10 >= 5 bp deletions weren't analyzed |
| MALY-DE::SP116665 | Blood | MALY-DE | 143 | 0 | 0 | 1 | 8 | Samples with <= 10 >= 5 bp deletions weren't analyzed |
| PRAD-CA::SP113001 | Prostate | PRAD-US | 282 | 0 | 0 | 9 | 11 | Samples with <= 10 >= 5 bp deletions weren't analyzed |
| CLLE-ES::SP13317 | Blood | CLLE-ES | 104 | 0 | 0 | 3 | 10.5 | Samples with <= 10 >= 5 bp deletions weren't analyzed |
| PACA-CA::SP125733 | Pancreas | PACA-CA | 438 | 0 | 0 | 6 | 11 | Samples with <= 10 >= 5 bp deletions weren't analyzed |
| THCA-US::SP87337 | Thyroid | THCA-US | 40 | 0 | 3 | 3 | 13 | Samples with <= 10 >= 5 bp deletions weren't analyzed |
| CLLE-ES::SP115128 | Blood | CLLE-ES | 97 | 0 | 0 | 2 | 7.5 | Samples with <= 10 >= 5 bp deletions weren't analyzed |
| CLLE-ES::SP116754 | Blood | CLLE-ES | 123 | 0 | 0 | 3 | 10 | Samples with <= 10 >= 5 bp deletions weren't analyzed |
| CLLE-ES::SP116766 | Blood | CLLE-ES | 357 | 0 | 0 | 3 | 6.5 | Samples with <= 10 >= 5 bp deletions weren't analyzed |
| LIRI-JP::SP99097 | Liver | LIRI-JP | 240 | 0 | 0 | 10 | 12 | Samples with <= 10 >= 5 bp deletions weren't analyzed |
| PACA-CA::SP78502 | Pancreas | PACA-CA | 135 | 0 | 19 | 8 | 16 | Samples with <= 10 >= 5 bp deletions weren't analyzed |
| CESC-US::SP96540 | Cervix | CESC-US | 229 | 0 | 0 | 1 | 10 | Samples with <= 10 >= 5 bp deletions weren't analyzed |
| PRAD-US::SP80217 | Prostate | PRAD-US | 9 | 0 | 0 | 0 | NA | Samples with <= 10 >= 5 bp deletions weren't analyzed |
| PAEN-AU::SP106750 | Pancreas | PAEN-AU | 58 | 0 | 7 | 0 | 11 | Samples with <= 10 >= 5 bp deletions weren't analyzed |
| GBM-US::SP23622 | Brain | GBM-US | 393 | 0 | 0 | 8 | 5 | Samples with <= 10 >= 5 bp deletions weren't analyzed |
| LIRI-JP::SP99113 | Liver | LIRI-JP | 79 | 0 | 12 | 4 | 10 | Samples with <= 10 >= 5 bp deletions weren't analyzed |
| GBM-US::SP27939 | Brain | GBM-US | 317 | 0 | 0 | 7 | 5 | Samples with <= 10 >= 5 bp deletions weren't analyzed |
| BRCA-US::SP12856 | Breast | BRCA-US | 230 | 0 | 50 | 9 | 10.5 | Samples with <= 10 >= 5 bp deletions weren't analyzed |
| CLLE-ES::SP116748 | Blood | CLLE-ES | 232 | 0 | 0 | 7 | 10 | Samples with <= 10 >= 5 bp deletions weren't analyzed |
| OV-AU::SP101658 | Ovary | OV-AU | 111 | 0 | 16 | 4 | 10 | Samples with <= 10 >= 5 bp deletions weren't analyzed |
| CLLE-ES::SP115070 | Blood | CLLE-ES | 97 | 0 | 0 | 5 | 13 | Samples with <= 10 >= 5 bp deletions weren't analyzed |
| LUAD-US::SP55004 | Lung | LUAD-US | 21 | 0 | 9 | 3 | 17.5 | Samples with <= 10 >= 5 bp deletions weren't analyzed |
| PAEN-AU::SP102511 | Pancreas | PAEN-AU | 185 | 29 | 0 | 9 | 9.5 | Samples with <= 10 >= 5 bp deletions weren't analyzed |
| MALY-DE::SP116604 | Blood | MALY-DE | 371 | 0 | 0 | 3 | 8.5 | Samples with <= 10 >= 5 bp deletions weren't analyzed |
| CLLE-ES::SP115131 | Blood | CLLE-ES | 95 | 0 | 0 | 1 | 18.5 | Samples with <= 10 >= 5 bp deletions weren't analyzed |
| LAML-KR::SP127706 | Blood | LAML-US | 157 | 0 | 0 | 7 | 11 | Samples with <= 10 >= 5 bp deletions weren't analyzed |
| PRAD-US::SP80160 | Prostate | PRAD-US | 17 | 0 | 0 | 0 | 15 | Samples with <= 10 >= 5 bp deletions weren't analyzed |
| ORCA-IN::SP117681 | Head&Neck | ORCA-IN | 143 | 0 | 14 | 4 | 13.5 | Samples with <= 10 >= 5 bp deletions weren't analyzed |
| CESC-US::SP114032 | Cervix | CESC-US | 317 | 0 | 0 | 10 | 8 | Samples with <= 10 >= 5 bp deletions weren't analyzed |
| UCEC-US::SP89519 | Endometrium | UCEC-US | 412 | 0 | 0 | 8 | 8 | Samples with <= 10 >= 5 bp deletions weren't analyzed |
| CLLE-ES::SP115097 | Blood | CLLE-ES | 128 | 0 | 0 | 9 | 9.5 | Samples with <= 10 >= 5 bp deletions weren't analyzed |
| OV-US::SP66687 | Ovary | OV-US | 453 | 0 | 0 | 9 | 11 | Samples with <= 10 >= 5 bp deletions weren't analyzed |
| MALY-DE::SP59376 | Blood | MALY-DE | 147 | 0 | 0 | 7 | 19 | Samples with <= 10 >= 5 bp deletions weren't analyzed |
| GACA-CN::SP135293 | Stomach | STAD-US | 5 | 3 | 0 | 1 | 18 | Samples with <= 10 >= 5 bp deletions weren't analyzed |
| LAML-KR::SP108999 | Blood | LAML-US | 113 | 0 | 0 | 7 | 9 | Samples with <= 10 >= 5 bp deletions weren't analyzed |
| GBM-US::SP23754 | Brain | GBM-US | 130 | 0 | 0 | 3 | 6 | Samples with <= 10 >= 5 bp deletions weren't analyzed |

| | | | | | | | | |
|---|---|---|---|---|---|---|---|---|
| BRCA-US::SP5820 | Breast | BRCA-US | 349 | 0 | 72 | 6 | 7 | Samples with <= 10 >= 5 bp deletions weren't analyzed |
| PRAD-CA::SP112853 | Prostate | PRAD-US | 239 | 0 | 0 | 7 | 11.5 | Samples with <= 10 >= 5 bp deletions weren't analyzed |
| CLLE-ES::SP115060 | Blood | CLLE-ES | 155 | 0 | 0 | 5 | 12.5 | Samples with <= 10 >= 5 bp deletions weren't analyzed |
| CLLE-ES::SP13594 | Blood | CLLE-ES | 161 | 0 | 0 | 4 | 12 | Samples with <= 10 >= 5 bp deletions weren't analyzed |
| GBM-US::SP29697 | Brain | GBM-US | 314 | 0 | 0 | 3 | 5 | Samples with <= 10 >= 5 bp deletions weren't analyzed |
| CLLE-ES::SP96834 | Blood | CLLE-ES | 138 | 0 | 0 | 7 | 13 | Samples with <= 10 >= 5 bp deletions weren't analyzed |
| CLLE-ES::SP115057 | Blood | CLLE-ES | 155 | 0 | 0 | 2 | 8 | Samples with <= 10 >= 5 bp deletions weren't analyzed |
| PRAD-CA::SP113005 | Prostate | PRAD-US | 99 | 13 | 0 | 4 | 6.5 | Samples with <= 10 >= 5 bp deletions weren't analyzed |
| MALY-DE::SP59272 | Blood | MALY-DE | 283 | 0 | 0 | 5 | 10 | Samples with <= 10 >= 5 bp deletions weren't analyzed |
| PRAD-US::SP80244 | Prostate | PRAD-US | 89 | 0 | 14 | 5 | 9.5 | Samples with <= 10 >= 5 bp deletions weren't analyzed |
| PRAD-CA::SP112957 | Prostate | PRAD-US | 269 | 0 | 30 | 10 | 10 | Samples with <= 10 >= 5 bp deletions weren't analyzed |
| CLLE-ES::SP116768 | Blood | CLLE-ES | 142 | 0 | 0 | 0 | 9 | Samples with <= 10 >= 5 bp deletions weren't analyzed |
| MALY-DE::SP116608 | Blood | MALY-DE | 132 | 0 | 0 | 2 | 6 | Samples with <= 10 >= 5 bp deletions weren't analyzed |
| PAEN-AU::SP117595 | Pancreas | PAEN-AU | 85 | 0 | 9 | 6 | 16.5 | Samples with <= 10 >= 5 bp deletions weren't analyzed |
| PRAD-CA::SP112783 | Prostate | PRAD-US | 293 | 0 | 32 | 10 | 7 | Samples with <= 10 >= 5 bp deletions weren't analyzed |
| MALY-DE::SP59396 | Blood | MALY-DE | 256 | 0 | 0 | 1 | 10 | Samples with <= 10 >= 5 bp deletions weren't analyzed |
| THCA-US::SP86130 | Thyroid | THCA-US | 73 | 0 | 13 | 4 | 8 | Samples with <= 10 >= 5 bp deletions weren't analyzed |
| CLLE-ES::SP13442 | Blood | CLLE-ES | 198 | 0 | 0 | 4 | 12 | Samples with <= 10 >= 5 bp deletions weren't analyzed |
| THCA-US::SP86425 | Thyroid | THCA-US | 138 | 0 | 28 | 9 | 11 | Samples with <= 10 >= 5 bp deletions weren't analyzed |
| CLLE-ES::SP13325 | Blood | CLLE-ES | 61 | 0 | 0 | 2 | 16.5 | Samples with <= 10 >= 5 bp deletions weren't analyzed |
| LAML-KR::SP109007 | Blood | LAML-US | 76 | 0 | 0 | 1 | 6 | Samples with <= 10 >= 5 bp deletions weren't analyzed |
| SKCM-US::SP82435 | Skin | SKCM-US | 256 | 0 | 35 | 10 | 12 | Samples with <= 10 >= 5 bp deletions weren't analyzed |
| PACA-CA::SP78533 | Pancreas | PACA-CA | 189 | 43 | 46 | 10 | 9 | Samples with <= 10 >= 5 bp deletions weren't analyzed |
| THCA-US::SP85495 | Thyroid | THCA-US | 31 | 0 | 3 | 0 | 13.5 | Samples with <= 10 >= 5 bp deletions weren't analyzed |
| PRAD-CA::SP112943 | Prostate | PRAD-US | 157 | 0 | 15 | 9 | 14 | Samples with <= 10 >= 5 bp deletions weren't analyzed |
| MALY-DE::SP116688 | Blood | MALY-DE | 317 | 0 | 0 | 3 | 8.5 | Samples with <= 10 >= 5 bp deletions weren't analyzed |
| COAD-US::SP17016 | CRC | COAD-US | 1545 | 0 | 0 | 9 | 10 | Samples with <= 10 >= 5 bp deletions weren't analyzed |
| PRAD-CA::SP112861 | Prostate | PRAD-US | 230 | 0 | 0 | 3 | 11 | Samples with <= 10 >= 5 bp deletions weren't analyzed |
| MALY-DE::SP59392 | Blood | MALY-DE | 221 | 0 | 0 | 1 | 8 | Samples with <= 10 >= 5 bp deletions weren't analyzed |
| PRAD-CA::SP112917 | Prostate | PRAD-US | 263 | 0 | 0 | 9 | 13 | Samples with <= 10 >= 5 bp deletions weren't analyzed |
| PACA-AU::SP70046 | Pancreas | PACA-AU | 311 | 0 | 64 | 6 | 9 | Samples with <= 10 >= 5 bp deletions weren't analyzed |
| PAEN-AU::SP117294 | Pancreas | PAEN-AU | 43 | 7 | 0 | 4 | 8.5 | Samples with <= 10 >= 5 bp deletions weren't analyzed |
| PAEN-AU::SP102499 | Pancreas | PAEN-AU | 123 | 0 | 27 | 9 | 10 | Samples with <= 10 >= 5 bp deletions weren't analyzed |
| PAEN-AU::SP117951 | Pancreas | PAEN-AU | 39 | 0 | 8 | 4 | 19.5 | Samples with <= 10 >= 5 bp deletions weren't analyzed |
| GBM-US::SP29331 | Brain | GBM-US | 456 | 0 | 0 | 10 | 5 | Samples with <= 10 >= 5 bp deletions weren't analyzed |
| LIRI-JP::SP99169 | Liver | LIRI-JP | 167 | 0 | 21 | 7 | 19 | Samples with <= 10 >= 5 bp deletions weren't analyzed |
| PRAD-US::SP79998 | Prostate | PRAD-US | 7 | 0 | 0 | 0 | NA | Samples with <= 10 >= 5 bp deletions weren't analyzed |
| COAD-US::SP17329 | CRC | COAD-US | 888 | 0 | 0 | 6 | 10 | Samples with <= 10 >= 5 bp deletions weren't analyzed |
| PAEN-AU::SP117085 | Pancreas | PAEN-AU | 48 | 10 | 11 | 5 | 8 | Samples with <= 10 >= 5 bp deletions weren't analyzed |
| PACA-CA::SP116985 | Pancreas | PACA-CA | 504 | 0 | 0 | 10 | 9 | Samples with <= 10 >= 5 bp deletions weren't analyzed |
| GBM-US::SP26475 | Brain | GBM-US | 410 | 0 | 0 | 10 | 6 | Samples with <= 10 >= 5 bp deletions weren't analyzed |
| PACA-CA::SP125769 | Pancreas | PACA-CA | 280 | 0 | 70 | 8 | 12 | Samples with <= 10 >= 5 bp deletions weren't analyzed |
| THCA-US::SP85787 | Thyroid | THCA-US | 43 | 0 | 4 | 2 | 10 | Samples with <= 10 >= 5 bp deletions weren't analyzed |
| CESC-US::SP107650 | Cervix | CESC-US | 356 | 0 | 0 | 9 | 12 | Samples with <= 10 >= 5 bp deletions weren't analyzed |
| LIHC-US::SP49449 | Liver | LIHC-US | 403 | 0 | 0 | 9 | 13 | Samples with <= 10 >= 5 bp deletions weren't analyzed |
| MALY-DE::SP59348 | Blood | MALY-DE | 393 | 0 | 0 | 10 | 11 | Samples with <= 10 >= 5 bp deletions weren't analyzed |
| PACA-AU::SP76353 | Pancreas | PACA-AU | 258 | 0 | 66 | 3 | 15 | Samples with <= 10 >= 5 bp deletions weren't analyzed |
| PACA-AU::SP69587 | Pancreas | PACA-AU | 217 | 0 | 0 | 5 | 16 | Samples with <= 10 >= 5 bp deletions weren't analyzed |
| PRAD-CA::SP112811 | Prostate | PRAD-US | 293 | 0 | 0 | 10 | 12 | Samples with <= 10 >= 5 bp deletions weren't analyzed |
| ORCA-IN::SP117408 | Head&Neck | ORCA-IN | 112 | 13 | 17 | 6 | 9 | Samples with <= 10 >= 5 bp deletions weren't analyzed |

| | | | | | | | | |
|---|---|---|---|---|---|---|---|---|
| CLLE-ES::SP116736 | Blood | CLLE-ES | 106 | 0 | 0 | 1 | 9 | Samples with <= 10 >= 5 bp deletions weren't analyzed |
| PACA-CA::SP125730 | Pancreas | PACA-CA | 255 | 0 | 0 | 5 | 11 | Samples with <= 10 >= 5 bp deletions weren't analyzed |
| GACA-CN::SP135415 | Stomach | STAD-US | 199 | 0 | 28 | 7 | 9 | Samples with <= 10 >= 5 bp deletions weren't analyzed |
| MALY-DE::SP59340 | Blood | MALY-DE | 367 | 0 | 0 | 8 | 16 | Samples with <= 10 >= 5 bp deletions weren't analyzed |
| MALY-DE::SP59380 | Blood | MALY-DE | 130 | 0 | 14 | 9 | 9 | Samples with <= 10 >= 5 bp deletions weren't analyzed |
| PAEN-AU::SP106669 | Pancreas | PAEN-AU | 111 | 0 | 15 | 6 | 10 | Samples with <= 10 >= 5 bp deletions weren't analyzed |
| CLLE-ES::SP13619 | Blood | CLLE-ES | 129 | 0 | 0 | 6 | 6 | Samples with <= 10 >= 5 bp deletions weren't analyzed |
| PRAD-CA::SP112909 | Prostate | PRAD-US | 306 | 0 | 0 | 10 | 10 | Samples with <= 10 >= 5 bp deletions weren't analyzed |
| HNSC-US::SP32222 | Head&Neck | HNSC-US | 5 | 0 | 0 | 0 | NA | Samples with <= 10 >= 5 bp deletions weren't analyzed |
| CLLE-ES::SP115111 | Blood | CLLE-ES | 289 | 0 | 0 | 5 | 12 | Samples with <= 10 >= 5 bp deletions weren't analyzed |
| PRAD-CA::SP112781 | Prostate | PRAD-US | 265 | 0 | 23 | 4 | 20.5 | Samples with <= 10 >= 5 bp deletions weren't analyzed |
| LIRI-JP::SP50141 | Liver | LIRI-JP | 32 | 0 | 10 | 3 | 15 | Samples with <= 10 >= 5 bp deletions weren't analyzed |
| PACA-CA::SP77848 | Pancreas | PACA-CA | 126 | 0 | 0 | 2 | 15 | Samples with <= 10 >= 5 bp deletions weren't analyzed |
| LGG-US::SP48073 | Brain | LGG-US | 155 | 0 | 0 | 4 | 5 | Samples with <= 10 >= 5 bp deletions weren't analyzed |
| GBM-US::SP28275 | Brain | GBM-US | 289 | 0 | 0 | 1 | 5 | Samples with <= 10 >= 5 bp deletions weren't analyzed |
| CLLE-ES::SP116760 | Blood | CLLE-ES | 122 | 0 | 0 | 3 | 8 | Samples with <= 10 >= 5 bp deletions weren't analyzed |
| GACA-CN::SP135310 | Stomach | STAD-US | 2 | 0 | 0 | 0 | NA | Samples with <= 10 >= 5 bp deletions weren't analyzed |
| CLLE-ES::SP13311 | Blood | CLLE-ES | 91 | 0 | 0 | 5 | 9.5 | Samples with <= 10 >= 5 bp deletions weren't analyzed |
| CLLE-ES::SP13509 | Blood | CLLE-ES | 96 | 0 | 0 | 1 | 14 | Samples with <= 10 >= 5 bp deletions weren't analyzed |
| BLCA-US::SP967 | Bladder | BLCA-US | 294 | 0 | 0 | 6 | 8.5 | Samples with <= 10 >= 5 bp deletions weren't analyzed |
| LGG-US::SP48008 | Brain | LGG-US | 586 | 0 | 0 | 9 | 5 | Samples with <= 10 >= 5 bp deletions weren't analyzed |
| OV-US::SP61343 | Ovary | OV-US | 276 | 0 | 44 | 10 | 10 | Samples with <= 10 >= 5 bp deletions weren't analyzed |
| THCA-US::SP86836 | Thyroid | THCA-US | 18 | 0 | 0 | 0 | 32 | Samples with <= 10 >= 5 bp deletions weren't analyzed |
| CLLE-ES::SP13539 | Blood | CLLE-ES | 80 | 0 | 0 | 1 | 7 | Samples with <= 10 >= 5 bp deletions weren't analyzed |
| CLLE-ES::SP13624 | Blood | CLLE-ES | 109 | 0 | 0 | 3 | 5 | Samples with <= 10 >= 5 bp deletions weren't analyzed |
| MALY-DE::SP59428 | Blood | MALY-DE | 176 | 0 | 22 | 3 | 9 | Samples with <= 10 >= 5 bp deletions weren't analyzed |
| READ-US::SP81440 | CRC | READ-US | 488 | 0 | 0 | 8 | 7.5 | Samples with <= 10 >= 5 bp deletions weren't analyzed |
| PAEN-AU::SP102581 | Pancreas | PAEN-AU | 31 | 0 | 0 | 1 | 7 | Samples with <= 10 >= 5 bp deletions weren't analyzed |
| PACA-AU::SP70226 | Pancreas | PACA-AU | 406 | 82 | 0 | 8 | 6.5 | Samples with <= 10 >= 5 bp deletions weren't analyzed |
| PACA-CA::SP125711 | Pancreas | PACA-CA | 584 | 0 | 146 | 8 | 16 | Samples with <= 10 >= 5 bp deletions weren't analyzed |
| THCA-US::SP89245 | Thyroid | THCA-US | 32 | 0 | 2 | 0 | 28 | Samples with <= 10 >= 5 bp deletions weren't analyzed |
| PRAD-CA::SP112993 | Prostate | PRAD-US | 302 | 0 | 34 | 9 | 13 | Samples with <= 10 >= 5 bp deletions weren't analyzed |
| GBM-US::SP28581 | Brain | GBM-US | 423 | 0 | 0 | 10 | 5.5 | Samples with <= 10 >= 5 bp deletions weren't analyzed |
| PRAD-CA::SP112827 | Prostate | PRAD-US | 243 | 0 | 36 | 9 | 11 | Samples with <= 10 >= 5 bp deletions weren't analyzed |
| CLLE-ES::SP116746 | Blood | CLLE-ES | 120 | 0 | 0 | 4 | 10 | Samples with <= 10 >= 5 bp deletions weren't analyzed |
| PRAD-CA::SP112855 | Prostate | PRAD-US | 183 | 0 | 17 | 7 | 14.5 | Samples with <= 10 >= 5 bp deletions weren't analyzed |
| MALY-DE::SP116624 | Blood | MALY-DE | 582 | 0 | 0 | 5 | 9 | Samples with <= 10 >= 5 bp deletions weren't analyzed |
| PRAD-US::SP80216 | Prostate | PRAD-US | 95 | 8 | 14 | 9 | 13 | Samples with <= 10 >= 5 bp deletions weren't analyzed |
| OV-AU::SP101572 | Ovary | OV-AU | 167 | 0 | 33 | 10 | 15.5 | Samples with <= 10 >= 5 bp deletions weren't analyzed |
| PAEN-AU::SP106718 | Pancreas | PAEN-AU | 191 | 0 | 27 | 3 | 6.5 | Samples with <= 10 >= 5 bp deletions weren't analyzed |
| LGG-US::SP48534 | Brain | LGG-US | 247 | 0 | 0 | 7 | 5 | Samples with <= 10 >= 5 bp deletions weren't analyzed |
| READ-US::SP80950 | CRC | READ-US | 806 | 0 | 0 | 8 | 10 | Samples with <= 10 >= 5 bp deletions weren't analyzed |
| CLLE-ES::SP13490 | Blood | CLLE-ES | 133 | 0 | 0 | 1 | 12 | Samples with <= 10 >= 5 bp deletions weren't analyzed |
| GBM-US::SP23639 | Brain | GBM-US | 473 | 0 | 0 | 9 | 5.5 | Samples with <= 10 >= 5 bp deletions weren't analyzed |
| MALY-DE::SP59432 | Blood | MALY-DE | 287 | 0 | 0 | 8 | 11 | Samples with <= 10 >= 5 bp deletions weren't analyzed |
| GACA-CN::SP135323 | Stomach | STAD-US | 149 | 0 | 25 | 7 | 7.5 | Samples with <= 10 >= 5 bp deletions weren't analyzed |
| CLLE-ES::SP116774 | Blood | CLLE-ES | 99 | 0 | 0 | 0 | 27 | Samples with <= 10 >= 5 bp deletions weren't analyzed |
| GACA-CN::SP135390 | Stomach | STAD-US | 45 | 0 | 5 | 2 | 7 | Samples with <= 10 >= 5 bp deletions weren't analyzed |
| LUAD-US::SP55309 | Lung | LUAD-US | 69 | 9 | 6 | 4 | 10.5 | Samples with <= 10 >= 5 bp deletions weren't analyzed |
| CLLE-ES::SP115105 | Blood | CLLE-ES | 102 | 0 | 0 | 2 | 5.5 | Samples with <= 10 >= 5 bp deletions weren't analyzed |

| | | | | | | | | |
|---|---|---|---|---|---|---|---|---|
| MALY-DE::SP116679 | Blood | MALY-DE | 104 | 13 | 0 | 1 | 7 | Samples with <= 10 >= 5 bp deletions weren't analyzed |
| THCA-US::SP88757 | Thyroid | THCA-US | 59 | 0 | 11 | 6 | 6.5 | Samples with <= 10 >= 5 bp deletions weren't analyzed |
| LIRI-JP::SP107028 | Liver | LIRI-JP | 134 | 0 | 22 | 10 | 13 | Samples with <= 10 >= 5 bp deletions weren't analyzed |
| MALY-DE::SP59360 | Blood | MALY-DE | 852 | 0 | 0 | 4 | 11 | Samples with <= 10 >= 5 bp deletions weren't analyzed |
| CLLE-ES::SP13278 | Blood | CLLE-ES | 97 | 0 | 0 | 4 | 11 | Samples with <= 10 >= 5 bp deletions weren't analyzed |
| PACA-AU::SP110824 | Pancreas | PACA-AU | 383 | 0 | 105 | 6 | 16 | Samples with <= 10 >= 5 bp deletions weren't analyzed |
| PRAD-CA::SP112833 | Prostate | PRAD-US | 147 | 0 | 16 | 8 | 8 | Samples with <= 10 >= 5 bp deletions weren't analyzed |
| PRAD-US::SP80042 | Prostate | PRAD-US | 122 | 0 | 23 | 8 | 8 | Samples with <= 10 >= 5 bp deletions weren't analyzed |
| LUAD-US::SP52607 | Lung | LUAD-US | 124 | 0 | 13 | 5 | 17 | Samples with <= 10 >= 5 bp deletions weren't analyzed |
| MALY-DE::SP59384 | Blood | MALY-DE | 516 | 0 | 0 | 1 | 5 | Samples with <= 10 >= 5 bp deletions weren't analyzed |
| CLLE-ES::SP116756 | Blood | CLLE-ES | 232 | 0 | 0 | 5 | 10 | Samples with <= 10 >= 5 bp deletions weren't analyzed |
| CLLE-ES::SP13319 | Blood | CLLE-ES | 38 | 0 | 0 | 0 | NA | Samples with <= 10 >= 5 bp deletions weren't analyzed |
| MALY-DE::SP59332 | Blood | MALY-DE | 221 | 0 | 0 | 4 | 15.5 | Samples with <= 10 >= 5 bp deletions weren't analyzed |
| LIRI-JP::SP99221 | Liver | LIRI-JP | 48 | 0 | 0 | 1 | 11 | Samples with <= 10 >= 5 bp deletions weren't analyzed |
| MELA-AU::SP124344 | Skin | SKCM-US | 227 | 0 | 34 | 7 | 11 | Samples with <= 10 >= 5 bp deletions weren't analyzed |
| PACA-CA::SP117037 | Pancreas | PACA-CA | 405 | 0 | 80 | 8 | 13 | Samples with <= 10 >= 5 bp deletions weren't analyzed |
| CLLE-ES::SP13280 | Blood | CLLE-ES | 140 | 0 | 0 | 4 | 11 | Samples with <= 10 >= 5 bp deletions weren't analyzed |
| MALY-DE::SP116616 | Blood | MALY-DE | 761 | 0 | 0 | 8 | 12 | Samples with <= 10 >= 5 bp deletions weren't analyzed |
| MALY-DE::SP124981 | Blood | MALY-DE | 477 | 0 | 0 | 9 | 12.5 | Samples with <= 10 >= 5 bp deletions weren't analyzed |
| COAD-US::SP96114 | CRC | COAD-US | 1462 | 0 | 0 | 10 | 9 | Samples with <= 10 >= 5 bp deletions weren't analyzed |
| PRAD-CA::SP112777 | Prostate | PRAD-US | 603 | 0 | 0 | 5 | 11 | Samples with <= 10 >= 5 bp deletions weren't analyzed |
| PAEN-AU::SP117331 | Pancreas | PAEN-AU | 227 | 0 | 28 | 10 | 8.5 | Samples with <= 10 >= 5 bp deletions weren't analyzed |
| BRCA-US::SP6115 | Breast | BRCA-US | 244 | 0 | 66 | 10 | 10.5 | Samples with <= 10 >= 5 bp deletions weren't analyzed |
| THCA-US::SP85733 | Thyroid | THCA-US | 100 | 0 | 39 | 5 | 7.5 | Samples with <= 10 >= 5 bp deletions weren't analyzed |
| PACA-CA::SP125795 | Pancreas | PACA-CA | 437 | 0 | 110 | 7 | 13 | Samples with <= 10 >= 5 bp deletions weren't analyzed |
| PACA-AU::SP108065 | Pancreas | PACA-AU | 495 | 0 | 0 | 8 | 9 | Samples with <= 10 >= 5 bp deletions weren't analyzed |
| MALY-DE::SP59420 | Blood | MALY-DE | 237 | 0 | 0 | 7 | 16 | Samples with <= 10 >= 5 bp deletions weren't analyzed |
| OV-US::SP60610 | Ovary | OV-US | 219 | 0 | 34 | 10 | 14 | Samples with <= 10 >= 5 bp deletions weren't analyzed |
| CLLE-ES::SP13363 | Blood | CLLE-ES | 86 | 0 | 0 | 3 | 10 | Samples with <= 10 >= 5 bp deletions weren't analyzed |
| MELA-AU::SP124348 | Skin | SKCM-US | 216 | 0 | 37 | 7 | 9 | Samples with <= 10 >= 5 bp deletions weren't analyzed |
| LINC-JP::SP98863 | Liver | LIHC-US | 230 | 0 | 39 | 10 | 9 | Samples with <= 10 >= 5 bp deletions weren't analyzed |
| GACA-CN::SP135209 | Stomach | STAD-US | 76 | 0 | 0 | 0 | 5 | Samples with <= 10 >= 5 bp deletions weren't analyzed |
| STAD-US::SP85222 | Stomach | STAD-US | 320 | 0 | 0 | 5 | 13 | Samples with <= 10 >= 5 bp deletions weren't analyzed |
| CLLE-ES::SP13347 | Blood | CLLE-ES | 112 | 0 | 0 | 5 | 10 | Samples with <= 10 >= 5 bp deletions weren't analyzed |
| CLLE-ES::SP116730 | Blood | CLLE-ES | 170 | 0 | 0 | 3 | 9.5 | Samples with <= 10 >= 5 bp deletions weren't analyzed |
| BRCA-US::SP2731 | Breast | BRCA-US | 193 | 0 | 51 | 10 | 13 | Samples with <= 10 >= 5 bp deletions weren't analyzed |
| GBM-US::SP28041 | Brain | GBM-US | 538 | 0 | 0 | 5 | 5 | Samples with <= 10 >= 5 bp deletions weren't analyzed |
| PACA-CA::SP125703 | Pancreas | PACA-CA | 510 | 0 | 0 | 8 | 16 | Samples with <= 10 >= 5 bp deletions weren't analyzed |
| LIRI-JP::SP50175 | Liver | LIRI-JP | 66 | 0 | 14 | 3 | 14 | Samples with <= 10 >= 5 bp deletions weren't analyzed |
| CLLE-ES::SP13365 | Blood | CLLE-ES | 79 | 0 | 0 | 0 | 7 | Samples with <= 10 >= 5 bp deletions weren't analyzed |
| MALY-DE::SP59308 | Blood | MALY-DE | 359 | 0 | 0 | 5 | 10 | Samples with <= 10 >= 5 bp deletions weren't analyzed |
| MALY-DE::SP59288 | Blood | MALY-DE | 865 | 0 | 0 | 10 | 9 | Samples with <= 10 >= 5 bp deletions weren't analyzed |
| PRAD-CA::SP112877 | Prostate | PRAD-US | 21 | 0 | 0 | 0 | 8 | Samples with <= 10 >= 5 bp deletions weren't analyzed |
| PRAD-CA::SP112871 | Prostate | PRAD-US | 217 | 0 | 22 | 10 | 9.5 | Samples with <= 10 >= 5 bp deletions weren't analyzed |
| PACA-AU::SP71576 | Pancreas | PACA-AU | 333 | 0 | 0 | 6 | 12 | Samples with <= 10 >= 5 bp deletions weren't analyzed |
| CLLE-ES::SP115099 | Blood | CLLE-ES | 167 | 0 | 0 | 3 | 12 | Samples with <= 10 >= 5 bp deletions weren't analyzed |
| CLLE-ES::SP116764 | Blood | CLLE-ES | 93 | 0 | 0 | 4 | 9 | Samples with <= 10 >= 5 bp deletions weren't analyzed |
| RECA-EU::SP103555 | Kidney | RECA-EU | 88 | 0 | 12 | 4 | 7 | Samples with <= 10 >= 5 bp deletions weren't analyzed |
| PACA-CA::SP117261 | Pancreas | PACA-CA | 272 | 0 | 59 | 8 | 13.5 | Samples with <= 10 >= 5 bp deletions weren't analyzed |
| MALY-DE::SP116622 | Blood | MALY-DE | 214 | 0 | 0 | 6 | 9 | Samples with <= 10 >= 5 bp deletions weren't analyzed |

| | | | | | | | | |
|---|---|---|---|---|---|---|---|---|
| COAD-US::SP19582 | CRC | COAD-US | 586 | 0 | 0 | 7 | 14.5 | Samples with <= 10 >= 5 bp deletions weren't analyzed |
| BRCA-UK::SP2143 | Breast | BRCA-US | 113 | 0 | 27 | 5 | 10 | Samples with <= 10 >= 5 bp deletions weren't analyzed |
| BRCA-UK::SP2150 | Breast | BRCA-US | 141 | 24 | 0 | 1 | 13 | Samples with <= 10 >= 5 bp deletions weren't analyzed |
| BRCA-UK::SP2152 | Breast | BRCA-US | 104 | 0 | 21 | 3 | 11 | Samples with <= 10 >= 5 bp deletions weren't analyzed |
| BRCA-UK::SP2159 | Breast | BRCA-US | 160 | 0 | 31 | 6 | 15.5 | Samples with <= 10 >= 5 bp deletions weren't analyzed |
| BRCA-UK::SP2151 | Breast | BRCA-US | 90 | 16 | 21 | 10 | 12 | Samples with <= 10 >= 5 bp deletions weren't analyzed |
| BRCA-UK::SP2157 | Breast | BRCA-US | 54 | 0 | 20 | 5 | 8 | Samples with <= 10 >= 5 bp deletions weren't analyzed |
| PRAD-US::SP80014 | Prostate | PRAD-US | 67 | 0 | 0 | 1 | 8 | Samples with <= 10 >= 5 bp deletions weren't analyzed |
| PACA-AU::SP74896 | Pancreas | PACA-AU | 460 | 0 | 128 | 4 | 13 | Samples with <= 10 >= 5 bp deletions weren't analyzed |
| CLLE-ES::SP116772 | Blood | CLLE-ES | 116 | 0 | 0 | 4 | 8 | Samples with <= 10 >= 5 bp deletions weren't analyzed |
| MALY-DE::SP59316 | Blood | MALY-DE | 323 | 0 | 0 | 8 | 8 | Samples with <= 10 >= 5 bp deletions weren't analyzed |
| CESC-US::SP114016 | Cervix | CESC-US | 317 | 53 | 0 | 8 | 8.5 | Samples with <= 10 >= 5 bp deletions weren't analyzed |
| PRAD-CA::SP112881 | Prostate | PRAD-US | 213 | 0 | 24 | 6 | 9 | Samples with <= 10 >= 5 bp deletions weren't analyzed |
| LGG-US::SP48010 | Brain | LGG-US | 138 | 0 | 0 | 3 | 5 | Samples with <= 10 >= 5 bp deletions weren't analyzed |
| PACA-CA::SP77972 | Pancreas | PACA-CA | 469 | 0 | 0 | 7 | 8.5 | Samples with <= 10 >= 5 bp deletions weren't analyzed |
| MALY-DE::SP59324 | Blood | MALY-DE | 260 | 0 | 31 | 9 | 9.5 | Samples with <= 10 >= 5 bp deletions weren't analyzed |
| PRAD-UK::SP125579 | Prostate | PRAD-US | 1 | 0 | 0 | 0 | NA | Samples with <= 10 >= 5 bp deletions weren't analyzed |
| PRAD-UK::SP111121 | Prostate | PRAD-US | 29 | 0 | 10 | 0 | 7 | Samples with <= 10 >= 5 bp deletions weren't analyzed |
| PRAD-UK::SP111124 | Prostate | PRAD-US | 149 | 0 | 0 | 5 | 11 | Samples with <= 10 >= 5 bp deletions weren't analyzed |
| PRAD-UK::SP111128 | Prostate | PRAD-US | 182 | 0 | 32 | 7 | 9 | Samples with <= 10 >= 5 bp deletions weren't analyzed |
| PRAD-UK::SP111130 | Prostate | PRAD-US | 144 | 0 | 21 | 9 | 8.5 | Samples with <= 10 >= 5 bp deletions weren't analyzed |
| BRCA-US::SP7456 | Breast | BRCA-US | 262 | 0 | 85 | 10 | 13.5 | Samples with <= 10 >= 5 bp deletions weren't analyzed |
| LAML-KR::SP109003 | Blood | LAML-US | 52 | 0 | 0 | 3 | 12.5 | Samples with <= 10 >= 5 bp deletions weren't analyzed |
| THCA-US::SP85737 | Thyroid | THCA-US | 27 | 0 | 27 | 4 | 7.5 | Samples with <= 10 >= 5 bp deletions weren't analyzed |
| CLLE-ES::SP96888 | Blood | CLLE-ES | 51 | 0 | 0 | 3 | 8.5 | Samples with <= 10 >= 5 bp deletions weren't analyzed |
| PACA-CA::SP77866 | Pancreas | PACA-CA | 72 | 0 | 3 | 2 | 9 | Samples with <= 10 >= 5 bp deletions weren't analyzed |
| CLLE-ES::SP13327 | Blood | CLLE-ES | 146 | 0 | 0 | 6 | 11 | Samples with <= 10 >= 5 bp deletions weren't analyzed |
| PAEN-AU::SP106758 | Pancreas | PAEN-AU | 140 | 0 | 27 | 4 | 8.5 | Samples with <= 10 >= 5 bp deletions weren't analyzed |
| BRCA-UK::SP116351 | Breast | BRCA-US | 267 | 28 | 23 | 10 | 6 | Samples with <= 10 >= 5 bp deletions weren't analyzed |
| THCA-US::SP85647 | Thyroid | THCA-US | 67 | 0 | 11 | 4 | 10 | Samples with <= 10 >= 5 bp deletions weren't analyzed |
| BRCA-UK::SP116347 | Breast | BRCA-US | 157 | 0 | 42 | 7 | 7.5 | Samples with <= 10 >= 5 bp deletions weren't analyzed |
| PRAD-UK::SP106835 | Prostate | PRAD-US | 76 | 0 | 12 | 5 | 8 | Samples with <= 10 >= 5 bp deletions weren't analyzed |
| PRAD-UK::SP106834 | Prostate | PRAD-US | 156 | 0 | 25 | 6 | 11 | Samples with <= 10 >= 5 bp deletions weren't analyzed |
| PRAD-UK::SP106841 | Prostate | PRAD-US | 79 | 15 | 12 | 7 | 9 | Samples with <= 10 >= 5 bp deletions weren't analyzed |
| PRAD-UK::SP106833 | Prostate | PRAD-US | 159 | 0 | 37 | 7 | 10.5 | Samples with <= 10 >= 5 bp deletions weren't analyzed |
| PRAD-UK::SP106839 | Prostate | PRAD-US | 296 | 0 | 29 | 9 | 11.5 | Samples with <= 10 >= 5 bp deletions weren't analyzed |
| PRAD-UK::SP106840 | Prostate | PRAD-US | 170 | 0 | 0 | 7 | 13 | Samples with <= 10 >= 5 bp deletions weren't analyzed |
| PRAD-UK::SP106828 | Prostate | PRAD-US | 134 | 0 | 17 | 6 | 10 | Samples with <= 10 >= 5 bp deletions weren't analyzed |
| PRAD-UK::SP106829 | Prostate | PRAD-US | 316 | 0 | 0 | 10 | 11 | Samples with <= 10 >= 5 bp deletions weren't analyzed |
| GBM-US::SP26649 | Brain | GBM-US | 436 | 0 | 38 | 7 | 5 | Samples with <= 10 >= 5 bp deletions weren't analyzed |
| PRAD-UK::SP106836 | Prostate | PRAD-US | 23 | 0 | 2 | 1 | 10.5 | Samples with <= 10 >= 5 bp deletions weren't analyzed |
| PRAD-UK::SP106837 | Prostate | PRAD-US | 95 | 0 | 15 | 7 | 12 | Samples with <= 10 >= 5 bp deletions weren't analyzed |
| PRAD-CA::SP112951 | Prostate | PRAD-US | 28 | 0 | 3 | 1 | 8.5 | Samples with <= 10 >= 5 bp deletions weren't analyzed |
| PRAD-CA::SP112845 | Prostate | PRAD-US | 247 | 0 | 26 | 7 | 8 | Samples with <= 10 >= 5 bp deletions weren't analyzed |
| CLLE-ES::SP116740 | Blood | CLLE-ES | 178 | 0 | 0 | 4 | 7 | Samples with <= 10 >= 5 bp deletions weren't analyzed |

*: For cohorts where no RNAseq data was available, we used data of the same tumor/tissue type.

#: For reference: the median size of deletions ≥ 5 bp in CCA_TH_19 is 8bp

Supplementary Table S3: The ID_TOP2A indel mutational signature

| Indel category Type | Indel category Subtype | Indel category Indel_size | Indel category Repeat_MH_size | Mean proportion of indels |
|---|---|---|---|---|
| DEL | C | 1 | 0 | 0.012463297 |
| DEL | C | 1 | 1 | 0.008977881 |
| DEL | C | 1 | 2 | 0.004135576 |
| DEL | C | 1 | 3 | 9.43E-05 |
| DEL | C | 1 | 4 | 7.07E-05 |
| DEL | C | 1 | 5+ | 0.00235192 |
| DEL | T | 1 | 0 | 0.024871668 |
| DEL | T | 1 | 1 | 0.013431147 |
| DEL | T | 1 | 2 | 0.005328606 |
| DEL | T | 1 | 3 | 0.002007652 |
| DEL | T | 1 | 4 | 0.000330087 |
| DEL | T | 1 | 5+ | 0.012340181 |
| INS | C | 1 | 0 | 0.005388998 |
| INS | C | 1 | 1 | 0.011689861 |
| INS | C | 1 | 2 | 0.013765039 |
| INS | C | 1 | 3 | 0.039728584 |
| INS | C | 1 | 4 | 0.004348358 |
| INS | C | 1 | 5+ | 0.009382251 |
| INS | T | 1 | 0 | 0.005977 |
| INS | T | 1 | 1 | 0.030892891 |
| INS | T | 1 | 2 | 0.025589339 |
| INS | T | 1 | 3 | 0.00407544 |
| INS | T | 1 | 4 | 0.001229006 |
| INS | T | 1 | 5+ | 0.010288818 |
| DEL | repeats | 2 | 0 | 0.025337649 |
| DEL | repeats | 2 | 1 | 0.013397023 |
| DEL | repeats | 2 | 2 | 0.00061302 |
| DEL | repeats | 2 | 3 | 0.000259354 |
| DEL | repeats | 2 | 4 | 4.72E-05 |
| DEL | repeats | 2 | 5+ | 0.005695835 |
| DEL | repeats | 3 | 0 | 0.020064376 |
| DEL | repeats | 3 | 1 | 0.008415445 |
| DEL | repeats | 3 | 2 | 0.000165044 |
| DEL | repeats | 3 | 3 | 2.36E-05 |
| DEL | repeats | 3 | 4 | 2.36E-05 |
| DEL | repeats | 3 | 5+ | 0 |
| DEL | repeats | 4 | 0 | 0.014268402 |
| DEL | repeats | 4 | 1 | 0.001060995 |
| DEL | repeats | 4 | 2 | 0.000117888 |
| DEL | repeats | 4 | 3 | 0 |
| DEL | repeats | 4 | 4 | 0 |
| DEL | repeats | 4 | 5+ | 4.72E-05 |
| DEL | repeats | 5+ | 0 | 0.058292306 |
| DEL | repeats | 5+ | 1 | 0.000400821 |
| DEL | repeats | 5+ | 2 | 0 |
| DEL | repeats | 5+ | 3 | 0 |
| DEL | repeats | 5+ | 4 | 0 |
| DEL | repeats | 5+ | 5+ | 0 |

| | | | | |
|---|---|---|---|---|
| INS | repeats | 2 | 0 | 0.021844021 |
| INS | repeats | 2 | 1 | 0.04879824 |
| INS | repeats | 2 | 2 | 0.030036737 |
| INS | repeats | 2 | 3 | 0.005648935 |
| INS | repeats | 2 | 4 | 0.000330087 |
| INS | repeats | 2 | 5+ | 0.000424398 |
| INS | repeats | 3 | 0 | 0.016244641 |
| INS | repeats | 3 | 1 | 0.060146355 |
| INS | repeats | 3 | 2 | 0.01480287 |
| INS | repeats | 3 | 3 | 0.000117888 |
| INS | repeats | 3 | 4 | 0 |
| INS | repeats | 3 | 5+ | 4.72E-05 |
| INS | repeats | 4 | 0 | 0.010433111 |
| INS | repeats | 4 | 1 | 0.132464465 |
| INS | repeats | 4 | 2 | 0.001913341 |
| INS | repeats | 4 | 3 | 2.36E-05 |
| INS | repeats | 4 | 4 | 4.72E-05 |
| INS | repeats | 4 | 5+ | 9.43E-05 |
| INS | repeats | 5+ | 0 | 0.004747412 |
| INS | repeats | 5+ | 1 | 0.006358615 |
| INS | repeats | 5+ | 2 | 2.36E-05 |
| INS | repeats | 5+ | 3 | 0 |
| INS | repeats | 5+ | 4 | 0 |
| INS | repeats | 5+ | 5+ | 2.36E-05 |
| DEL | MH | 2 | 1 | 0.016614098 |
| DEL | MH | 3 | 1 | 0.010074818 |
| DEL | MH | 3 | 2 | 0.010303221 |
| DEL | MH | 4 | 1 | 0.022355151 |
| DEL | MH | 4 | 2 | 0.010595344 |
| DEL | MH | 4 | 3 | 0.003944571 |
| DEL | MH | 5+ | 1 | 0.057719429 |
| DEL | MH | 5+ | 2 | 0.049373485 |
| DEL | MH | 5+ | 3 | 0.036807903 |
| DEL | MH | 5+ | 4 | 0.018588775 |
| DEL | MH | 5+ | 5+ | 0.012064464 |

| Standard error of the mean |
|---|
| 0.003603105 |
| 0.002988211 |
| 0.002159397 |
| 9.43E-05 |
| 7.07E-05 |
| 0.001485854 |
| 0.004485143 |
| 0.0051933 |
| 0.00236681 |
| 0.001529873 |
| 0.000330087 |
| 0.010263182 |
| 0.002130772 |
| 0.003288221 |
| 0.005535873 |
| 0.010812186 |
| 0.001877687 |
| 0.006651297 |
| 0.002412896 |
| 0.003959889 |
| 0.004411779 |
| 0.002294441 |
| 0.000941213 |
| 0.004036416 |
| 0.007793629 |
| 0.004911909 |
| 0.00061302 |
| 0.000259354 |
| 4.72E-05 |
| 0.004520649 |
| 0.004486309 |
| 0.00392341 |
| 0.000165044 |
| 2.36E-05 |
| 2.36E-05 |
| 0 |
| 0.003580444 |
| 0.001060995 |
| 0.000117888 |
| 0 |
| 0 |
| 4.72E-05 |
| 0.008558464 |
| 0.000400821 |
| 0 |
| 0 |
| 0 |
| 0 |

| |
|---|
| 0.002945519 |
| 0.003767752 |
| 0.007748055 |
| 0.002258035 |
| 0.000330087 |
| 0.000424398 |
| 0.005019985 |
| 0.010257671 |
| 0.006041408 |
| 0.000117888 |
| 0 |
| 4.72E-05 |
| 0.004221822 |
| 0.012573779 |
| 0.001519457 |
| 2.36E-05 |
| 4.72E-05 |
| 9.43E-05 |
| 0.002439243 |
| 0.002728227 |
| 2.36E-05 |
| 0 |
| 0 |
| 2.36E-05 |
| 0.003557306 |
| 0.003133076 |
| 0.002989503 |
| 0.006160346 |
| 0.002888141 |
| 0.003050584 |
| 0.006256792 |
| 0.009036998 |
| 0.007069812 |
| 0.002670225 |
| 0.007265672 |

Supplementary Table S4: ID_TOP2A mutations in cancer gene census genes

| Chr | Start | End | Ref | Alt | Ref.Genom | Func.refGene | Gene.refGene | GeneDetai | ExonicFun | AAChange | sample | CGC_tier |
|---|---|---|---|---|---|---|---|---|---|---|---|---|
| 1 | 201981535 | 201981535 | - | C | GRCh37 | exonic | ELF3 | . | frameshift | ELF3:NM_( | TCGA-CG-4477 | 2 |
| 1 | 202015216 | 202015217 | TA | - | GRCh38 | exonic | ELF3 | . | frameshift | ELF3:NM_( | CCA_TH_19 | 2 |
| 2 | 60687597 | 60687609 | TTCTCCAGGGTAC | - | GRCh37 | exonic | BCL11A | . | frameshift | BCL11A:NI | TCGA-VQ-A8DL | 1 |
| 2 | 70315933 | 70315933 | - | G | GRCh37 | exonic | PCBP1 | . | frameshift | PCBP1:NM | TCGA-KB-A93J | 2 |
| 3 | 52407999 | 52408002 | GGGT | - | GRCh38 | exonic | BAP1 | . | stopgain | BAP1:NM_ | CCA_TH_19 | 1 |
| 3 | 53836195 | 53836209 | CATTAGGTATGTGCT | - | GRCh37 | exonic | CACNA1D | . | stopgain | CACNA1D: | TCGA-CG-5725 | 1 |
| 3 | 121207265 | 121207265 | - | TAAT | GRCh37 | exonic | POLQ | . | frameshift | POLQ:NM_ | TCGA-CG-4477 | 1 |
| 3 | 128622178 | 128622178 | - | AGAG | GRCh38 | exonic | RPN1 | . | stopgain | RPN1:NM_ | 04-112 | 1 |
| 5 | 142258963 | 142258963 | - | TCAC | GRCh37 | exonic | ARHGAP26 | . | frameshift | ARHGAP2( | TCGA-KB-A93J | 1 |
| 5 | 149459703 | 149459707 | CTTGG | - | GRCh37 | exonic | CSF1R | . | frameshift | CSF1R:NM | TCGA-KB-A93J | 2 |
| 6 | 168290003 | 168290020 | AAAGGTAGTAACCTTATT | - | GRCh37 | exonic | AFDN | . | frameshift | AFDN:NM_ | TCGA-KB-A93J | 1 |
| 7 | 140778025 | 140778039 | ACTGCTGAGGTGTAG | - | GRCh38 | exonic | BRAF | . | nonframes | BRAF:NM_ | 04-112 | 1 |
| 8 | 42321912 | 42321924 | GAGCTGTACAGGA | - | GRCh38 | exonic | IKBKB | . | frameshift | IKBKB:NM_ | 04-112 | 1 |
| 9 | 96030931 | 96030931 | - | T | GRCh37 | exonic | WNK2 | . | frameshift | WNK2:NM | TCGA-CG-4477 | 2 |
| 9 | 110248111 | 110248111 | - | GGT | GRCh37 | exonic | KLF4 | . | nonframes | KLF4:NM_( | TCGA-CG-4477 | 1 |
| 9 | 121144961 | 121144961 | - | CAG | GRCh38 | splicing | CNTRL | NM_0013( | . | . | T10 | 1 |
| 9 | 136901394 | 136901424 | CTTCCTCTGAGTCGTAGGAGGCAGATGCCTG | - | GRCh37 | exonic | BRD3 | . | frameshift | BRD3:NM_ | TCGA-CG-5725 | 1 |
| 10 | 843022 | 843023 | CA | - | GRCh38 | exonic | LARP4B | . | frameshift | LARP4B:NI | T10 | 2 |
| 10 | 59906342 | 59906342 | - | TGCT | GRCh38 | exonic | CCDC6 | . | stopgain | CCDC6:NM | CCA_TH_19 | 1 |
| 10 | 87952122 | 87952122 | - | A | GRCh38 | exonic | PTEN | . | frameshift | PTEN:NM_ | 04-112 | 1 |
| 10 | 87952124 | 87952126 | ACT | - | GRCh38 | exonic | PTEN | . | nonframes | PTEN:NM_ | 04-112 | 1 |
| 10 | 89711879 | 89711879 | - | A | GRCh37 | exonic | PTEN | . | frameshift | PTEN:NM_ | TCGA-CG-5725 | 1 |
| 10 | 89720779 | 89720779 | - | C | GRCh37 | exonic | PTEN | . | frameshift | PTEN:NM_ | TCGA-VQ-A8DL | 1 |
| 12 | 22671278 | 22671283 | TACAGT | - | GRCh38 | exonic | ETNK1 | . | nonframes | ETNK1:NM | T10 | 1 |
| 12 | 49042874 | 49042874 | - | AGTT | GRCh38 | exonic | KMT2D | . | frameshift | KMT2D:NI | T10 | 1 |
| 12 | 56495820 | 56495820 | - | TAA | GRCh37 | exonic | ERBB3 | . | nonframes | ERBB3:NM | TCGA-KB-A93J | 1 |
| 12 | 56720159 | 56720164 | AAAGGT | - | GRCh38 | exonic | NACA | . | nonframes | NACA:NM_ | CCA_TH_19 | 2 |
| 12 | 121805842 | 121805845 | AGTT | - | GRCh38 | exonic | SETD1B | . | frameshift | SETD1B:NI | 04-112 | 2 |
| 13 | 35628223 | 35628248 | GTATCACTGATCCTGTGCTCAGGGAG | - | GRCh38 | exonic | NBEA | . | frameshift | NBEA:NM_ | CCA_TH_19 | 2 |
| 14 | 51202240 | 51202240 | - | AGCT | GRCh37 | exonic | NIN | . | frameshift | NIN:NM_0 | TCGA-CG-4477 | 1 |
| 15 | 57192232 | 57192232 | - | TACTATTC | GRCh38 | exonic | TCF12 | . | frameshift | TCF12:NM | CCA_TH_19 | 1 |
| 15 | 57251347 | 57251353 | CAGGTAC | - | GRCh38 | exonic | TCF12 | . | frameshift | TCF12:NM | T10 | 1 |
| 17 | 7577565 | 7577566 | TT | - | GRCh37 | exonic | TP53 | . | frameshift | TP53:NM_ | TCGA-VQ-A8DL | 1 |
| 17 | 7578183 | 7578187 | CGGCT | - | GRCh37 | exonic | TP53 | . | frameshift | TP53:NM_ | TCGA-CG-4477 | 1 |
| 17 | 7579528 | 7579528 | - | CATT | GRCh37 | exonic | TP53 | . | stopgain | TP53:NM_ | TCGA-KB-A93J | 1 |
| 17 | 7674246 | 7674246 | - | T | GRCh38 | exonic | TP53 | . | frameshift | TP53:NM_ | CCA_TH_19 | 1 |
| 17 | 20000065 | 20000065 | - | T | GRCh37 | exonic | SPECC1 | . | frameshift | SPECC1:NI | TCGA-CG-4477 | 2 |
| 17 | 57743978 | 57743978 | G | - | GRCh37 | exonic | CLTC | . | frameshift | CLTC:NM_ | TCGA-CG-4477 | 1 |
| 18 | 42449206 | 42449206 | - | AGTG | GRCh37 | exonic | SETBP1 | . | frameshift | SETBP1:NI | TCGA-VQ-A8DL | 1 |
| 19 | 6829798 | 6829798 | - | ATGC | GRCh37 | exonic | VAV1 | . | frameshift | VAV1:NM_ | TCGA-CG-5725 | 2 |
| 20 | 32435123 | 32435139 | GACCCACCGTTCCTGCA | - | GRCh38 | exonic | ASXL1 | . | frameshift | ASXL1:NM | T10 | 1 |
| 20 | 32435124 | 32435124 | - | C | GRCh38 | exonic | ASXL1 | . | frameshift | ASXL1:NM | 04-112 | 1 |
| 20 | 62161438 | 62161439 | GG | - | GRCh37 | exonic | PTK6 | . | frameshift | PTK6:NM_ | TCGA-CG-5725 | 1 |
| X | 44949103 | 44949106 | GCAG | - | GRCh37 | exonic | KDM6A | . | frameshift | KDM6A:NI | TCGA-KB-A93J | 1 |
| X | 66766578 | 66766578 | - | T | GRCh37 | exonic | AR | . | frameshift | AR:NM_0C | TCGA-CG-4477 | 1 |
